# Uncovering the hidden RNA virus diversity in Lake Nam Co: evolutionary insights from an extreme high-altitude environment

**DOI:** 10.1101/2024.07.04.601995

**Authors:** Lilin Wu, Yongqin Liu, Wenqing Shi, Tianyi Chang, Pengfei Liu, Keshao Liu, Yong He, Zhaorong Li, Mang Shi, Nianzhi Jiao, Andrew S. Lang, Xiyang Dong, Qiang Zheng

## Abstract

Alpine lakes, characterized by isolation, low temperatures, oligotrophic conditions, and intense ultraviolet radiation, remain a poorly explored ecosystem for RNA viruses. Here, we present the first comprehensive metatranscriptomic study of RNA viruses in Lake Nam Co, a high-altitude alkaline saline lake on the Tibetan Plateau. Using a combination of sequence– and structure-based homology searches, we identified 742 RNA virus species, including 383 novel genus-level groups and 84 novel family-level groups exclusively found in Lake Nam Co. These findings significantly expand the known diversity of the *Orthornavirae*, uncovering evolutionary adaptations such as permutated RNA-dependent RNA polymerase (RdRP) motifs and distinct RNA secondary structures. Notably, 14 previously unknown RNA virus families potentially infecting prokaryotes were predicted, broadening the known host range of RNA viruses and questioning the traditional assumption that RNA viruses predominantly target eukaryotes. The presence of auxiliary metabolic genes (AMGs) in viral genomes suggested that RNA viruses (families *f*.*0102* and Nam-Co_family_51) exploit host energy production mechanisms in energy-limited alpine lakes. Low nucleotide diversity, SNP frequencies, and pN/pS ratios indicate strong purifying selection in Nam Co viral populations. Our findings offer insights into RNA virus evolution and ecology, highlighting the importance of extreme environments in uncovering hidden viral diversity, and further shed light into their potential ecological implications, particularly in the context of climate change.

**Significance Statement:** This study unveils the hidden diversity and complexity of RNA viruses in the high-altitude ecosystem of Lake Nam Co, significantly expanding the known RNA virome. The discovery of novel viral clades, expanded potential host ranges, and unique evolutionary adaptations emphasizes the importance of exploring extreme environments to uncover viral diversity and evolutionary pathways. Our findings offer critical insights into viral evolution, host-virus interactions, and the ecological roles of RNA viruses, particularly in the context of climate change and environmental resilience.

## Introduction

RNA viruses are among the most abundant and diverse biological entities, yet our understanding of their global diversity and evolutionary dynamics remains limited(1). Since the first aquatic RNA virus was isolated in 1966, significant advancements have been made in the study of the RNA virome(2). Early efforts primarily focused on isolating virus particles, while recent studies have utilized high-throughput sequencing data to uncover new viruses more efficiently(3). The methodologies for discovering RNA viruses have evolved from traditional sequence similarity searches to advanced techniques using deep learning frameworks(4–8). Over the past two decades, comprehensive omics surveys have led to exponential growth in identifying RNA viruses, particularly through the identification of RNA-dependent RNA polymerases (RdRPs) encoding sequences in specific organisms(9, 10) and environmental samples(11–15). For instance, an analysis of over 5000 metatranscriptomes has identified vast clades of RNA bacteriophages(16). Additionally, a deep learning algorithm named LucaProt has proposed a hidden realm of viral diversity(8). Although these groups have not been officially recognized, their significance in RNA virus retrieval and identification is undeniable. Despite these advancements, RNA viruses in extreme and isolated ecosystems, such as high-altitude lakes, remain underexplored(3, 8, 17), which likely accounts for the relatively small fraction of RNA viral diversity discovered so far(11). Moreover, areas beyond RNA virus classification and genomics, such as the adaptive and structural characteristics of RNA viruses in environmental contexts, have been largely overlooked and remain rarely explored(18, 19).

Freshwater ecosystems, particularly those in cold regions, have been shown to harbor a higher diversity of RNA viruses compared to other environments, likely due to their favorable conditions for viral persistence(8, 20, 21). Among these, alpine lakes, characterized by extreme conditions such as low temperatures, high ultraviolet radiation, and oligotrophic waters, serve as reservoirs of previously unknown microbial and viral diversity. These communities often show specific adaptation strategies to their harsh environments(22–24). However, no comprehensive study of the RNA virome in such environments has been conducted. Current knowledge of RNA viruses has largely been derived from studies in more accessible habitats, such as terrestrial(13), estuarine(12), and oceanic ecosystems(14). The Tibetan Plateau, often referred to as the “Third Pole” due to its extensive coverage of glaciers and high-altitude lakes beyond the polar regions(25), boasts over 1,400 lakes larger than 1.0 km^2^ (26), representing a largely untapped reservoir of viral diversity(22). Lake Nam Co, a large enclosed lake primarily fed by glacial meltwater(27, 28), situated at an elevation of 4,718 meters, spans a surface area of around 2,000 square kilometers, making it the second-largest lake on the Tibet Plateau(29). Its isolation, extreme environmental conditions, and the unique microbial communities make it an ideal candidate for uncovering novel RNA viruses(21, 30, 31).

In this study, we conducted a comprehensive metatranscriptomic analysis of Lake Nam Co, employing both traditional sequence similarity searches—BLAST and HMM—and advanced deep learning method LucaProt to uncover RNA viruses. Our findings have revealed a significant array of novel and endemic RNA viruses, substantially broadening our understanding of RNA viral diversity in this large enclosed lake. We also identified protein and genome characteristics that contribute to viral adaptation strategies to extreme habitats. This work provides new insights into the viral ecology of extreme aquatic environments, highlighting the importance of studying isolated ecosystems to understand the pressures shaping RNA viruses. By characterizing the features of RNA viruses in Lake Nam Co, we aim to advance knowledge of viral evolution, host-virus interactions, and the ecological roles of viruses in high-altitude lakes.

## Results

### Diverse cellular life in Lake Nam Co’s cold, alkaline, and oligotrophic waters

Environmental analysis of the 5 water samples from four distinct depths (surface, upper-middle, lower-middle, and bottom) in Lake Nam Co revealed its high-altitude, cold, alkaline, moderately saline, and oligotrophic conditions (**Figure 1 and Tables S1, S2**). The lake’s significant exposure to intense sunlight is marked by a peak photosynthetically active radiation of 1,379 W m^−2^. Temperatures at the surface drop significantly with depth, from 12.9□ to 3.6□ over an 80 m range, even during summer, while the pH levels remained stable between 9.3 and 9.4 at the LC station. Salinity showed minor fluctuations, ranging from 1.05 to 1.08‰. Dissolved oxygen levels fluctuated with depth, from 11.1 to 13.8 mg L^−1^. Dissolved organic carbon concentrations were narrowly constrained between 3.39 to 3.50 mg L^−1^. Nutrient concentrations, including nitrate, phosphates, and ammonium, were below 1 μM, and generally increased from the surface to the bottom, with the lower-middle layer showing the highest concentrations (**Table S2**).

**Figure 1.**
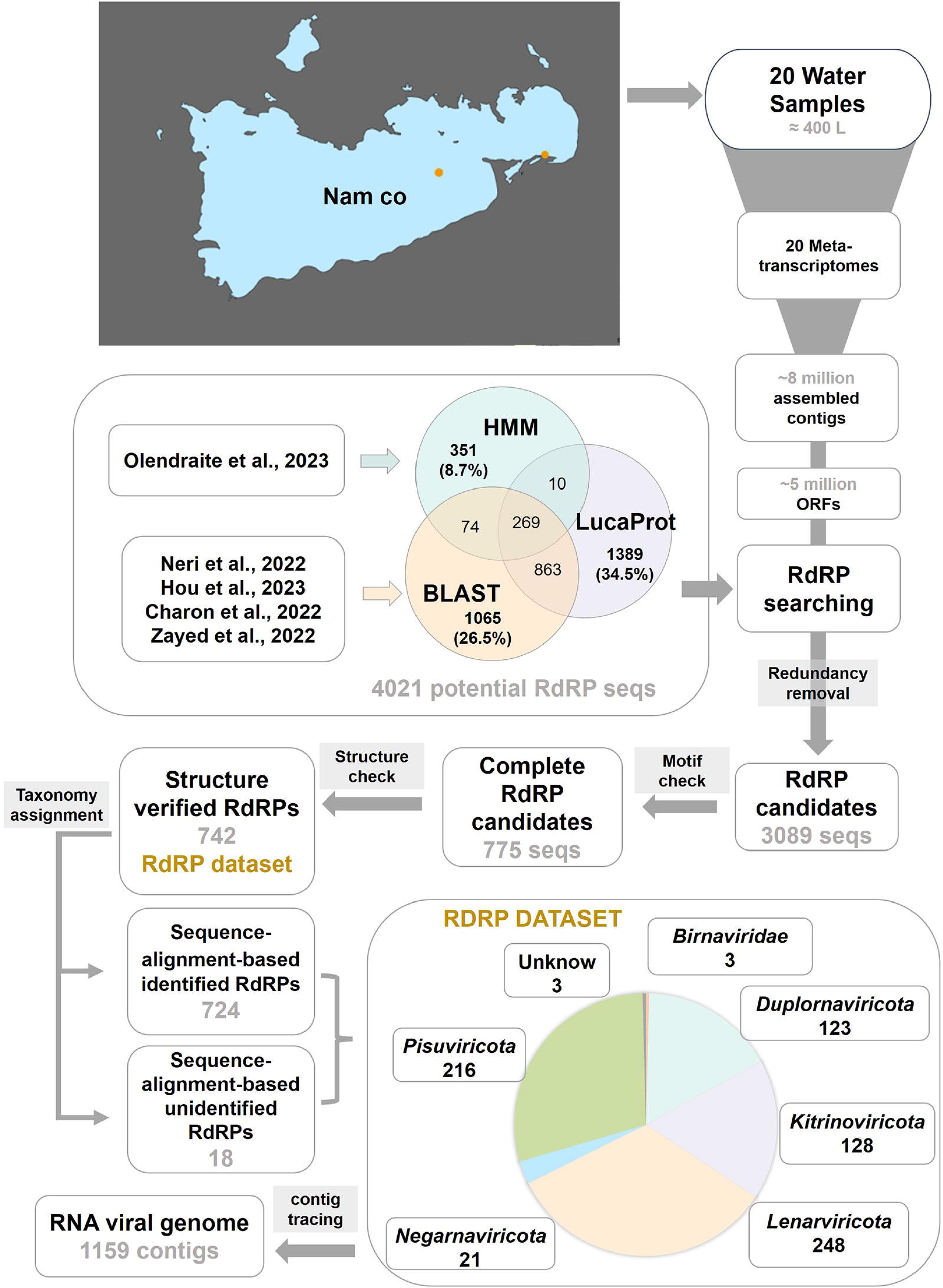
Sampling locations and RNA virus detection process. The diagram outlines the workflow from data collection and processing to the construction of the RdRP dataset for Lake Nam Co. The methods used for retrieving RdRP sequences include BLASTp, HMM searches, and the deep learning algorithm LucaProt. In total, 742 non-redundant complete RdRP sequences were identified from 20 metatranscriptomes. Of these, 724 RdRPs were identified through both protein structure-based clustering and sequence-alignment-based analyses, while the remaining 18 were detected exclusively via structure-based clustering. The RdRP dataset consists of RNA viruses from the five orthornaviran phyla and the *Birnaviridae* family, which is not assigned to the five major phyla. The pie chart displays the distribution of RNA virus species across the five orthornaviran phyla and *Birnaviridae*, based on combined sequence-alignment and protein structure clustering analyses.

Sequencing of 16S rRNA and 18S rRNA transcripts in Lake Nam Co revealed an active microbial landscape with high bacterial and eukaryotic richness (**Figures S1–S4**). At the class level, *Cyanobacteriia* dominantly comprised 28.0% to 68.7% of the bacterial community. Other notable contributors to the bacterial diversity include *Gammaproteobacteria*, *Bacteroidia*, *Alphaproteobacteria*, *Planctomycetes*, *Verrucomicrobiales*, *Actinobacteria*, *Bdellovibrionota*, *Desulfuromonadia*, and *Acidimicrobiia*. For the eukaryotic community, diversity spans several groups with *Spirotrichea* (ranging from 2.8% to 59.5%) and *Dinophyceae* (17.5% to 55.5%) among the most prominent. Other key eukaryotic classes include *Bacillariophyta*, *Cryptophyceae*, *Arthropoda*, *Heterotrichea*, *Choanoflagellatea*, *MAST-2*, *Trebouxiophyceae*, *CONThreeP*, *Chrysophyceae*, *Rotifera*, *Zygnemophyceae*, *Litostomatea*, *Chytridiomycota*, and *Filosa-Sarcomonadea*.

### Discovery of 14 novel RNA viral clades

Using BLAST and HMM, along with the deep learning algorithm LucaProt, followed by a series of validation processes (**Figure 1**), we identified 742 non-redundant complete RdRPs containing three motifs forming the catalytic core from 20 metatranscriptomes of Lake Nam Co. Among these, 724 RdRPs, which were subsequently used for phylogenetic analysis, were identified through both sequence-alignment-based and protein structure-based clustering analyses (**Figure 2A**), with the remaining 18 detected solely via structure-based clustering (**Figure 2B**). For the former 724 RdRPs, 334 RdRP sequences were assigned to taxa currently defined by ICTV (International Committee on Taxonomy of Viruses), while another 390 belonged to taxa identified in the RVMT database but not officially classified at this time (**Figures S5-S20 and Table S3**). For those RdRPs belonging to the proposed classifications in the RVMT database derived from the work by Neri et al., we have retained the classification names(16). Of the 18 RdRPs identified exclusively by structure-based clustering, 15 were identified to current phyla within *Orthornavirae* without classification into more refined taxonomic levels, while the remaining three could not be assigned to any existing *Orthornavirae* taxonomy (**Figures S21-S31**). From the total RdRP dataset, 602 newly identified RdRPs formed 383 novel genus-level groups at a 75% amino acid identity (AAI), and 118 RdRPs formed 84 novel family-level groups at a 40% AAI.

**Figure 2.**
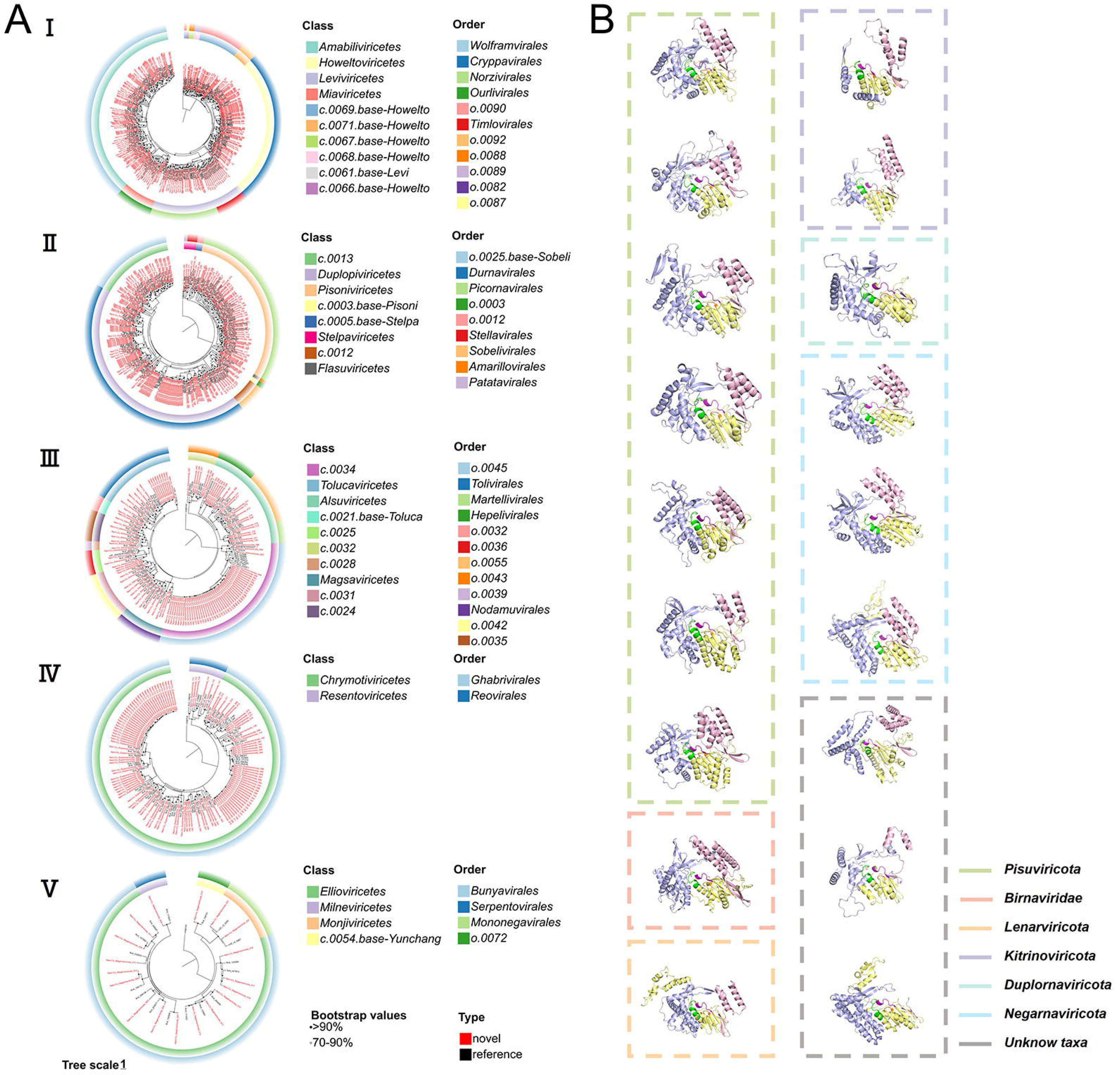
Classification and structural information of RdRP dataset in Lake Nam Co. (A) Phylogenetic trees of 724 sequence-alignment-based identified RdRPs. From □ to □, they represent phyla *Lenarviricota*, *Pisuviricota*, *Kitrinoviricota*, *Duplornaviricota*, and *Negarnaviricota*, respectively. RdRPs found in Lake Nam Co are highlighted in red, while the reference sequences are shown in black. (B) Schematic representation of 18 sequence-alignment-based unidentified RdRPs that detected solely via protein structure-based clustering. The “thumb”, “palm”, and “fingers” regions of RdRP are colored in pink, yellow, and blue, respectively. Within the “palm” region, catalytic center motifs A, B, and C are colored in purple, green, and orange, respectively.

Based on sequence and structural similarities (**Table S3 and Figure 1**), except for the three previously mentioned unclassified RdRPs, the RdRP dataset comprised viruses from the five established phyla: 248 from *Lenarviricota*, 216 from *Pisuviricota*, 128 from *Kitrinoviricota*, 123 from *Duplornaviricota*, and 21 from *Negarnaviricota*. Additionally, the dataset includes three *Birnaviridae* viruses that could not be assigned to any of the 5 major phyla due to the permutation of three essential motifs forming the RdRP catalytic core (**Figure 1**).

Expansions were observed in various phyla through the phylogenetic analysis, and 14 novel and Lake Nam Co-specific phylogenetic clades (with ≥ 3 representative sequences) at the sub-family or family level were identified. These clades shared less than 50% AAI with RdRP reference sequences from other environments(8, 14, 16, 32) (**Table S4 and Figures S5-S20**). In *Lenarviricota*, although the majority of newly identified RdRP sequences were found (**Figure 2A** D), only two novel clades (*Lenarviricota*_Clades_I-II) were discovered, corresponding to the families *Narnaviridae* and proposed family *f.0388* from the RVMT database(16) (**Table S4 and Figures S5, S8**). For *Pisuviricota*, six newly identified clades were observed: one in *Picornavirales* (*Pisuviricota*_Clade_I) within *Pisoniviricetes* and five in *Durnavirales* (*Pisuviricota*_Clades_II-VI) within *Duplopiviricetes* (**Figure 2A** D) (**Table S4 and Figures S9, S12, S13**). In *Kitrinoviricota*, the primary expansion was observed in the proposed order *o.0045* of class *c.0034* from the RVMT database(16) (**Figure 2A** D), where 45 sequences formed an independent clade (*Kitrinoviricota*_Clade_I) (**Figure S14**). These sequences diverged from the closest reference sequences in family *f.0176* identified in the RVMT database(16) with similarities ranging from 29.4% to 36.3% (**Table S4**). Among them, three clades (*Duplornaviricota*_Clades_III, IV and V) corresponded to families *f.0295*, *f.0286* and *f.0287* in the RVMT database(16), exhibiting low AAI (<40%) with other sequences and were distantly clustered (**Figures S18-S20**). In *Negarnaviricota*, the fewest newly identified RdRP sequences were observed, predominantly belonging to *Bunyavirales* within *Ellioviricetes* of *Polyploviricotina* (**Figure 2A** D), with no novel clades discovered.

### Novel RdRPs with unique structural characteristics in Lake Nam Co

The polymerase domain of RdRPs consists of three subdomains: palm, fingers, and thumb(33). To determine the taxonomy of sequence-alignment-based unclassified RdRPs, which could not be recognized through sequence alignment, we predicted the structures of the polymerase domain of all Lake Nam Co RdRPs and their closely related reference sequences, and conducted structural clustering (**Figures 3 and 4**).

**Figure 3.**
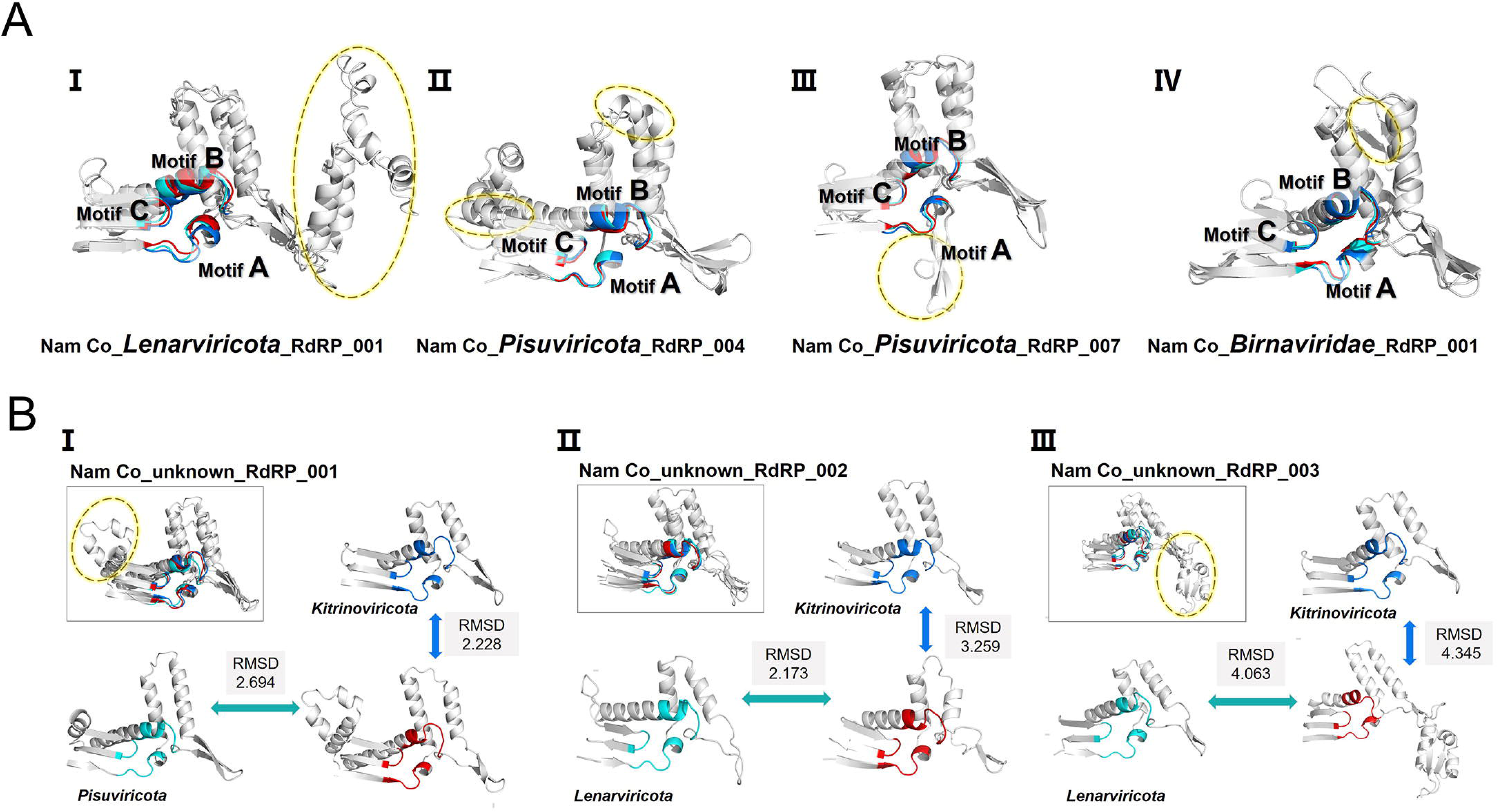
Schematic representation of catalytic core in the sequence-alignment-based unidentified RdRP structures. (A) Unique structures of the “palm” region of sequence-alignment-based unidentified RdRPs that could be identified at the phyla level based on protein structural clustering. (B) Structures of the “palm” region of RdRPs that remain unidentified at the phyla level through either sequence-alignment-based or protein structural-based clustering analyses. Motifs from sequence-alignment-based unidentified RdRPs are colored in red, motifs from the RVMT reference structure are colored in dark blue, and motifs representing sequence-alignment-based identified RdRPs in Lake Nam Co are colored in light blue. The yellow dashed circle indicates unique structural regions of sequence-alignment-based unidentified RdRPs from Lake Nam Co.

**Figure 4.**
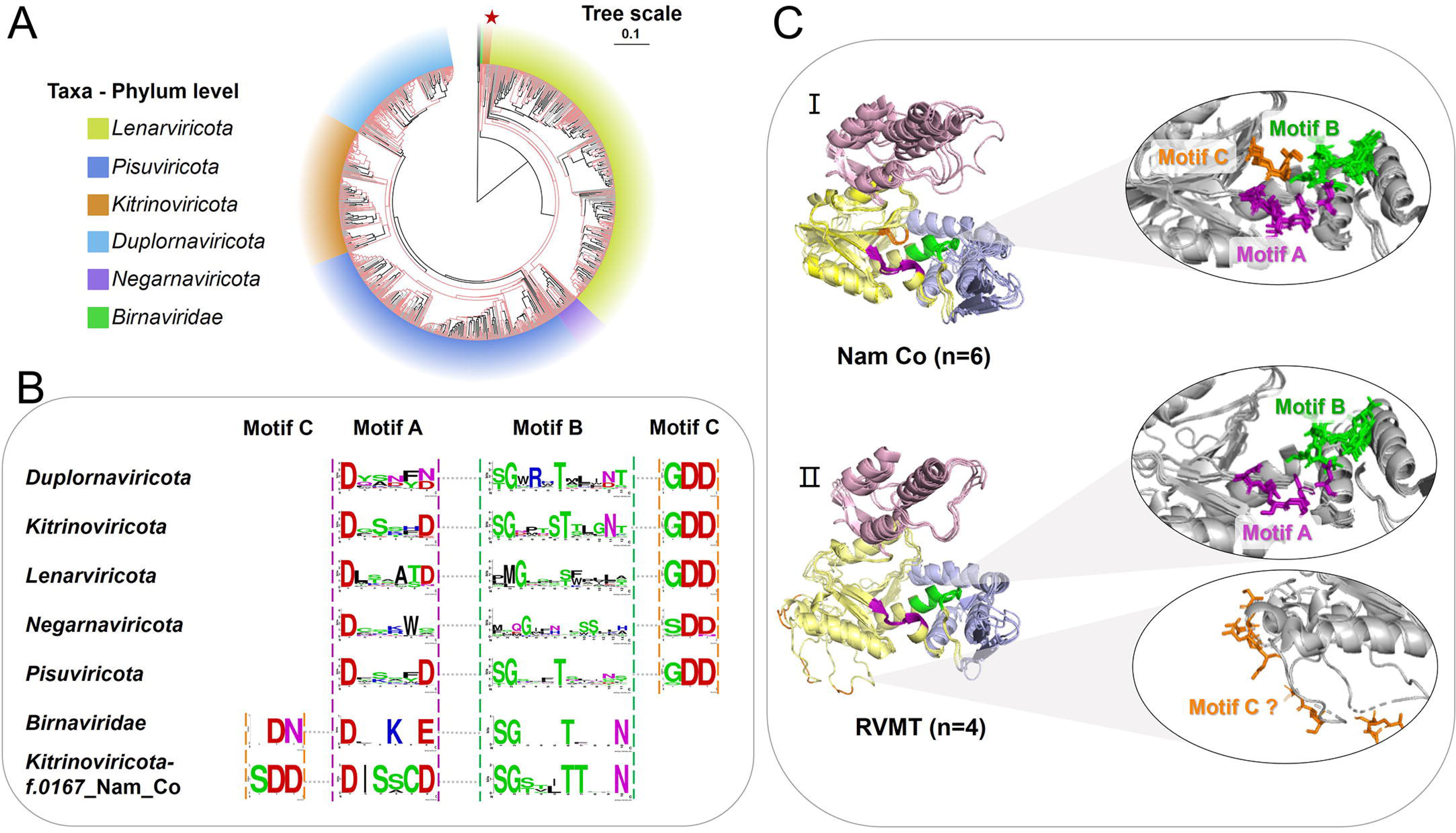
RdRP structure-based clustering, motif arrangements and structural features. (A) Structure-based clustering results of the RdRP dataset in Lake Nam Co and references. The position of *f.0167*_Nam_Co is marked with a red star. (B) Composition and arrangement of RdRP motifs at the phylum level in Lake Nam Co, with the motifs of *f.0167*_Nam_Co from *Kitrinoviricota* exhibiting a permuted arrangement shown at the bottom. (C) Schematic representation of the RdRP structure of *f.0167* from *Kitrinoviricota*, with overlaid structures from Lake Nam Co (□) and the RVMT database (□), annotated with the number of overlaid sequences. The “thumb,” “palm,” and “fingers” regions of RdRP are colored in pink, yellow, and blue, respectively. Within the “palm” region, catalytic center motifs A, B, and C are colored in purple, green, and orange, respectively. Motifs in the enlarged details are displayed in stick mode.

Through structural clustering and root mean squared deviation values (RMSD) comparison, we assigned the phyla of 15 sequence-alignment-based unidentified RdRPs, seven of which exhibited unique structural features in their core regions (**Figures 3A and S21-S28**). For instance, Nam-Co_*Birnaviridae*_001 had a special β-sheet structure between two typical α-helices in the non-conserved region between motifs A and B (**Figures 3A** D**, S21**). Nam-Co_*Lenarviricota*_001 featured multiple α-helices and irregular coils with distinctive amino acids in motifs A and B (**Figures 3A** D**, S22**). Nam-Co_*Pisuviricota*_004 and Nam-Co_*Pisuviricota*_007 exhibited extra helix and distinctive β-sheet structures, respectively (**Figures 3A** D, D**, S24B, S25B**). Nam-Co_*Kitrinoviricota*_001 showed a shorter first α-helix between motifs A and B, with an extra α-helix at the β-turn between motifs B and C (**Figure S26A**). These unique features highlight the diversity and endemism of RdRPs in Lake Nam Co.

Three sequence-alignment-based unidentified RdRPs remain unclassified based on structural clustering. Their RMSD values closely align with reference structures from different phyla, indicating their unique structural features. For Taxa_unknown_RdRP_001, the RMSD values of its structure compared to relatively close reference structures belonging to *Pisuviricota* and *Kitrinoviricota* both exceed 2 (**Figures 3B** D **and S29**). It shared β-turn features with *Kitrinoviricota* reference and similar α-helical structures with the *Pisuviricota* reference, though its helical structure was longer and divided into three segments. Taxa_unknown_RdRP_002 referenced *Lenarviricota* and *Kitrinoviricota*, with RMSD values of 2.173 and 3.259, respectively. It shared more similarities with *Kitrinoviricota* reference in the composition of motifs A and B, along with the β-turns at the edges of motifs A and C (**Figures 3B** D **and S30**). Taxa_unknown_RdRP_003 was associated with *Lenarviricota* and *Kitrinoviricota* references, with both RMSD values greater than 4, suggesting a high degree of specificity. It presented a highly unique area comprising two α-helices and two β-folds between motifs A and B (**Figures 3B** D **and S31**). These three RdRPs, identified exclusively by structure-based clustering with remote sequence and structural similarity, suggest the potential emergence of new phyla.

Besides the confirmation of phyla of sequence-alignment-based unidentified RdRPs, structural clustering unveiled a unique RNA virus group in Lake Nam Co. Six RdRP sequences, initially identified as *Kitrinoviricota* based on sequence alignments from Lake Nam Co (referred to as *f.0167*_Nam_Co), did not cluster with other *Kitrinoviricota* sequences. Instead, they formed a distinct cluster situated between *Birnaviridae* and the five orthornaviran phyla according to protein structural clustering analysis (**Figure 4A**). The RdRPs of *Birnaviridae* or related viruses are permutated, meaning that their motifs might shift places (**Figure 4B**). The low RMSD values (RMSD = 2.16) between *f.0167*_Nam_Co and *Birnaviridae* references, along with the similar rearranged motif sequences (motifs C-A-B) suggest a similarity between *f.0167*_Nam Co and *Birnaviridae* (**Figures 4B and 4C** D). Reference sequences of *f.0167*_Nam_Co, previously classified in the family *f.0167* within the class *c.0032* of *Kitrinoviricota* identified from the RVMT database(16), were thought to contain the SDD sequence as motif C with the general arrangement. However, structural predictions revealed that these sequences did not possess motif C in the palm region of the polymerase domain (**Figure 4C** D). The distinct arrangement of motifs in *f.0167* between Lake Nam Co and other aquatic environments(16, 34) reveals the evolutionary process of motif C positions, reflecting reconstitution and loss in the conserved core region.

### Genome and protein features of RNA viruses adapting to alpine environments

A total of 1,159 RNA viral contigs (**Figure 1**), representing viral genomes, were identified based on the presence of genes encoding RdRPs. Among these, 166 contigs contained more than three open reading frames (ORFs), with genome lengths ranging from 1,371 to 16,715 bases. Besides genes encoding RdRP and capsid proteins, the majority of these RNA virus genomes contained predominantly unannotated genes (**Figure S32**). Other annotated viral proteins encoded by RNA viruses in Lake Nam Co included movement proteins (MP), glycoproteins, helper-component proteinases (HCP), replicases, methyltransferases (MT), packaging NTPase P4, and peptidases S39 and A21 (**Figures 5A, S33A**).

**Figure 5.**
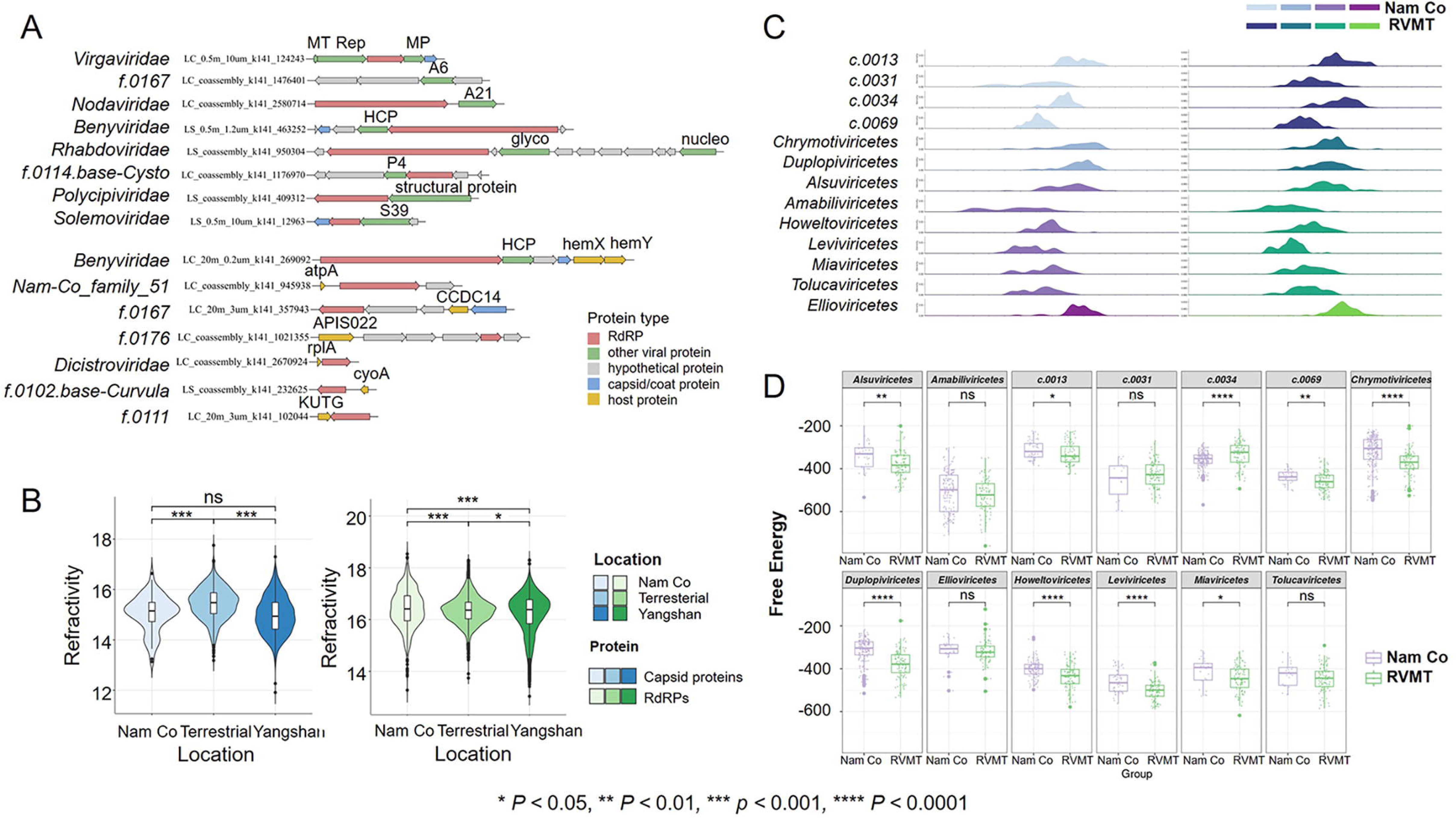
Features of genomes and proteins of RNA viruses in Lake Nam Co. (A) The RNA virus genomes containing viral and host genes in Lake Nam Co. Genes encoding RdRPs are colored in red, genes encoding capsid or coat proteins are colored in blue, other viral genes were colored in green, host genes were colored in yellow. (B) Comparison of refractivity indices of RNA virus proteins from Lake Nam Co with those from terrestrial and estuarine environments. RdRPs and capsid proteins of RNA viruses from different locations are shown in gradient blue and green, respectively. The capsid proteins of Lake Nam Co RNA viruses and those from estuarine environments show no significant difference in refractivity, but they do differ significantly from terrestrial environments. The RdRPs of Lake Nam Co RNA viruses show significant refractivity differences from both estuarine and terrestrial environments. (C) The distribution of normalized minimum free energy (MFE) of RNA secondary structures with respect to viral classes. MFE distributions of RNA viruses from different classes in Lake Nam Co (colored in gradient purple) and from the RVMT database (colored in gradient blue-green) show consistent trends across classes. (D) Comparison of normalized MFE of RNA secondary structures between Lake Nam Co and the RVMT database at the level of viral classes. MFE of RNA viruses from the same class in Lake Nam Co (colored in purple) and the RVMT database (colored in green) show significant differences between nine classes.

Six of viral genomes contained a total of seven auxiliary metabolic genes (AMGs), impacting various metabolic processes in hosts, including energy generation (*atpA*, *cyoA*), translation (*rplA*), phospholipid metabolism (*ugpQ*), and heme biosynthesis (*hemX*, *hemY*) (**Figure 5A and Table S9**). Among them, a virus belonging to the family *f.0102* within *Pisuviricota* identified in the RVMT database(16) carries a gene encoding cytochrome c oxidase subunit II in *Achlya* (**Figure 5A, Table S9**). An RNA virus identified as Nam-Co_family_51 within *Kitrinoviricota* carries a gene encoding ATP synthase CF1 alpha subunit of an unknown host (**Figure 5A, Table S9**). These two AMGs encode proteins that have been confirmed to function in the host’s mitochondria or chloroplasts, can affect the energy metabolism processes, which reflects a viral strategy to exploit host energy production. Genes related to other metabolism were also identified, such as AMGs encoding uroporphyrinogen-□ C-methyltransferase and heme biosynthesis-associated TPR protein in *Hydrogenobacter*, are possibly associated with multiple processes like electron transfer and oxygen transport(35). Besides AMGs, two genes associated with anti-prokaryotic immune systems, like APIS022 and AcrVIA7, suggest an ongoing arms race between viruses and their hosts(36) (**Figure 5A and Table S9**).

To illustrate the RNA virus protein characteristics in Lake Nam Co, we compared the physicochemical properties of capsid proteins and RdRPs from RNA viruses in Lake Nam Co, various terrestrial sites (16 provinces and regions in China)(13), and estuarine environments (Yangshan Port)(12), including hydrophobicity, polarity, mutability, transmembrane tendency, and refractivity (**Figure 5B and S34**). Capsid proteins from the two aquatic environments showed no obvious difference (wilcox test, *p* > 0.05) but were significantly different from terrestrial capsid proteins (wilcox test, *p* < 0.05). Comparing to capsid proteins, Lake Nam Co RdRPs significantly differed from both estuarine and terrestrial environments across all five indices (wilcox test, *p* < 0.001) (**Figure 5B and S34**).

Besides characteristics of given proteins, the genome features of RNA viruses in Lake Nam Co were also shaped by its environment and hosts. We compared the RNA secondary structural features of RNA viruses from Lake Nam Co with RVMT references within the same classes, analyzing hairpins, stems, interior loops, multiloops, unpaired ratios, and minimum free energy (MFE)(37) (**Figures 5C, 5D and S35**). We focused on taxonomic groups at the class level that had > 10 relatively complete RNA genomes, averaging 90 sequences per class, including the *Alsuviricetes*, *Amabiliviricetes*, *Chrymotiviricetes*, *Duplopiviricetes*, *Ellioviricetes*, *Howeltoviricetes*, *Leviviricetes*, *Miaviricetes*, *Tolucaviricetes*, and proposed classes *c.0013*, *c.0031*, *c.0034*, *c.0069* in RVMT database(**Figures S36-41**). Overall, the variation trends of secondary structural features along classes of Lake Nam Co viral RNAs aligned with those of RVMT viral RNAs, indicating stable differences across taxa (**Figures 5C and S36-41A**). However, within viral classes, except *Ellioviricetes*, the secondary structural features of Lake Nam Co viral RNAs exhibit distinct density distributions, indicating some habitat specificity compared to RVMT references. (**Figures 5D and S36-41B**). For example, compared to RVMT references, *Amabiliviricetes* viruses of Lake Nam Co have fewer hairpins (wilcox test, *p* < 0.05) (**Figure S36B**) but more stems (wilcox test, *p* < 0.01) (**Figure S37B**) and interior loops (wilcox test, *p* < 0.05) (**Figure S38B**). In comparison to RVMT references, *Chrymotiviricetes* viruses of Lake Nam Co have fewer stems (wilcox test, *p* < 0.0001) (**Figure S37B**), interior loops (wilcox test, *p* < 0.01) (**Figure S38B**), multiloops (wilcox test, *p* < 0.05) (**Figures S39B**) but higher unpaired ratio (wilcox test, *p* < 0.001) (**Figure S40B**) and MFE (wilcox test, *p* < 0.0001) (**Figure S41B**).

We also analyzed the microdiversity of RNA viral genomes from Lake Nam Co by assessing nucleotide diversity (π), single nucleotide polymorphisms (SNPs), the non-synonymous to synonymous substitution ratio (pN/pS ratio), and fixation indices (F_ST_) (**Figure S42**). Nucleotide diversity ranged from zero to 5.15 × 10□³, with an average of 1.77 × 10□□ (**Figure S42 A**). SNP frequencies of Lake Nam Co RNA viral genomes averaged 0.47 SNPs per 1000 bp (**Figure S42 B**). Notably, 98% of the RNA viral genes exhibited pN/pS ratios below 0.4, with an average of 0.024 (**Figure S42 C**), indicating a predominance of synonymous substitutions and strong purifying selection in this high-altitude alpine lake. The F_ST_ values averaged 0.052, with 80% of pairwise fixation indices being zero (**Figure S42 D**), suggesting that the RNA viruses are conserved and homogeneous across different stations and depths.

### Host range of RNA viruses predominantly targeting prokaryotic organisms

We employed various methods to determine the relationship between RNA viruses and their hosts in Lake Nam Co. A total of eighty-five RNA viruses were identified with host evidence supported by the CRISPR-Cas spacer evidence, iPhop random forest algorithm, host genes, and related viruses (see Materials and Methods). Of these viruses, nine were predicted to infect eukaryotes, 75 to target bacteria, and one to potentially infect archaea (**Figure 6A**). Previous studies have identified only three groups known to infect prokaryotes: *Leviviricetes*, *Cystoviridae*, and *Picobirnaviridae*(38–40). However, host evidence in our study revealed 14 additional families with potential prokaryotic hosts in Lake Nam Co (**Table S7, S8**). These 14 families included *Totiviridae*, *Mitoviridae*, *Narnaviridae*, *Benyviridae*, *Marnaviridae* currently defined by ICTV, as well as *f.0028*, *f.0076*, *f.0102*, *f.0111*, *f.0176*, *f.0283*, *f.0286*, *f.0295*, and *f.0296* identified in the RVMT database but not yet officially proposed(16). Among these families, *Benyviridae* and *f.0176* are within *Kitrinoviricota*, a phylum previously unrecognized for bacterial infection. An RNA virus (contig ID: LS_coassembly_k141_1097885) from the family *Marnaviridae* was predicted to infect prokaryotic hosts according to the iPhop random forest algorithm (**Figure S43**). Further results indicate that it is most likely to infect archaea (confidence score: 94.7) and may also infect bacteria (confidence score: 91.4) (**Table S8**).

**Figure 6.**
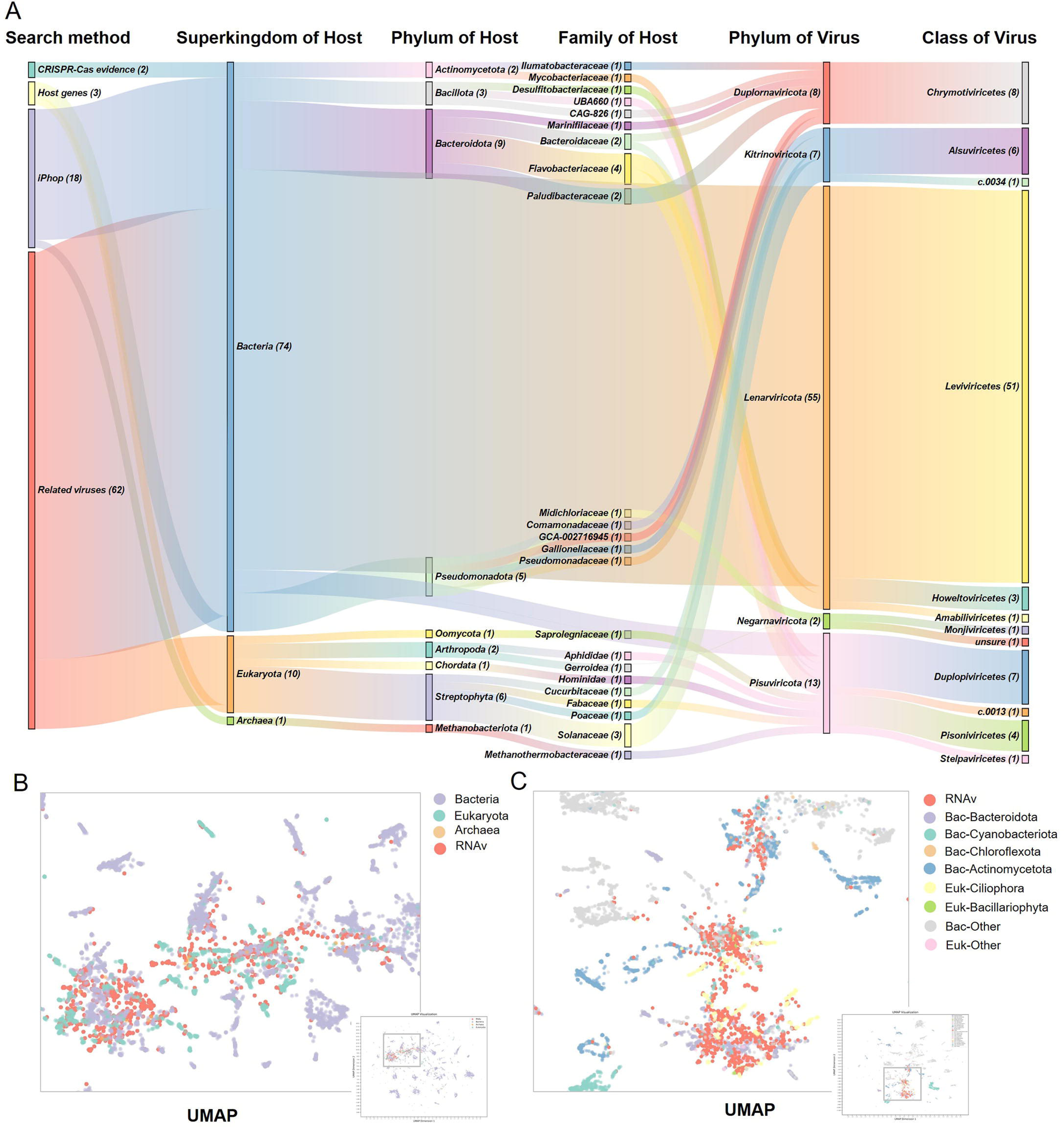
Host prediction for RNA viruses in Lake Nam Co. (A) Virus-host classification of RNA viruses with specific hosts. Most of the determined hosts are prokaryotes (74 hosts), and nine and one eukaryotic and archaeal hosts, respectively, were identified. (B) UMAP visualization of tetra-nucleotide usage frequency (TUF) of RNA viruses and microbes in Lake Nam Co at the super-kingdom level. Possible hosts with similar tetra-nucleotide usage frequency to RNA viruses cluster together in UMAP visualization. Most of RNA viruses cluster with prokaryotes (purple dots), while a limited portion of RNA viruses cluster with eukaryotes (blue dots). (C) UMAP visualization of TUF of RNA viruses and microbes in Lake Nam Co at the phylum level. Possible hosts with similar tetra-nucleotide usage frequency to RNA viruses cluster together in UMAP visualization. RNA viruses infecting bacteria primarily comprise *Bacteroidota*, *Cyanobacteriota*, *Chloroflexota*, and *Actinomycetota*.

To predict the overall potential host range of RNA viruses in Lake Nam Co that lack host evidence, we assessed their infection tendencies by calculating and visualizing amino acid usage frequency (AAUF) (**Figure S44A**) and di-/tetra-nucleotide usage frequency (D/TUF)(41, 42) (**Figures 6B, 6C and S44B**). Visualization of AAUF showed that viruses predominantly clustered independently, with a few overlapping with bacteria (**Figure S44A**). In contrast, RNA viral D/TUF closely matched their potential hosts, particularly bacteria, indicating an underlying preference for infecting prokaryotes (**Figures 6B, S44B**). According to the TUF aggregation of RNA viruses with cellular microbes at the phylum level, we predict that they primarily infected bacteria from *Bacteroidota*, *Actinomycetota*, *Chloroflexota*, and *Cyanobacteriota* phyla (**Figure 6C**).

We analyzed the relative abundance of prokaryotic RNA viruses across samples to investigate potential host and environmental influences (**Figures S45 and S46**). Prokaryotic viruses were most abundant in upper-middle water layers and decreased in deeper layers **(Figure S45)**. Distribution of individual prokaryotic viruses varied across samples **(Figure S46)**, with the most significant differences observed between sampling sites, followed by water layers. These differences align with the distribution of prokaryotic host groups **(Figures S1 and S47).**

## Discussion

This study represents the first comprehensive exploration of the RNA virome in a high-altitude alpine lake, revealing significant viral diversity and expanding the boundaries of the known RNA virosphere. Our findings unveil novel viral clades and unique adaptation strategies of RNA viruses. These discoveries offer new insights into RNA viral ecology and evolution in extreme environments.

### Novel RNA viruses and their evolutionary adaptations in Lake Nam Co

The abundance and diversity of RNA viruses depend on their environment and hosts, leading to significant differences across habitats, especially with unobserved taxa in extreme environments(17, 43). Our study identified 383 novel genus-level groups and 84 novel family-level groups, along with 14 unique phylogenetic clades and hints of three potential novel phyla. These findings emphasize significant differences in the viral composition of RNA viruses in Lake Nam Co compared to the global RNA virome(8, 14, 16, 32). The RNA viral diversity is mainly composed of *Lenarviricota*, *Pisuviricota*, and *Kitrinoviricota* viruses, whereas *Negarnaviricota* and *Duplornaviricota* make up a smaller portion of the global RNA virome(16). Similarly, the three diverse phyla are also prominent in Lake Nam Co, but the typically rare *Duplornaviricota* also shows high diversity here, comparable to *Kitrinoviricota* at the species level. This highlights the distinct and complex viral components of high-altitude lake ecosystems(27, 29, 31).

The conditions of Lake Nam Co—characterized by low temperatures, high ultraviolet radiation, and oligotrophic waters(31)—likely drive the evolution of these distinct viral communities. These viruses, which are distantly related in lineage of current RNA virome, exhibit genomic isolation and contain many genes encoding proteins of unknown function(44–47), contributing to the ‘dark matter’ of their genomes whose origins remain unclear(48). In addition, these RNA viral genomes exclusive to Lake Nam Co contain auxiliary metabolic genes (AMGs) acquired from their hosts, such as genes encoding cytochrome c oxidase subunit II and uroporphyrinogen-□ C-methyltransferase, along with the genes related to anti-prokaryotic immune systems. Viruses acquire host genes through exaptation to aid in replication and virus-host interactions, while host cells also gain viral genes for defense and optimize cellular functions(48). This gene flow is accompanied by an “arms race” between antivirus defense mechanisms of hosts and viral counter-defense systems adaptation, which is the core concept in virus evolution(48). Thus, such viral gene acquisitions in Lake Nam Co, which have not been previously reported in studies of orthornaviran viruses, indirectly related to energy production and nutrient metabolism of hosts, and directly linked to host immune systems, are considered one of the driving forces behind viral evolution in the enclosed and extreme environment(48). As a consequence of viral genome differentiation, the RNA viruses in Lake Nam Co have evolved distinct RNA secondary structures divergence from those of known RNA viruses(16) within the same viral class, including composition of loops and stems, as well as the MFE of these structures.

Moreover, compared to other ecosystems(12, 13, 16), the RNA viruses in Lake Nam Co have evolved significant differences at the protein level, particularly in distinctive secondary structures of proteins and permutated motifs in RdRP catalytic cores, which underscore the emergence of novel viral lineages adapted to extreme environments. The catalytic core of RdRP contains a “palmprint” region within the “palm” domain that includes three well-conserved catalytic motifs: A, B, and C(33, 49–51). According to previous studies, permutations of motif C have been observed in families *Birnaviridae* and *Permutotetraviridae*, which are distinct from the five major phyla of orthornavirans(11, 32). A recent global study revealed that these permutated families are associated with *Pisuviricota*, leading to their classification under the broader group of *Pisuviricota*(16). In contrast, our research reveals that the motifs permutation of *f.0167*_Nam_Co, which belongs to *Kitrinoviricota*, positions it on an intermediate evolutionary stage, indicating that *Birnaviridae* and *Permutotetraviridae* are related not only to *Pisuviricota* but also to *Kitrinoviricota*. The further comparison of physicochemical properties of RdRPs and capsid proteins(52, 53) of RNA viruses from Lake Nam Co, estuarine(12), and terrestrial environments(13) represents one of the adaptive consequences of the RNA virus proteins.

Microdiversity analysis of RNA viral populations from Lake Nam Co reveals evolutionary trajectories distinct from those of DNA viruses in other environments(54–60), as limited comparable RNA viral data are currently available. The nucleotide diversity of RNA viral populations in Lake Nam Co is significantly lower than that observed in DNA viral populations from global seawater(56) (3.78 × 10□□ on average) and soils with various land uses(57) (6.54 × 10□³ on average) (*p* < 0.0001). SNP frequencies in Lake Nam Co RNA viral genomes are also relatively low compared to those of the SARS-CoV-2 coronavirus(60), virophages from North American freshwater lakes(59), and dominant dsDNA viruses in the oceans(58). However, the nucleotide diversity and SNP frequencies of these RNA viruses are comparable to those of deep-sea cold seep viruses, which also inhabit low-temperature, high-salinity environments(54). The exceptionally low pN/pS ratios of Lake Nam Co RNA viruses, compared to both cold seep viruses(54) (0.13 on average) and viruses from underground saline waters(55) (0.86 on average) (*p* < 0.0001), indicate that the majority of the RNA viruses are under strong purifying selection pressures(61, 62). The low F_ST_ values between samples reveal genetic conservation, reflecting homogeneous viral genomes across different stations and depths in such enclosed high-altitude lake(63).

### Expansion of the RNA viral host range towards prokaryotes

The expansion of the RNA viral host range to prokaryotes, including bacteria and potentially archaea, is one of the striking findings of this study. Traditionally, RNA viruses were thought to predominantly infect eukaryotes, with relatively few known examples of RNA bacteriophages(16, 64). Although in recent years, the diversity of RNA viruses infecting bacteria has significantly increased within the class *Leviviricetes* of the phylum *Lenarviricota*, the family *Cystoviridae* of the phylum *Duplornaviricota*, and the family *Picobirnaviridae* of the phylum *Pisuviricota*(16, 38–40), the known RNA viruses that can infect bacteria are still limited to the aforementioned classifications(11). However, the evidence for the 14 previously unknown putative RNA virus families that infect prokaryotes in Lake Nam Co, along with the similarity of the overall RNA virus AAUF and D/TUF(65) in Lake Nam Co to those of prokaryotes, challenges this notion and suggests that the host range of RNA viruses is far broader than previously recognized(66). The inclusion of diverse bacteriophage clades in the RNA virome underscores extensive diversity across environments(16, 17, 64). In alpine extreme environments, where prokaryotes dominate(24, 67), RNA viruses may preferentially target bacterial hosts, potentially leading to underlying pathways of cross-species transmission, as these bacterial hosts could act like vectors, similar to arthropods in eukaryotic RNA virus transmission(68–70). This opens new avenues for research into virus-host co-evolution and viral contributions to microbial community structure in extreme environments(16, 17).

### Ecological implications of RNA viruses in extreme alpine environments

The ecological implications of our findings transcend the specific context of Lake Nam Co. With climate change exerting growing pressures on extreme environments(71), unraveling the roles of RNA viruses in these ecosystems becomes increasingly critical, as their influence on microbial dynamics could have far-reaching consequences for ecosystem resilience and global biogeochemical processes(72). Viruses in aquatic environments shape host community composition and diversity through lysis(73), gene transfer(74), and the expression of AMGs, which facilitate specific host metabolism(75–77). Additionally, viruses affect the efficiency of the biological pump(78) and the microbial carbon pump(79) through processes like the viral shunt(75, 80, 81), thereby influencing microbial-driven biogeochemical cycles in their habitat. The discovery of novel viral clades, expansion of host ranges, and AMGs all point to the potential for RNA viruses to play a central role in ecosystem resilience and adaptation, similar to demonstrated DNA viruses(82, 83). The presence of AMGs related to host energy metabolism in the alpine RNA viral genomes suggests that these viruses may actively modulate host metabolic processes to ensure replication in nutrient-limited environments(25, 31, 84), suggesting a previously unrecognized role of RNA viruses in influencing microbial-driven nutrient cycles and energy flow in such ecosystems(25, 31, 85). The evolutionary features such as the permutation of conserved RdRP motifs within *f.0167*_Nam_Co, reflect the pressures faced by viruses in extreme environments, highlighting their potential for evolving mechanisms of replication and survival(86). Overall, the complex and highly diverse assemblages of Lake Nam Co RNA viruses—ranging from variations in genetic material and protein structures to differences in host preferences—contributes to the unique and isolated ecological composition of high-altitude lakes.

Moreover, the potential for cross-species transmission, coupled with the adaptations of viruses in isolated environments, raises concerns about the emergence of novel viral pathogens(18, 87). There are significant variations in the viral communities in glaciers on the Tibetan Plateau under different climate conditions(88). Glacier melt induced by global warming has led to changes in hydrological conditions, such as increased water levels in Lake Nam Co. By combining immediate changes with potential delayed responses(89), these shifts result in more complex microbial inputs(31) and create conditions for the continued co-evolution of viruses and hosts. In light of global concerns over pandemics like COVID-19(90, 91), it is crucial to consider the potential risks posed by ancient or isolated viruses, especially in extreme environments like permafrost or alpine lakes(92). It has been reported that climate change thaws permafrost or alter isolated ecosystems, potentially releasing ancient viral pathogens that may pose a threat to public health and global ecosystems(71, 87). Our findings underscore the importance of viral surveillance in such environments to proactively address emerging risks.

## Materials and Methods

### Sample collection

Water samples were collected from two sampling stations in Lake Nam Co, Tibet, China (**Figure 1 and Table S1**) **-** the LS station near the lakeshore (90.98°E, 30.79°N) and the LC station near the lake center (90.48°E, 30.46°N). At the LS station, 20 l of surface water (at a water depth of 0.5 m) was collected. Simultaneously, at the LC station, samples were taken from four different layers – surface (0.5 m), upper-middle (20 m), lower-middle (50 m), and bottom (80 m), with 20 l collected from each layer. Surface water samples at 0.5 m depth were manually collected using 25L PC barrels. For samples at 20 m, 50 m, and 80 m depths, a deep well pump (Xinmutian Electrical & Mechanical Co., Ltd., Zhejiang, China) was used to collect the water, which was then transferred into 25L PC barrels. All water samples were pre-filtered using nylon sieves with a pore size of 60 µm and then filtered through polycarbonate membranes (Millipore, New Jersey, USA) of different pore sizes within 5 hours after collection, using 142 mm acrylic filter discs (Tianxiang Ocean Technology Co., Ltd., Xiamen, China). Membranes with pore sizes of 10 µm, 3 µm, 1.2 µm, and 0.2 µm were collected, rapidly frozen with liquid nitrogen, and stored at –80□ until RNA extraction. Additionally, 40 ml of filtrate was transferred to a 50 ml clean tube and stored at –20□ for inorganic nutrient analysis. For dissolved organic carbon (DOC) analysis, water samples were filtered through 0.7-µm GF/F glass fiber filters (pre-combusted at 450□ for 4 hours, Whatman, USA). The 20 ml filtered water samples were collected directly into two 40 ml glass vials (acid-washed, rinsed with Milli-Q water, and pre-combusted, CNW, Germany) and immediately stored at –20□ for DOC concentration measurements.

### Measurement of environmental parameters

Solar radiation intensity was measured using Handheld TES-1333 solar power meter digital radiation detector. Water temperature, pH, dissolved oxygen (DO), and salinity were measured using in situ Conductivity-Temperature-Depth (CTD) sensors (Sea-Bird Electronics, Bellevue, WA, USA). Inorganic nutrient concentrations, including phosphate, nitrate, nitrite, and silicate, were measured using the PowerMon Kolorimeter AA3 (Bran + Luebbe, Charlotte, NC, USA) following the spectrophotometric methods(93). The DOC concentrations were determined using a Shimadzu TOC-VCPH analyzer through high-temperature (680□) catalytic oxidation. Ammonium concentrations were measured spectrophotometrically using the indophenol blue method.

### 16S/18S rRNA transcripts amplicon sequencing and data processing

Total RNA was extracted using the DNA/RNA Miniprep Kit (Zymo Research Corporation) according to the manufacturer’s instructions and subsequently stored at –80□. One part of this RNA was reverse transcribed into cDNA using random hexamer primers and SuperScript III Reverse Transcriptase (Invitrogen), as per the manufacturer’s guidelines. To identify active microbial communities, the V4-V5 regions of prokaryotic 16S rRNA were amplified using forward (515F 5’-GTGCCAGCMGCCGCGGTAA-3’) and reverse (907R 5’-CCGTCAATTCMTTTRAGTTT-3’) primers. Similarly, the V4 region of microeukaryotic 18S rRNA was amplified with forward (TAReuk454FWD1 5’-CCAGCASCYGCGGTAATTCC-3’) and reverse (TAReukREV3 5’-ACTTTCGTTCTTGATYRA-3’) primers. Sequencing of the 16S/18S rRNA transcripts was conducted on an Illumina NovaSeq 6000 system (pair-end sequencing, 2 × 250 bp) by Magigene (Guangdong, China). For the analysis, Qiime2 v2021.8.0 was employed to process the 16S/18S rRNA sequencing data from the 20 samples(94). This process included merging, quality filtering, denoising, and the exclusion of sequences originating from chloroplasts and mitochondria. Additionally, the software supported the clustering and annotation of operational taxonomic units (OTUs), enabling the profiling of the taxonomic composition.

### Metatranscriptomic sequencing and data processing

Part of total RNA was subjected to rRNA depletion using the Ribo-off® rRNA Depletion Kit V2. This rRNA-depleted RNA was then reverse-transcribed into cDNA using random hexamer primers and SuperScript III Reverse Transcriptase (Invitrogen). Metatranscriptomic sequencing was performed on the Illumina NovaSeq 6000 platform (pair-end sequencing, 2 × 150 bp) by Magigene (Guangdong, China). For the metatranscriptomic sequencing data, Fastp v0.23.2 was utilized for trimming and quality control, retaining only high-quality clean reads(95). These clean reads were then mapped to the SortMeRNA database v4.3.4(96) for separating rRNA reads from non-rRNA reads using Bowtie2 v2.4.3(97). Non-rRNA reads were assembled using MEGAHIT v1.2.9(98) to generate assembled contigs, employing both single-sample assembly and multiple-sample co-assembly (LC + LS), resulting in a total of 22 assemblies. The getORF package from EMBOSS v6.6.0.0(99) and Prodigal v2.6.3(100) were used for open reading frame (ORF) prediction from the assembled contigs (using the 11^th^ codon table), preserving ORFs longer than 200 amino acids.

### Identification of RdRP sequences

To enhance the detection of RNA viruses within our datasets, we applied a comprehensive approach combining traditional sequence alignment, hidden Markov model (HMM) searches, and a deep learning algorithm (**Figure 1**). Specifically, we utilized: (1) BLASTp analysis (Diamond v2.0.15.15363)(101) to compare our sequences against a custom database comprising RdRP sequences gathered from four recent studies on RNA virus diversity(8, 14, 16, 32); (2) HMM searches (HMMER v3.3.2)(102) targeting 77 different RdRP family profiles to identify potential matches(103); (3) deep learning analysis through LucaProt (https://github.com/alibaba/LucaProt)(8), focusing on ORFs longer than 300 amino acids. Through this multifaceted approach, we initially identified 4,021 potential RdRP sequences (**Figure 1**).

To refine our results for species-level analysis of RdRP sequences, we applied CD-HIT version 4.8.1 to eliminate duplicates from the sequences, using a 90% amino acid identity (AAI) similarity threshold(49, 104, 105), reducing the count to 3,089 non-redundant candidate RdRP sequences. Further analysis for identifying three core RdRP motifs (Motifs A, B, and C) was conducted using PalmScan2(49, 50). Sequences exhibiting all three motifs were considered viable RdRP candidates, resulting in a total of 809 sequences. These candidates underwent an additional check against the non-redundant (nr) database using Diamond BLASTp v2.0.15.15363 to ensure their viral origin. We excluded sequences whose top BLAST hits (e-value < 0.001) did not match known RNA virus proteins, thereby eliminating false positives and potential endogenous viral elements(10). This careful filtering process yielded 742 sequences that were identified as RdRPs and selected for further analysis (**Figure 1**).

### Phylogenetic tree construction and taxonomic assignment of RdRPs

The taxonomic classification of newly discovered RdRPs was determined through sequence-alignment-based and protein structure-based clustering analyses. For the sequence alignment approach, the determination of RdRP taxonomy was informed by a synthesis of the top BLASTp hits across a range of databases, incorporating the valuable contributions from the RVMT database(16), as well as research conducted by Hou et al.(8), Charon et al.(32), and Zayed et al.(14). For the names of taxa that have not been officially proposed, we have retained the provisional classifications from the RVMT database, which were derived from the work of Neri et al.(16). RdRPs with taxonomy references here were considered to be sequence-alignment-based identified RdRPs, while others were identified as sequence-alignment-based unidentified RdRPs.

For the phylogenetic analysis of sequence-alignment-based identified RdRPs, due to the rapid evolution of RNA viruses and substantial differences in RdRp sequences among different classes, multiple gaps occurred during alignment, hindering the construction of a robust phylogenetic tree. Therefore, separate alignments were performed for the five phyla of RNA viruses: *Duplornaviricota*, *Kitrinoviricota*, *Pisuviricota*, *Lenarviricota*, and *Negarnaviricota*. Subsequently, maximum likelihood phylogenetic trees for RdRp proteins were constructed. For classes with over 100 identified RdRp sequences, sub-alignments were performed based on evolutionary distances and protein clustering numbers, followed by sub-trees construction. Based on different classifications, the identified complete RdRPs was aligned with reference sequences using MAFFT v 7.508(106) with the “--auto” option. Initial trimming of aligned regions was performed using trimAl v 1.4.rev15(107) with the options “-resoverlap 0.70” and “-seqoverlap 75”. Sequences were manually trimmed to the adjacent area of core RdRP region (including the three conserved motifs A, B, and C) and sequences causing alignment confusion were manually removed using Jalview v 2.11.3.2(108). FastTree v 2.1.10 was used to construct maximum likelihood phylogenetic trees for each phylum and class(109), the trees were then visualized using the Chiplot online website (https://www.chiplot.online/)(110). New taxonomy groups of Lake Nam Co were then identified according to the amino acid similarity.

To conduct the structure verification of RdRP candidates and ascertain the approximate clades of sequence-alignment-based unidentified RdRPs, Alphafold2 v dtk22.04(111) was used to predict the structures of all RdRP candidates in Lake Nam Co and their reference sequences(8, 14, 16, 32, 103). All candidates with pLDDT (predicted Local Distance Difference Test) values below 60 were excluded from further analysis. We conducted the paired structure alignment based on template modeling score (TM-score) method via US-align v 20230609(112), which outputted the TM-scores and the root-mean-square deviation (RMSD) values for comparisons between structures. Subsequently, we performed the structure-based clustering of all predicted RdRP structures based on the TM-score distance matrix(113). RdRP candidates with an RMSD value over 5 compared to all references were excluded (Establishment of RdRP dataset part). To eliminate the potential impact of non-core region sequences bias (too much amino acid on the left or right) on structural similarity, all sequence-alignment-based identified RdRP sequences were trimmed to the relative conserved domain according to the sequence alignment results, while the sequence-alignment-based unidentified RdRPs remain untrimmed before re-clustering implemented the same way as described above. For a sequence-alignment-based unidentified RdRP, the phylum was determined by the RdRP structure with the lowest RMSD value. If the structure with the lowest RMSD value was a sequence-alignment-based identified RdRP from Lake Nam Co, an additional reference structure from RdRP identification databases(8, 14, 16, 32, 103) with the lowest RMSD value was also included. We used the align command in PyMOL v2.5.5(114, 115) to compare and visualize the catalytic core regions(49) comprising three essential motifs of sequence-alignment-based unidentified RdRP structures against reference structures. The phylum of sequence-alignment-based unidentified RdRPs was then determined based on the RMSD values of catalytic core regions output by this command, with lower RMSD values indicating higher similarity to the reference structure(116). After integrating the phylum classification of sequence-alignment-based identified RdRPs and sequence-alignment-based unidentified RdRPs, the conservative motifs A, B, and C for different phyla of RNA viruses were visualized using WebLogo v 2.1.10(117).

All reference sequences and newly identified RdRPs were subjected to palmprint recognition by PalmScan2(49, 50) with default parameters. Palmprints were clustered using MMseq2 v 14.7e284(118) at 75% and 40% identity thresholds to identify clusters at the genus-level and family-level(49) composed solely of newly identified RdRPs from Lake Nam Co.

### RNA virus genome annotation and distribution

All contigs containing the identified RdRP of Lake Nam Co were considered as RNA viral contigs. To ensure that the contigs represent indeed viral genomes and not assembly errors, we used metagenomic sequencing reads from the same environment to map to these contigs. Contigs that that had more than 10 reads mapped from DNA sequencing data were excluded(8). After redundancy reduction of nucleotide sequences with a threshold of 0.95 using CD-HIT v4.8.(104), these genomes were annotated using tools such as Cenote-Taker v3.0.0(44), eggNOG-mapper v2.1.12(45), and dbCAN3 v4.0.0(46), along with annotation using DRAM v1.2.0(47) after preprocessing with Virsorter2 v2.2.4(119). The dbAPIS database was also utilized for the annotation of Anti-prokaryotic Immune System (APIS) proteins in these RNA viruses(36), using Diamond BLASTx v2.0.15.15363 with the default threshold (e-value < 0.001) (101).

### Comparison of physicochemical property indices of RdRP and capsid proteins in RNA viruses from different environments

RdRP and capsid proteins of Yangshan estuary environment(12) and terrestrial environment across China(13), together with the RdRP and capsid proteins annotated in Lake Nam Co were collected. The physicochemical properties indices, including hydrophobicity, polarity, relative mutability, transmembrane tendency, and refractivity were calculated refer to ProtScale website (https://web.expasy.org/protscale/). A linear weighted model (window size of 9) was applied to calculate scores at different positions in the protein sequences for each of the physicochemical properties. The variation of individual protein indicators across positions were depicted by line plots, while the comparison of all physicochemical properties indices between different locations were depicted by violin plots drawing by R v 4.2.0.

### Comparison of RNA secondary structures between RNA viruses in Lake Nam Co and global data

Classes containing over 10 RNA virus sequences in Lake Nam Co were selected, and 100 viral contigs from these classes were randomly extracted from the RVMT database(16) for comparison. RNA secondary structure features (**Figure S35**) of these classes, including hairpins, stems, interior loops, multiloops, unpaired ratios, and minimum free energy (MFE), were analyzed and visualized refer to the script published by Uri Neri’s laboratory (https://github.com/UriNeri/RNAfold_virus_Rima).

### Assessment of genome-wide evolutionary metrics of Lake Nam Co RNA viral populations

The non-rRNA reads of all 20 samples were mapped to the identified genomes using Bowtie2 v 2.4.3(97), and the genes of genomes were annotated using Prodigal v2.6.3(100). Subsequently, the abundance and diversity of RNA viruses in Lake Nam Co were determined by MetaPop v 1.0.0(120) with default parameters and the “--snp_scale both” option. To ensure sufficient coverage for accurate SNP calling and evaluation of contig-level microdiversity, viral genomes with less than 70% length of genome covered or an average read depth coverage below 10× were excluded for microdiversity analyses(121). SNP frequencies were subsampled to 10× coverage for the evaluation of nucleotide diversity (π) at gene/genome levels, as well as fixation indices (F_ST_) of genomes and selective pressures on genes (pN/pS)(54). Visualizations and comparisons of the SNP frequencies, π, pN/pS ratios, and F_ST_ values of RNA viral contigs were generated using the ggplot2 and ggpubr packages in R v 4.2.0.

### RNA Virus-Host relationships

To find CRISPR-Cas spacer evidence, we used Diamond BLASTx v2.0.15.15363 with the default threshold (e-value < 0.001)(101) to search for corresponding CRISPR-Cas spacers in a CRISPR spacers dataset released with IMG/VR v4. Furthermore, leveraging iPhop v1.3.3(122) for viral host prediction, in addition to host information of closely related viruses identified through sequence alignments, and annotated viral sequences containing host genes, we identified potential hosts for Lake Nam Co RNA viruses. Kaiju v 1.10.0 was employed to predict the taxonomic classification of all contigs from Lake Nam Co, using the default threshold (e-value < 0.01). Amino acid usage frequency (AAUF) and the di-/tetra-nucleotide usage frequency (D/TUF) of the contigs exceeding 10,000 nucleotides along with the identified RNA viral contigs identified were then predicted and visualized utilizing scripts published in previous research(65).

## Acknowledgments

We acknowledge the financial support provided by the National Natural Science Foundation of China (42188102, 42222604, 92351303 and 92251306), National Key Research and Development Project of the Ministry of Science and Technology of China (Grant No. 2021QZKK0102). We thank Dr. Fabai Wu, Dr. Shengwei Hou, Dr. Xin Hou and Dr. Donald Pan for providing valuable comments. We thank Lin Zang, Yunlan Yang, Yunxuan Li, Tianhui Wang, and Wenqiang Wang for their assistance during the field experiments and support in providing environmental parameters. We thank Yongyi Peng, Jing Liao, Yingchun Han, Xiaomeng Wang, Qiuyun Jiang, and Xingyu Wang for their guidance in data analysis.

## Author contributions

Z.Q., D.X., and L.Y. initiated and designed the study. W.L., L.K., X.J., and L.P. carried out field sampling. W.L. performed total RNA extraction. W.L., Z.Q., D.X., and S.W. performed all analyses of metagenomes, metatranscriptomes, and RNA viral communities with the help of L.Z., H.Y. and S.M.. W.L., Z.Q., D.X., and others led the writing of the manuscript, with result interpretations from J.N. and C.T.. C.T. and S.M. helped revise the paper. All authors contributed to editing the text and approved the final version.

## Classification

Biological Sciences/ Microbiology

