## Supplementary material for "Uncovering the hidden RNA virus diversity in Lake Nam Co: evolutionary insights from an extreme high-altitude environment": Figures S1 to S47 & Tables S1 to S9: Supplementary Information.docx

**This file includes:**

Supporting text

Figures S1 to S47

Tables S1 to S9

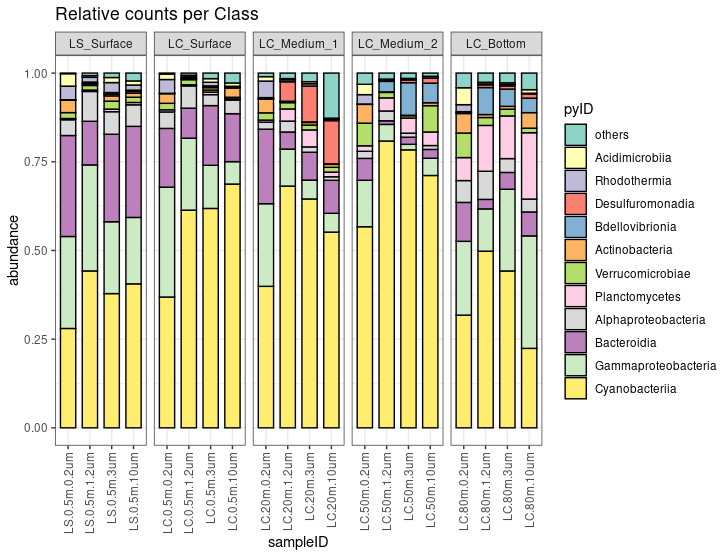

**Figure S1. Prokaryotic community compositions (class) for each water samples.** Relative proportions were determined by the normalized number of reads from 16S rRNA amplicon sequencing. The layers of the water samples are shown at the top of the bar. LS, lakeshore; LC, lake center.

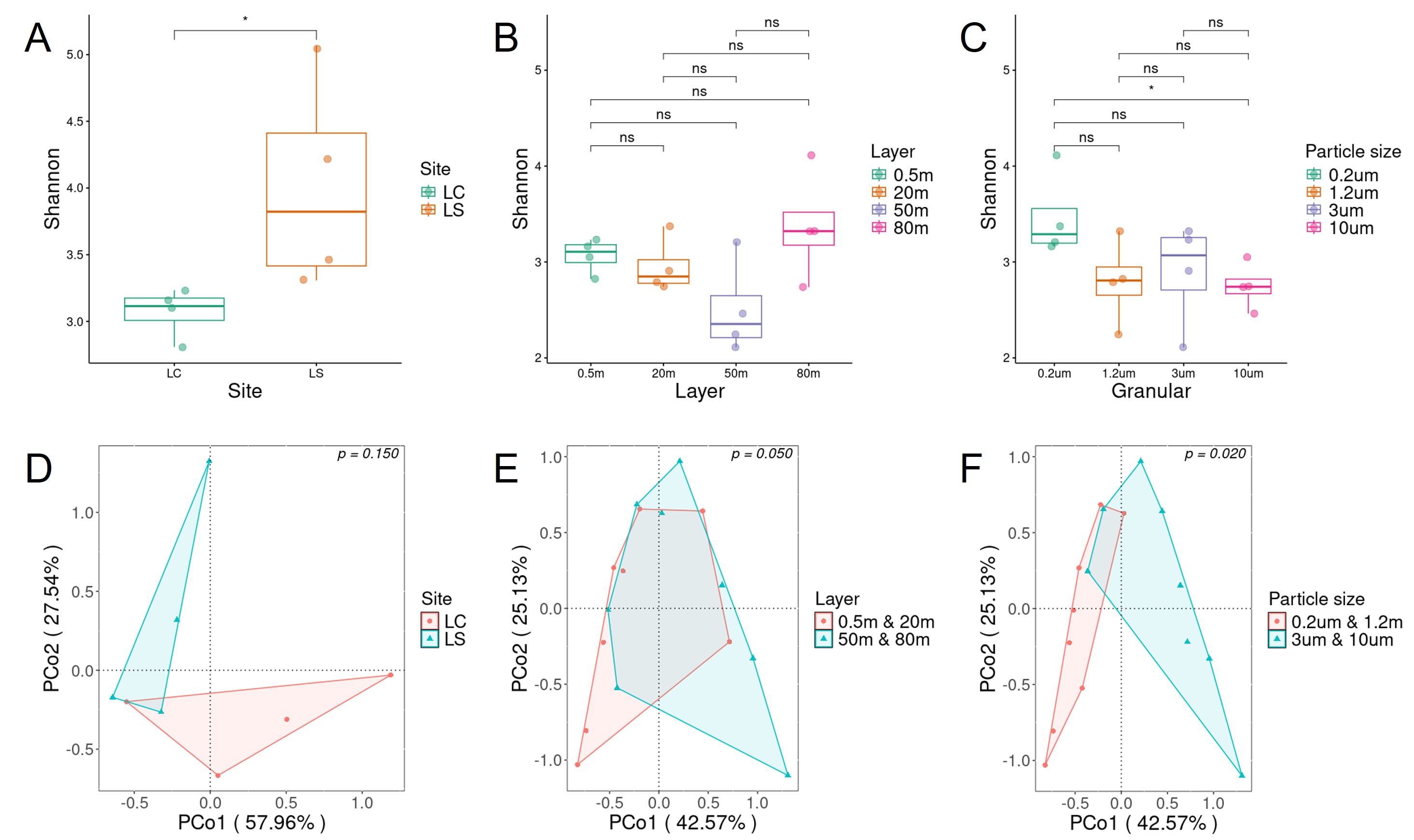

**Figure S2. Alpha and beta diversity of prokaryotic communities in Lake Nam Co across different particle sizes, water layers, and locations.** Alpha diversity is represented by the Shannon index, while beta diversity is based on Bray-Curtis distance. (A) Shannon index of prokaryotic communities at different sites in Lake Nam Co. (B) Shannon index of prokaryotic communities across different layers of LC station in Lake Nam Co. (C) Shannon index of prokaryotic communities for different particle sizes of LC station in Lake Nam Co. (D) PCoA visualization of the Bray-Curtis distances between prokaryotic communities of different sites in Lake Nam Co. (E) PCoA visualization of the Bray-Curtis distances between prokaryotic communities of different layers of LC station in Lake Nam Co. (F) PCoA visualization of the Bray-Curtis distances between prokaryotic communities of different particle sizes of LC station in Lake Nam Co. LS, lakeshore; LC, lake center.

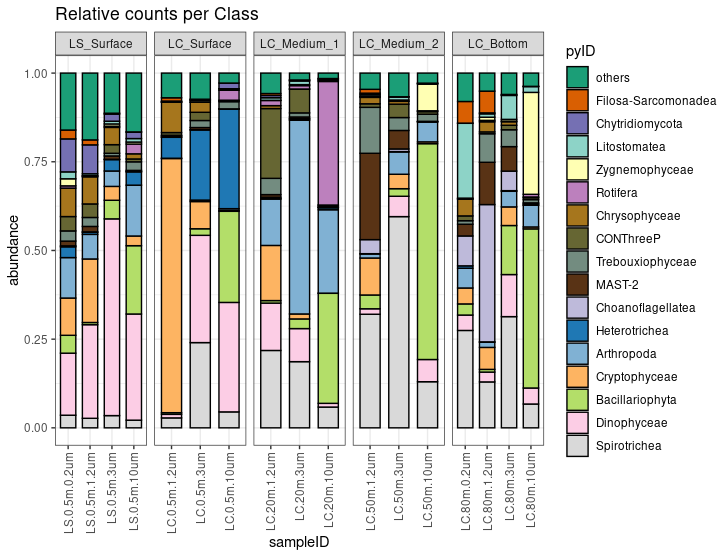

**Figure S3. Eukaryotic community compositions (class) for each water samples.** Relative proportions were determined by the normalized number of reads from 18S rRNA amplicon sequencing. The layers of the water samples are shown at the top of the bar. LS, lakeshore; LC, lake center.

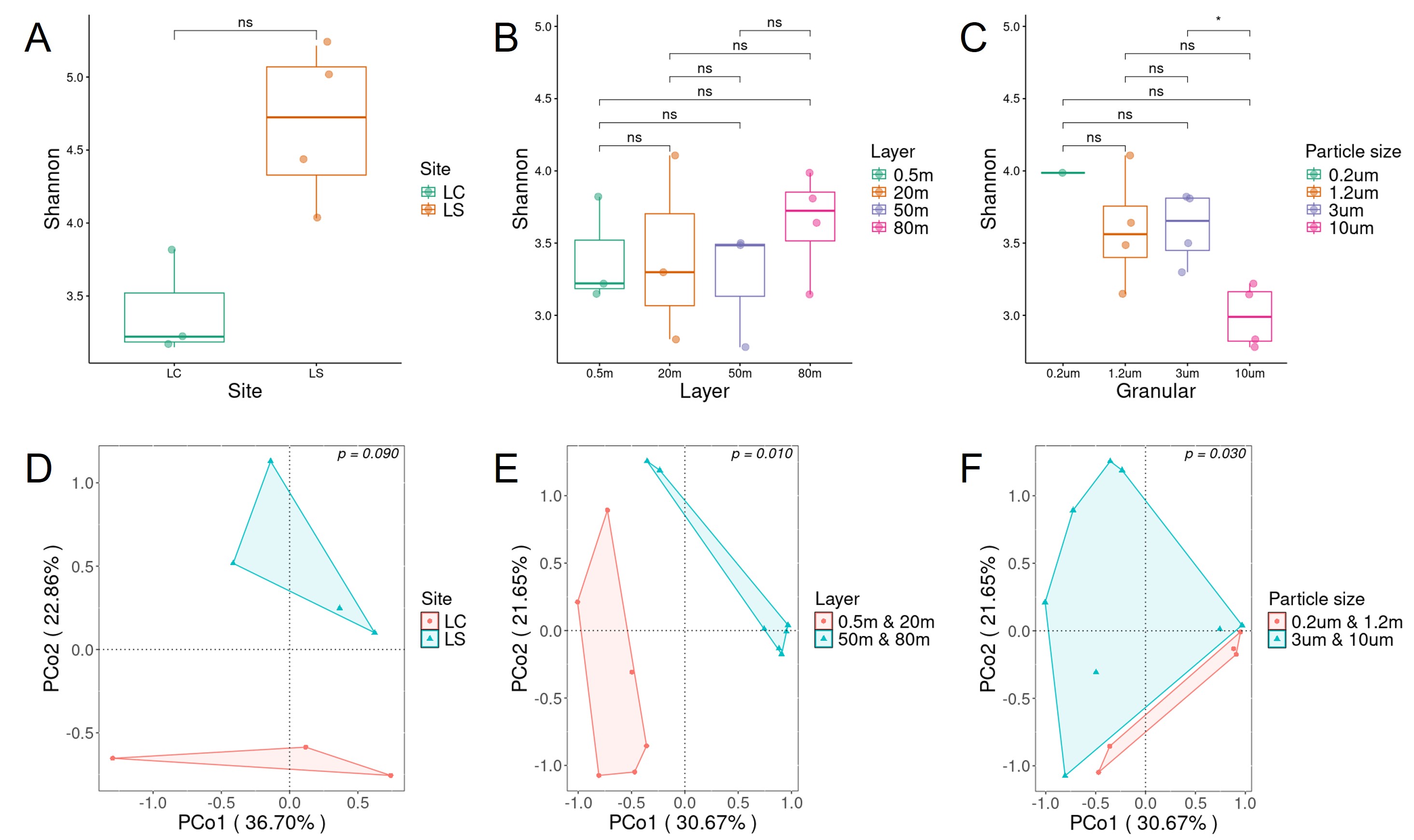

**Figure S4. Alpha and beta diversity of eukaryotic communities in Lake Nam Co across different particle sizes, water layers, and locations.** Alpha diversity is represented by the Shannon index, while beta diversity is based on Bray-Curtis distance. (A) Shannon index of eukaryotic communities at different sites in Lake Nam Co. (B) Shannon index of eukaryotic communities across different layers of LC station in Lake Nam Co. (C) Shannon index of eukaryotic communities for different particle sizes of LC station in Lake Nam Co. (D) PCoA visualization of the Bray-Curtis distances between eukaryotic communities of different sites in Lake Nam Co. (E) PCoA visualization of the Bray-Curtis distances between eukaryotic communities of different layers of LC station in Lake Nam Co. (F) PCoA visualization of the Bray-Curtis distances between eukaryotic communities of different particle sizes of LC station in Lake Nam Co. LS, lakeshore; LC, lake center.

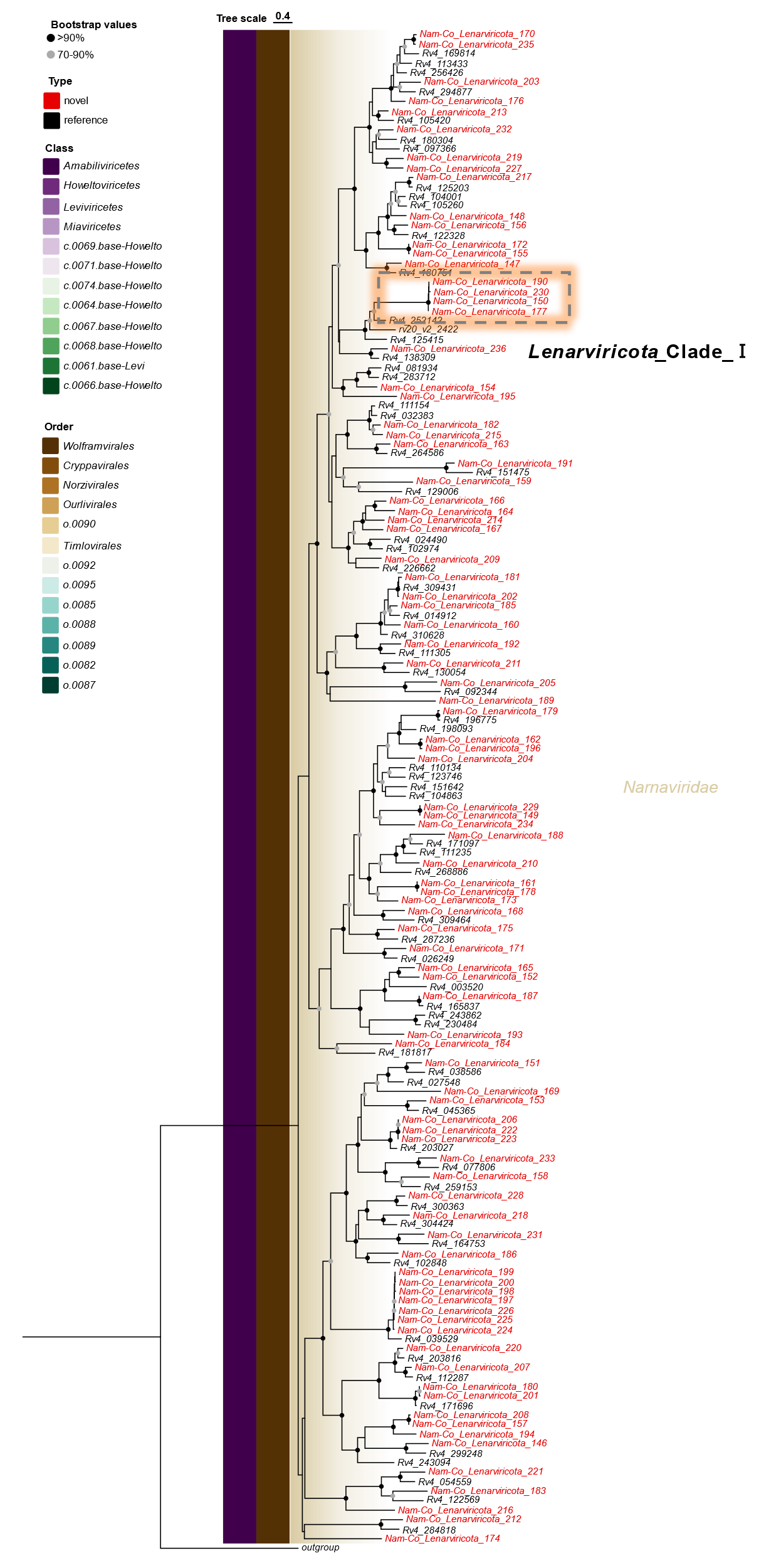

**Figure S5. The subtree I for *Lenarviricota* including family *Narnaviridae*.** Names of RdRP identified in Nam Co are highlighted in red. Color blocks from left to right denote the classes and orders of Nam Co RdRPs’ top hits in BLASTp, with gradient blocks and their labels indicating the families of Nam Co RdRPs’ top hits in BLASTp. The names of RdRPs correspond to specific ORF sequences in Table S4.

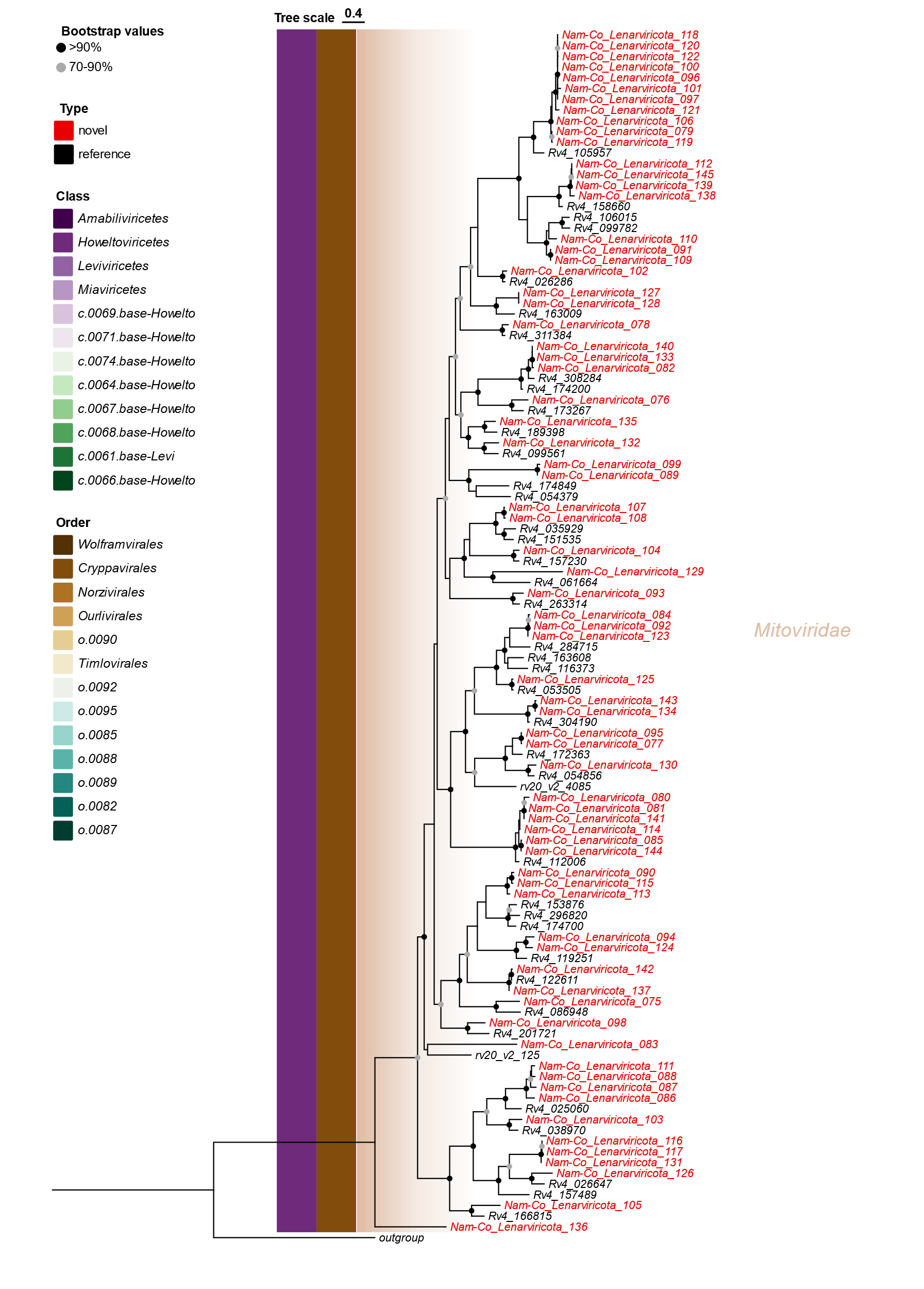

**Figure S6. The subtree II for *Lenarviricota* including family *Mitoviridae*.** Names of RdRP identified in Nam Co are highlighted in red. Color blocks from left to right denote the classes and orders of Nam Co RdRPs’ top hits in BLASTp, with gradient blocks and their labels indicating the families of Nam Co RdRPs’ top hits in BLASTp. The names of RdRPs correspond to specific ORF sequences in Table S4.

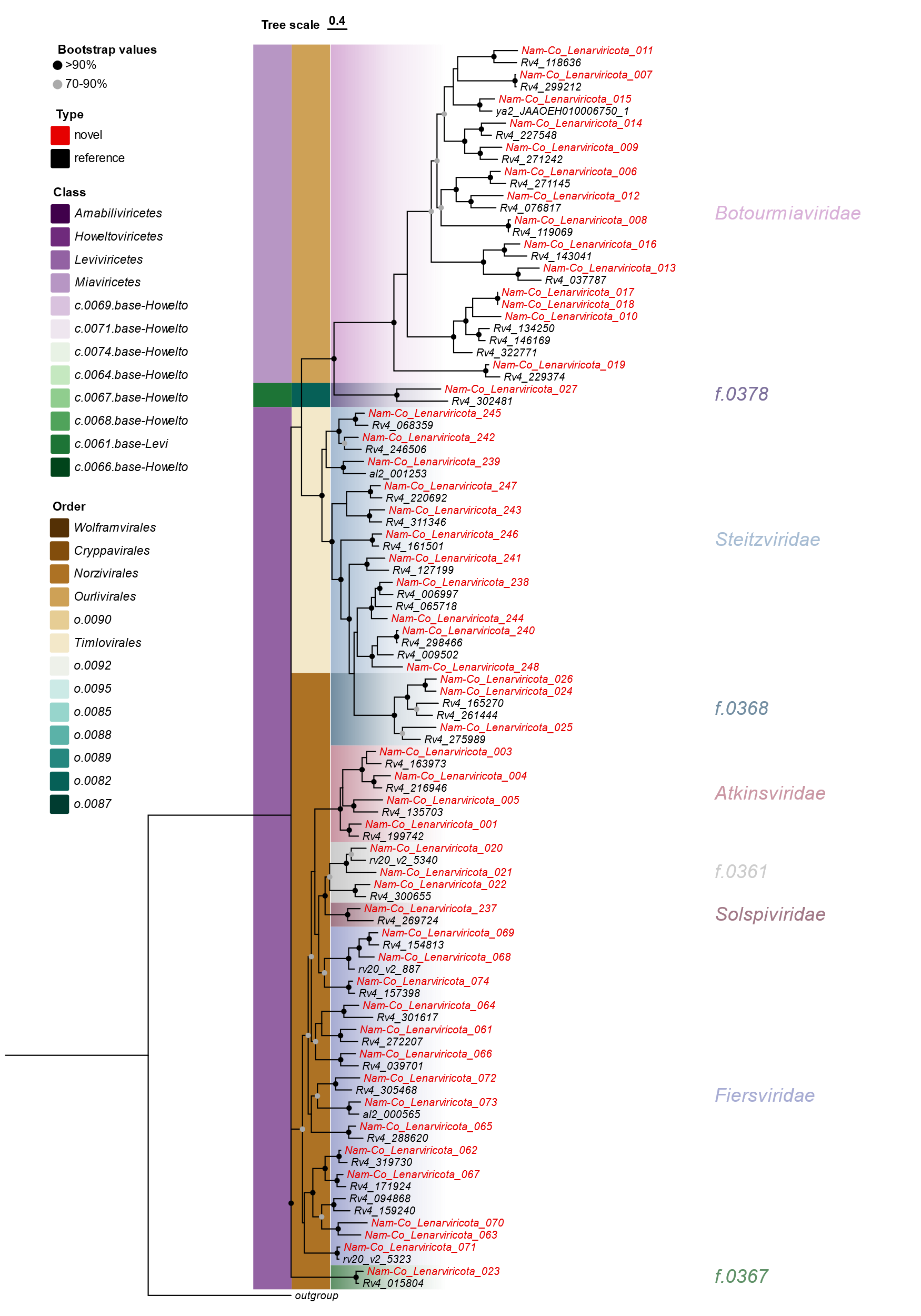

**Figure S7. The subtree III for *Lenarviricota* including families *Steitzviridae*, *f.0368*, *Atkinsviridae*, *f.0361*, *Solspiviridae*, *Fiersviridae* and *f.0367*.** Names of RdRP identified in Nam Co are highlighted in red. Color blocks from left to right denote the classes and orders of Nam Co RdRPs’ top hits in BLASTp, with gradient blocks and their labels indicating the families of Nam Co RdRPs’ top hits in BLASTp. The names of RdRPs correspond to specific ORF sequences in Table S4.

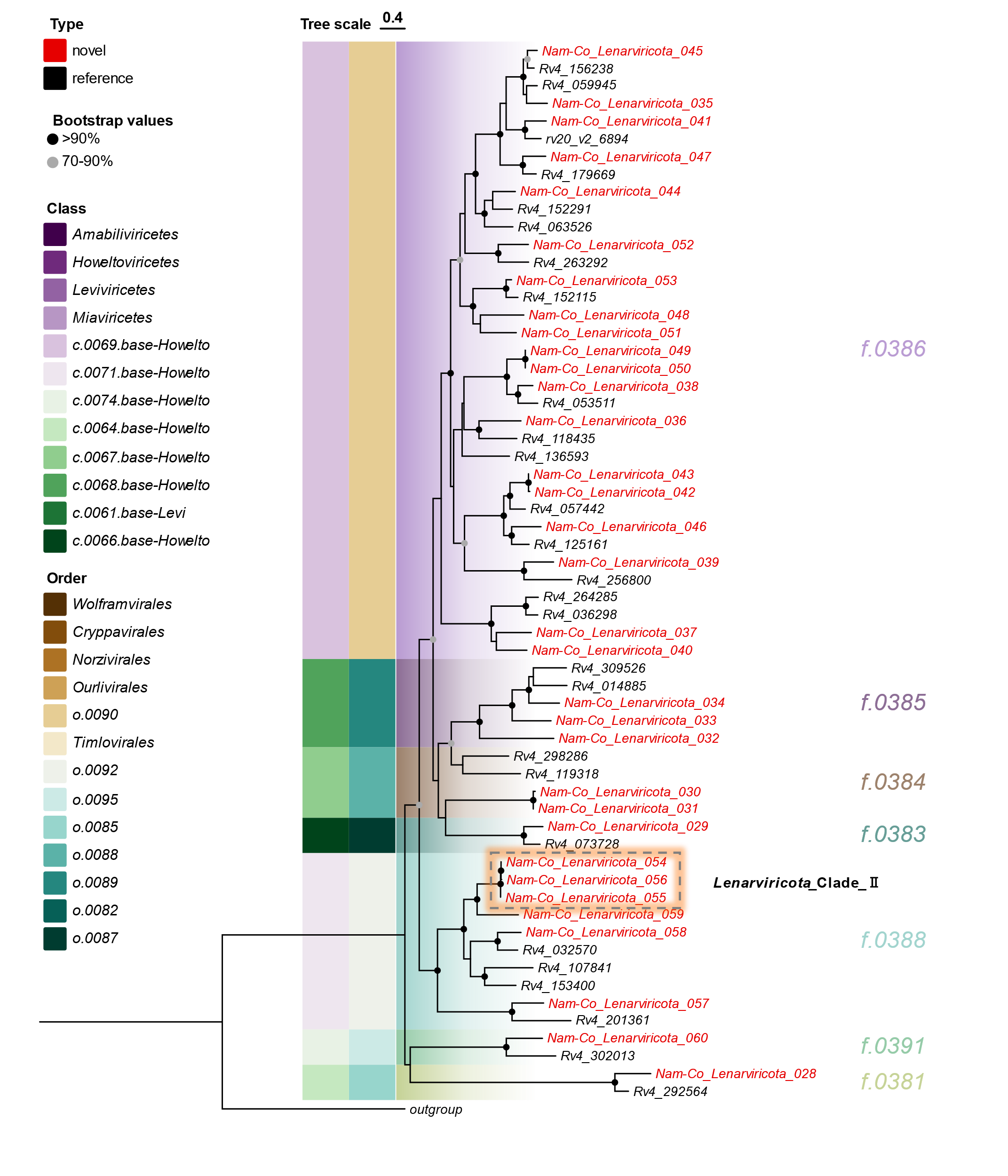

**Figure S8. The subtrees Ⅳ for *Lenarviricota*** **including families *f.0381*, *f.0383*, *f.0384*, *f.0385*, *f.0386*, *f.0388* and *f.0391*.** Names of RdRP identified in Nam Co are highlighted in red. Color blocks from left to right denote the classes and orders of Nam Co RdRPs’ top hits in BLASTp, with gradient blocks and their labels indicating the families of Nam Co RdRPs’ top hits in BLASTp. The names of RdRPs correspond to specific ORF sequences in Table S4.

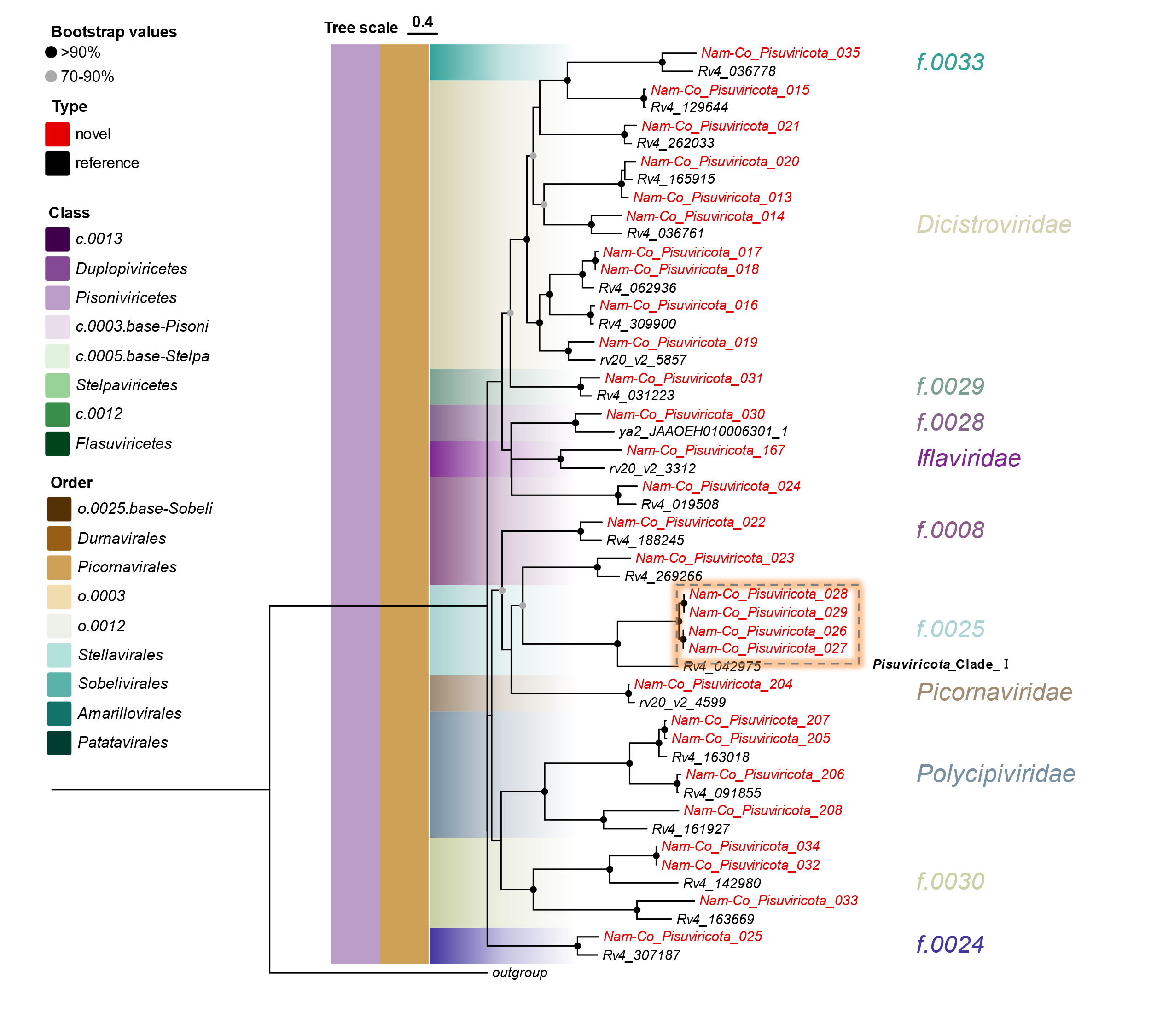

**Figure S9. The subtrees Ⅰ for *Pisuviricota* including families *Dicistroviridae*, *Iflaviridae*, *Picornaviridae*, *Polycipiviridae*, *f.0008*, *f.0024*, *f.0025*, *f.0028*, *f*.0029, *f.0030* and *f.0033*.** Names of RdRP identified in Nam Co are highlighted in red. Color blocks from left to right denote the classes and orders of Nam Co RdRPs’ top hits in BLASTp, with gradient blocks and their labels indicating the families of Nam Co RdRPs’ top hits in BLASTp. The names of RdRPs correspond to specific ORF sequences in Table S4.

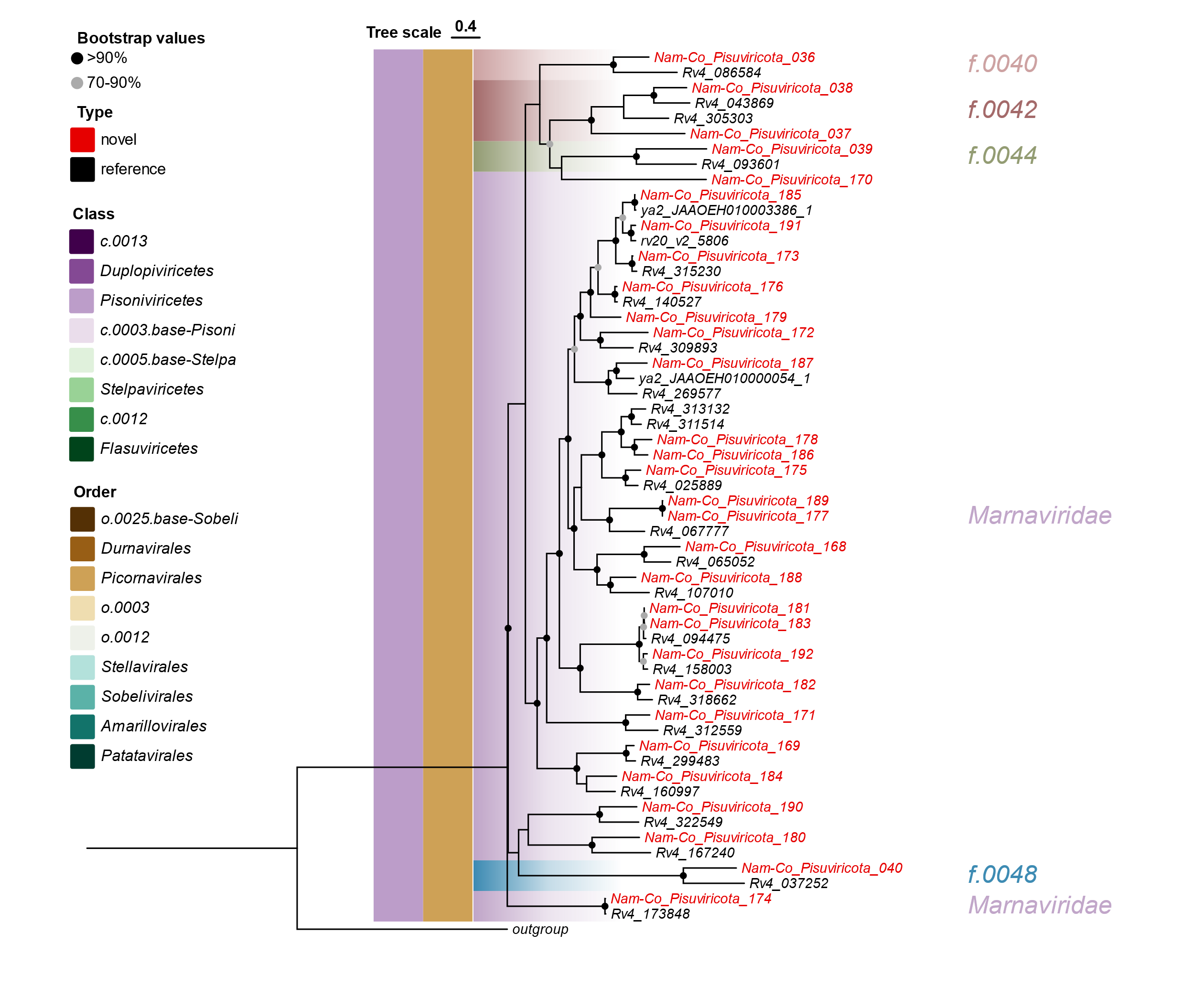

**Figure S10. The subtrees Ⅱ for *Pisuviricota* including families *Marnaviridae*, *f.0040*, *f.0042*, *f.0044* and *f.0048*.** Names of RdRP identified in Nam Co are highlighted in red. Color blocks from left to right denote the classes and orders of Nam Co RdRPs’ top hits in BLASTp, with gradient blocks and their labels indicating the families of Nam Co RdRPs’ top hits in BLASTp. The names of RdRPs correspond to specific ORF sequences in Table S4.

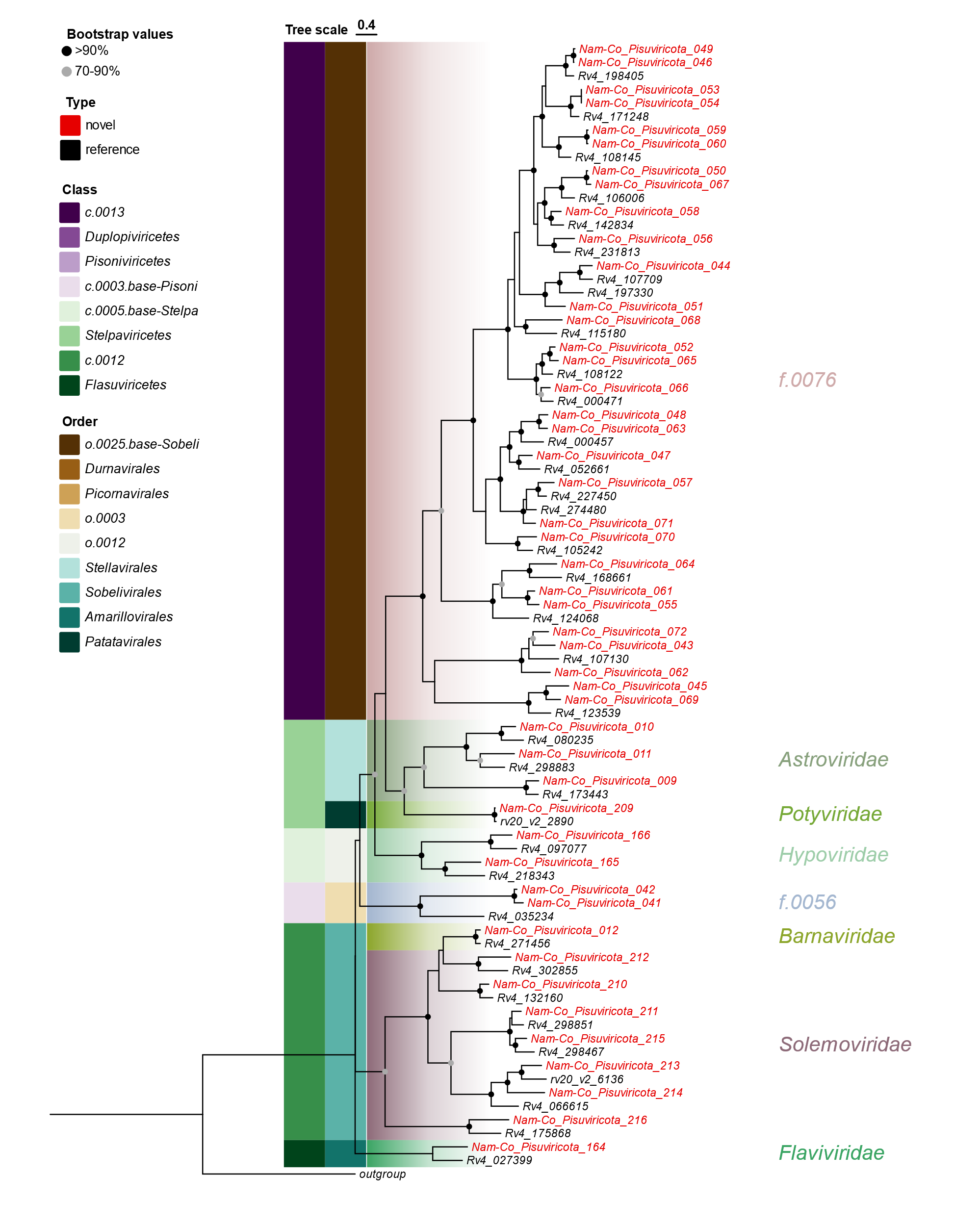

**Figure S11. The subtree Ⅲ for *Pisuviricota* including families *Astroviridae*, *Barnaviridae*, *Flaviviridae*, *Hypoviridae*, *Potyviridae*, *Solemoviridae*, *f.0056* and *f.0076*.** Names of RdRP identified in Nam Co are highlighted in red. Color blocks from left to right denote the classes and orders of Nam Co RdRPs’ top hits in BLASTp, with gradient blocks and their labels indicating the families of Nam Co RdRPs’ top hits in BLASTp. The names of RdRPs correspond to specific ORF sequences in Table S4.

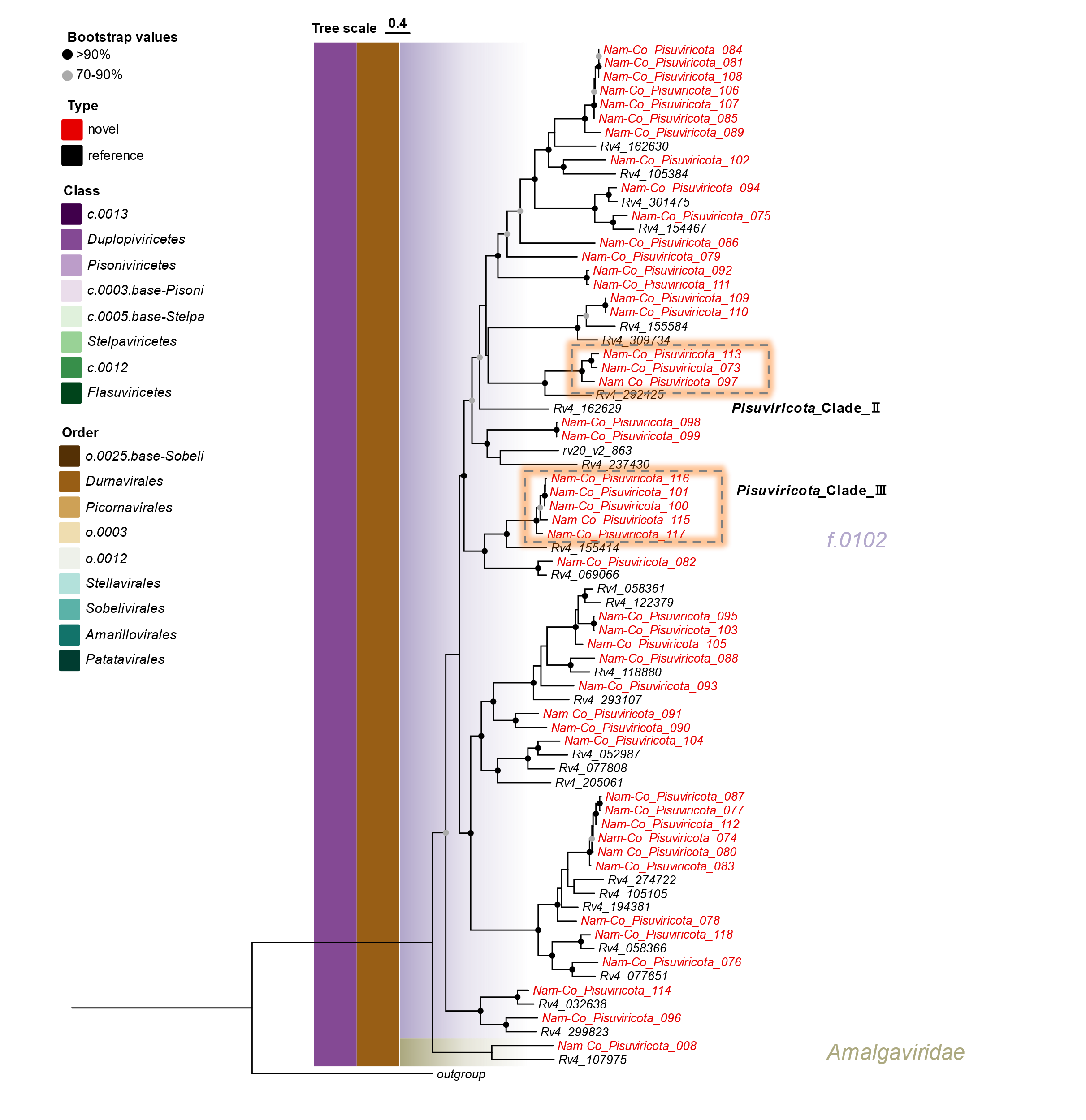

**Figure S12. The subtree Ⅳ for *Pisuviricota* including families *Amalgaviridae* and *f.0102*.** Names of RdRP identified in Nam Co are highlighted in red. Color blocks from left to right denote the classes and orders of Nam Co RdRPs’ top hits in BLASTp, with gradient blocks and their labels indicating the families of Nam Co RdRPs’ top hits in BLASTp. The names of RdRPs correspond to specific ORF sequences in Table S4.

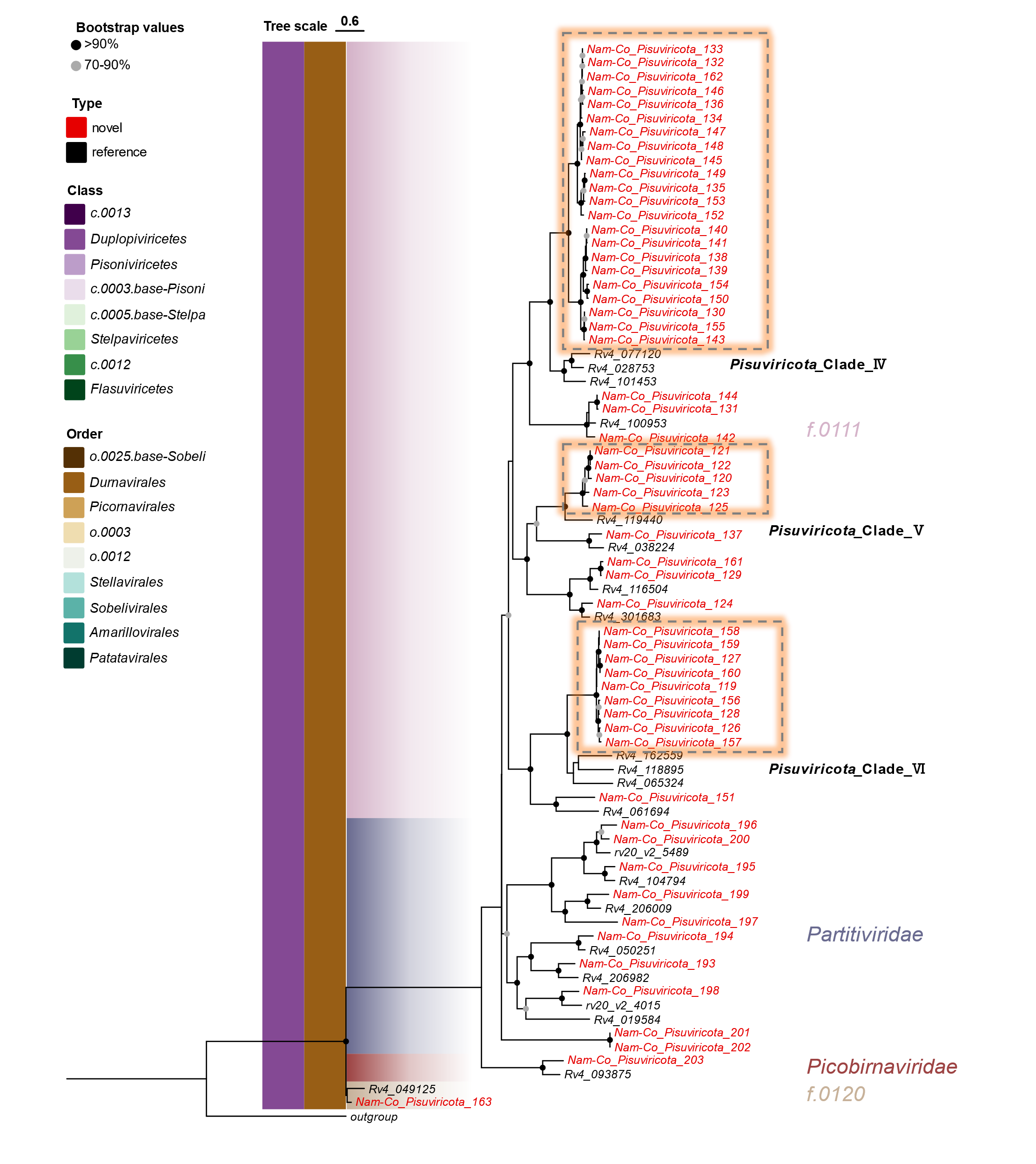

**Figure S13. The subtree Ⅴ for *Pisuviricota* including families *Partitiviridae*, *Picobirnaviridae*, *f.0111* and *f.0120*.** Names of RdRP identified in Nam Co are highlighted in red. Color blocks from left to right denote the classes and orders of Nam Co RdRPs’ top hits in BLASTp, with gradient blocks and their labels indicating the families of Nam Co RdRPs’ top hits in BLASTp. The names of RdRPs correspond to specific ORF sequences in Table S4.

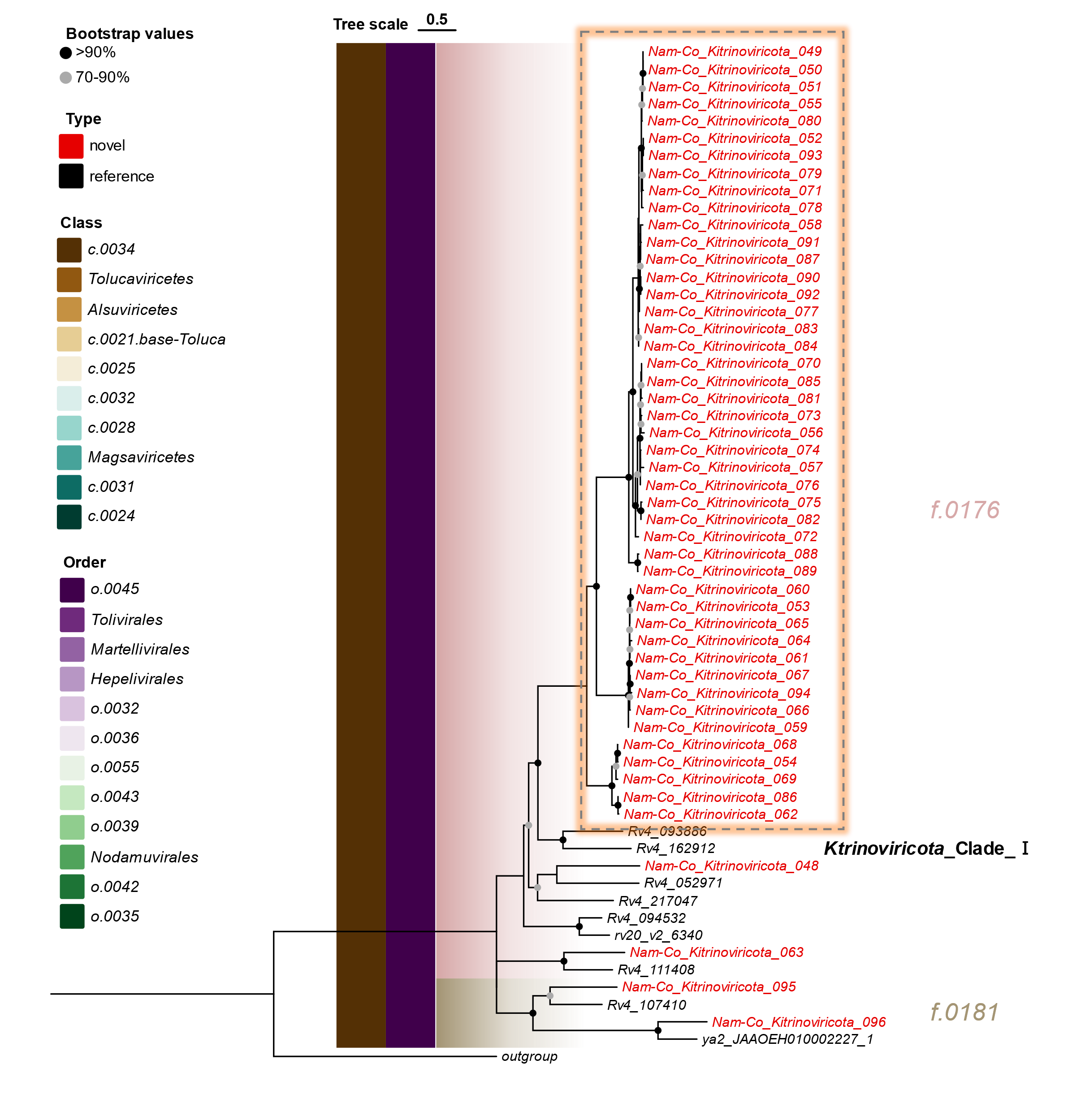

**Figure S14. The subtree Ⅰ for *Kitrinoviricota* including families *f.0176* and *f.0181*.** Names of RdRP identified in Nam Co are highlighted in red. Color blocks from left to right denote the classes and orders of Nam Co RdRPs’ top hits in BLASTp, with gradient blocks and their labels indicating the families of Nam Co RdRPs’ top hits in BLASTp. The names of RdRPs correspond to specific ORF sequences in Table S4.

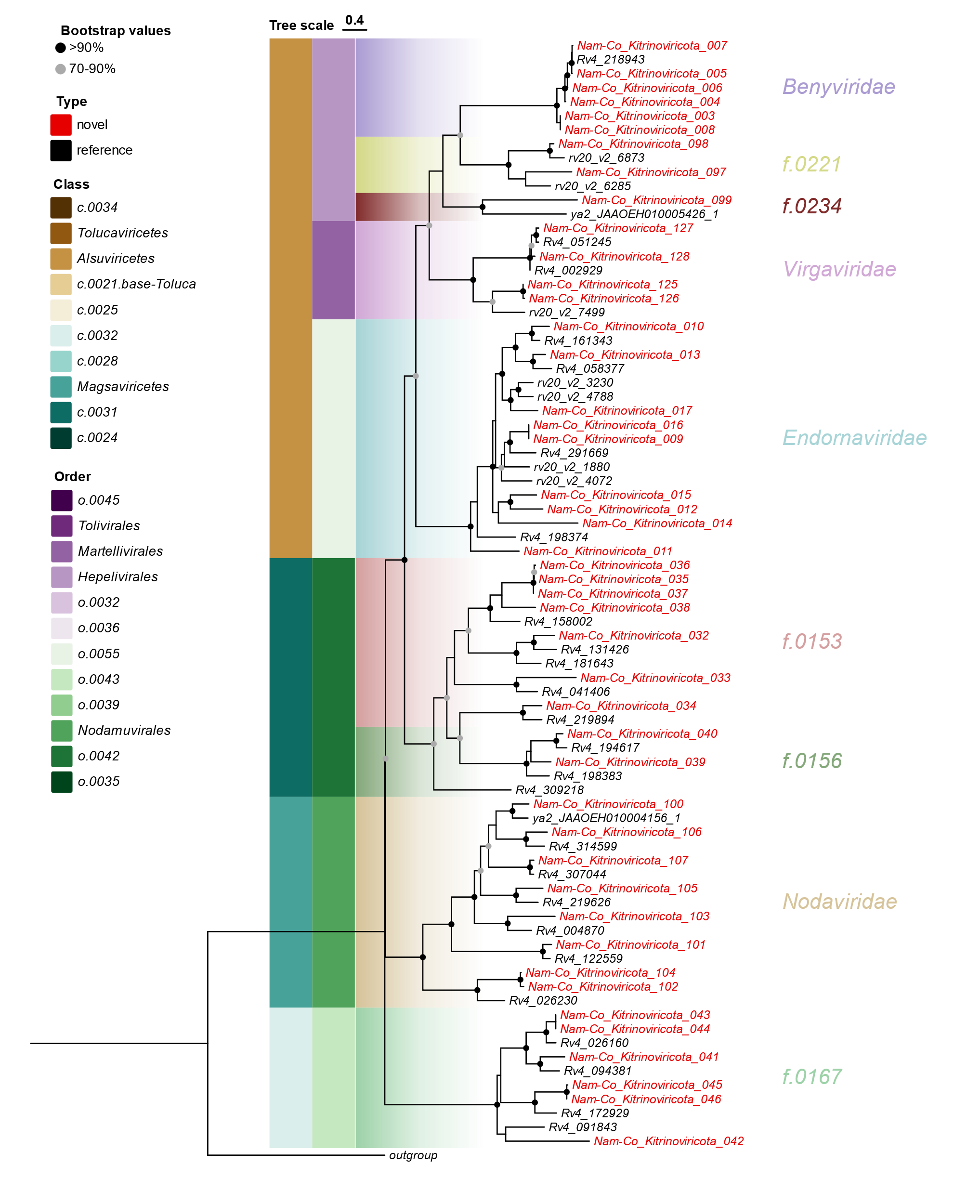

**Figure S15. The subtree Ⅱ for *Kitrinoviricota* including families *Benyviridae*, *Endornaviridae*, *Nodaviridae*, *Virgaviridae*, *f.0153*, *f.0156*, *f.0167*, *f.0221* and *f.0234*.** Names of RdRP identified in Nam Co are highlighted in red. Color blocks from left to right denote the classes and orders of Nam Co RdRPs’ top hits in BLASTp, with gradient blocks and their labels indicating the families of Nam Co RdRPs’ top hits in BLASTp. The names of RdRPs correspond to specific ORF sequences in Table S4.

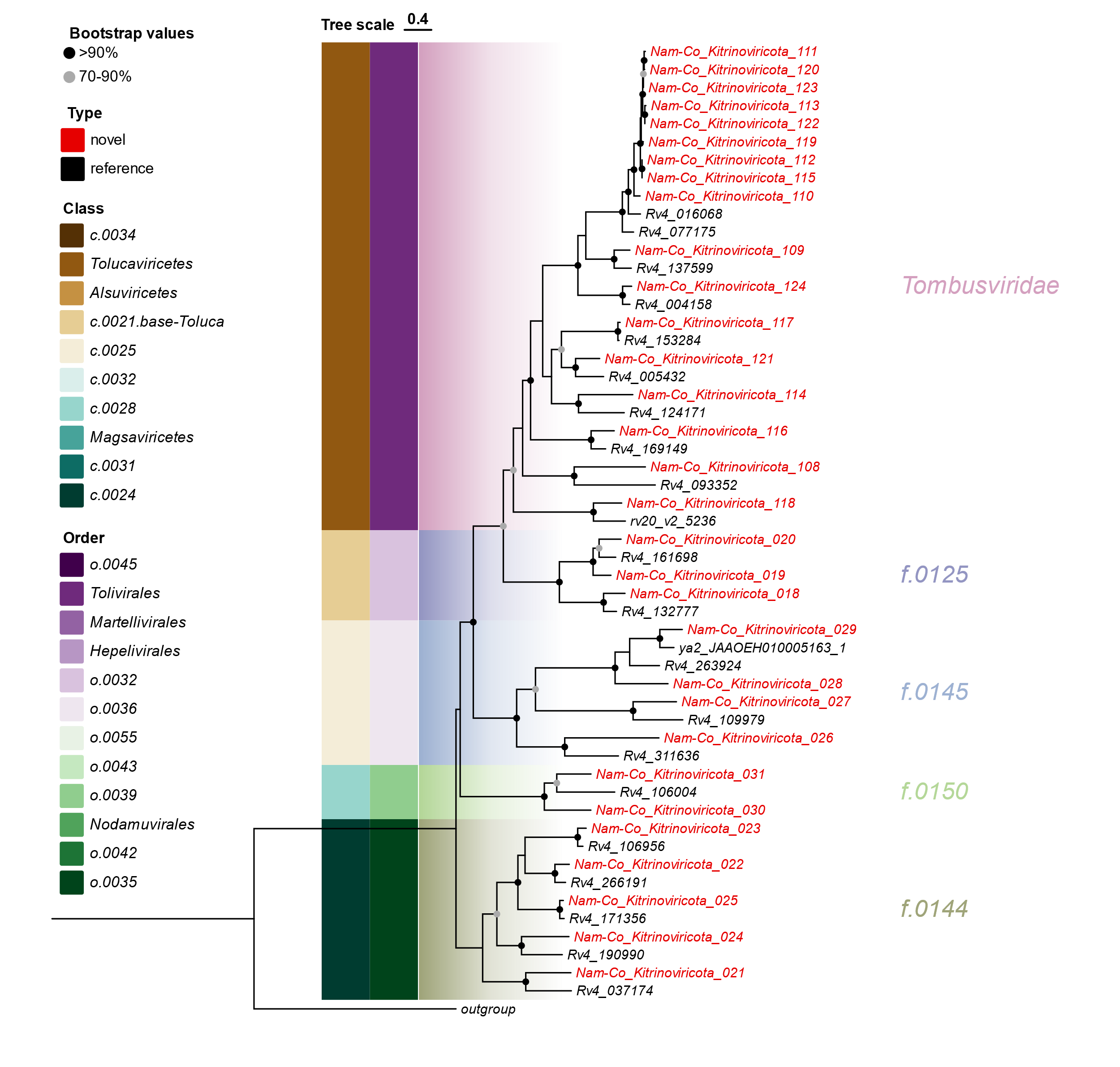

**Figure S16. The subtree Ⅲ for *Kitrinoviricota* including families *Tombusviridae*, *f.0125*, *f.0144*, *f.0145* and *f.0150*.** Names of RdRP identified in Nam Co are highlighted in red. Color blocks from left to right denote the classes and orders of Nam Co RdRPs’ top hits in BLASTp, with gradient blocks and their labels indicating the families of Nam Co RdRPs’ top hits in BLASTp. The names of RdRPs correspond to specific ORF sequences in Table S4.

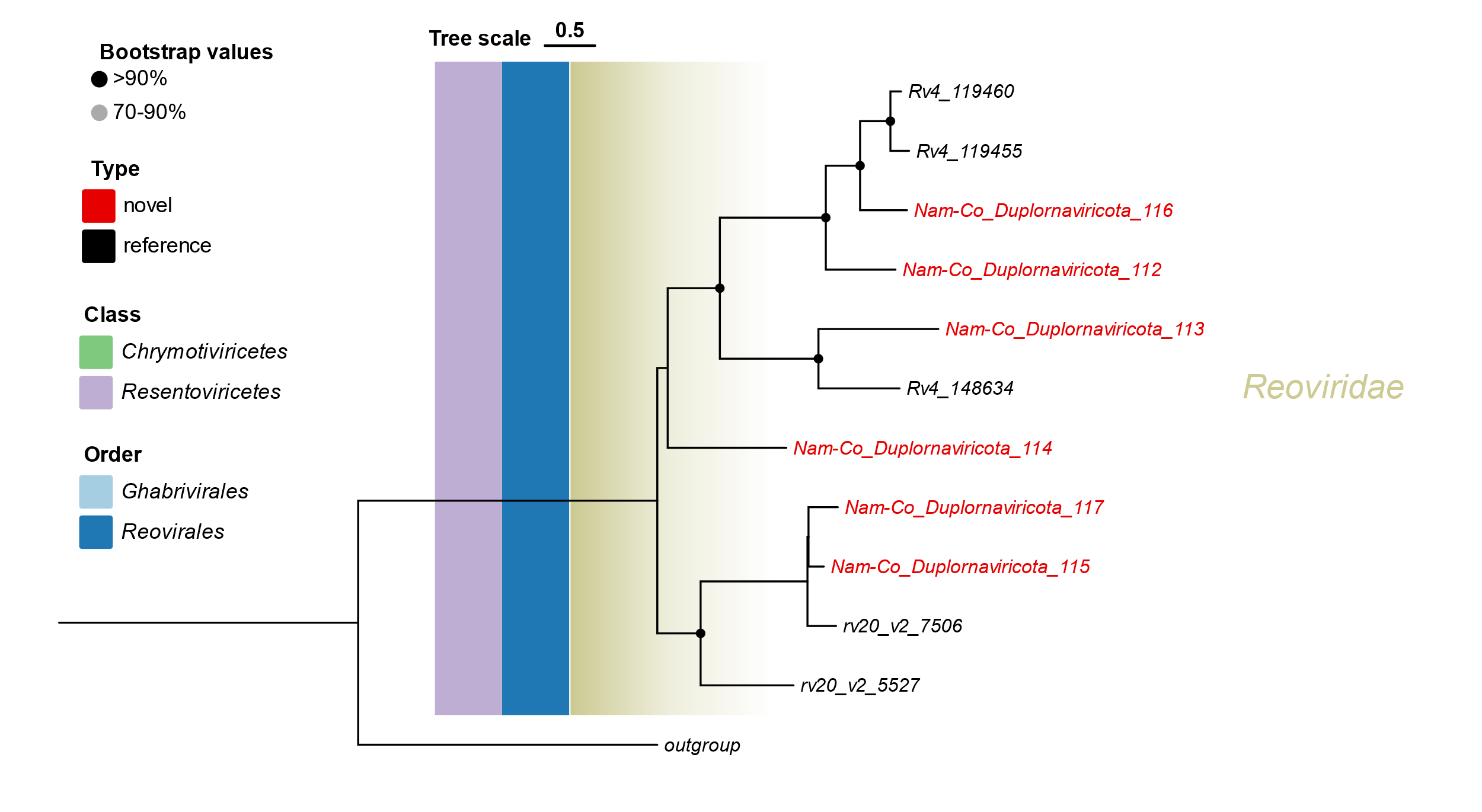

**Figure S17. The subtree Ⅰ for *Duplornaviricota* including family *Reoviridae***. Names of RdRP identified in Nam Co are highlighted in red. Color blocks from left to right denote the classes and orders of Nam Co RdRPs’ top hits in BLASTp, with gradient blocks and their labels indicating the families of Nam Co RdRPs’ top hits in BLASTp. The names of RdRPs correspond to specific ORF sequences in Table S4.

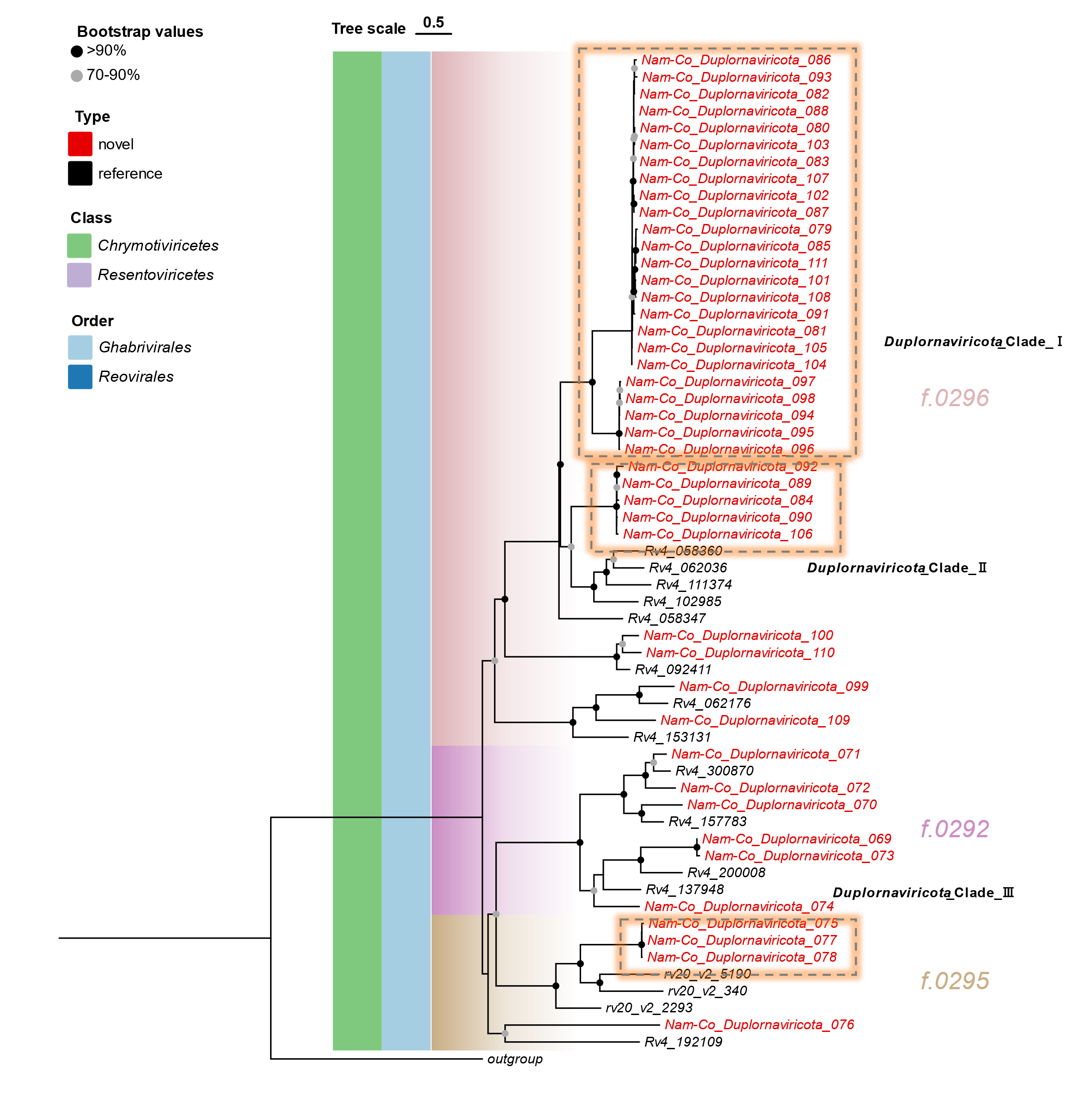

**Figure S18. The subtree Ⅱ for *Duplornaviricota* including families *f.0292*, *f.0295* and *f.0296*.** Names of RdRP identified in Nam Co are highlighted in red. Color blocks from left to right denote the classes and orders of Nam Co RdRPs’ top hits in BLASTp, with gradient blocks and their labels indicating the families of Nam Co RdRPs’ top hits in BLASTp. The names of RdRPs correspond to specific ORF sequences in Table S4.

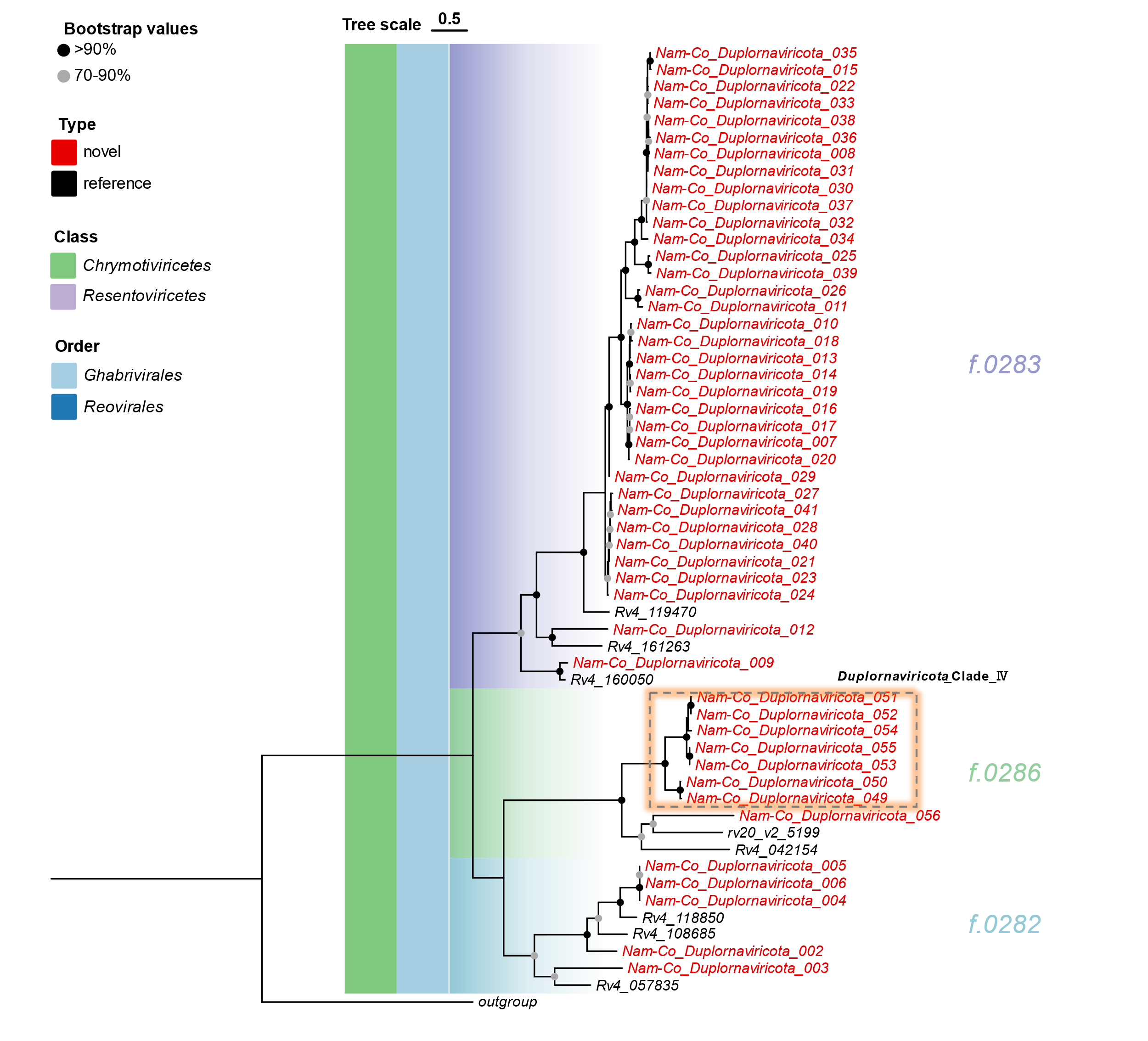

**Figure S19. The subtree Ⅲ for *Duplornaviricota* including families *f.0282*, *f.0283* and *f.0286*.** Names of RdRP identified in Nam Co are highlighted in red. Color blocks from left to right denote the classes and orders of Nam Co RdRPs’ top hits in BLASTp, with gradient blocks and their labels indicating the families of Nam Co RdRPs’ top hits in BLASTp. The names of RdRPs correspond to specific ORF sequences in Table S4.

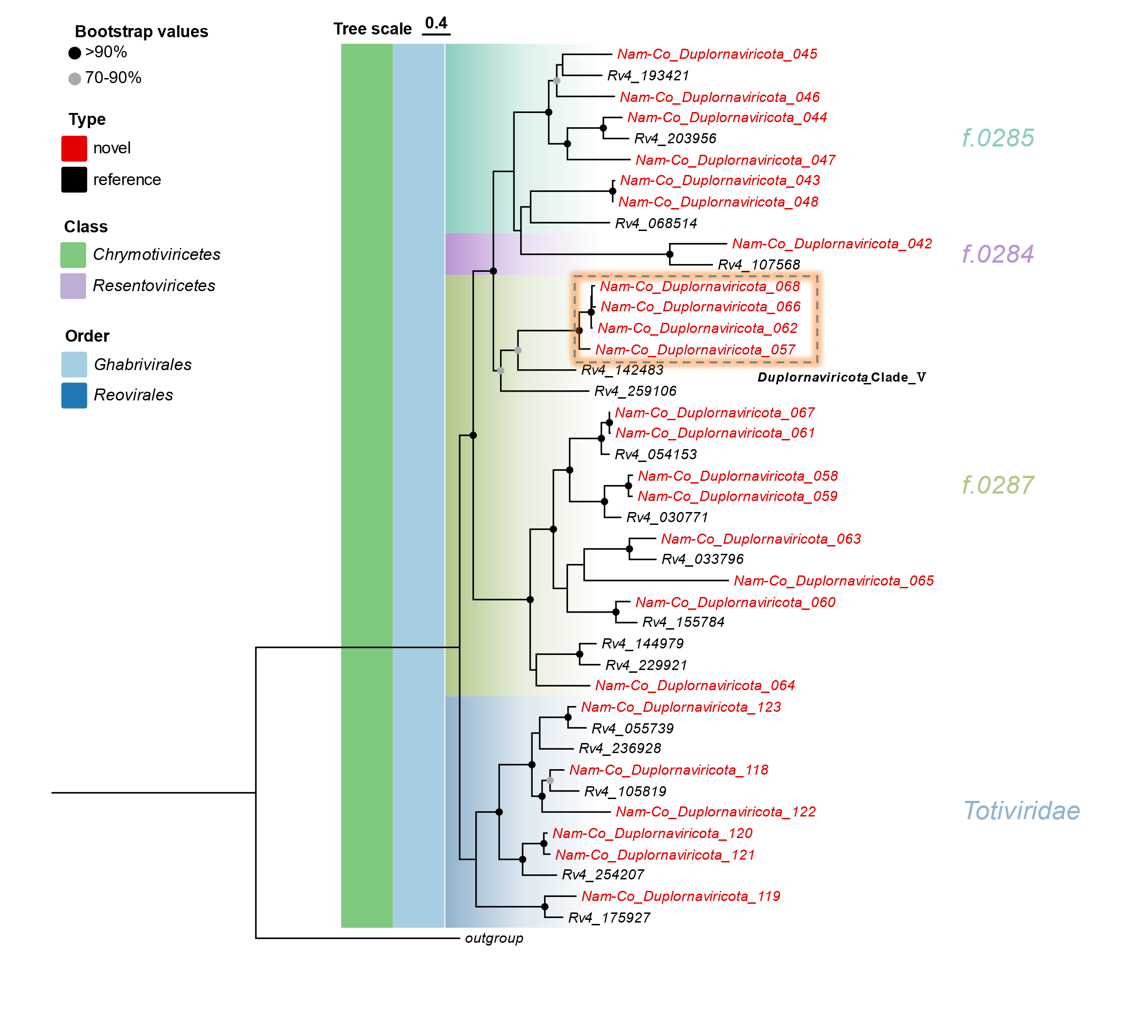

**Figure S20. The subtree Ⅳ for *Duplornaviricota* including families *Totiviridae*, *f.0284*, *f.0285* and *f.0287*.** Names of RdRP identified in Nam Co are highlighted in red. Color blocks from left to right denote the classes and orders of Nam Co RdRPs’ top hits in BLASTp, with gradient blocks and their labels indicating the families of Nam Co RdRPs’ top hits in BLASTp. The names of RdRPs correspond to specific ORF sequences in Table S4.

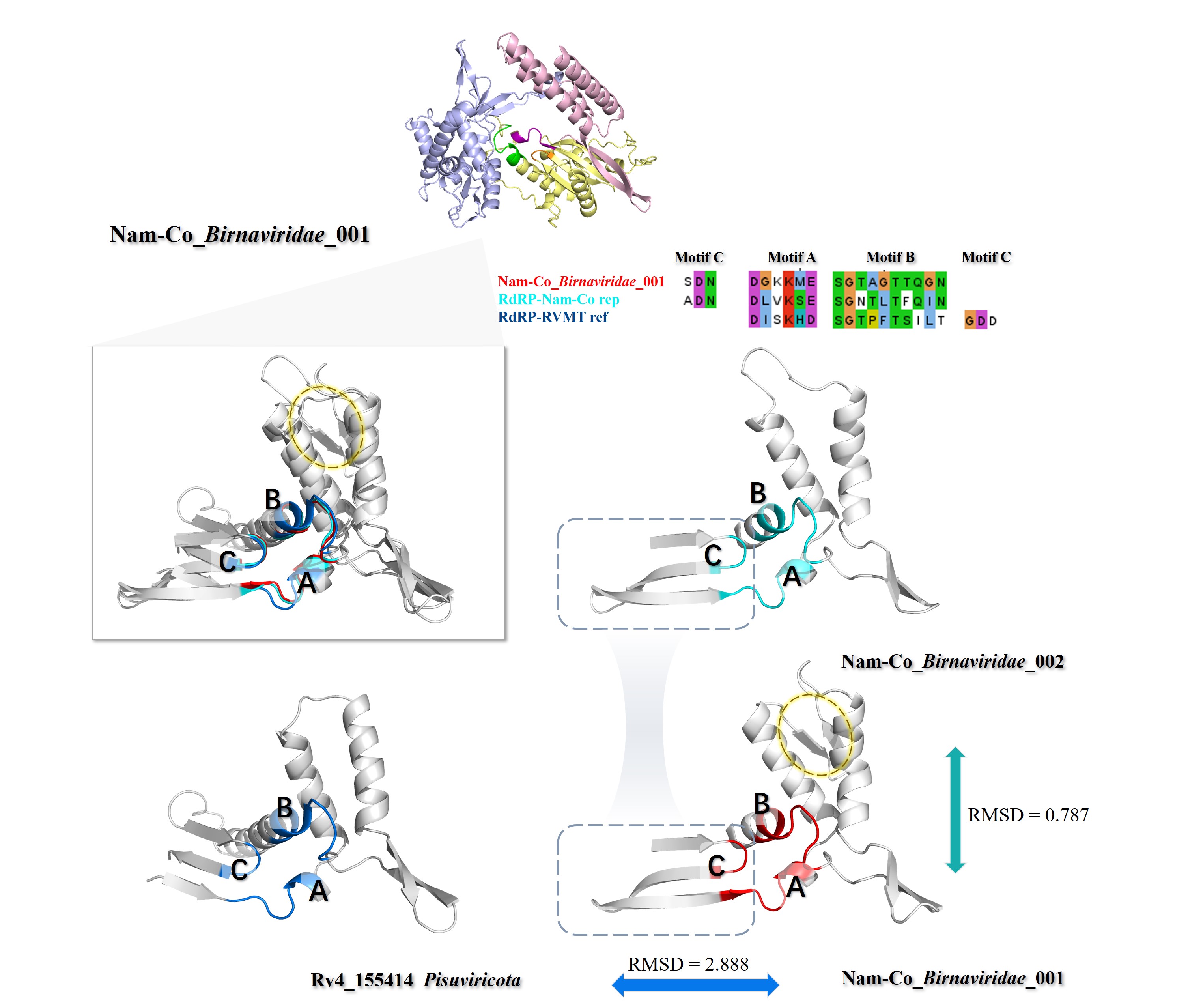

**Figure S21. The catalytic center portion of the "palm" region in the structure of Nam-Co_*Birnaviridae*_RdRP_001.** The amino acid composition of motifs for the compared structures are listed in the actual arrangement sequence, shown below the schematic diagram. Motifs from Nam-Co_*Birnaviridae*_RdRP_001 are colored in red, motifs from the RVMT reference structure are colored in dark blue, and motifs of the Nam Co representative are colored in light blue. The RMSD values between structures are indicated by arrows of the same color as the referential structures. Blue dashed boxes highlight regions of similarity between the two structures, while yellow dashed circles denote structural features unique to Nam-Co_*Birnaviridae*_RdRP_001.

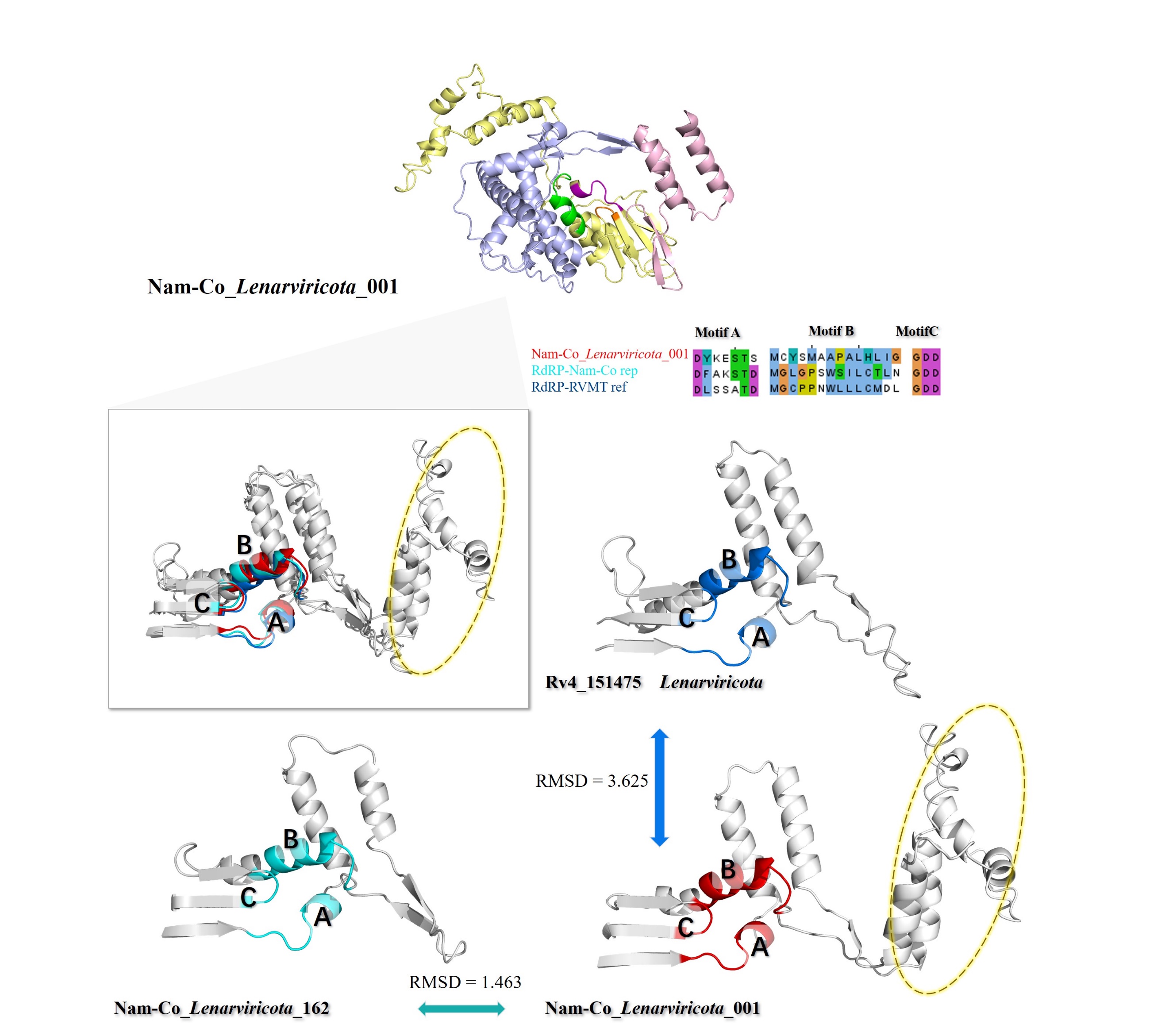

**Figure S22. The catalytic center portion of the "palm" region in the structure of Nam-Co_*Lenarviricota*_RdRP_001.** The amino acid composition of motifs for the compared structures are listed in the actual arrangement sequence, shown below the schematic diagram. Motifs from Nam-Co_*Lenarviricota*_RdRP_001 are colored in red, motifs from the RVMT reference structure are colored in dark blue, and motifs of the Nam Co representative are colored in light blue. The RMSD values between structures are indicated by arrows of the same color as the referential structures. Yellow dashed circles denote structural features unique to Nam-Co_*Lenarviricota*_RdRP_001.

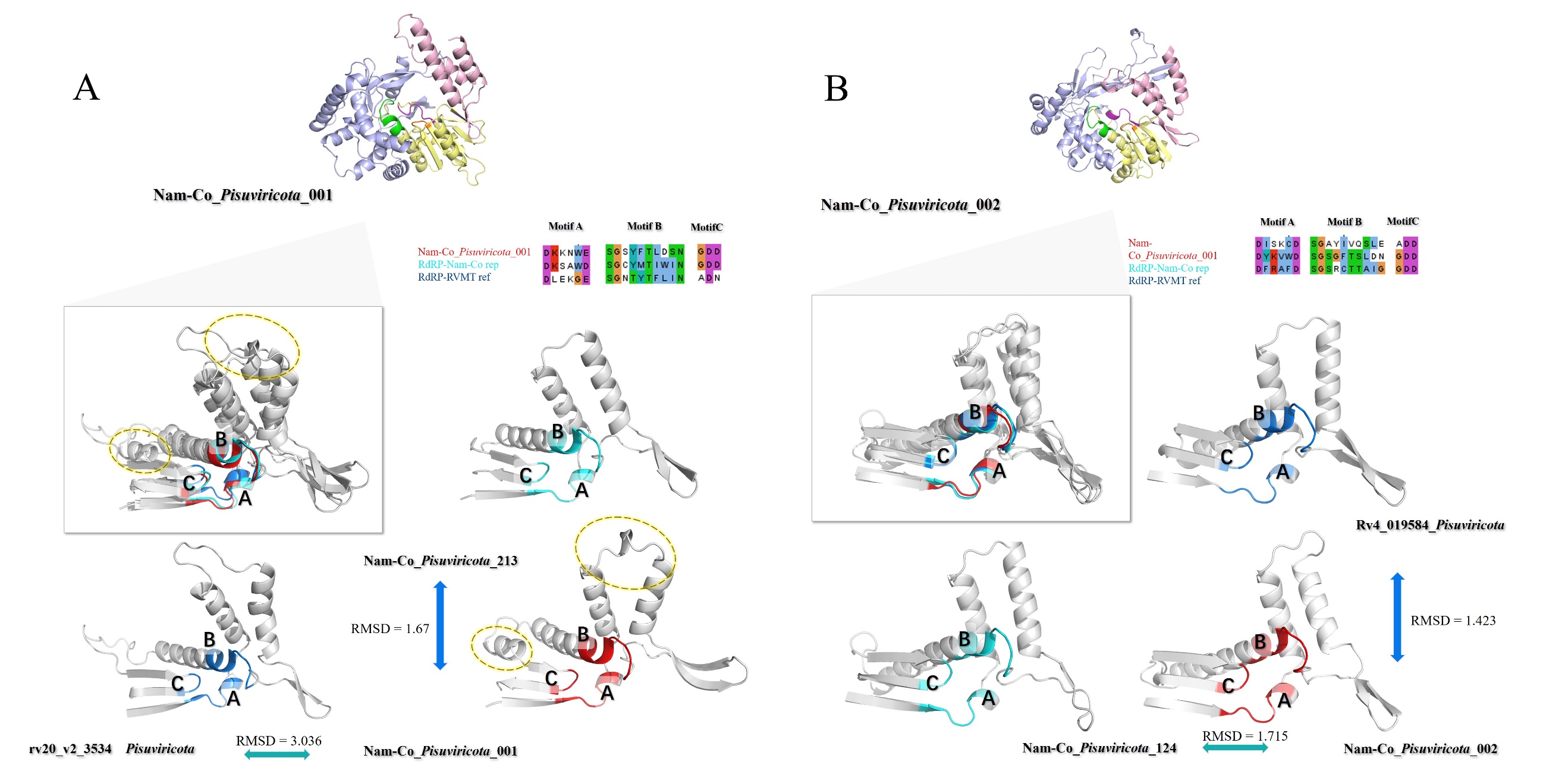

**Figure S23. The catalytic center portion of the "palm" region in the structure of Nam-Co_*Pisuviricota*_RdRP_001-002.** The amino acid composition of motifs for the compared structures are listed in the actual arrangement sequence, shown below the schematic diagram. Motifs from Nam-Co_*Pisuviricota*_RdRP_001-002 are colored in red, motifs from the RVMT reference structures are colored in dark blue, and motifs of the Nam Co representatives are colored in light blue. The RMSD values between structures are indicated by arrows of the same color as the referential structures. Yellow dashed circles denote structural features unique to Nam-Co_*Pisuviricota*_RdRP_001.

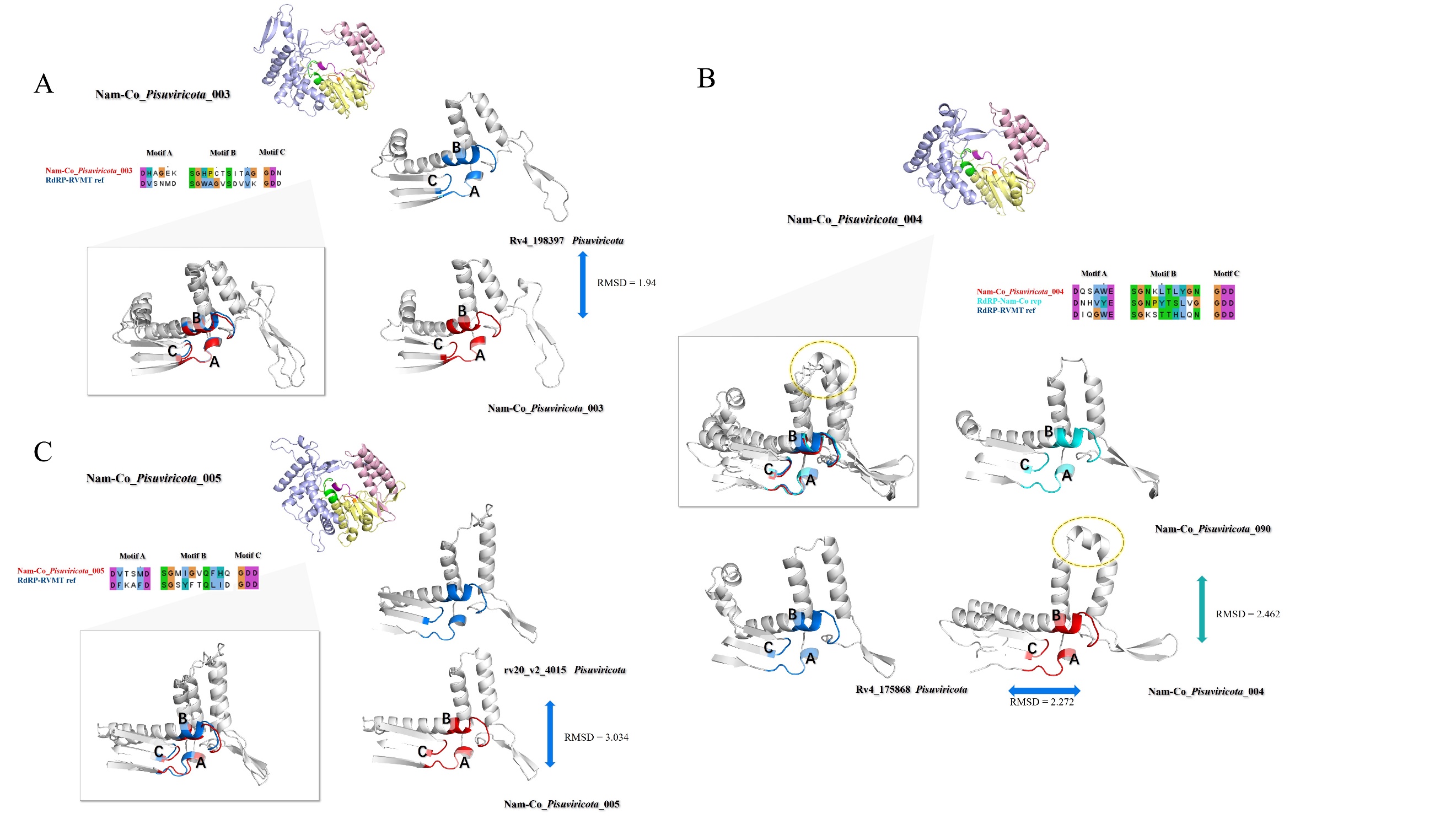

**Figure S24. The catalytic center portion of the "palm" region in the structure of Nam-Co_*Pisuviricota*_RdRP_003-005.** The amino acid composition of motifs for the compared structures are listed in the actual arrangement sequence, shown below the schematic diagram. Motifs from Nam-Co_*Pisuviricota*_RdRP_003-005 are colored in red, motifs from the RVMT reference structures are colored in dark blue, and motifs of the Nam Co representatives are colored in light blue. The RMSD values between structures are indicated by arrows of the same color as the referential structures. Yellow dashed circles denote structural features unique to Nam-Co_*Pisuviricota*_RdRP_004.

**Figure S25. The catalytic center portion of the "palm" region in the structure of Nam-Co_*Pisuviricota*_RdRP_006-007.** The amino acid composition of motifs for the compared structures are listed in the actual arrangement sequence, shown below the schematic diagram. Motifs from Nam-Co_*Pisuviricota*_RdRP_006-007 are colored in red, motifs from the RVMT reference structures are colored in dark blue, and motifs of the Nam Co representatives are colored in light blue. The RMSD values between structures are indicated by arrows of the same color as the referential structures. Yellow dashed circles denote structural features unique to Nam-Co_*Pisuviricota*_RdRP_007.

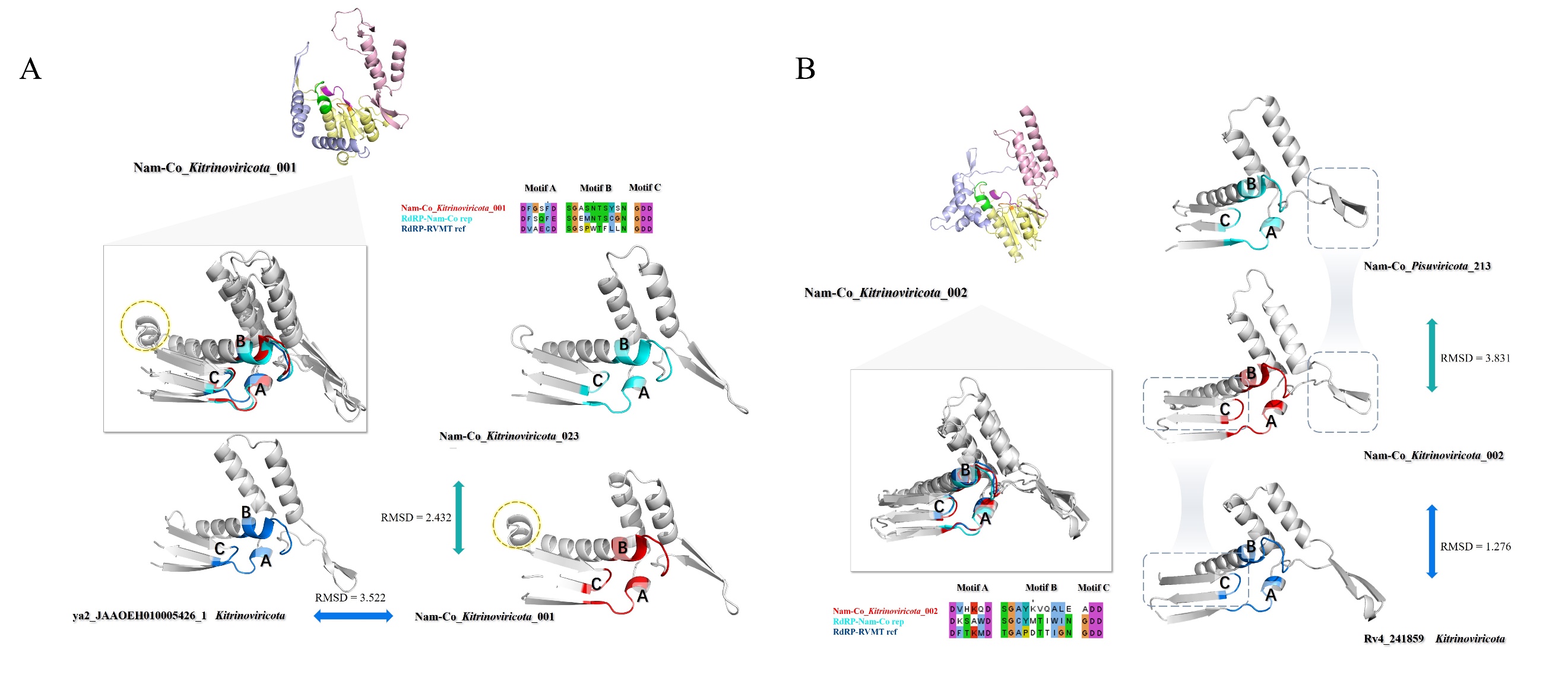

**Figure S26. The catalytic center portion of the "palm" region in the structure of Nam-Co_*Kitrinoviricota*_RdRP_001-002.** The amino acid composition of motifs for the compared structures are listed in the actual arrangement sequence, shown below the schematic diagram. Motifs from Nam-Co_*Kitrinoviricota*_RdRP_001-002 are colored in red, motifs from the RVMT reference structures are colored in dark blue, and motifs of the Nam Co representatives are colored in light blue. The RMSD values between structures are indicated by arrows of the same color as the referential structure. Blue dashed boxes highlight regions of similarity between the two structures of Nam-Co_*Kitrinoviricota*_RdRP_002, while yellow dashed circles denote structural features unique to Nam-Co_*Kitrinoviricota*_RdRP_001.

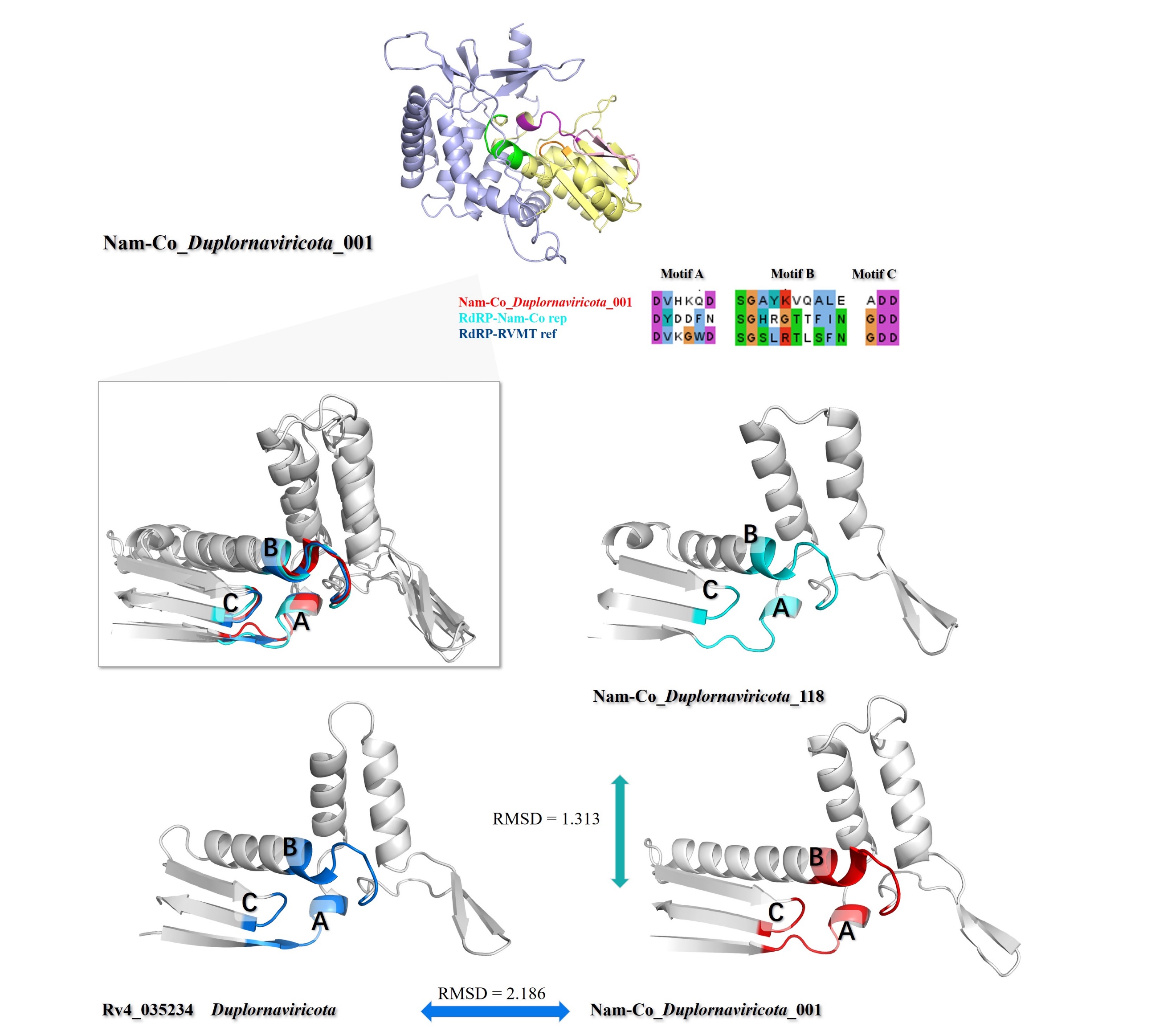

**Figure S27. The catalytic center portion of the "palm" region in the structure of Nam-Co_*Duplornaviricota*_RdRP_001.** The amino acid composition of motifs for the compared structures are listed in the actual arrangement sequence, shown below the schematic diagram. Motifs from Nam-Co_*Duplornaviricota*_RdRP_001 are colored in red, motifs from the RVMT reference structure are colored in dark blue, and motifs of the Nam Co representative are colored in light blue. The RMSD values between structures are indicated by arrows of the same color as the referential structures.

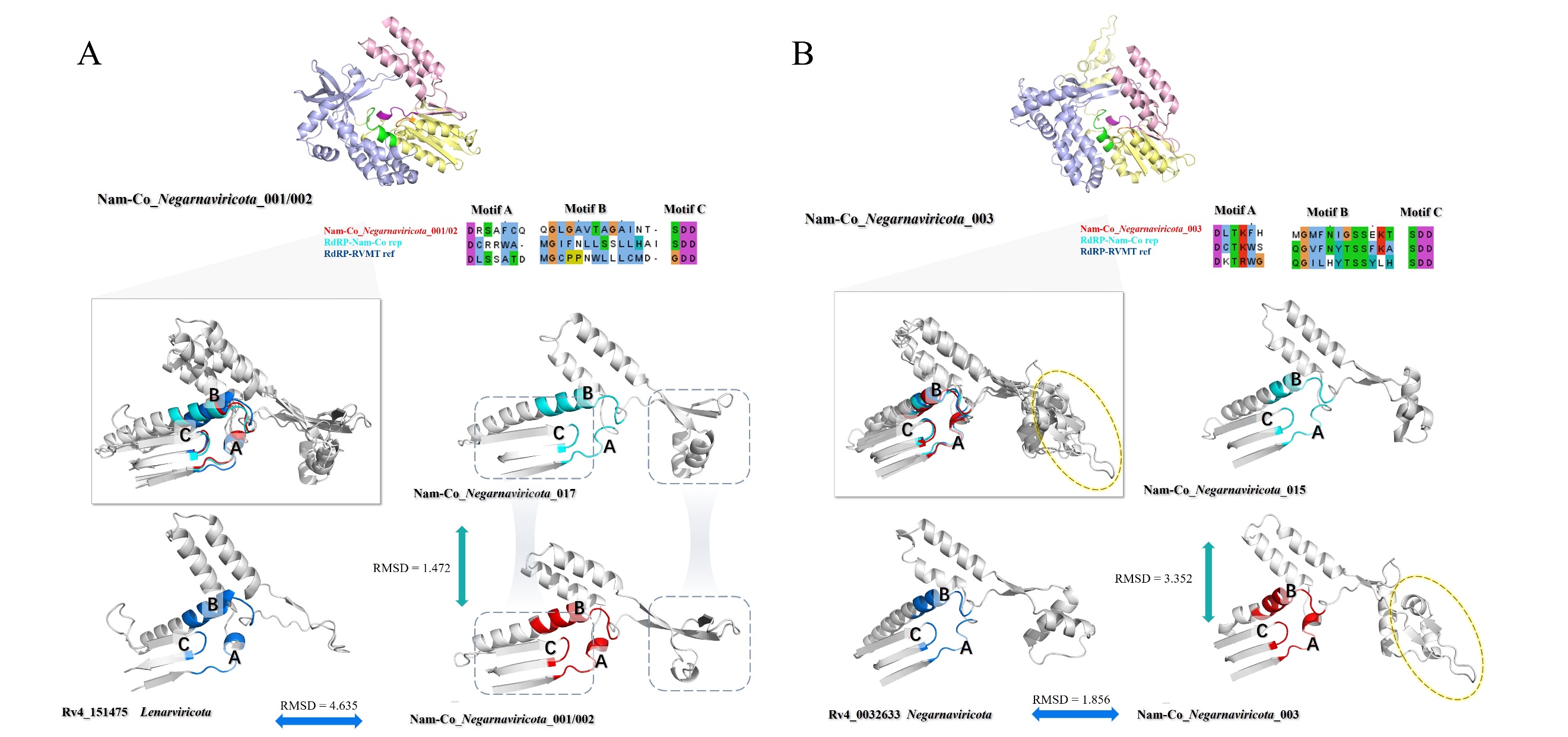

**Figure S28. The catalytic center portion of the "palm" region in the structure of Nam-Co_*Negarnaviricota*_RdRP_001-003.** Since the RMSD between Nam-Co_*Negarnaviricota*_001 and Nam-Co_*Negarnaviricota*_002 <0.5, the structure of Nam-Co_*Negarnaviricota*_001 is representative of both. The amino acid composition of motifs for the compared structures are listed in the actual arrangement sequence, shown below the schematic diagram. Motifs from Nam-Co_ *Negarnaviricota*_RdRP_001-003 are colored in red, motifs from the RVMT reference structures are colored in dark blue, and motifs of the Nam Co representatives are colored in light blue. The RMSD values between structures are indicated by arrows of the same color as the referential structures. Blue dashed boxes highlight regions of similarity between the two structures of Nam-Co_*Negarnaviricota*_001/002, while yellow dashed circles denote structural features unique to Nam-Co_ *Negarnaviricota*_RdRP_003.

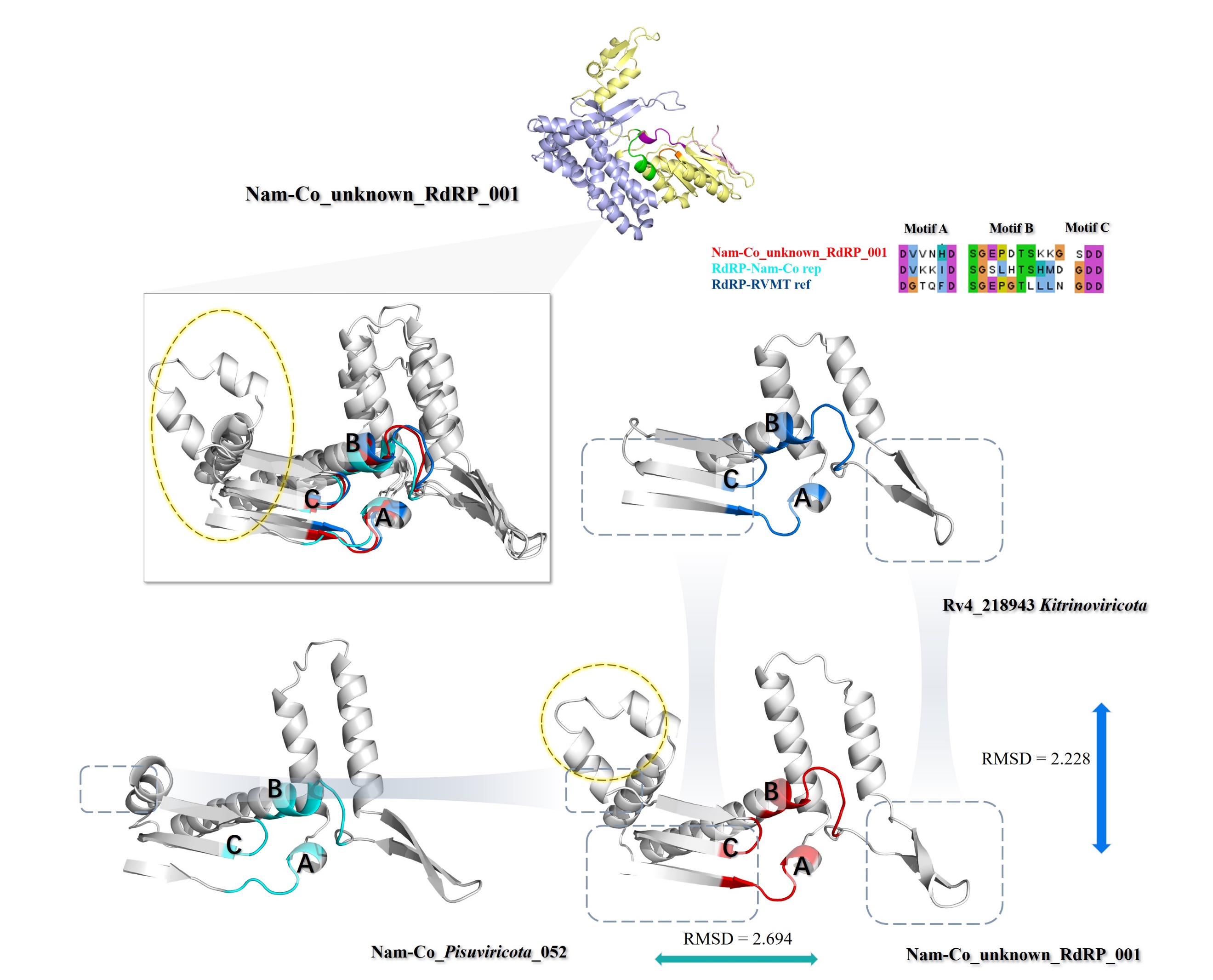

**Figure S29. The catalytic center portion of the "palm" region in the structure of Nam-Co_unknown_RdRP_001.** The amino acid composition of motifs for the compared structures are listed in the actual arrangement sequence, shown below the schematic diagram. Motifs from Nam-Co_unknown_RdRP_001 are colored in red, motifs from the RVMT reference structure are colored in dark blue, and motifs of the Nam Co representative are colored in light blue. The RMSD values between structures are indicated by arrows of the same color as the referential structures. Blue dashed boxes highlight regions of similarity between the two structures, while yellow dashed circles denote structural features unique to Nam-Co_unknown_RdRP_001.

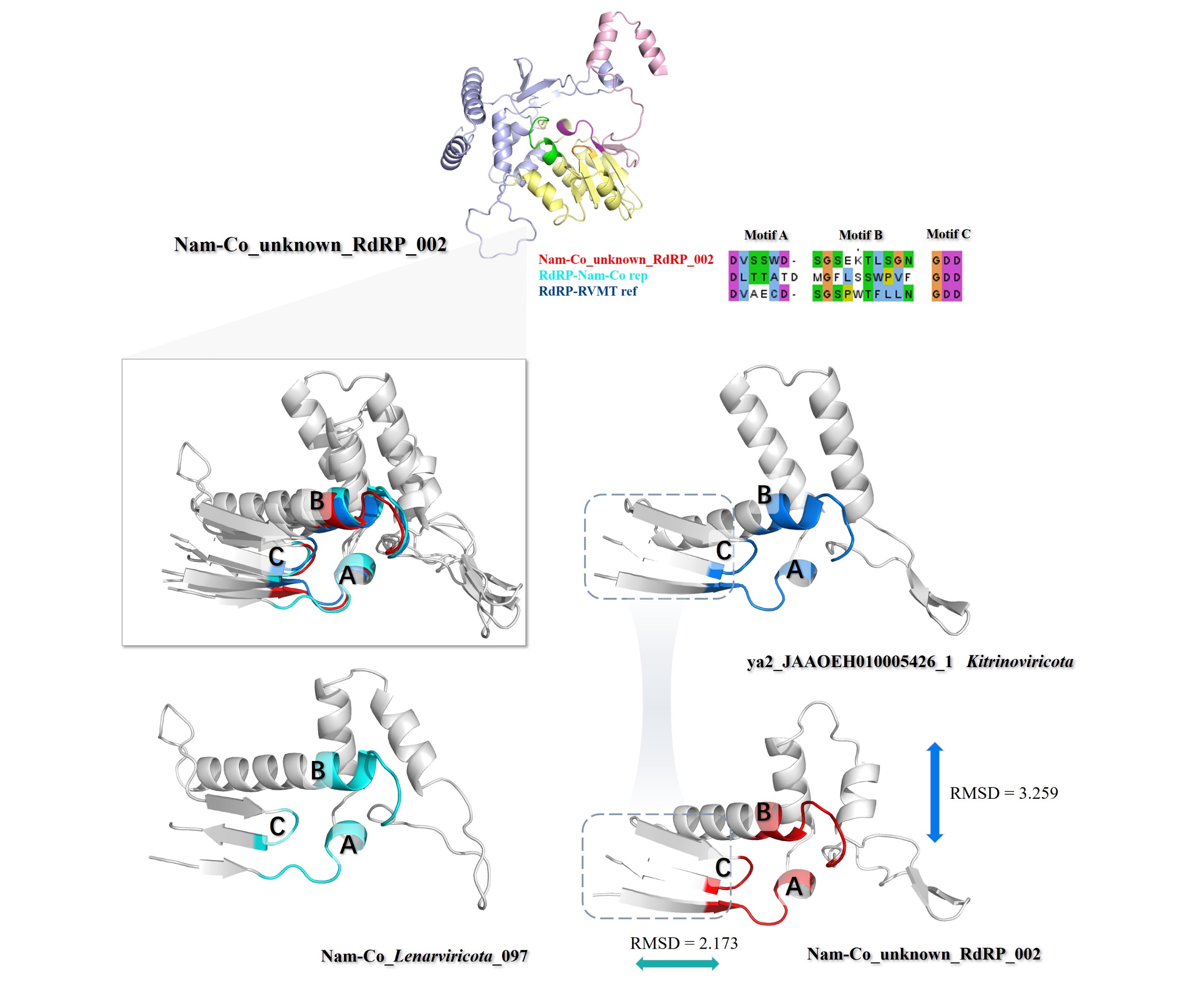

**Figure S30. The catalytic center portion of the "palm" region in the structure of Nam-Co_unknown_RdRP_002.** The amino acid composition of motifs for the compared structures are listed in the actual arrangement sequence, shown below the schematic diagram. Motifs from Nam-Co_unknown_RdRP_002 are colored in red, motifs from the RVMT reference structure are colored in dark blue, and motifs of the Nam Co representative are colored in light blue. The RMSD values between structures are indicated by arrows of the same color as the referential structures. Blue dashed boxes highlight regions of similarity between the two structures of Nam-Co_unknown_RdRP_002.

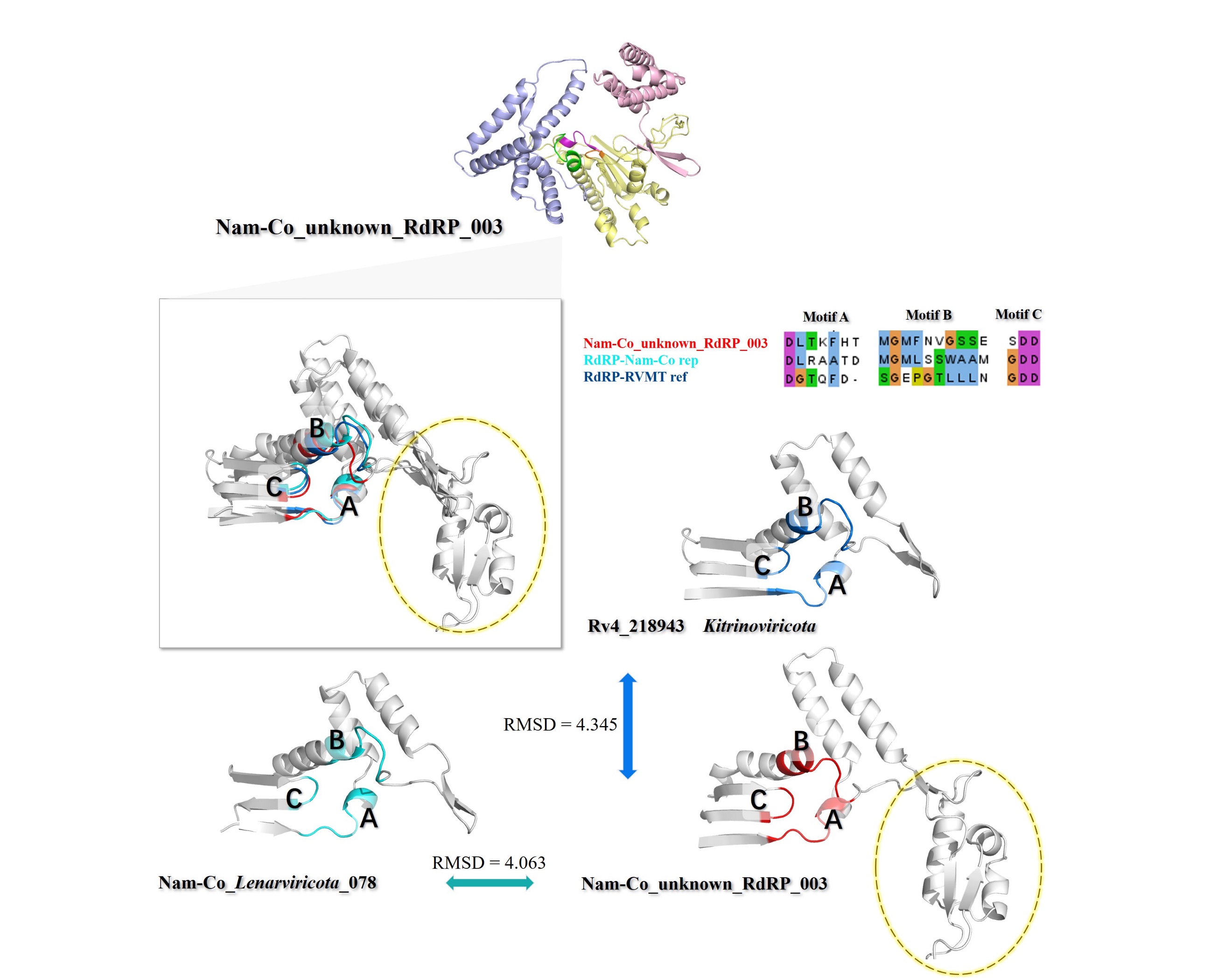

**Figure S31. The catalytic center portion of the "palm" region in the structure of Nam-Co_unknown_RdRP_003.** The amino acid composition of motifs for the compared structures are listed in the actual arrangement sequence, shown below the schematic diagram. Motifs from Nam-Co_unknown_RdRP_003 are colored in red, motifs from the RVMT reference structure are colored in dark blue, and motifs of the Nam Co representative are colored in light blue. The RMSD values between structures are indicated by arrows of the same color as the referential structures. Blue dashed boxes highlight regions of similarity between the two structures, while yellow dashed circles denote structural features unique to Nam-Co_unknown_RdRP_003.

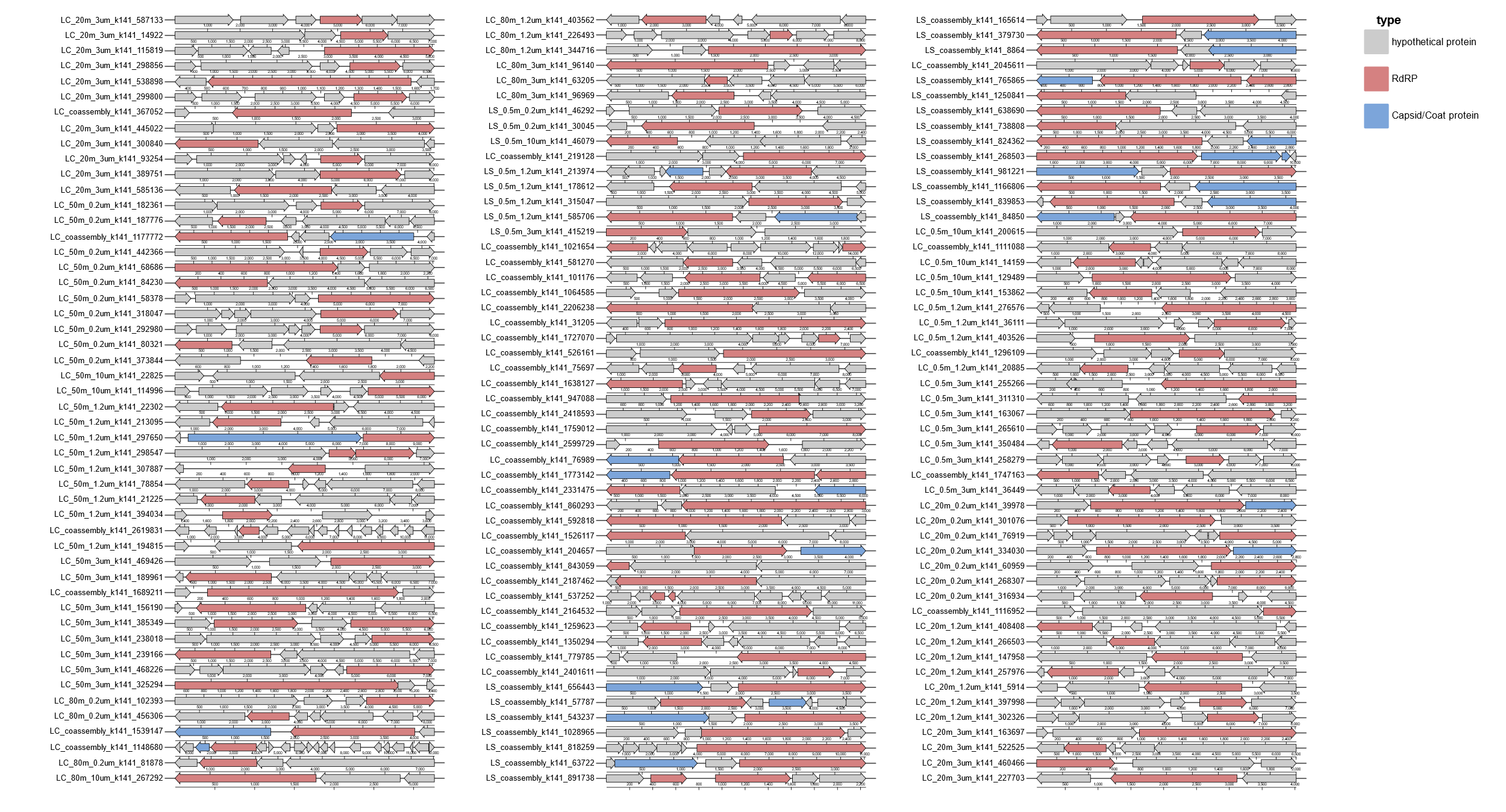

**Figure S32. Most genes in the genomes of RNA viruses in Lake Nam Co encode unknown proteins except for RdRPs and capsid proteins.** ORFs encoding RdRPs are colored red, those for capsid proteins are colored blue, and those for unknown proteins are colored grey.

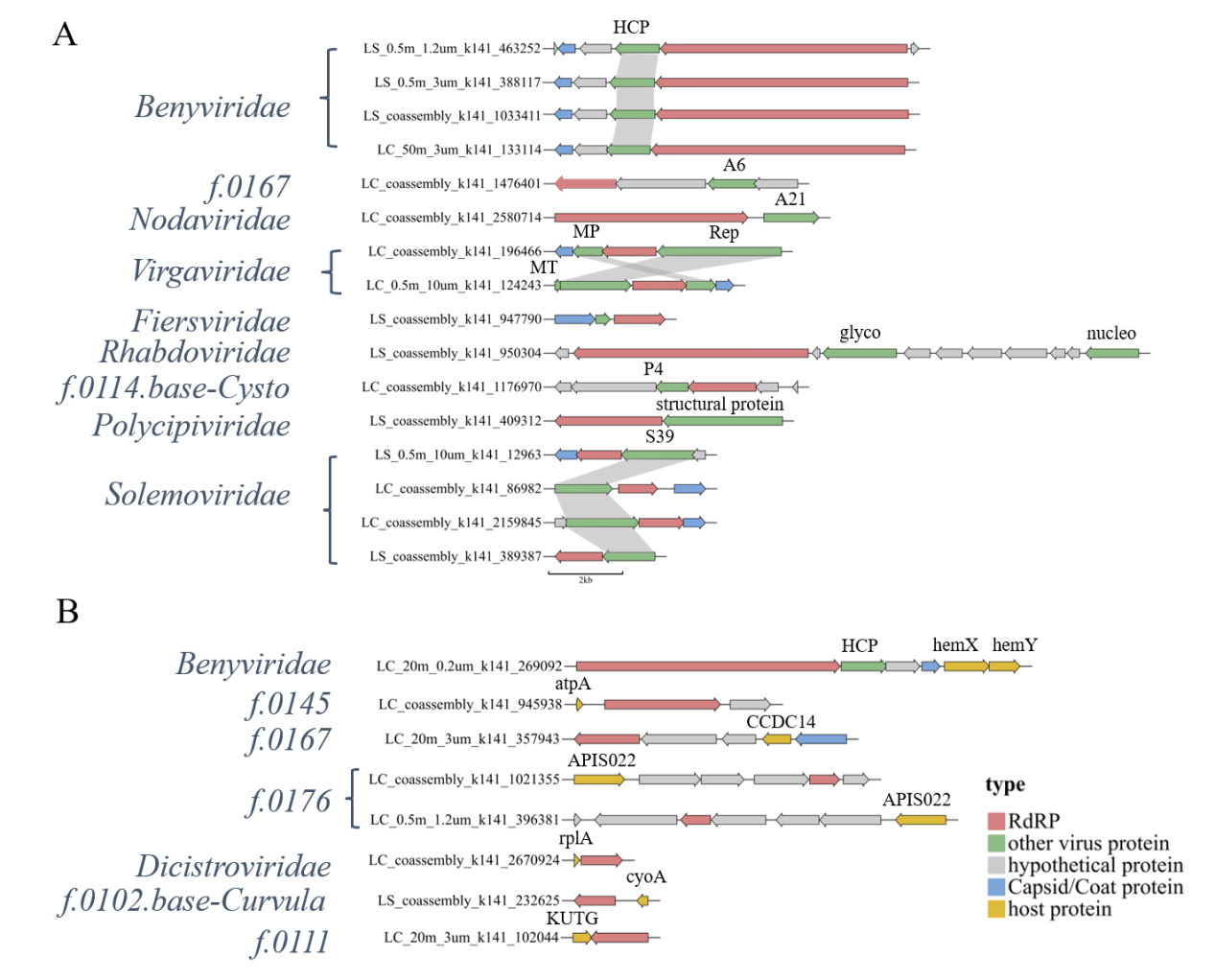

**Figure S33. The RNA virus genome from Nam Co, containing viral and host genes, labeled with the respective family affiliations on the left.** (A) Genome containing genes encoding viral proteins other than capsid proteins and RdRPs. (B) Genome containing host genes. ORFs encoding RdRPs are colored red, those for capsid proteins are colored blue, and those for unknown proteins are colored grey. ORFs encoding other viral proteins are colored blue, and ORFs encoding host proteins are colored yellow.

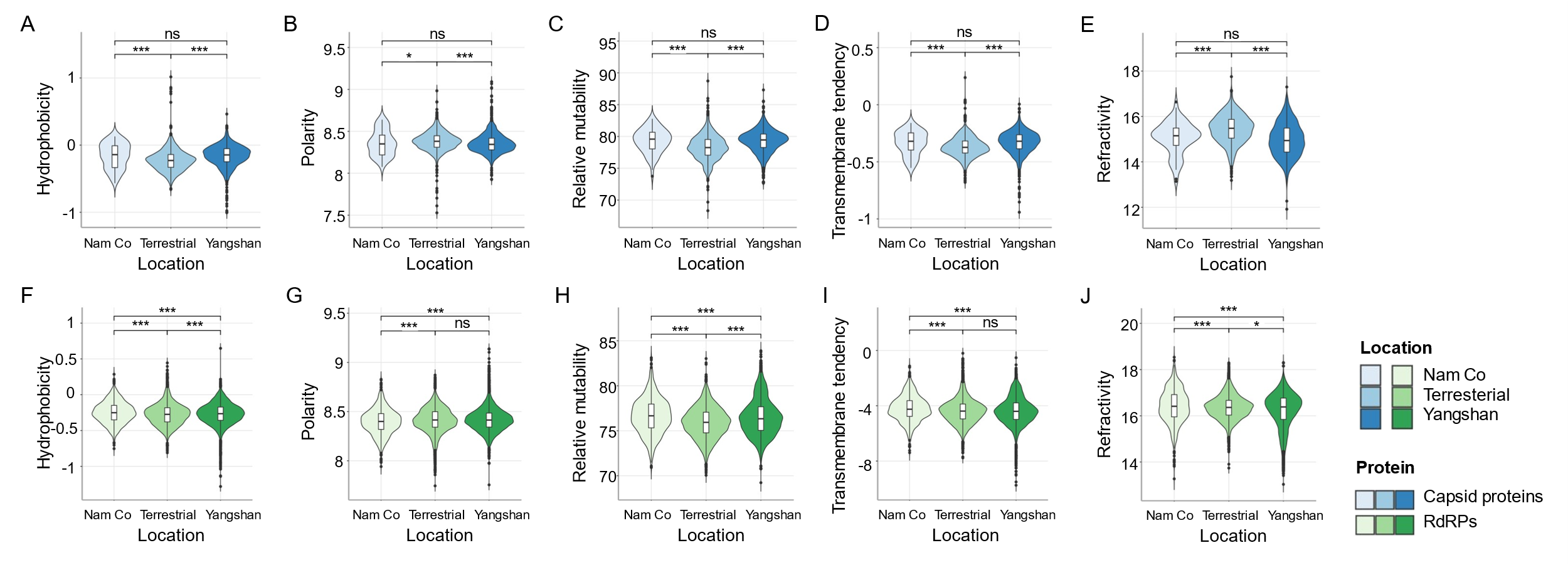

**Figure S34. Comparison of physicochemical property indices of RNA virus proteins from Nam Co Lake with those from terrestrial and estuarine environments.** (A) The comparisons of hydrophobicity of capsid proteins among different habitats. (B) The comparisons of polarity of capsid proteins among different habitats. (C) The comparisons of relative mutability of capsid proteins among different habitats. (D) The comparisons of transmembrane tendency of capsid proteins among different habitats. (E) The comparisons of refractivity of capsid proteins among different habitats. (F) The comparisons of hydrophobicity of RdRPs among different habitats. (G) The comparisons of polarity of RdRPs among different habitats. (H) The comparisons of relative mutability of RdRPs among different habitats. (I) The comparisons of transmembrane tendency of RdRPs among different habitats. (J) The comparisons of refractivity of RdRPs among different habitats.

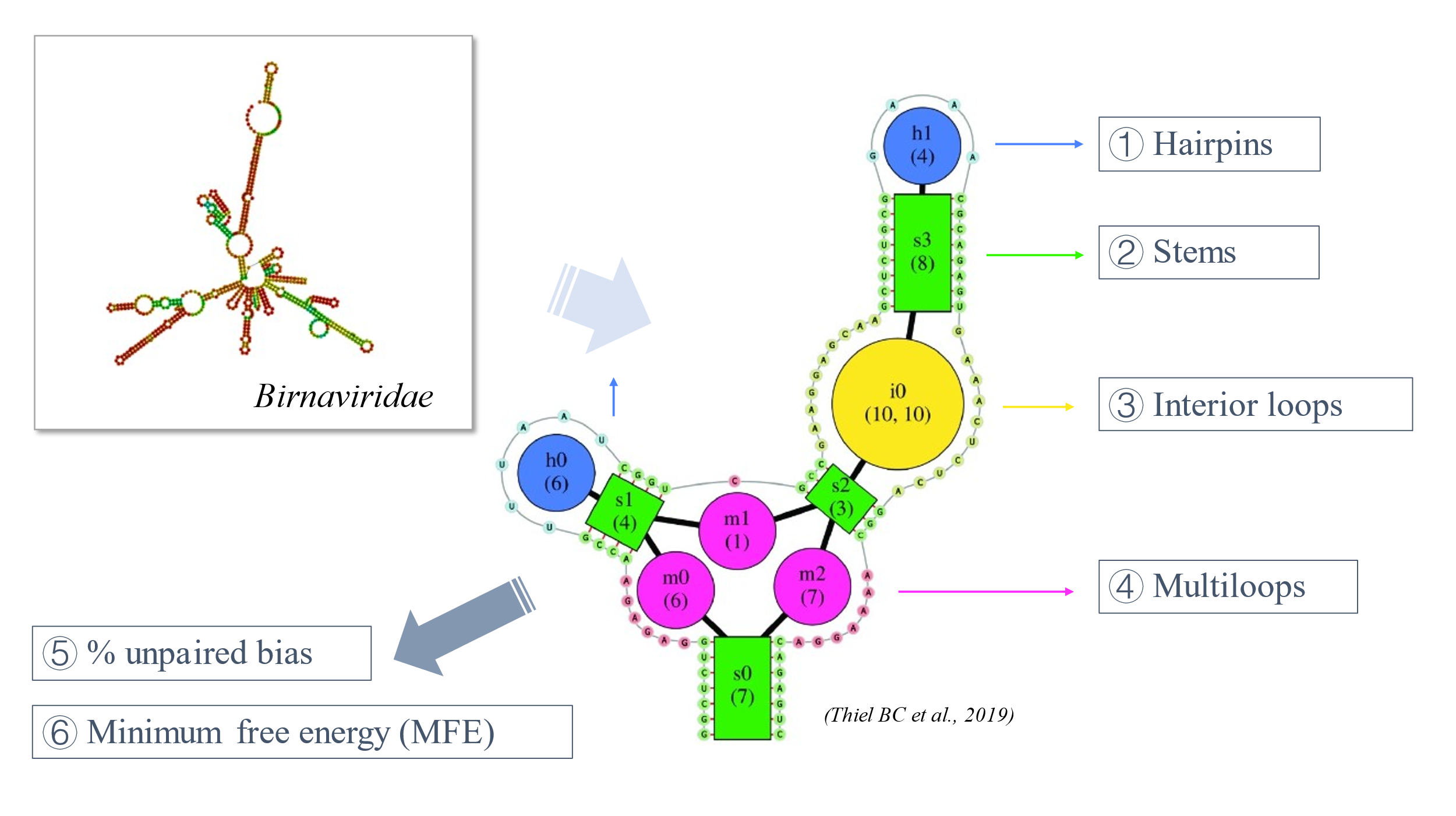

**Figure S35. Illustration of RNA secondary structure features.** The upper left corner depicts the secondary structure of a virus of *Birnaviridae*. The schematic is adapted from Thiel BC et al., 2019. In the RNA secondary structure, hairpins are colored in blue, stems in green, interior loops in yellow, and multiloops in pink. Features that are not elements include unpaired bases and minimum free energy.

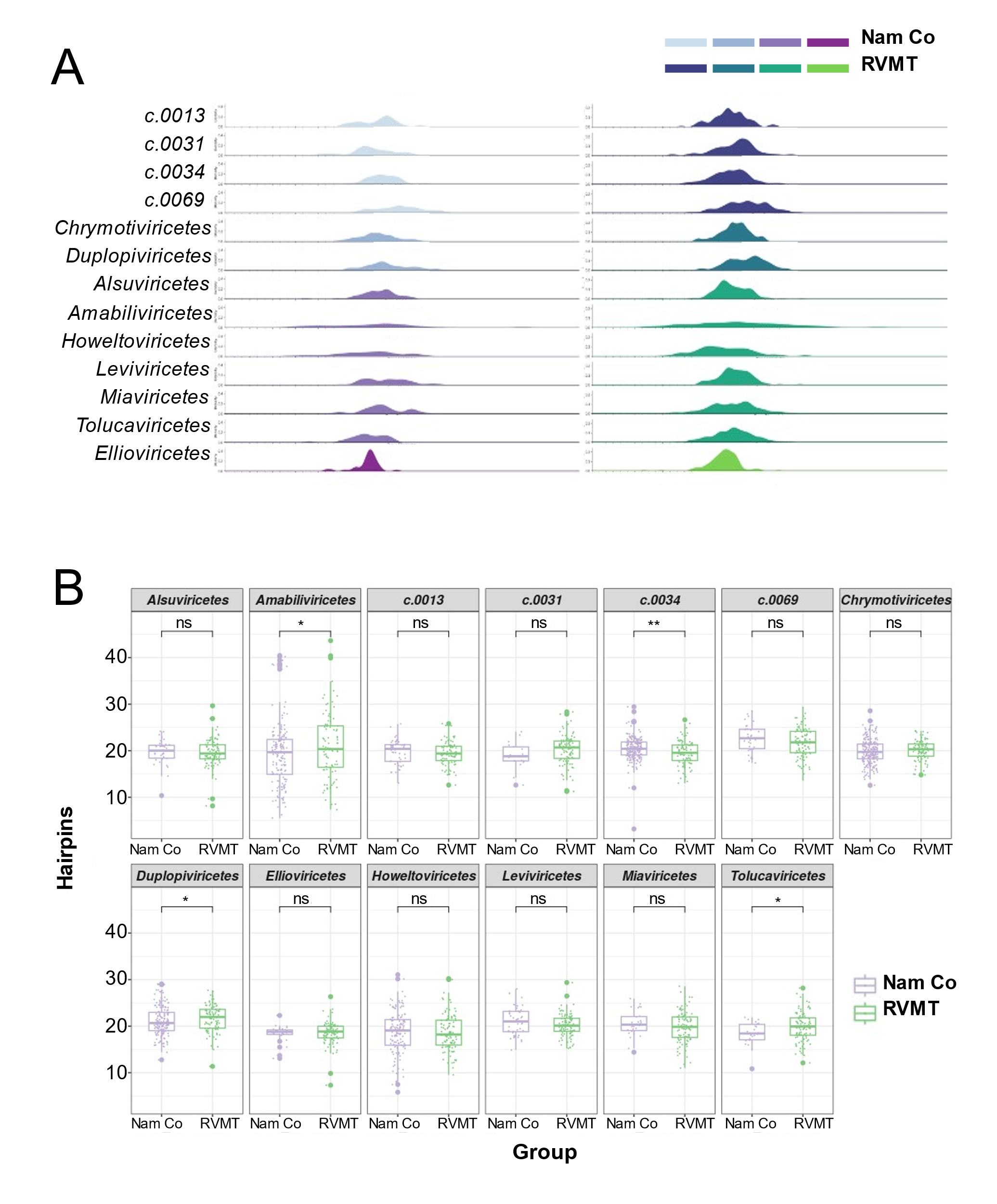

**Figure S36. The secondary structure (hairpins) of RNA viruses in Nam Co Lake compared to the RVMT database.** (A) The trend of hairpins with respect to viral classes. (B) Comparison of the hairpins between Nam Co Lake and the RVMT database at the level of viral classes.

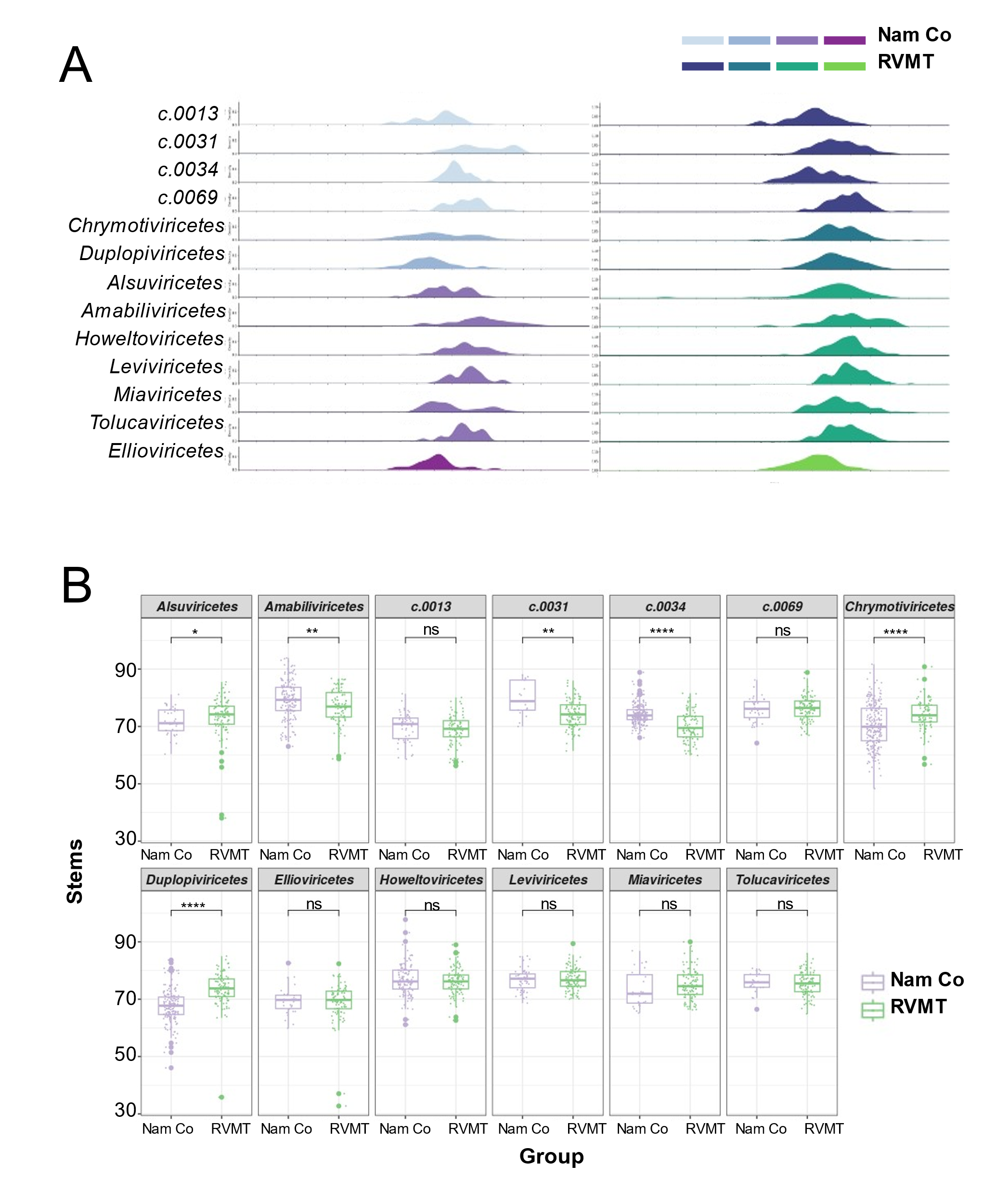

**Figure S37. The secondary structure (stems) of RNA viruses in Nam Co Lake compared to the RVMT database.** (A)The trend of stems with respect to viral classes. (B) Comparison of stems between Nam Co Lake and the RVMT database at the level of viral classes.

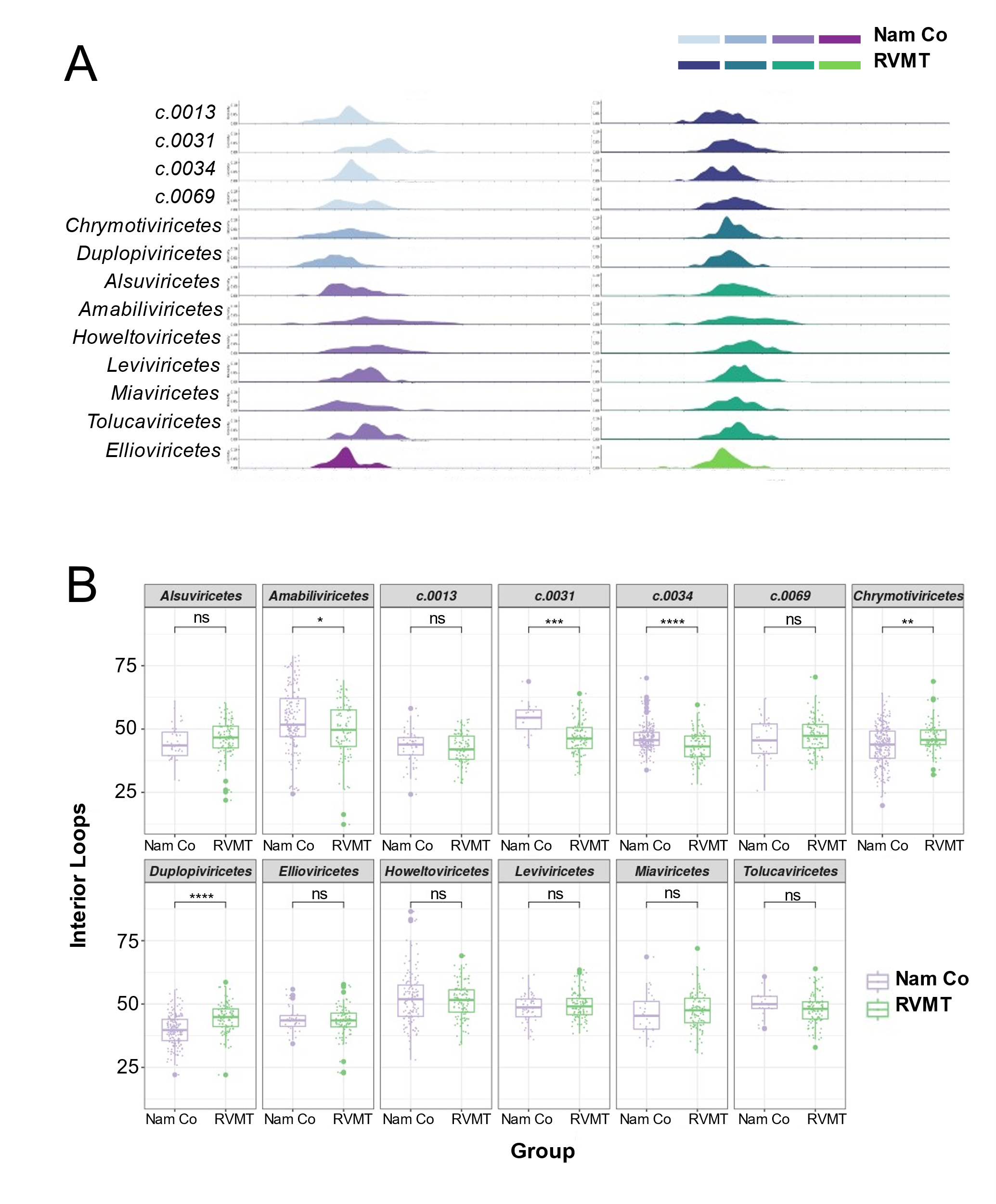

**Figure S38. The secondary structure (interior loops) of RNA viruses in Nam Co Lake compared to the RVMT database.** (A)The trend of interior loops with respect to viral classes. (B) Comparison of interior loops between Nam Co Lake and the RVMT database at the level of viral classes.

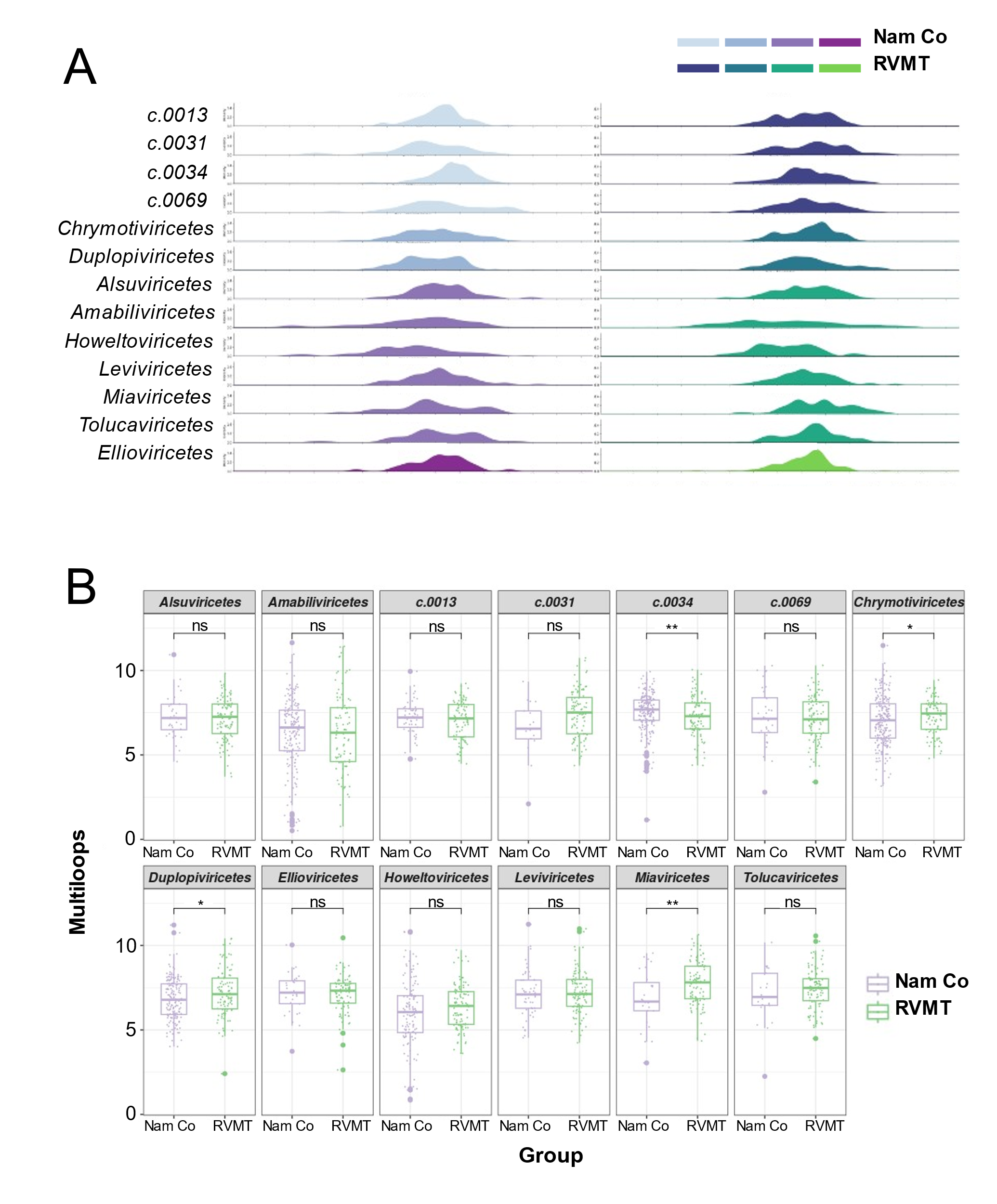

**Figure S39. The secondary structure (multiloops) of RNA viruses in Nam Co Lake compared to the RVMT database.** (A)The trend of multiloops with respect to viral classes. (B) Comparison of multiloops between Nam Co Lake and the RVMT database at the level of viral classes.

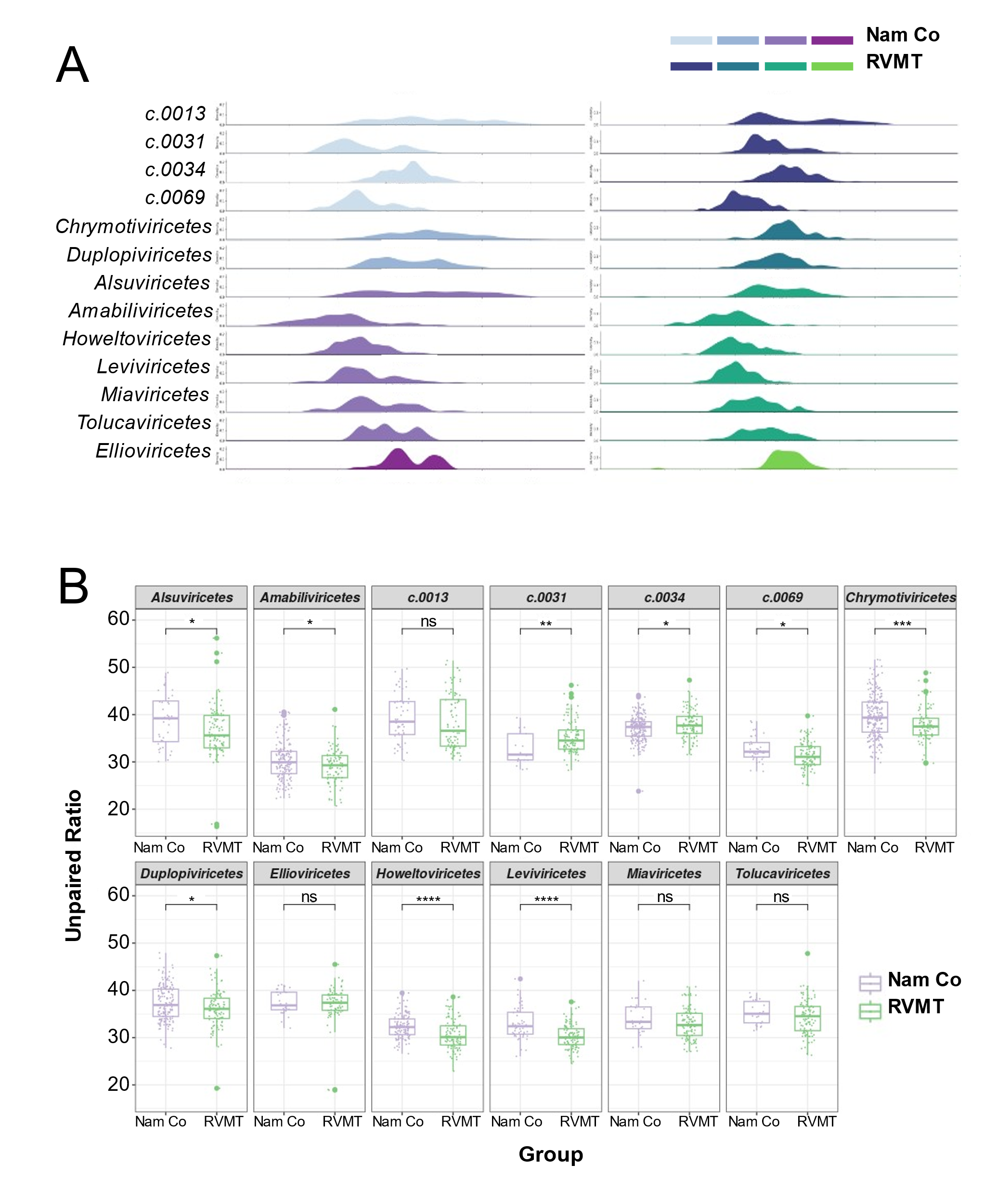

**Figure S40. The secondary structure (unpaired bias) of RNA viruses in Nam Co Lake compared to the RVMT database.** (A)The trend of unpaired bias with respect to viral classes. (B) Comparison of unpaired bias between Nam Co Lake and the RVMT database at the level of viral classes.

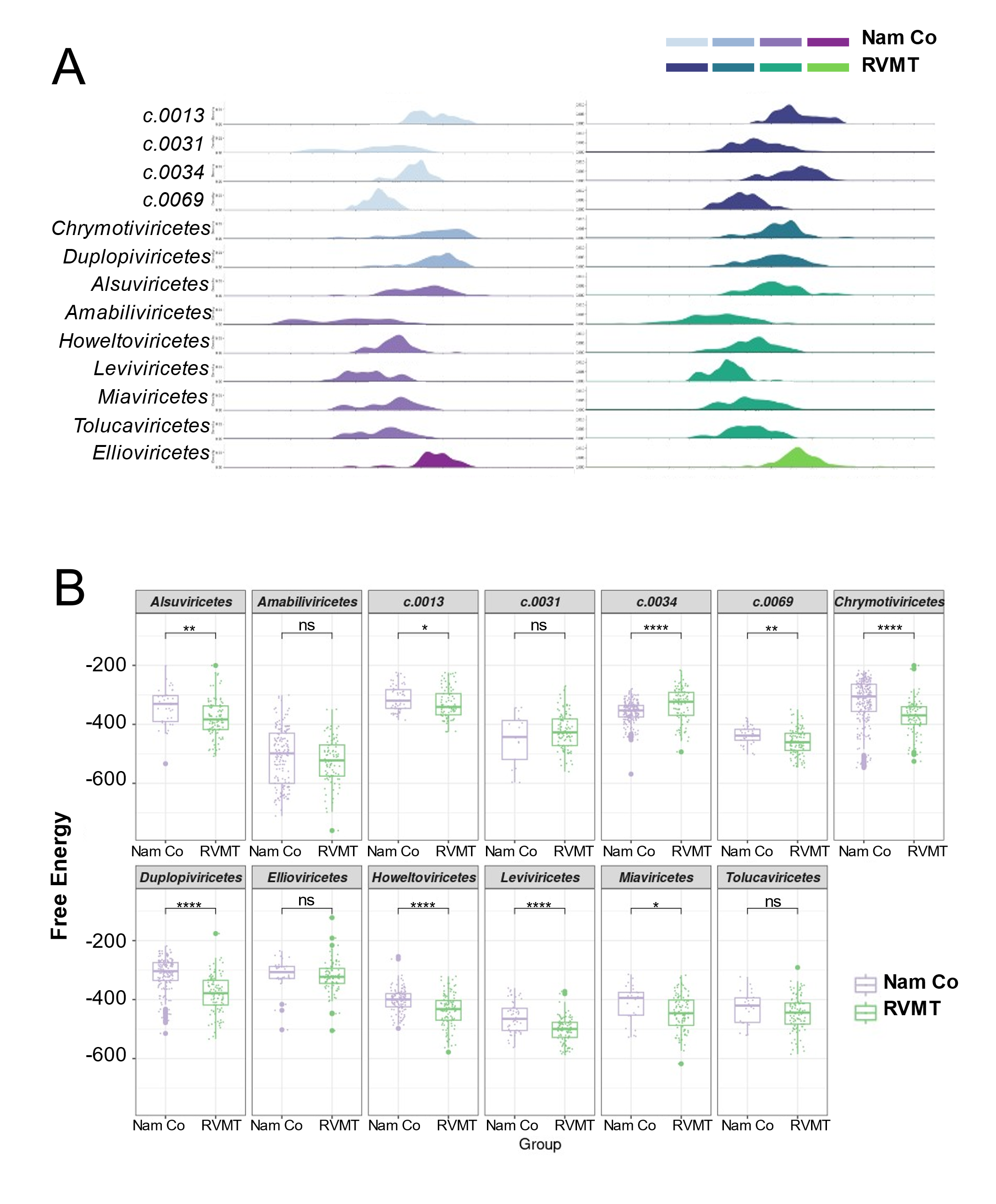

**Figure S41. The secondary structure (minimum free energy, MFE) of RNA viruses in Nam Co Lake compared to the RVMT database.** (A)The trend of MFE with respect to viral classes. (B) Comparison of MFE between Nam Co Lake and the RVMT database at the level of viral classes.

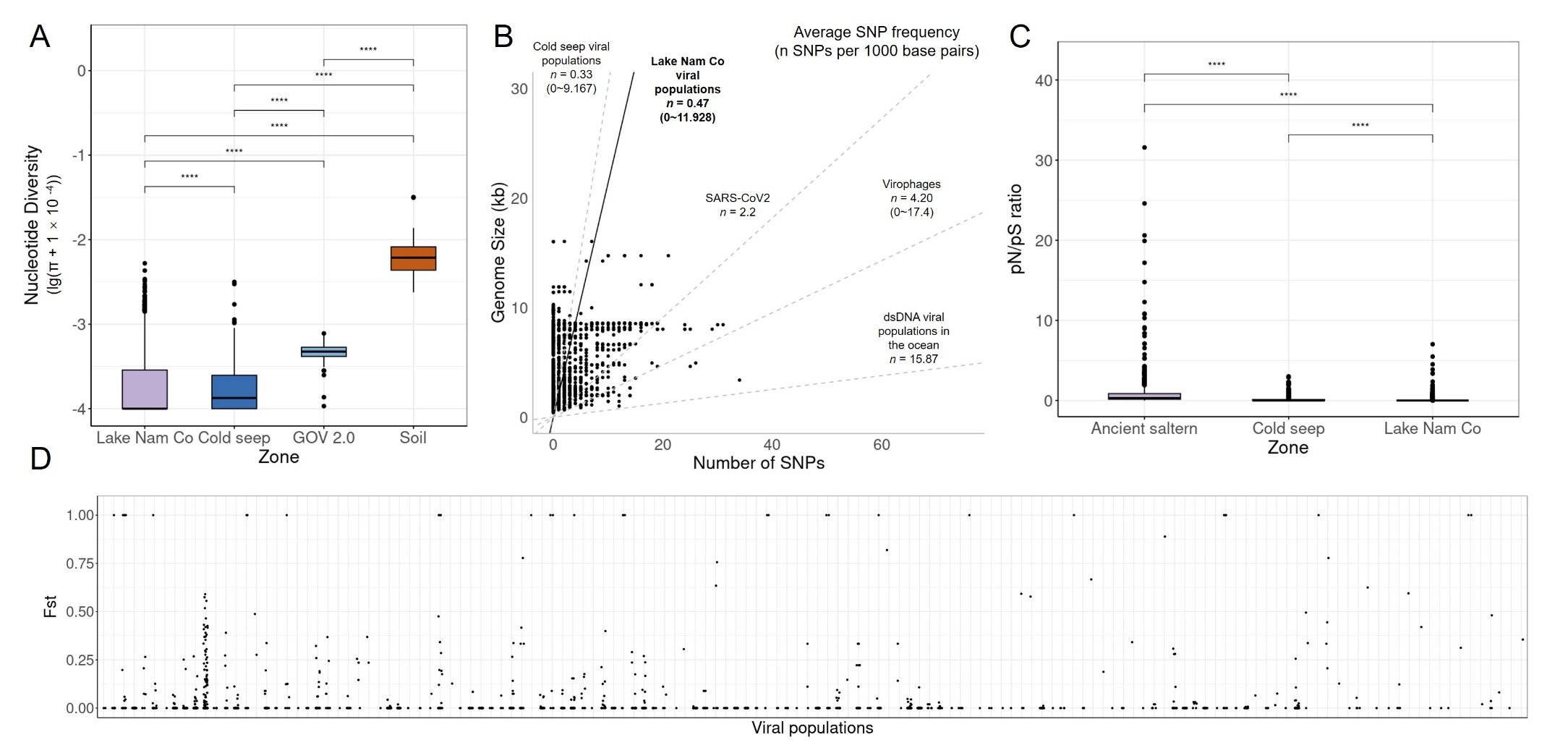

**Figure S42. Genome-wide evolutionary metrics of Lake Nam Co RNA viral populations.** (A) Nucleotide diversity of RNA viruses from Lake Nam Co (this study), viral populations from cold seeps, global seawater (GOV 2.0) and soils from various land-use types. (B) Comparisons of average SNP frequency among Lake Nam Co RNA viruses (this study), cold seeps viruses, SARS-CoV-2, virophages from North American freshwater lakes, and dominant dsDNA viruses in the oceans. Linear regressions are indicated for different viral populations derived from studies. (C) pN/pS ratio of viral genes from Lake Nam Co (this study), cold seeps and high-salinity groundwater of an ancient saltern. (D) Scatter diagram showing the F_ST_ values, representing the genetic differences between identical viral populations across various samples from Lake Nam Co.

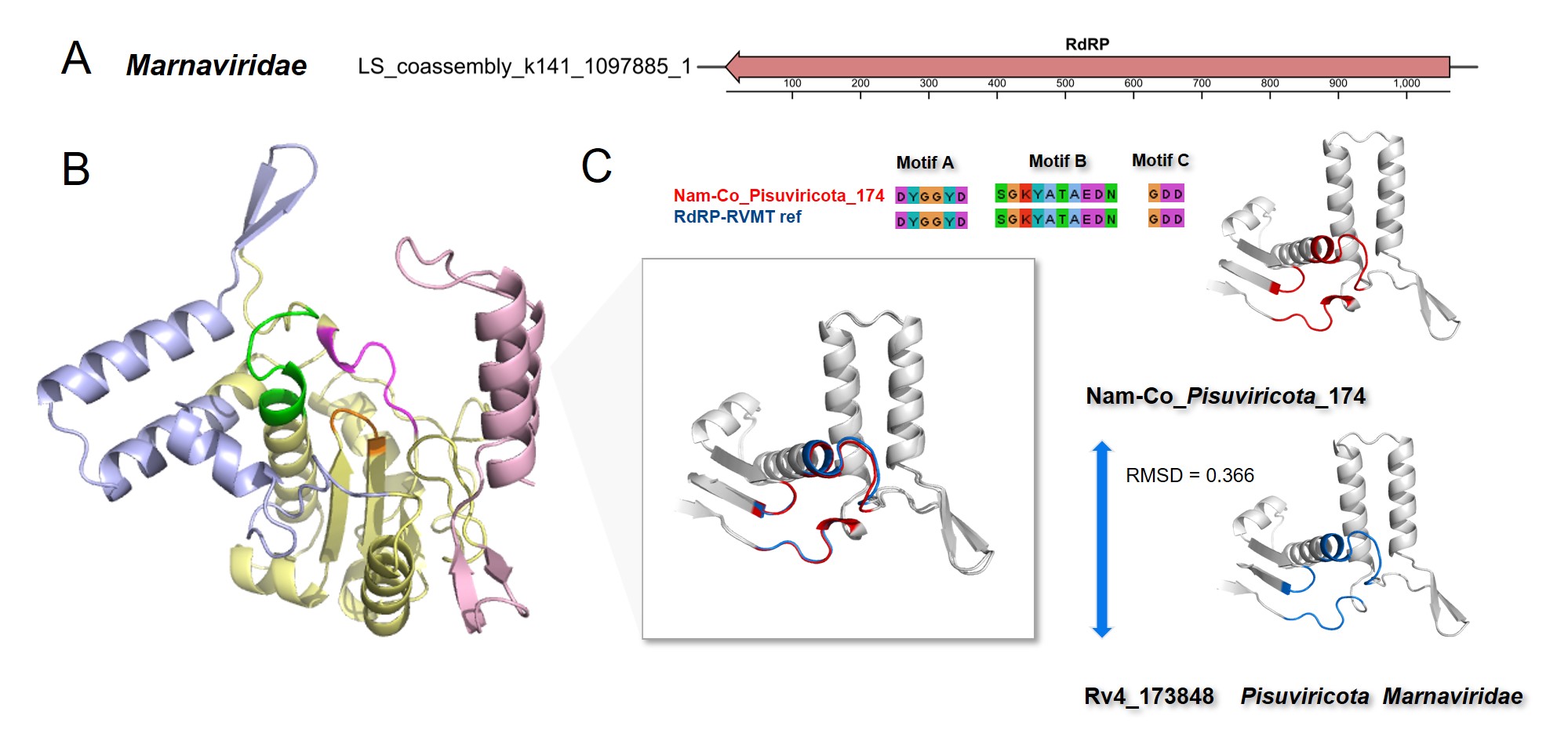

**Figure S43. Genomic composition and RdRP structure of the putative archaeal-infecting RNA virus Nam-Co_*Pisuviricota*_174 in the family *Marnaviridae*.** (A) Genomic Composition of Nam-Co_*Pisuviricota*_174. The genomic assembly of Nam-Co_*Pisuviricota*_174 yielded only a fragment encoding the RdRP. (B) The RdRP structure of Nam-Co_*Pisuviricota*_174. The “thumb”, “palm”, and “fingers” regions of RdRP are colored in pink, yellow, and blue, respectively. Within the “palm” region, catalytic center motifs A, B, and C are colored in purple, green, and orange. (C) Structural and motif composition of the catalytic center in the "palm" region of Nam-Co_*Pisuviricota*_174 compared with reference virus. The amino acid composition of motifs for the compared structures are listed in the actual arrangement sequence, shown above the schematic diagram. Motifs from the RdRP of Nam-Co_*Pisuviricota*_174 are colored in red, motifs from the RVMT reference structures are colored in dark blue. The RMSD values between structures are indicated by arrows of the same color as the referential structures.

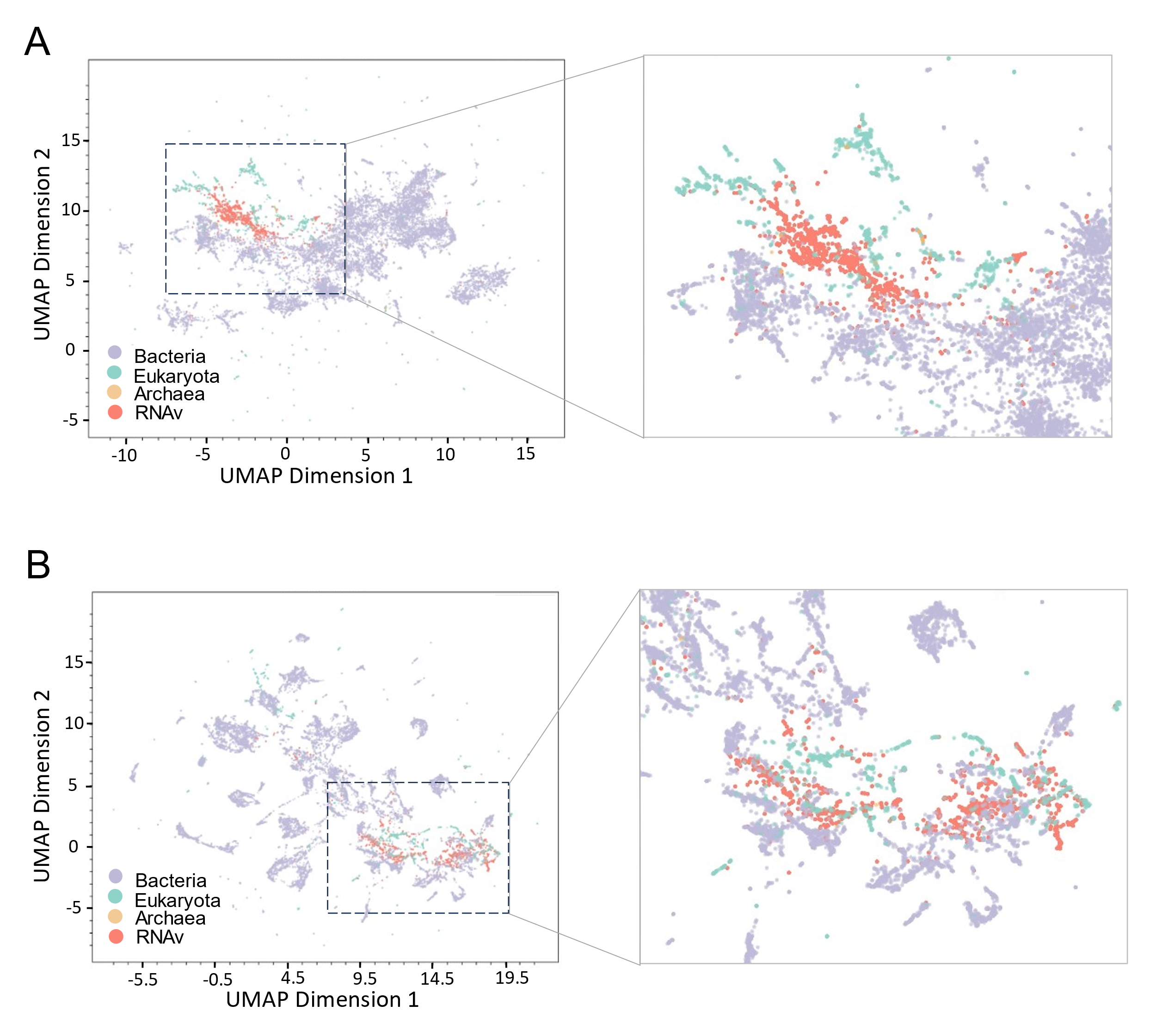

**Figure S44. The visualized UMAP dimensionality reduction maps of AAUF/DUF of RNA viruses and organisms in Lake Nam Co, superkingdom level.** Each dot represents a genomic sequence. Different organisms are indicated in different colors. RNA viruses are colored in red, bacteria are colored in purple, eukaryotes are colored in green, archaea are colored in orange. Regions including RNA viruses and related genomes are marked using a dashed line. (A) UMAP visualization of amino acid usage frequency (AAUF). (B) UMAP visualization of di-nucleotide usage frequency (DUF).

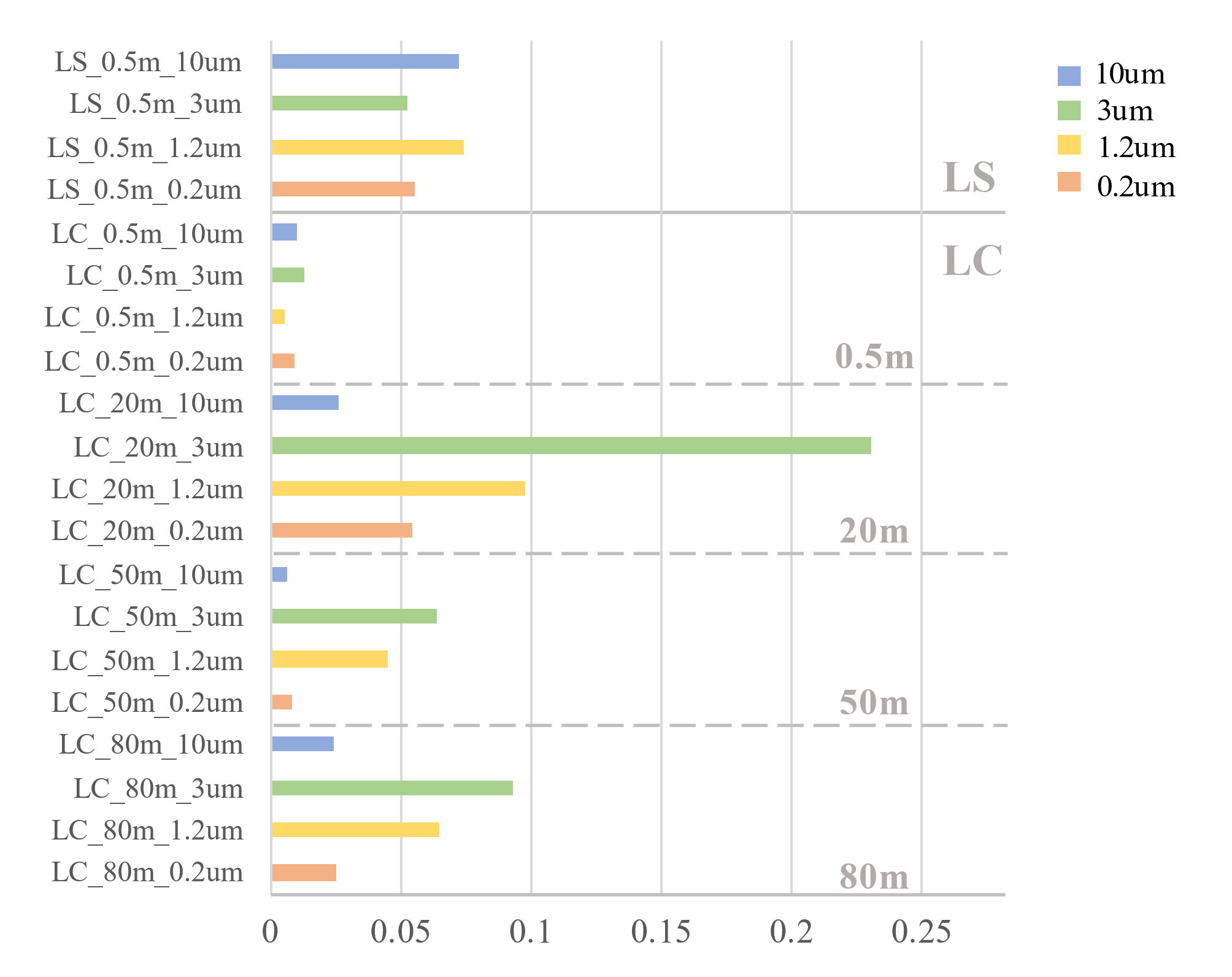

**Figure S45. Relative abundance of procaryotic viruses among RNA viruses (reads mapping).** The mapping results are arranged according to sampling site, depth, and particle size. Different sites are separated by solid lines, different depths by dashed lines, and different particle size samples are distinguished by color. LS, lakeshore; LC, lake center.

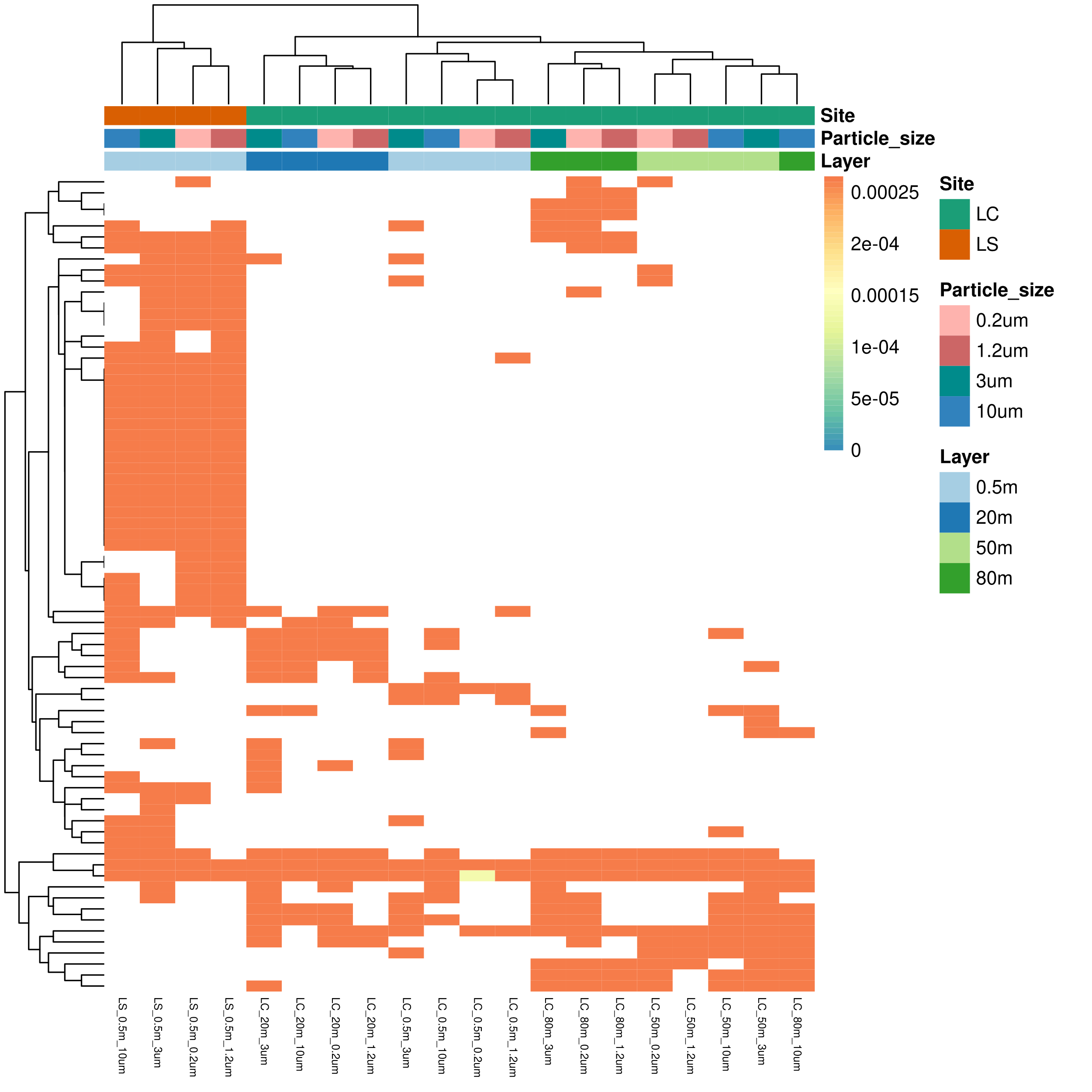

**Figure S46. The distribution of prokaryotic RNA viruses in different samples (reads mapping).** The relative abundance values of prokaryotic RNA virus contigs in a specific sample above the 75th percentile have been replaced by the 75th percentile. The site, particle size, and layer to which a specific sample belongs are marked at the top. LS, lakeshore; LC, lake center.

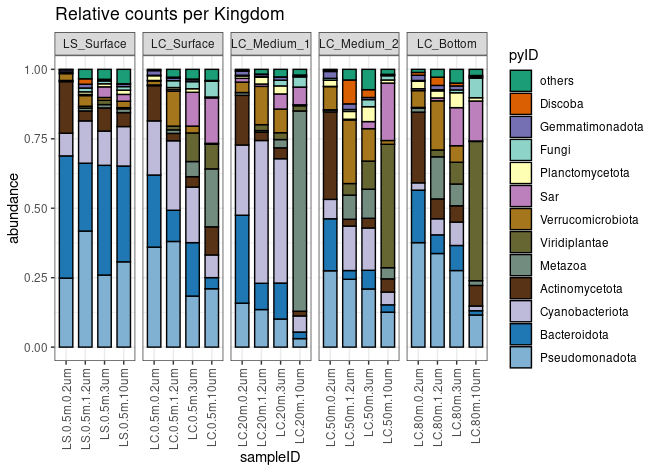

**Figure S47.** **Total compositions (kingdom/phylum) for each water samples.** Relative proportions were determined by the normalized number of hits related to rpsB from metatranscriptomes. The layers of the water samples are shown at the top of the bar. With the increase of water layer, the relative abundance of eukaryotic groups increased and that of prokaryotic groups decreased.

| **Table S1: Information of samples collected in Lake Nam Co.** | | | | | |
| --- | --- | --- | --- | --- | --- |
| **Stations** | **Samples** | **Longitude (°E)** | **Latitude (°N)** | **Depth (m)** | **Particle size (um)** |
| ^1^LS_Surface (0.5m) | LS-1 | 90.98 | 30.79 | 0.5 | 0.2 |
|  | LS-2 | 90.98 | 30.79 | 0.5 | 1.2 |
|  | LS-3 | 90.98 | 30.79 | 0.5 | 3 |
|  | LS-4 | 90.98 | 30.79 | 0.5 | 10 |
| ^2^LC_Surface (0.5m) | LC-1 | 90.48 | 30.46 | 0.5 | 0.2 |
|  | LC-2 | 90.48 | 30.46 | 0.5 | 1.2 |
|  | LC-3 | 90.48 | 30.46 | 0.5 | 3 |
|  | LC-4 | 90.48 | 30.46 | 0.5 | 10 |
| LC_Medium_1 (20m) | LC-5 | 90.48 | 30.46 | 20 | 0.2 |
|  | LC-6 | 90.48 | 30.46 | 20 | 1.2 |
|  | LC-7 | 90.48 | 30.46 | 20 | 3 |
|  | LC-8 | 90.48 | 30.46 | 20 | 10 |
| LC_Medium_2 (50m) | LC-9 | 90.48 | 30.46 | 50 | 0.2 |
|  | LC-10 | 90.48 | 30.46 | 50 | 1.2 |
|  | LC-11 | 90.48 | 30.46 | 50 | 3 |
|  | LC-12 | 90.48 | 30.46 | 50 | 10 |
| LC_Bottom (80m) | LC-13 | 90.48 | 30.46 | 80 | 0.2 |
|  | LC-14 | 90.48 | 30.46 | 80 | 1.2 |
|  | LC-15 | 90.48 | 30.46 | 80 | 3 |
|  | LC-16 | 90.48 | 30.46 | 80 | 10 |
| 1.LS: The station near the lakeshore (90.98°E, 30.79°N) | | | |  |  |
| 2.LC: The station near the lake center (90.48°E, 30.46°N) | | | |  |  |

| **Table S2. Environmental parameters of the water column and nutrient concentrations in different water layers.** | | | | | | | | | | | | | | |
| --- | --- | --- | --- | --- | --- | --- | --- | --- | --- | --- | --- | --- | --- | --- |
| **Station** | **Depth**  **(m)** | **Temperature**  **(℃)** | **pH** | **Salinity**  **(‰)** | **ODO**  **(mg/L)** | **DOC**  **(mg/L)** | **TN**  **(mg/L)** | **DTN**  **(mg/L)** | **NH4N**  **(mg/L)** | **NO3N**  **(mg/L)** | **NO2N**  **(mg/L)** | **TP**  **(mg/L)** | **DTP**  **(mg/L)** | **PO4P**  **(mg/L)** |
| ^1^LS | 0 | 12.888 | 9.17 | 1.05 | 11.41 | NA | NA | NA | 0.0094 | 0.0093 | 0.00069 | NA | NA | 0.007 |
| ^2^LC | 0 | 12.911 | 9.29 | 1.05 | 11.14 | 3.437 | 0.382 | 0.2834 | 0.029 | 0.01 | 1E-04 | 0.0114 | 0.0096 | 0.0078 |
|  | 2 | 12.905 | 9.32 | 1.05 | 11.29 | 3.423 | 0.3699 | 0.2647 | 0.02 | 0.0187 | 1E-04 | 0.0124 | 0.0123 | 0.007 |
|  | 5 | 12.87 | 9.32 | 1.05 | 11.23 | 3.427 | 0.4313 | 0.2729 | 0.0376 | 0.0094 | 1E-04 | 0.0131 | 0.0117 | 0.0066 |
|  | 10 | 12.795 | 9.3 | 1.05 | 11.08 | 3.498 | 0.5387 | 0.2679 | 0.0142 | 0.0266 | 0 | 0.0325 | 0.008 | 0.0039 |
|  | 20 | 11.161 | 9.3 | 1.08 | 11.12 | 3.451 | 0.3816 | 0.2562 | 0.0178 | 0.009 | 0 | 0.0225 | 0.0122 | 0.0083 |
|  | 30 | 6.404 | 9.39 | 1.05 | 13.33 | 3.43 | 0.3565 | 0.2965 | 0.0674 | 0.0111 | 0 | 0.0418 | 0.0122 | 0.0087 |
|  | 40 | 4.57 | 9.4 | 1.05 | 13.78 | 3.453 | 0.4486 | 0.2827 | 0.0254 | 0.0168 | 0.0003 | 0.0131 | 0.0123 | 0.0083 |
|  | 50 | 3.808 | 9.4 | 1.05 | 13.61 | 3.421 | 0.6414 | 0.3195 | 0.0242 | 0.0154 | 0.0003 | 0.0588 | 0.0126 | 0.0086 |
|  | 60 | 3.608 | 9.38 | 1.05 | 12.98 | 3.393 | 0.4454 | 0.2756 | 0.0488 | 0.0199 | 0.0002 | 0.0133 | 0.012 | 0.0092 |
|  | 70 | 3.59 | 9.37 | 1.05 | 12.76 | 3.393 | 0.354 | 0.2834 | 0.0296 | 0.0172 | 0.0003 | 0.0182 | 0.0131 | 0.0085 |
|  | 80 | 3.576 | 9.37 | 1.05 | 12.55 | 3.433 | 0.2627 | 0.2338 | 0.0276 | 0.0151 | 1E-04 | 0.0113 | 0.0109 | 0.0091 |
|  | 90 | 3.577 | 9.36 | 1.05 | 12.37 | 3.493 | 0.7071 | 0.4628 | 0.0568 | 0.0183 | 0.0003 | 0.2163 | 0.0135 | 0.0087 |
| 1.LS: The station near the lakeshore (90.98°E, 30.79°N) | | | | |  |  |  |  |  |  |  |  |  |  |
| 2.LC: The station near the lake center (90.48°E, 30.46°N) | | | | |  |  |  |  |  |  |  |  |  |  |

| **Table S3. Information of 742 newly identified RdRPs in Nam Co Lake.** | | | | | | | | | |
| --- | --- | --- | --- | --- | --- | --- | --- | --- | --- |
| **RdRP (90%AAI)** | **Phylum** | **Class** | **Order** | **Family** | **40%AAI new groups**  **(family_level)** | **75%AAI new groups**  **(Genus_level)** | **Method** | **Name** | **pLDDT (pdb)** |
| LS_coassembly_k141_910916_1 | *Birnaviridae* |  |  |  | *Nam-Co_family_01* | *Nam-Co_genus_001* | structure | *Nam-Co_Birnaviridae_001* | 60.91 |
| LC_coassembly_k141_309995_1 | *Birnaviridae* |  |  |  | *Nam-Co_family_02* | *Nam-Co_genus_002* | blastp | *Nam-Co_Birnaviridae_002* | 84.64 |
| LC_coassembly_k141_330298_1 | *Birnaviridae* |  |  |  | *Nam-Co_family_03* | *Nam-Co_genus_003* | blastp | *Nam-Co_Birnaviridae_003* | 84.62 |
| LC_coassembly_k141_76639_2 | *Lenarviricota* |  |  |  | *Nam-Co_family_04* | *Nam-Co_genus_004* | structure | *Nam-Co_Lenarviricota_001* | 65.77 |
| LC_coassembly_k141_1177772_3 | *Lenarviricota* | *Leviviricetes* | *Norzivirales* | *Atkinsviridae* |  | *Nam-Co_genus_113* | blastp | *Nam-Co_Lenarviricota_002* | 94.07 |
| LC_coassembly_k141_1968509_1 | *Lenarviricota* | *Leviviricetes* | *Norzivirales* | *Atkinsviridae* |  | *Nam-Co_genus_114* | blastp | *Nam-Co_Lenarviricota_003* | 94.16 |
| LC_coassembly_k141_2315390_1 | *Lenarviricota* | *Leviviricetes* | *Norzivirales* | *Atkinsviridae* |  | *Nam-Co_genus_115* | blastp | *Nam-Co_Lenarviricota_004* | 95.45 |
| LC_coassembly_k141_1817896_1 | *Lenarviricota* | *Leviviricetes* | *Norzivirales* | *Atkinsviridae* |  | *Nam-Co_genus_116* | blastp | *Nam-Co_Lenarviricota_005* | 95.77 |
| LS_coassembly_k141_54629_1 | *Lenarviricota* | *Miaviricetes* | *Ourlivirales* | *Botourmiaviridae* |  | *Nam-Co_genus_146* | blastp | *Nam-Co_Lenarviricota_006* | 94.81 |
| LS_coassembly_k141_604047_1 | *Lenarviricota* | *Miaviricetes* | *Ourlivirales* | *Botourmiaviridae* |  |  | blastp | *Nam-Co_Lenarviricota_007* | 92.91 |
| LS_coassembly_k141_199685_1 | *Lenarviricota* | *Miaviricetes* | *Ourlivirales* | *Botourmiaviridae* |  |  | blastp | *Nam-Co_Lenarviricota_008* | 94.74 |
| LS_coassembly_k141_30722_1 | *Lenarviricota* | *Miaviricetes* | *Ourlivirales* | *Botourmiaviridae* |  | *Nam-Co_genus_147* | blastp | *Nam-Co_Lenarviricota_009* | 95.22 |
| LC_coassembly_k141_2698212_1 | *Lenarviricota* | *Miaviricetes* | *Ourlivirales* | *Botourmiaviridae* |  | *Nam-Co_genus_145* | blastp | *Nam-Co_Lenarviricota_010* | 91.35 |
| LS_coassembly_k141_347998_3 | *Lenarviricota* | *Miaviricetes* | *Ourlivirales* | *Botourmiaviridae* | *Nam-Co_family_19* | *Nam-Co_genus_148* | blastp | *Nam-Co_Lenarviricota_011* | 91.18 |
| LC_coassembly_k141_14177_1 | *Lenarviricota* | *Miaviricetes* | *Ourlivirales* | *Botourmiaviridae* |  | *Nam-Co_genus_143* | blastp | *Nam-Co_Lenarviricota_012* | 92.13 |
| LS_coassembly_k141_980808_1 | *Lenarviricota* | *Miaviricetes* | *Ourlivirales* | *Botourmiaviridae* |  | *Nam-Co_genus_142* | blastp | *Nam-Co_Lenarviricota_013* | 90.3 |
| LS_coassembly_k141_667750_1 | *Lenarviricota* | *Miaviricetes* | *Ourlivirales* | *Botourmiaviridae* |  | *Nam-Co_genus_150* | blastp | *Nam-Co_Lenarviricota_014* | 91.75 |
| LC_coassembly_k141_227913_1 | *Lenarviricota* | *Miaviricetes* | *Ourlivirales* | *Botourmiaviridae* |  | *Nam-Co_genus_144* | blastp | *Nam-Co_Lenarviricota_015* | 93.65 |
| LS_coassembly_k141_1013649_1 | *Lenarviricota* | *Miaviricetes* | *Ourlivirales* | *Botourmiaviridae* |  | *Nam-Co_genus_149* | blastp | *Nam-Co_Lenarviricota_016* | 91.44 |
| LC_coassembly_k141_1498540_1 | *Lenarviricota* | *Miaviricetes* | *Ourlivirales* | *Botourmiaviridae* |  | *Nam-Co_genus_141* | blastp | *Nam-Co_Lenarviricota_017* | 94.08 |
| LC_50m_10um_k141_96448_1 | *Lenarviricota* | *Miaviricetes* | *Ourlivirales* | *Botourmiaviridae* |  | *Nam-Co_genus_141* | blastp | *Nam-Co_Lenarviricota_018* | 93.88 |
| LC_coassembly_k141_303976_1 | *Lenarviricota* | *Miaviricetes* | *Ourlivirales* | *Botourmiaviridae* |  |  | blastp | *Nam-Co_Lenarviricota_019* | 92.54 |
| LS_coassembly_k141_1183900_1 | *Lenarviricota* | *Leviviricetes* | *Norzivirales* | *f.0361* |  |  | blastp | *Nam-Co_Lenarviricota_020* | 95.09 |
| LS_coassembly_k141_981221_2 | *Lenarviricota* | *Leviviricetes* | *Norzivirales* | *f.0361* |  | *Nam-Co_genus_118* | blastp | *Nam-Co_Lenarviricota_021* | 94.35 |
| LS_0.5m_1.2um_k141_623158_1 | *Lenarviricota* | *Leviviricetes* | *Norzivirales* | *f.0361* |  | *Nam-Co_genus_117* | blastp | *Nam-Co_Lenarviricota_022* | 94.28 |
| LS_coassembly_k141_169245_1 | *Lenarviricota* | *Leviviricetes* | *Norzivirales* | *f.0367* |  |  | blastp | *Nam-Co_Lenarviricota_023* | 95.97 |
| LS_coassembly_k141_112997_1 | *Lenarviricota* | *Leviviricetes* | *Norzivirales* | *f.0368* |  | *Nam-Co_genus_121* | blastp | *Nam-Co_Lenarviricota_024* | 96.07 |
| LS_coassembly_k141_379730_2 | *Lenarviricota* | *Leviviricetes* | *Norzivirales* | *f.0368* | *Nam-Co_family_18* | *Nam-Co_genus_119* | blastp | *Nam-Co_Lenarviricota_025* | 93.34 |
| LS_coassembly_k141_1338665_1 | *Lenarviricota* | *Leviviricetes* | *Norzivirales* | *f.0368* |  | *Nam-Co_genus_120* | blastp | *Nam-Co_Lenarviricota_026* | 96.04 |
| LS_coassembly_k141_150032_1 | *Lenarviricota* | *c.0061.base-Levi* | *o.0082* | *f.0378* | *Nam-Co_family_14* | *Nam-Co_genus_066* | blastp | *Nam-Co_Lenarviricota_027* | 85.18 |
| LC_0.5m_10um_k141_140412_1 | *Lenarviricota* | *c.0064.base-Howelto* | *o.0085* | *f.0381* |  | *Nam-Co_genus_067* | blastp | *Nam-Co_Lenarviricota_028* | 77.39 |
| LS_coassembly_k141_723190_1 | *Lenarviricota* | *c.0066.base-Howelto* | *o.0087* | *f.0383* |  | *Nam-Co_genus_068* | blastp | *Nam-Co_Lenarviricota_029* | 85.67 |
| LS_coassembly_k141_399847_1 | *Lenarviricota* | *c.0067.base-Howelto* | *o.0088* | *f.0384* |  | *Nam-Co_genus_069* | blastp | *Nam-Co_Lenarviricota_030* | 87.56 |
| LS_0.5m_1.2um_k141_340418_1 | *Lenarviricota* | *c.0067.base-Howelto* | *o.0088* | *f.0384* |  | *Nam-Co_genus_069* | blastp | *Nam-Co_Lenarviricota_031* | 87.73 |
| LS_coassembly_k141_640025_2 | *Lenarviricota* | *c.0068.base-Howelto* | *o.0089* | *f.0385* |  | *Nam-Co_genus_071* | blastp | *Nam-Co_Lenarviricota_032* | 95.6 |
| LS_coassembly_k141_280312_1 | *Lenarviricota* | *c.0068.base-Howelto* | *o.0089* | *f.0385* |  | *Nam-Co_genus_072* | blastp | *Nam-Co_Lenarviricota_033* | 95.51 |
| LS_coassembly_k141_1046808_2 | *Lenarviricota* | *c.0068.base-Howelto* | *o.0089* | *f.0385* |  | *Nam-Co_genus_070* | blastp | *Nam-Co_Lenarviricota_034* | 95.02 |
| LS_coassembly_k141_886170_1 | *Lenarviricota* | *c.0069.base-Howelto* | *o.0090* | *f.0386* |  |  | blastp | *Nam-Co_Lenarviricota_035* | 95.62 |
| LS_coassembly_k141_786484_1 | *Lenarviricota* | *c.0069.base-Howelto* | *o.0090* | *f.0386* |  | *Nam-Co_genus_073* | blastp | *Nam-Co_Lenarviricota_036* | 96.27 |
| LS_coassembly_k141_562058_1 | *Lenarviricota* | *c.0069.base-Howelto* | *o.0090* | *f.0386* |  | *Nam-Co_genus_080* | blastp | *Nam-Co_Lenarviricota_037* | 90.82 |
| LS_coassembly_k141_1348429_1 | *Lenarviricota* | *c.0069.base-Howelto* | *o.0090* | *f.0386* |  |  | blastp | *Nam-Co_Lenarviricota_038* | 94.73 |
| LS_coassembly_k141_1016243_1 | *Lenarviricota* | *c.0069.base-Howelto* | *o.0090* | *f.0386* |  | *Nam-Co_genus_084* | blastp | *Nam-Co_Lenarviricota_039* | 87.73 |
| LS_coassembly_k141_1122728_1 | *Lenarviricota* | *c.0069.base-Howelto* | *o.0090* | *f.0386* |  | *Nam-Co_genus_086* | blastp | *Nam-Co_Lenarviricota_040* | 93.11 |
| LS_coassembly_k141_244743_1 | *Lenarviricota* | *c.0069.base-Howelto* | *o.0090* | *f.0386* |  | *Nam-Co_genus_075* | blastp | *Nam-Co_Lenarviricota_041* | 94.85 |
| LS_coassembly_k141_1216739_1 | *Lenarviricota* | *c.0069.base-Howelto* | *o.0090* | *f.0386* |  | *Nam-Co_genus_079* | blastp | *Nam-Co_Lenarviricota_042* | 96 |
| LC_coassembly_k141_1595254_1 | *Lenarviricota* | *c.0069.base-Howelto* | *o.0090* | *f.0386* |  | *Nam-Co_genus_079* | blastp | *Nam-Co_Lenarviricota_043* | 95.67 |
| LS_coassembly_k141_95774_1 | *Lenarviricota* | *c.0069.base-Howelto* | *o.0090* | *f.0386* |  | *Nam-Co_genus_077* | blastp | *Nam-Co_Lenarviricota_044* | 95.42 |
| LS_coassembly_k141_532808_1 | *Lenarviricota* | *c.0069.base-Howelto* | *o.0090* | *f.0386* |  |  | blastp | *Nam-Co_Lenarviricota_045* | 95.55 |
| LS_coassembly_k141_1176845_1 | *Lenarviricota* | *c.0069.base-Howelto* | *o.0090* | *f.0386* |  | *Nam-Co_genus_083* | blastp | *Nam-Co_Lenarviricota_046* | 94.97 |
| LS_coassembly_k141_967088_1 | *Lenarviricota* | *c.0069.base-Howelto* | *o.0090* | *f.0386* |  | *Nam-Co_genus_078* | blastp | *Nam-Co_Lenarviricota_047* | 92.45 |
| LS_coassembly_k141_1115973_1 | *Lenarviricota* | *c.0069.base-Howelto* | *o.0090* | *f.0386* |  | *Nam-Co_genus_085* | blastp | *Nam-Co_Lenarviricota_048* | 95.4 |
| LC_0.5m_1.2um_k141_132419_1 | *Lenarviricota* | *c.0069.base-Howelto* | *o.0090* | *f.0386* |  | *Nam-Co_genus_081* | blastp | *Nam-Co_Lenarviricota_049* | 95.44 |
| LC_20m_0.2um_k141_72846_1 | *Lenarviricota* | *c.0069.base-Howelto* | *o.0090* | *f.0386* |  | *Nam-Co_genus_081* | blastp | *Nam-Co_Lenarviricota_050* | 95.43 |
| LS_coassembly_k141_1113875_1 | *Lenarviricota* | *c.0069.base-Howelto* | *o.0090* | *f.0386* |  | *Nam-Co_genus_082* | blastp | *Nam-Co_Lenarviricota_051* | 95.23 |
| LS_coassembly_k141_588663_1 | *Lenarviricota* | *c.0069.base-Howelto* | *o.0090* | *f.0386* |  | *Nam-Co_genus_074* | blastp | *Nam-Co_Lenarviricota_052* | 88.99 |
| LS_coassembly_k141_18251_1 | *Lenarviricota* | *c.0069.base-Howelto* | *o.0090* | *f.0386* | *Nam-Co_family_15* | *Nam-Co_genus_076* | blastp | *Nam-Co_Lenarviricota_053* | 95.62 |
| LC_20m_0.2um_k141_107269_1 | *Lenarviricota* | *c.0071.base-Howelto* | *o.0092* | *f.0388* |  | *Nam-Co_genus_090* | blastp | *Nam-Co_Lenarviricota_054* | 94.11 |
| LC_50m_1.2um_k141_238221_1 | *Lenarviricota* | *c.0071.base-Howelto* | *o.0092* | *f.0388* |  | *Nam-Co_genus_090* | blastp | *Nam-Co_Lenarviricota_055* | 94.07 |
| LS_coassembly_k141_122735_1 | *Lenarviricota* | *c.0071.base-Howelto* | *o.0092* | *f.0388* |  | *Nam-Co_genus_090* | blastp | *Nam-Co_Lenarviricota_056* | 94.03 |
| LC_coassembly_k141_860293_1 | *Lenarviricota* | *c.0071.base-Howelto* | *o.0092* | *f.0388* |  | *Nam-Co_genus_088* | blastp | *Nam-Co_Lenarviricota_057* | 90.18 |
| LC_coassembly_k141_1002275_1 | *Lenarviricota* | *c.0071.base-Howelto* | *o.0092* | *f.0388* |  | *Nam-Co_genus_089* | blastp | *Nam-Co_Lenarviricota_058* | 94.28 |
| LC_coassembly_k141_41569_1 | *Lenarviricota* | *c.0071.base-Howelto* | *o.0092* | *f.0388* |  | *Nam-Co_genus_087* | blastp | *Nam-Co_Lenarviricota_059* | 93.8 |
| LC_coassembly_k141_1669236_1 | *Lenarviricota* | *c.0074.base-Howelto* | *o.0095* | *f.0391* | *Nam-Co_family_16* | *Nam-Co_genus_091* | blastp | *Nam-Co_Lenarviricota_060* | 89.33 |
| LS_coassembly_k141_447615_1 | *Lenarviricota* | *Leviviricetes* | *Norzivirales* | *Fiersviridae* |  | *Nam-Co_genus_131* | blastp | *Nam-Co_Lenarviricota_061* | 96.97 |
| LS_coassembly_k141_808804_1 | *Lenarviricota* | *Leviviricetes* | *Norzivirales* | *Fiersviridae* |  |  | blastp | *Nam-Co_Lenarviricota_062* | 96.73 |
| LS_coassembly_k141_543237_2 | *Lenarviricota* | *Leviviricetes* | *Norzivirales* | *Fiersviridae* |  | *Nam-Co_genus_127* | blastp | *Nam-Co_Lenarviricota_063* | 95.29 |
| LC_coassembly_k141_864888_1 | *Lenarviricota* | *Leviviricetes* | *Norzivirales* | *Fiersviridae* |  | *Nam-Co_genus_129* | blastp | *Nam-Co_Lenarviricota_064* | 96.6 |
| LS_coassembly_k141_947790_2 | *Lenarviricota* | *Leviviricetes* | *Norzivirales* | *Fiersviridae* |  | *Nam-Co_genus_124* | blastp | *Nam-Co_Lenarviricota_065* | 96.61 |
| LS_coassembly_k141_709523_1 | *Lenarviricota* | *Leviviricetes* | *Norzivirales* | *Fiersviridae* |  | *Nam-Co_genus_126* | blastp | *Nam-Co_Lenarviricota_066* | 95.11 |
| LC_coassembly_k141_1805655_1 | *Lenarviricota* | *Leviviricetes* | *Norzivirales* | *Fiersviridae* |  | *Nam-Co_genus_130* | blastp | *Nam-Co_Lenarviricota_067* | 96.96 |
| LS_coassembly_k141_1121902_1 | *Lenarviricota* | *Leviviricetes* | *Norzivirales* | *Fiersviridae* |  | *Nam-Co_genus_128* | blastp | *Nam-Co_Lenarviricota_068* | 95.31 |
| LS_coassembly_k141_839961_1 | *Lenarviricota* | *Leviviricetes* | *Norzivirales* | *Fiersviridae* |  |  | blastp | *Nam-Co_Lenarviricota_069* | 96.23 |
| LS_coassembly_k141_656443_2 | *Lenarviricota* | *Leviviricetes* | *Norzivirales* | *Fiersviridae* |  | *Nam-Co_genus_125* | blastp | *Nam-Co_Lenarviricota_070* | 94.87 |
| LS_coassembly_k141_259450_1 | *Lenarviricota* | *Leviviricetes* | *Norzivirales* | *Fiersviridae* |  |  | blastp | *Nam-Co_Lenarviricota_071* | 96.5 |
| LC_coassembly_k141_1827790_1 | *Lenarviricota* | *Leviviricetes* | *Norzivirales* | *Fiersviridae* |  | *Nam-Co_genus_123* | blastp | *Nam-Co_Lenarviricota_072* | 96.74 |
| LS_coassembly_k141_63722_2 | *Lenarviricota* | *Leviviricetes* | *Norzivirales* | *Fiersviridae* |  | *Nam-Co_genus_122* | blastp | *Nam-Co_Lenarviricota_073* | 89.53 |
| LS_coassembly_k141_8864_3 | *Lenarviricota* | *Leviviricetes* | *Norzivirales* | *Fiersviridae* |  |  | blastp | *Nam-Co_Lenarviricota_074* | 96.06 |
| LC_coassembly_k141_2695273_1 | *Lenarviricota* | *Howeltoviricetes* | *Cryppavirales* | *Mitoviridae* |  | *Nam-Co_genus_105* | blastp | *Nam-Co_Lenarviricota_075* | 81.96 |
| LS_coassembly_k141_108362_1 | *Lenarviricota* | *Howeltoviricetes* | *Cryppavirales* | *Mitoviridae* |  | *Nam-Co_genus_098* | blastp | *Nam-Co_Lenarviricota_076* | 86.86 |
| LS_0.5m_3um_k141_161223_1 | *Lenarviricota* | *Howeltoviricetes* | *Cryppavirales* | *Mitoviridae* |  | *Nam-Co_genus_099* | blastp | *Nam-Co_Lenarviricota_077* | 91.43 |
| LC_coassembly_k141_2721539_1 | *Lenarviricota* | *Howeltoviricetes* | *Cryppavirales* | *Mitoviridae* |  |  | blastp | *Nam-Co_Lenarviricota_078* | 93.52 |
| LC_coassembly_k141_328812_1 | *Lenarviricota* | *Howeltoviricetes* | *Cryppavirales* | *Mitoviridae* |  | *Nam-Co_genus_110* | blastp | *Nam-Co_Lenarviricota_079* | 87.77 |
| LC_coassembly_k141_787612_1 | *Lenarviricota* | *Howeltoviricetes* | *Cryppavirales* | *Mitoviridae* |  |  | blastp | *Nam-Co_Lenarviricota_080* | 87.03 |
| LC_20m_3um_k141_402748_1 | *Lenarviricota* | *Howeltoviricetes* | *Cryppavirales* | *Mitoviridae* |  |  | blastp | *Nam-Co_Lenarviricota_081* | 88.4 |
| LC_80m_3um_k141_132991_1 | *Lenarviricota* | *Howeltoviricetes* | *Cryppavirales* | *Mitoviridae* |  |  | blastp | *Nam-Co_Lenarviricota_082* | 91.5 |
| LC_coassembly_k141_1873206_1 | *Lenarviricota* | *Howeltoviricetes* | *Cryppavirales* | *Mitoviridae* | *Nam-Co_family_17* | *Nam-Co_genus_096* | blastp | *Nam-Co_Lenarviricota_083* | 87.78 |
| LC_0.5m_10um_k141_28114_1 | *Lenarviricota* | *Howeltoviricetes* | *Cryppavirales* | *Mitoviridae* |  | *Nam-Co_genus_108* | blastp | *Nam-Co_Lenarviricota_084* | 91.74 |
| LC_0.5m_3um_k141_309296_1 | *Lenarviricota* | *Howeltoviricetes* | *Cryppavirales* | *Mitoviridae* |  |  | blastp | *Nam-Co_Lenarviricota_085* | 88.44 |
| LC_coassembly_k141_2106280_1 | *Lenarviricota* | *Howeltoviricetes* | *Cryppavirales* | *Mitoviridae* |  | *Nam-Co_genus_093* | blastp | *Nam-Co_Lenarviricota_086* | 93.87 |
| LC_coassembly_k141_2111917_1 | *Lenarviricota* | *Howeltoviricetes* | *Cryppavirales* | *Mitoviridae* |  | *Nam-Co_genus_093* | blastp | *Nam-Co_Lenarviricota_087* | 93.1 |
| LC_coassembly_k141_709835_1 | *Lenarviricota* | *Howeltoviricetes* | *Cryppavirales* | *Mitoviridae* |  | *Nam-Co_genus_093* | blastp | *Nam-Co_Lenarviricota_088* | 93.46 |
| LC_coassembly_k141_2428978_1 | *Lenarviricota* | *Howeltoviricetes* | *Cryppavirales* | *Mitoviridae* |  | *Nam-Co_genus_106* | blastp | *Nam-Co_Lenarviricota_089* | 89.55 |
| LC_20m_3um_k141_343813_1 | *Lenarviricota* | *Howeltoviricetes* | *Cryppavirales* | *Mitoviridae* |  | *Nam-Co_genus_102* | blastp | *Nam-Co_Lenarviricota_090* | 87.38 |
| LC_coassembly_k141_1074524_2 | *Lenarviricota* | *Howeltoviricetes* | *Cryppavirales* | *Mitoviridae* |  | *Nam-Co_genus_109* | blastp | *Nam-Co_Lenarviricota_091* | 85.44 |
| LC_50m_10um_k141_200829_1 | *Lenarviricota* | *Howeltoviricetes* | *Cryppavirales* | *Mitoviridae* |  | *Nam-Co_genus_108* | blastp | *Nam-Co_Lenarviricota_092* | 93.55 |
| LS_coassembly_k141_1052634_1 | *Lenarviricota* | *Howeltoviricetes* | *Cryppavirales* | *Mitoviridae* |  |  | blastp | *Nam-Co_Lenarviricota_093* | 93.63 |
| LC_coassembly_k141_2723452_1 | *Lenarviricota* | *Howeltoviricetes* | *Cryppavirales* | *Mitoviridae* |  | *Nam-Co_genus_092* | blastp | *Nam-Co_Lenarviricota_094* | 87.38 |
| LS_coassembly_k141_502318_1 | *Lenarviricota* | *Howeltoviricetes* | *Cryppavirales* | *Mitoviridae* |  | *Nam-Co_genus_099* | blastp | *Nam-Co_Lenarviricota_095* | 91.86 |
| LC_coassembly_k141_125988_1 | *Lenarviricota* | *Howeltoviricetes* | *Cryppavirales* | *Mitoviridae* |  | *Nam-Co_genus_110* | blastp | *Nam-Co_Lenarviricota_096* | 89.99 |
| LS_0.5m_10um_k141_171521_1 | *Lenarviricota* | *Howeltoviricetes* | *Cryppavirales* | *Mitoviridae* |  | *Nam-Co_genus_110* | blastp | *Nam-Co_Lenarviricota_097* | 91.41 |
| LS_coassembly_k141_460169_1 | *Lenarviricota* | *Howeltoviricetes* | *Cryppavirales* | *Mitoviridae* |  | *Nam-Co_genus_101* | blastp | *Nam-Co_Lenarviricota_098* | 90.43 |
| LC_coassembly_k141_2752873_1 | *Lenarviricota* | *Howeltoviricetes* | *Cryppavirales* | *Mitoviridae* |  | *Nam-Co_genus_106* | blastp | *Nam-Co_Lenarviricota_099* | 90.8 |
| LC_0.5m_10um_k141_4899_1 | *Lenarviricota* | *Howeltoviricetes* | *Cryppavirales* | *Mitoviridae* |  | *Nam-Co_genus_110* | blastp | *Nam-Co_Lenarviricota_100* | 87.7 |
| LC_20m_3um_k141_56124_1 | *Lenarviricota* | *Howeltoviricetes* | *Cryppavirales* | *Mitoviridae* |  | *Nam-Co_genus_110* | blastp | *Nam-Co_Lenarviricota_101* | 86.71 |
| LS_coassembly_k141_335016_1 | *Lenarviricota* | *Howeltoviricetes* | *Cryppavirales* | *Mitoviridae* |  |  | blastp | *Nam-Co_Lenarviricota_102* | 92.41 |
| LC_coassembly_k141_2686142_1 | *Lenarviricota* | *Howeltoviricetes* | *Cryppavirales* | *Mitoviridae* |  | *Nam-Co_genus_097* | blastp | *Nam-Co_Lenarviricota_103* | 89.18 |
| LS_coassembly_k141_919836_1 | *Lenarviricota* | *Howeltoviricetes* | *Cryppavirales* | *Mitoviridae* |  |  | blastp | *Nam-Co_Lenarviricota_104* | 93.57 |
| LS_coassembly_k141_891738_1 | *Lenarviricota* | *Howeltoviricetes* | *Cryppavirales* | *Mitoviridae* |  | *Nam-Co_genus_095* | blastp | *Nam-Co_Lenarviricota_105* | 93.95 |
| LC_0.5m_10um_k141_193132_1 | *Lenarviricota* | *Howeltoviricetes* | *Cryppavirales* | *Mitoviridae* |  | *Nam-Co_genus_110* | blastp | *Nam-Co_Lenarviricota_106* | 88.39 |
| LC_coassembly_k141_1900159_1 | *Lenarviricota* | *Howeltoviricetes* | *Cryppavirales* | *Mitoviridae* |  | *Nam-Co_genus_100* | blastp | *Nam-Co_Lenarviricota_107* | 92.51 |
| LC_50m_10um_k141_292869_1 | *Lenarviricota* | *Howeltoviricetes* | *Cryppavirales* | *Mitoviridae* |  | *Nam-Co_genus_100* | blastp | *Nam-Co_Lenarviricota_108* | 91.7 |
| LC_20m_3um_k141_369318_1 | *Lenarviricota* | *Howeltoviricetes* | *Cryppavirales* | *Mitoviridae* |  | *Nam-Co_genus_109* | blastp | *Nam-Co_Lenarviricota_109* | 89.76 |
| LS_0.5m_1.2um_k141_350481_1 | *Lenarviricota* | *Howeltoviricetes* | *Cryppavirales* | *Mitoviridae* |  | *Nam-Co_genus_109* | blastp | *Nam-Co_Lenarviricota_110* | 91.3 |
| LC_coassembly_k141_1727227_1 | *Lenarviricota* | *Howeltoviricetes* | *Cryppavirales* | *Mitoviridae* |  | *Nam-Co_genus_093* | blastp | *Nam-Co_Lenarviricota_111* | 93.49 |
| LS_coassembly_k141_690095_1 | *Lenarviricota* | *Howeltoviricetes* | *Cryppavirales* | *Mitoviridae* |  |  | blastp | *Nam-Co_Lenarviricota_112* | 91.06 |
| LC_coassembly_k141_1486921_1 | *Lenarviricota* | *Howeltoviricetes* | *Cryppavirales* | *Mitoviridae* |  | *Nam-Co_genus_102* | blastp | *Nam-Co_Lenarviricota_113* | 89.42 |
| LC_coassembly_k141_2488396_1 | *Lenarviricota* | *Howeltoviricetes* | *Cryppavirales* | *Mitoviridae* |  |  | blastp | *Nam-Co_Lenarviricota_114* | 83.54 |
| LC_50m_10um_k141_274640_1 | *Lenarviricota* | *Howeltoviricetes* | *Cryppavirales* | *Mitoviridae* |  | *Nam-Co_genus_102* | blastp | *Nam-Co_Lenarviricota_115* | 87.67 |
| LC_coassembly_k141_2273513_1 | *Lenarviricota* | *Howeltoviricetes* | *Cryppavirales* | *Mitoviridae* |  | *Nam-Co_genus_111* | blastp | *Nam-Co_Lenarviricota_116* | 94.5 |
| LS_0.5m_1.2um_k141_285352_1 | *Lenarviricota* | *Howeltoviricetes* | *Cryppavirales* | *Mitoviridae* |  | *Nam-Co_genus_111* | blastp | *Nam-Co_Lenarviricota_117* | 90.58 |
| LC_coassembly_k141_213387_1 | *Lenarviricota* | *Howeltoviricetes* | *Cryppavirales* | *Mitoviridae* |  | *Nam-Co_genus_110* | blastp | *Nam-Co_Lenarviricota_118* | 90.08 |
| LC_0.5m_1.2um_k141_379937_1 | *Lenarviricota* | *Howeltoviricetes* | *Cryppavirales* | *Mitoviridae* |  | *Nam-Co_genus_110* | blastp | *Nam-Co_Lenarviricota_119* | 87.83 |
| LC_0.5m_1.2um_k141_232019_1 | *Lenarviricota* | *Howeltoviricetes* | *Cryppavirales* | *Mitoviridae* |  | *Nam-Co_genus_110* | blastp | *Nam-Co_Lenarviricota_120* | 86.67 |
| LC_0.5m_3um_k141_81056_1 | *Lenarviricota* | *Howeltoviricetes* | *Cryppavirales* | *Mitoviridae* |  | *Nam-Co_genus_110* | blastp | *Nam-Co_Lenarviricota_121* | 89.07 |
| LS_coassembly_k141_371343_1 | *Lenarviricota* | *Howeltoviricetes* | *Cryppavirales* | *Mitoviridae* |  | *Nam-Co_genus_110* | blastp | *Nam-Co_Lenarviricota_122* | 87.86 |
| LC_coassembly_k141_1897578_1 | *Lenarviricota* | *Howeltoviricetes* | *Cryppavirales* | *Mitoviridae* |  | *Nam-Co_genus_108* | blastp | *Nam-Co_Lenarviricota_123* | 92.98 |
| LC_coassembly_k141_2728316_1 | *Lenarviricota* | *Howeltoviricetes* | *Cryppavirales* | *Mitoviridae* |  | *Nam-Co_genus_092* | blastp | *Nam-Co_Lenarviricota_124* | 82.72 |
| LS_coassembly_k141_1106676_1 | *Lenarviricota* | *Howeltoviricetes* | *Cryppavirales* | *Mitoviridae* |  |  | blastp | *Nam-Co_Lenarviricota_125* | 92.25 |
| LS_coassembly_k141_582323_2 | *Lenarviricota* | *Howeltoviricetes* | *Cryppavirales* | *Mitoviridae* |  | *Nam-Co_genus_104* | blastp | *Nam-Co_Lenarviricota_126* | 92.79 |
| LC_coassembly_k141_63423_1 | *Lenarviricota* | *Howeltoviricetes* | *Cryppavirales* | *Mitoviridae* |  | *Nam-Co_genus_103* | blastp | *Nam-Co_Lenarviricota_127* | 88.78 |
| LC_80m_10um_k141_67259_1 | *Lenarviricota* | *Howeltoviricetes* | *Cryppavirales* | *Mitoviridae* |  | *Nam-Co_genus_103* | blastp | *Nam-Co_Lenarviricota_128* | 89.13 |
| LC_coassembly_k141_2070815_1 | *Lenarviricota* | *Howeltoviricetes* | *Cryppavirales* | *Mitoviridae* |  | *Nam-Co_genus_094* | blastp | *Nam-Co_Lenarviricota_129* | 86.9 |
| LS_coassembly_k141_880328_1 | *Lenarviricota* | *Howeltoviricetes* | *Cryppavirales* | *Mitoviridae* |  |  | blastp | *Nam-Co_Lenarviricota_130* | 92.27 |
| LS_coassembly_k141_182501_1 | *Lenarviricota* | *Howeltoviricetes* | *Cryppavirales* | *Mitoviridae* |  | *Nam-Co_genus_111* | blastp | *Nam-Co_Lenarviricota_131* | 93.39 |
| LC_50m_0.2um_k141_232096_1 | *Lenarviricota* | *Howeltoviricetes* | *Cryppavirales* | *Mitoviridae* |  | *Nam-Co_genus_107* | blastp | *Nam-Co_Lenarviricota_132* | 91.27 |
| LC_20m_3um_k141_118459_1 | *Lenarviricota* | *Howeltoviricetes* | *Cryppavirales* | *Mitoviridae* |  |  | blastp | *Nam-Co_Lenarviricota_133* | 88.75 |
| LS_0.5m_10um_k141_134516_1 | *Lenarviricota* | *Howeltoviricetes* | *Cryppavirales* | *Mitoviridae* |  |  | blastp | *Nam-Co_Lenarviricota_134* | 89.51 |
| LC_50m_10um_k141_55109_1 | *Lenarviricota* | *Howeltoviricetes* | *Cryppavirales* | *Mitoviridae* |  |  | blastp | *Nam-Co_Lenarviricota_135* | 89.47 |
| LC_coassembly_k141_961667_1 | *Lenarviricota* | *Howeltoviricetes* | *Cryppavirales* | *Mitoviridae* |  | *Nam-Co_genus_112* | blastp | *Nam-Co_Lenarviricota_136* | 86.76 |
| LC_coassembly_k141_242337_1 | *Lenarviricota* | *Howeltoviricetes* | *Cryppavirales* | *Mitoviridae* |  |  | blastp | *Nam-Co_Lenarviricota_137* | 84.99 |
| LC_coassembly_k141_398455_1 | *Lenarviricota* | *Howeltoviricetes* | *Cryppavirales* | *Mitoviridae* |  |  | blastp | *Nam-Co_Lenarviricota_138* | 84.88 |
| LC_0.5m_3um_k141_130993_1 | *Lenarviricota* | *Howeltoviricetes* | *Cryppavirales* | *Mitoviridae* |  |  | blastp | *Nam-Co_Lenarviricota_139* | 88.41 |
| LC_20m_10um_k141_19732_1 | *Lenarviricota* | *Howeltoviricetes* | *Cryppavirales* | *Mitoviridae* |  |  | blastp | *Nam-Co_Lenarviricota_140* | 89.18 |
| LC_20m_1.2um_k141_43823_1 | *Lenarviricota* | *Howeltoviricetes* | *Cryppavirales* | *Mitoviridae* |  |  | blastp | *Nam-Co_Lenarviricota_141* | 89.09 |
| LC_50m_10um_k141_221720_1 | *Lenarviricota* | *Howeltoviricetes* | *Cryppavirales* | *Mitoviridae* |  |  | blastp | *Nam-Co_Lenarviricota_142* | 85.52 |
| LS_coassembly_k141_777368_1 | *Lenarviricota* | *Howeltoviricetes* | *Cryppavirales* | *Mitoviridae* |  |  | blastp | *Nam-Co_Lenarviricota_143* | 89.44 |
| LS_coassembly_k141_851065_1 | *Lenarviricota* | *Howeltoviricetes* | *Cryppavirales* | *Mitoviridae* |  |  | blastp | *Nam-Co_Lenarviricota_144* | 88.34 |
| LS_0.5m_1.2um_k141_105741_1 | *Lenarviricota* | *Howeltoviricetes* | *Cryppavirales* | *Mitoviridae* |  |  | blastp | *Nam-Co_Lenarviricota_145* | 91.38 |
| LC_coassembly_k141_1170521_1 | *Lenarviricota* | *Amabiliviricetes* | *Wolframvirales* | *Narnaviridae* |  | *Nam-Co_genus_039* | blastp | *Nam-Co_Lenarviricota_146* | 91.55 |
| LC_20m_3um_k141_442215_1 | *Lenarviricota* | *Amabiliviricetes* | *Wolframvirales* | *Narnaviridae* |  |  | blastp | *Nam-Co_Lenarviricota_147* | 90.29 |
| LS_coassembly_k141_327859_1 | *Lenarviricota* | *Amabiliviricetes* | *Wolframvirales* | *Narnaviridae* |  | *Nam-Co_genus_022* | blastp | *Nam-Co_Lenarviricota_148* | 90.22 |
| LS_0.5m_3um_k141_409801_1 | *Lenarviricota* | *Amabiliviricetes* | *Wolframvirales* | *Narnaviridae* |  | *Nam-Co_genus_018* | blastp | *Nam-Co_Lenarviricota_149* | 87.93 |
| LC_50m_10um_k141_154418_1 | *Lenarviricota* | *Amabiliviricetes* | *Wolframvirales* | *Narnaviridae* | *Nam-Co_family_07* | *Nam-Co_genus_008* | blastp | *Nam-Co_Lenarviricota_150* | 91.99 |
| LS_coassembly_k141_1070766_1 | *Lenarviricota* | *Amabiliviricetes* | *Wolframvirales* | *Narnaviridae* |  | *Nam-Co_genus_062* | blastp | *Nam-Co_Lenarviricota_151* | 90.81 |
| LS_coassembly_k141_995692_1 | *Lenarviricota* | *Amabiliviricetes* | *Wolframvirales* | *Narnaviridae* |  | *Nam-Co_genus_041* | blastp | *Nam-Co_Lenarviricota_152* | 91.56 |
| LS_0.5m_1.2um_k141_104181_1 | *Lenarviricota* | *Amabiliviricetes* | *Wolframvirales* | *Narnaviridae* | *Nam-Co_family_09* | *Nam-Co_genus_031* | blastp | *Nam-Co_Lenarviricota_153* | 89.7 |
| LC_coassembly_k141_1754048_1 | *Lenarviricota* | *Amabiliviricetes* | *Wolframvirales* | *Narnaviridae* |  | *Nam-Co_genus_051* | blastp | *Nam-Co_Lenarviricota_154* | 88.92 |
| LC_0.5m_1.2um_k141_251615_2 | *Lenarviricota* | *Amabiliviricetes* | *Wolframvirales* | *Narnaviridae* |  | *Nam-Co_genus_054* | blastp | *Nam-Co_Lenarviricota_155* | 86.56 |
| LS_coassembly_k141_650154_1 | *Lenarviricota* | *Amabiliviricetes* | *Wolframvirales* | *Narnaviridae* |  | *Nam-Co_genus_056* | blastp | *Nam-Co_Lenarviricota_156* | 93.11 |
| LS_0.5m_1.2um_k141_563554_1 | *Lenarviricota* | *Amabiliviricetes* | *Wolframvirales* | *Narnaviridae* | *Nam-Co_family_13* | *Nam-Co_genus_064* | blastp | *Nam-Co_Lenarviricota_157* | 88.4 |
| LC_coassembly_k141_2261207_1 | *Lenarviricota* | *Amabiliviricetes* | *Wolframvirales* | *Narnaviridae* |  | *Nam-Co_genus_057* | blastp | *Nam-Co_Lenarviricota_158* | 92.92 |
| LC_coassembly_k141_1814031_1 | *Lenarviricota* | *Amabiliviricetes* | *Wolframvirales* | *Narnaviridae* |  | *Nam-Co_genus_046* | blastp | *Nam-Co_Lenarviricota_159* | 91.43 |
| LC_coassembly_k141_2638362_1 | *Lenarviricota* | *Amabiliviricetes* | *Wolframvirales* | *Narnaviridae* |  | *Nam-Co_genus_060* | blastp | *Nam-Co_Lenarviricota_160* | 88.75 |
| LC_0.5m_1.2um_k141_19237_1 | *Lenarviricota* | *Amabiliviricetes* | *Wolframvirales* | *Narnaviridae* |  | *Nam-Co_genus_050* | blastp | *Nam-Co_Lenarviricota_161* | 89.85 |
| LS_coassembly_k141_186661_1 | *Lenarviricota* | *Amabiliviricetes* | *Wolframvirales* | *Narnaviridae* |  | *Nam-Co_genus_055* | blastp | *Nam-Co_Lenarviricota_162* | 89.5 |
| LS_coassembly_k141_604560_1 | *Lenarviricota* | *Amabiliviricetes* | *Wolframvirales* | *Narnaviridae* |  | *Nam-Co_genus_030* | blastp | *Nam-Co_Lenarviricota_163* | 91.77 |
| LC_coassembly_k141_2368185_1 | *Lenarviricota* | *Amabiliviricetes* | *Wolframvirales* | *Narnaviridae* |  | *Nam-Co_genus_019* | blastp | *Nam-Co_Lenarviricota_164* | 90.47 |
| LS_coassembly_k141_1186789_1 | *Lenarviricota* | *Amabiliviricetes* | *Wolframvirales* | *Narnaviridae* |  | *Nam-Co_genus_010* | blastp | *Nam-Co_Lenarviricota_165* | 91.14 |
| LC_coassembly_k141_2261759_1 | *Lenarviricota* | *Amabiliviricetes* | *Wolframvirales* | *Narnaviridae* |  | *Nam-Co_genus_015* | blastp | *Nam-Co_Lenarviricota_166* | 90.45 |
| LC_coassembly_k141_1498943_1 | *Lenarviricota* | *Amabiliviricetes* | *Wolframvirales* | *Narnaviridae* |  | *Nam-Co_genus_016* | blastp | *Nam-Co_Lenarviricota_167* | 91.59 |
| LS_coassembly_k141_112006_1 | *Lenarviricota* | *Amabiliviricetes* | *Wolframvirales* | *Narnaviridae* |  | *Nam-Co_genus_005* | blastp | *Nam-Co_Lenarviricota_168* | 86.03 |
| LC_coassembly_k141_1544798_1 | *Lenarviricota* | *Amabiliviricetes* | *Wolframvirales* | *Narnaviridae* | *Nam-Co_family_12* | *Nam-Co_genus_047* | blastp | *Nam-Co_Lenarviricota_169* | 87.47 |
| LS_coassembly_k141_1328847_1 | *Lenarviricota* | *Amabiliviricetes* | *Wolframvirales* | *Narnaviridae* |  | *Nam-Co_genus_038* | blastp | *Nam-Co_Lenarviricota_170* | 92.49 |
| LS_coassembly_k141_1340706_1 | *Lenarviricota* | *Amabiliviricetes* | *Wolframvirales* | *Narnaviridae* |  | *Nam-Co_genus_020* | blastp | *Nam-Co_Lenarviricota_171* | 86.79 |
| LC_coassembly_k141_2586978_1 | *Lenarviricota* | *Amabiliviricetes* | *Wolframvirales* | *Narnaviridae* |  | *Nam-Co_genus_054* | blastp | *Nam-Co_Lenarviricota_172* | 87.02 |
| LS_coassembly_k141_118321_1 | *Lenarviricota* | *Amabiliviricetes* | *Wolframvirales* | *Narnaviridae* |  | *Nam-Co_genus_037* | blastp | *Nam-Co_Lenarviricota_173* | 94.38 |
| LC_coassembly_k141_1956227_1 | *Lenarviricota* | *Amabiliviricetes* | *Wolframvirales* | *Narnaviridae* | *Nam-Co_family_06* | *Nam-Co_genus_014* | blastp | *Nam-Co_Lenarviricota_174* | 87.88 |
| LS_coassembly_k141_969611_1 | *Lenarviricota* | *Amabiliviricetes* | *Wolframvirales* | *Narnaviridae* |  | *Nam-Co_genus_006* | blastp | *Nam-Co_Lenarviricota_175* | 91.88 |
| LC_coassembly_k141_2234358_1 | *Lenarviricota* | *Amabiliviricetes* | *Wolframvirales* | *Narnaviridae* |  | *Nam-Co_genus_065* | blastp | *Nam-Co_Lenarviricota_176* | 92.38 |
| LC_coassembly_k141_19190_1 | *Lenarviricota* | *Amabiliviricetes* | *Wolframvirales* | *Narnaviridae* | *Nam-Co_family_07* | *Nam-Co_genus_008* | blastp | *Nam-Co_Lenarviricota_177* | 91.74 |
| LC_0.5m_1.2um_k141_98739_2 | *Lenarviricota* | *Amabiliviricetes* | *Wolframvirales* | *Narnaviridae* |  | *Nam-Co_genus_050* | blastp | *Nam-Co_Lenarviricota_178* | 91.67 |
| LS_coassembly_k141_1156677_1 | *Lenarviricota* | *Amabiliviricetes* | *Wolframvirales* | *Narnaviridae* |  |  | blastp | *Nam-Co_Lenarviricota_179* | 91.19 |
| LC_coassembly_k141_1036729_1 | *Lenarviricota* | *Amabiliviricetes* | *Wolframvirales* | *Narnaviridae* |  |  | blastp | *Nam-Co_Lenarviricota_180* | 91.81 |
| LS_coassembly_k141_814387_2 | *Lenarviricota* | *Amabiliviricetes* | *Wolframvirales* | *Narnaviridae* |  |  | blastp | *Nam-Co_Lenarviricota_181* | 91.68 |
| LC_coassembly_k141_2292250_1 | *Lenarviricota* | *Amabiliviricetes* | *Wolframvirales* | *Narnaviridae* |  |  | blastp | *Nam-Co_Lenarviricota_182* | 91.18 |
| LS_coassembly_k141_192514_1 | *Lenarviricota* | *Amabiliviricetes* | *Wolframvirales* | *Narnaviridae* |  | *Nam-Co_genus_011* | blastp | *Nam-Co_Lenarviricota_183* | 82.23 |
| LS_coassembly_k141_1350286_1 | *Lenarviricota* | *Amabiliviricetes* | *Wolframvirales* | *Narnaviridae* | *Nam-Co_family_10* | *Nam-Co_genus_033* | blastp | *Nam-Co_Lenarviricota_184* | 92.99 |
| LC_coassembly_k141_1001983_1 | *Lenarviricota* | *Amabiliviricetes* | *Wolframvirales* | *Narnaviridae* |  |  | blastp | *Nam-Co_Lenarviricota_185* | 88.39 |
| LS_coassembly_k141_647081_1 | *Lenarviricota* | *Amabiliviricetes* | *Wolframvirales* | *Narnaviridae* |  | *Nam-Co_genus_040* | blastp | *Nam-Co_Lenarviricota_186* | 87.71 |
| LS_coassembly_k141_368163_1 | *Lenarviricota* | *Amabiliviricetes* | *Wolframvirales* | *Narnaviridae* |  |  | blastp | *Nam-Co_Lenarviricota_187* | 88.88 |
| LS_0.5m_1.2um_k141_567195_2 | *Lenarviricota* | *Amabiliviricetes* | *Wolframvirales* | *Narnaviridae* |  | *Nam-Co_genus_048* | blastp | *Nam-Co_Lenarviricota_188* | 87.07 |
| LC_coassembly_k141_2489571_1 | *Lenarviricota* | *Amabiliviricetes* | *Wolframvirales* | *Narnaviridae* | *Nam-Co_family_11* | *Nam-Co_genus_012* | blastp | *Nam-Co_Lenarviricota_189* |  |
| LC_20m_10um_k141_76613_1 | *Lenarviricota* | *Amabiliviricetes* | *Wolframvirales* | *Narnaviridae* | *Nam-Co_family_07* | *Nam-Co_genus_008* | blastp | *Nam-Co_Lenarviricota_190* | 91.81 |
| LS_coassembly_k141_345578_1 | *Lenarviricota* | *Amabiliviricetes* | *Wolframvirales* | *Narnaviridae* |  | *Nam-Co_genus_024* | blastp | *Nam-Co_Lenarviricota_191* | 85.24 |
| LC_coassembly_k141_2383019_1 | *Lenarviricota* | *Amabiliviricetes* | *Wolframvirales* | *Narnaviridae* |  | *Nam-Co_genus_017* | blastp | *Nam-Co_Lenarviricota_192* | 89.2 |
| LS_coassembly_k141_483032_1 | *Lenarviricota* | *Amabiliviricetes* | *Wolframvirales* | *Narnaviridae* | *Nam-Co_family_08* | *Nam-Co_genus_042* | blastp | *Nam-Co_Lenarviricota_193* | 86.66 |
| LC_coassembly_k141_635520_1 | *Lenarviricota* | *Amabiliviricetes* | *Wolframvirales* | *Narnaviridae* | *Nam-Co_family_13* | *Nam-Co_genus_043* | blastp | *Nam-Co_Lenarviricota_194* | 92.1 |
| LC_coassembly_k141_775494_1 | *Lenarviricota* | *Amabiliviricetes* | *Wolframvirales* | *Narnaviridae* |  | *Nam-Co_genus_032* | blastp | *Nam-Co_Lenarviricota_195* | 87.97 |
| LS_coassembly_k141_73853_1 | *Lenarviricota* | *Amabiliviricetes* | *Wolframvirales* | *Narnaviridae* |  | *Nam-Co_genus_055* | blastp | *Nam-Co_Lenarviricota_196* | 87.88 |
| LC_20m_0.2um_k141_367775_1 | *Lenarviricota* | *Amabiliviricetes* | *Wolframvirales* | *Narnaviridae* |  | *Nam-Co_genus_009* | blastp | *Nam-Co_Lenarviricota_197* | 85.78 |
| LC_20m_1.2um_k141_4741_2 | *Lenarviricota* | *Amabiliviricetes* | *Wolframvirales* | *Narnaviridae* |  | *Nam-Co_genus_009* | blastp | *Nam-Co_Lenarviricota_198* | 84.77 |
| LC_50m_1.2um_k141_37463_1 | *Lenarviricota* | *Amabiliviricetes* | *Wolframvirales* | *Narnaviridae* |  | *Nam-Co_genus_009* | blastp | *Nam-Co_Lenarviricota_199* | 84.04 |
| LC_80m_3um_k141_157262_2 | *Lenarviricota* | *Amabiliviricetes* | *Wolframvirales* | *Narnaviridae* |  | *Nam-Co_genus_009* | blastp | *Nam-Co_Lenarviricota_200* | 85.08 |
| LC_0.5m_10um_k141_178784_1 | *Lenarviricota* | *Amabiliviricetes* | *Wolframvirales* | *Narnaviridae* |  |  | blastp | *Nam-Co_Lenarviricota_201* | 92.26 |
| LS_coassembly_k141_575813_1 | *Lenarviricota* | *Amabiliviricetes* | *Wolframvirales* | *Narnaviridae* |  |  | blastp | *Nam-Co_Lenarviricota_202* | 88.89 |
| LS_coassembly_k141_677412_1 | *Lenarviricota* | *Amabiliviricetes* | *Wolframvirales* | *Narnaviridae* |  | *Nam-Co_genus_025* | blastp | *Nam-Co_Lenarviricota_203* | 91.3 |
| LS_coassembly_k141_186086_1 | *Lenarviricota* | *Amabiliviricetes* | *Wolframvirales* | *Narnaviridae* |  | *Nam-Co_genus_044* | blastp | *Nam-Co_Lenarviricota_204* | 86.67 |
| LC_coassembly_k141_431617_1 | *Lenarviricota* | *Amabiliviricetes* | *Wolframvirales* | *Narnaviridae* | *Nam-Co_family_05* | *Nam-Co_genus_036* | blastp | *Nam-Co_Lenarviricota_205* | 88.16 |
| LC_80m_0.2um_k141_579292_1 | *Lenarviricota* | *Amabiliviricetes* | *Wolframvirales* | *Narnaviridae* |  | *Nam-Co_genus_026* | blastp | *Nam-Co_Lenarviricota_206* | 91.1 |
| LS_coassembly_k141_965508_1 | *Lenarviricota* | *Amabiliviricetes* | *Wolframvirales* | *Narnaviridae* |  |  | blastp | *Nam-Co_Lenarviricota_207* | 92.73 |
| LS_coassembly_k141_70220_1 | *Lenarviricota* | *Amabiliviricetes* | *Wolframvirales* | *Narnaviridae* | *Nam-Co_family_13* | *Nam-Co_genus_064* | blastp | *Nam-Co_Lenarviricota_208* | 88.61 |
| LC_coassembly_k141_1350414_1 | *Lenarviricota* | *Amabiliviricetes* | *Wolframvirales* | *Narnaviridae* |  | *Nam-Co_genus_061* | blastp | *Nam-Co_Lenarviricota_209* | 90.62 |
| LS_coassembly_k141_300460_1 | *Lenarviricota* | *Amabiliviricetes* | *Wolframvirales* | *Narnaviridae* |  | *Nam-Co_genus_023* | blastp | *Nam-Co_Lenarviricota_210* | 92.17 |
| LS_coassembly_k141_19643_2 | *Lenarviricota* | *Amabiliviricetes* | *Wolframvirales* | *Narnaviridae* |  | *Nam-Co_genus_052* | blastp | *Nam-Co_Lenarviricota_211* | 89.75 |
| LS_coassembly_k141_1109689_1 | *Lenarviricota* | *Amabiliviricetes* | *Wolframvirales* | *Narnaviridae* |  | *Nam-Co_genus_063* | blastp | *Nam-Co_Lenarviricota_212* | 83.45 |
| LC_coassembly_k141_1396795_2 | *Lenarviricota* | *Amabiliviricetes* | *Wolframvirales* | *Narnaviridae* |  | *Nam-Co_genus_053* | blastp | *Nam-Co_Lenarviricota_213* | 94.61 |
| LC_coassembly_k141_2371561_1 | *Lenarviricota* | *Amabiliviricetes* | *Wolframvirales* | *Narnaviridae* |  | *Nam-Co_genus_049* | blastp | *Nam-Co_Lenarviricota_214* | 89.68 |
| LC_coassembly_k141_1828284_1 | *Lenarviricota* | *Amabiliviricetes* | *Wolframvirales* | *Narnaviridae* |  |  | blastp | *Nam-Co_Lenarviricota_215* | 90.17 |
| LC_coassembly_k141_659737_1 | *Lenarviricota* | *Amabiliviricetes* | *Wolframvirales* | *Narnaviridae* |  | *Nam-Co_genus_059* | blastp | *Nam-Co_Lenarviricota_216* | 86.65 |
| LS_coassembly_k141_174926_1 | *Lenarviricota* | *Amabiliviricetes* | *Wolframvirales* | *Narnaviridae* |  |  | blastp | *Nam-Co_Lenarviricota_217* | 92.07 |
| LS_0.5m_1.2um_k141_450766_1 | *Lenarviricota* | *Amabiliviricetes* | *Wolframvirales* | *Narnaviridae* |  | *Nam-Co_genus_028* | blastp | *Nam-Co_Lenarviricota_218* | 89.18 |
| LC_coassembly_k141_1120492_2 | *Lenarviricota* | *Amabiliviricetes* | *Wolframvirales* | *Narnaviridae* |  | *Nam-Co_genus_013* | blastp | *Nam-Co_Lenarviricota_219* | 94.53 |
| LS_coassembly_k141_1032564_1 | *Lenarviricota* | *Amabiliviricetes* | *Wolframvirales* | *Narnaviridae* |  |  | blastp | *Nam-Co_Lenarviricota_220* | 92.36 |
| LC_coassembly_k141_781199_1 | *Lenarviricota* | *Amabiliviricetes* | *Wolframvirales* | *Narnaviridae* |  | *Nam-Co_genus_007* | blastp | *Nam-Co_Lenarviricota_221* | 89.11 |
| LC_coassembly_k141_502336_1 | *Lenarviricota* | *Amabiliviricetes* | *Wolframvirales* | *Narnaviridae* |  | *Nam-Co_genus_026* | blastp | *Nam-Co_Lenarviricota_222* | 90.77 |
| LC_80m_3um_k141_194697_1 | *Lenarviricota* | *Amabiliviricetes* | *Wolframvirales* | *Narnaviridae* |  | *Nam-Co_genus_026* | blastp | *Nam-Co_Lenarviricota_223* | 90.63 |
| LC_0.5m_3um_k141_257773_1 | *Lenarviricota* | *Amabiliviricetes* | *Wolframvirales* | *Narnaviridae* |  | *Nam-Co_genus_009* | blastp | *Nam-Co_Lenarviricota_224* | 84.78 |
| LC_50m_10um_k141_296581_1 | *Lenarviricota* | *Amabiliviricetes* | *Wolframvirales* | *Narnaviridae* |  | *Nam-Co_genus_009* | blastp | *Nam-Co_Lenarviricota_225* | 84.37 |
| LC_80m_1.2um_k141_246966_1 | *Lenarviricota* | *Amabiliviricetes* | *Wolframvirales* | *Narnaviridae* |  | *Nam-Co_genus_009* | blastp | *Nam-Co_Lenarviricota_226* | 85.47 |
| LS_coassembly_k141_1278143_1 | *Lenarviricota* | *Amabiliviricetes* | *Wolframvirales* | *Narnaviridae* |  | *Nam-Co_genus_021* | blastp | *Nam-Co_Lenarviricota_227* | 94.34 |
| LS_coassembly_k141_881219_1 | *Lenarviricota* | *Amabiliviricetes* | *Wolframvirales* | *Narnaviridae* |  | *Nam-Co_genus_034* | blastp | *Nam-Co_Lenarviricota_228* | 91.33 |
| LS_0.5m_0.2um_k141_234728_1 | *Lenarviricota* | *Amabiliviricetes* | *Wolframvirales* | *Narnaviridae* |  | *Nam-Co_genus_018* | blastp | *Nam-Co_Lenarviricota_229* | 87.49 |
| LC_50m_3um_k141_487618_1 | *Lenarviricota* | *Amabiliviricetes* | *Wolframvirales* | *Narnaviridae* | *Nam-Co_family_07* | *Nam-Co_genus_008* | blastp | *Nam-Co_Lenarviricota_230* | 89.49 |
| LC_coassembly_k141_1773347_1 | *Lenarviricota* | *Amabiliviricetes* | *Wolframvirales* | *Narnaviridae* |  | *Nam-Co_genus_029* | blastp | *Nam-Co_Lenarviricota_231* | 90.05 |
| LS_coassembly_k141_1023795_1 | *Lenarviricota* | *Amabiliviricetes* | *Wolframvirales* | *Narnaviridae* |  | *Nam-Co_genus_045* | blastp | *Nam-Co_Lenarviricota_232* | 95.88 |
| LC_coassembly_k141_147022_1 | *Lenarviricota* | *Amabiliviricetes* | *Wolframvirales* | *Narnaviridae* |  | *Nam-Co_genus_058* | blastp | *Nam-Co_Lenarviricota_233* | 92.13 |
| LS_coassembly_k141_1328102_1 | *Lenarviricota* | *Amabiliviricetes* | *Wolframvirales* | *Narnaviridae* |  | *Nam-Co_genus_027* | blastp | *Nam-Co_Lenarviricota_234* | 87.21 |
| LS_coassembly_k141_1124569_1 | *Lenarviricota* | *Amabiliviricetes* | *Wolframvirales* | *Narnaviridae* |  | *Nam-Co_genus_038* | blastp | *Nam-Co_Lenarviricota_235* | 93.51 |
| LC_coassembly_k141_934910_1 | *Lenarviricota* | *Amabiliviricetes* | *Wolframvirales* | *Narnaviridae* |  | *Nam-Co_genus_035* | blastp | *Nam-Co_Lenarviricota_236* | 91.3 |
| LS_coassembly_k141_4481_1 | *Lenarviricota* | *Leviviricetes* | *Norzivirales* | *Solspiviridae* |  | *Nam-Co_genus_132* | blastp | *Nam-Co_Lenarviricota_237* | 96.85 |
| LS_coassembly_k141_631699_1 | *Lenarviricota* | *Leviviricetes* | *Timlovirales* | *Steitzviridae* |  | *Nam-Co_genus_135* | blastp | *Nam-Co_Lenarviricota_238* | 96.04 |
| LS_coassembly_k141_1166806_2 | *Lenarviricota* | *Leviviricetes* | *Timlovirales* | *Steitzviridae* |  | *Nam-Co_genus_140* | blastp | *Nam-Co_Lenarviricota_239* | 95.8 |
| LS_coassembly_k141_542386_1 | *Lenarviricota* | *Leviviricetes* | *Timlovirales* | *Steitzviridae* |  |  | blastp | *Nam-Co_Lenarviricota_240* | 97.18 |
| LS_coassembly_k141_638690_2 | *Lenarviricota* | *Leviviricetes* | *Timlovirales* | *Steitzviridae* |  | *Nam-Co_genus_134* | blastp | *Nam-Co_Lenarviricota_241* | 96.83 |
| LC_coassembly_k141_2541618_1 | *Lenarviricota* | *Leviviricetes* | *Timlovirales* | *Steitzviridae* |  | *Nam-Co_genus_138* | blastp | *Nam-Co_Lenarviricota_242* | 95.67 |
| LC_coassembly_k141_635711_1 | *Lenarviricota* | *Leviviricetes* | *Timlovirales* | *Steitzviridae* |  | *Nam-Co_genus_139* | blastp | *Nam-Co_Lenarviricota_243* | 95.06 |
| LS_coassembly_k141_824362_1 | *Lenarviricota* | *Leviviricetes* | *Timlovirales* | *Steitzviridae* |  | *Nam-Co_genus_136* | blastp | *Nam-Co_Lenarviricota_244* | 96.32 |
| LC_coassembly_k141_1271871_1 | *Lenarviricota* | *Leviviricetes* | *Timlovirales* | *Steitzviridae* |  |  | blastp | *Nam-Co_Lenarviricota_245* | 96.68 |
| LS_coassembly_k141_152525_1 | *Lenarviricota* | *Leviviricetes* | *Timlovirales* | *Steitzviridae* |  |  | blastp | *Nam-Co_Lenarviricota_246* | 91.76 |
| LS_coassembly_k141_370211_1 | *Lenarviricota* | *Leviviricetes* | *Timlovirales* | *Steitzviridae* |  | *Nam-Co_genus_133* | blastp | *Nam-Co_Lenarviricota_247* | 95.63 |
| LS_coassembly_k141_839853_2 | *Lenarviricota* | *Leviviricetes* | *Timlovirales* | *Steitzviridae* |  | *Nam-Co_genus_137* | blastp | *Nam-Co_Lenarviricota_248* | 96.5 |
| LC_20m_3um_k141_267098_1 | *Pisuviricota* |  |  |  | *Nam-Co_family_24* | *Nam-Co_genus_153* | structure | *Nam-Co_Pisuviricota_001* | 79.14 |
| LC_coassembly_k141_1176970_4 | *Pisuviricota* |  |  |  | *Nam-Co_family_26* | *Nam-Co_genus_156* | structure | *Nam-Co_Pisuviricota_002* | 92.59 |
| LC_coassembly_k141_788326_1 | *Pisuviricota* |  |  |  | *Nam-Co_family_25* | *Nam-Co_genus_154* | structure | *Nam-Co_Pisuviricota_003* | 79.23 |
| LC_coassembly_k141_1892157_1 | *Pisuviricota* |  |  |  | *Nam-Co_family_22* | *Nam-Co_genus_157* | structure | *Nam-Co_Pisuviricota_004* | 84.58 |
| LS_coassembly_k141_547291_1 | *Pisuviricota* |  |  |  | *Nam-Co_family_21* | *Nam-Co_genus_152* | structure | *Nam-Co_Pisuviricota_005* | 61.69 |
| LS_coassembly_k141_374349_1 | *Pisuviricota* |  |  |  | *Nam-Co_family_20* | *Nam-Co_genus_151* | structure | *Nam-Co_Pisuviricota_006* | 80.81 |
| LS_0.5m_1.2um_k141_172551_1 | *Pisuviricota* |  |  |  | *Nam-Co_family_23* | *Nam-Co_genus_155* | structure | *Nam-Co_Pisuviricota_007* | 72.89 |
| LC_coassembly_k141_2206238_2 | *Pisuviricota* | *Duplopiviricetes* | *Durnavirales* | *Amalgaviridae* | *Nam-Co_family_36* | *Nam-Co_genus_184* | blastp | *Nam-Co_Pisuviricota_008* | 88.7 |
| LS_coassembly_k141_908476_1 | *Pisuviricota* | *Stelpaviricetes* | *Stellavirales* | *Astroviridae* |  | *Nam-Co_genus_255* | blastp | *Nam-Co_Pisuviricota_009* | 90.83 |
| LS_coassembly_k141_705223_2 | *Pisuviricota* | *Stelpaviricetes* | *Stellavirales* | *Astroviridae* |  | *Nam-Co_genus_254* | blastp | *Nam-Co_Pisuviricota_010* | 93.33 |
| LC_coassembly_k141_1848390_2 | *Pisuviricota* | *Stelpaviricetes* | *Stellavirales* | *Astroviridae* |  | *Nam-Co_genus_253* | blastp | *Nam-Co_Pisuviricota_011* | 90.72 |
| LS_0.5m_1.2um_k141_674011_1 | *Pisuviricota* | *c.0012* | *Sobelivirales* | *Barnaviridae* |  |  | blastp | *Nam-Co_Pisuviricota_012* | 95.1 |
| LS_coassembly_k141_677142_1 | *Pisuviricota* | *Pisoniviricetes* | *Picornavirales* | *Dicistroviridae* |  |  | blastp | *Nam-Co_Pisuviricota_013* | 93.39 |
| LC_coassembly_k141_2139485_1 | *Pisuviricota* | *Pisoniviricetes* | *Picornavirales* | *Dicistroviridae* |  | *Nam-Co_genus_226* | blastp | *Nam-Co_Pisuviricota_014* | 93.8 |
| LS_coassembly_k141_611362_2 | *Pisuviricota* | *Pisoniviricetes* | *Picornavirales* | *Dicistroviridae* |  |  | blastp | *Nam-Co_Pisuviricota_015* | 90.46 |
| LC_coassembly_k141_1539147_1 | *Pisuviricota* | *Pisoniviricetes* | *Picornavirales* | *Dicistroviridae* |  |  | blastp | *Nam-Co_Pisuviricota_016* | 93.96 |
| LC_coassembly_k141_2670924_1 | *Pisuviricota* | *Pisoniviricetes* | *Picornavirales* | *Dicistroviridae* |  |  | blastp | *Nam-Co_Pisuviricota_017* | 94.91 |
| LS_coassembly_k141_268503_1 | *Pisuviricota* | *Pisoniviricetes* | *Picornavirales* | *Dicistroviridae* |  |  | blastp | *Nam-Co_Pisuviricota_018* | 94.81 |
| LS_coassembly_k141_625058_1 | *Pisuviricota* | *Pisoniviricetes* | *Picornavirales* | *Dicistroviridae* |  | *Nam-Co_genus_225* | blastp | *Nam-Co_Pisuviricota_019* | 94.01 |
| LC_80m_3um_k141_208137_1 | *Pisuviricota* | *Pisoniviricetes* | *Picornavirales* | *Dicistroviridae* |  |  | blastp | *Nam-Co_Pisuviricota_020* | 92.93 |
| LS_coassembly_k141_734239_1 | *Pisuviricota* | *Pisoniviricetes* | *Picornavirales* | *Dicistroviridae* |  |  | blastp | *Nam-Co_Pisuviricota_021* | 94.1 |
| LS_coassembly_k141_882900_1 | *Pisuviricota* | *Pisoniviricetes* | *Picornavirales* | *f.0008* |  |  | blastp | *Nam-Co_Pisuviricota_022* | 89.47 |
| LS_coassembly_k141_434311_1 | *Pisuviricota* | *Pisoniviricetes* | *Picornavirales* | *f.0008* |  | *Nam-Co_genus_227* | blastp | *Nam-Co_Pisuviricota_023* | 95.99 |
| LC_coassembly_k141_415235_1 | *Pisuviricota* | *Pisoniviricetes* | *Picornavirales* | *f.0008* |  |  | blastp | *Nam-Co_Pisuviricota_024* | 92.3 |
| LS_coassembly_k141_673941_1 | *Pisuviricota* | *Pisoniviricetes* | *Picornavirales* | *f.0024* |  | *Nam-Co_genus_228* | blastp | *Nam-Co_Pisuviricota_025* | 93.83 |
| LC_coassembly_k141_51246_1 | *Pisuviricota* | *Pisoniviricetes* | *Picornavirales* | *f.0025* |  | *Nam-Co_genus_229* | blastp | *Nam-Co_Pisuviricota_026* | 92.51 |
| LC_80m_3um_k141_52117_2 | *Pisuviricota* | *Pisoniviricetes* | *Picornavirales* | *f.0025* |  | *Nam-Co_genus_229* | blastp | *Nam-Co_Pisuviricota_027* | 93.24 |
| LC_coassembly_k141_365724_1 | *Pisuviricota* | *Pisoniviricetes* | *Picornavirales* | *f.0025* |  | *Nam-Co_genus_229* | blastp | *Nam-Co_Pisuviricota_028* | 92.27 |
| LC_80m_3um_k141_145146_1 | *Pisuviricota* | *Pisoniviricetes* | *Picornavirales* | *f.0025* |  | *Nam-Co_genus_229* | blastp | *Nam-Co_Pisuviricota_029* | 92.07 |
| LC_coassembly_k141_2427120_1 | *Pisuviricota* | *Pisoniviricetes* | *Picornavirales* | *f.0028* |  | *Nam-Co_genus_230* | blastp | *Nam-Co_Pisuviricota_030* | 92.83 |
| LC_coassembly_k141_737588_1 | *Pisuviricota* | *Pisoniviricetes* | *Picornavirales* | *f.0029* |  |  | blastp | *Nam-Co_Pisuviricota_031* | 94.02 |
| LS_coassembly_k141_559789_1 | *Pisuviricota* | *Pisoniviricetes* | *Picornavirales* | *f.0030* |  | *Nam-Co_genus_232* | blastp | *Nam-Co_Pisuviricota_032* | 92.12 |
| LC_coassembly_k141_682946_1 | *Pisuviricota* | *Pisoniviricetes* | *Picornavirales* | *f.0030* |  | *Nam-Co_genus_231* | blastp | *Nam-Co_Pisuviricota_033* | 93 |
| LC_coassembly_k141_2048581_1 | *Pisuviricota* | *Pisoniviricetes* | *Picornavirales* | *f.0030* |  | *Nam-Co_genus_232* | blastp | *Nam-Co_Pisuviricota_034* | 92.43 |
| LS_coassembly_k141_790503_1 | *Pisuviricota* | *Pisoniviricetes* | *Picornavirales* | *f.0033* |  |  | blastp | *Nam-Co_Pisuviricota_035* | 93.95 |
| LC_coassembly_k141_1091995_1 | *Pisuviricota* | *Pisoniviricetes* | *Picornavirales* | *f.0040* |  | *Nam-Co_genus_233* | blastp | *Nam-Co_Pisuviricota_036* | 92.28 |
| LC_coassembly_k141_2414581_1 | *Pisuviricota* | *Pisoniviricetes* | *Picornavirales* | *f.0042* | *Nam-Co_family_45* | *Nam-Co_genus_235* | blastp | *Nam-Co_Pisuviricota_037* | 91.95 |
| LC_coassembly_k141_567048_2 | *Pisuviricota* | *Pisoniviricetes* | *Picornavirales* | *f.0042* |  | *Nam-Co_genus_234* | blastp | *Nam-Co_Pisuviricota_038* | 90.84 |
| LC_coassembly_k141_1606298_1 | *Pisuviricota* | *Pisoniviricetes* | *Picornavirales* | *f.0044* | *Nam-Co_family_44* | *Nam-Co_genus_236* | blastp | *Nam-Co_Pisuviricota_039* | 87.04 |
| LS_coassembly_k141_303746_1 | *Pisuviricota* | *Pisoniviricetes* | *Picornavirales* | *f.0048* |  | *Nam-Co_genus_237* | blastp | *Nam-Co_Pisuviricota_040* | 91.29 |
| LC_50m_0.2um_k141_198770_2 | *Pisuviricota* | *c.0003.base-Pisoni* | *o.0003* | *f.0056* | *Nam-Co_family_27* | *Nam-Co_genus_158* | blastp | *Nam-Co_Pisuviricota_041* | 70.05 |
| LC_coassembly_k141_1686962_1 | *Pisuviricota* | *c.0003.base-Pisoni* | *o.0003* | *f.0056* | *Nam-Co_family_27* | *Nam-Co_genus_158* | blastp | *Nam-Co_Pisuviricota_042* | 63 |
| LC_coassembly_k141_1998157_1 | *Pisuviricota* | *c.0013* | *o.0025.base-Sobeli* | *f.0076* |  | *Nam-Co_genus_183* | blastp | *Nam-Co_Pisuviricota_043* | 80.16 |
| LC_coassembly_k141_964946_1 | *Pisuviricota* | *c.0013* | *o.0025.base-Sobeli* | *f.0076* |  |  | blastp | *Nam-Co_Pisuviricota_044* | 92.94 |
| LC_coassembly_k141_1030321_1 | *Pisuviricota* | *c.0013* | *o.0025.base-Sobeli* | *f.0076* |  | *Nam-Co_genus_172* | blastp | *Nam-Co_Pisuviricota_045* | 89.28 |
| LS_coassembly_k141_296363_1 | *Pisuviricota* | *c.0013* | *o.0025.base-Sobeli* | *f.0076* |  | *Nam-Co_genus_167* | blastp | *Nam-Co_Pisuviricota_046* | 92.35 |
| LS_coassembly_k141_127221_1 | *Pisuviricota* | *c.0013* | *o.0025.base-Sobeli* | *f.0076* |  | *Nam-Co_genus_168* | blastp | *Nam-Co_Pisuviricota_047* | 94.29 |
| LC_20m_0.2um_k141_332893_1 | *Pisuviricota* | *c.0013* | *o.0025.base-Sobeli* | *f.0076* |  | *Nam-Co_genus_170* | blastp | *Nam-Co_Pisuviricota_048* | 94.5 |
| LS_coassembly_k141_613763_1 | *Pisuviricota* | *c.0013* | *o.0025.base-Sobeli* | *f.0076* |  | *Nam-Co_genus_167* | blastp | *Nam-Co_Pisuviricota_049* | 90.9 |
| LC_coassembly_k141_1841635_1 | *Pisuviricota* | *c.0013* | *o.0025.base-Sobeli* | *f.0076* |  | *Nam-Co_genus_182* | blastp | *Nam-Co_Pisuviricota_050* | 90.65 |
| LC_coassembly_k141_2512221_1 | *Pisuviricota* | *c.0013* | *o.0025.base-Sobeli* | *f.0076* |  | *Nam-Co_genus_166* | blastp | *Nam-Co_Pisuviricota_051* | 93.48 |
| LC_coassembly_k141_421224_1 | *Pisuviricota* | *c.0013* | *o.0025.base-Sobeli* | *f.0076* |  |  | blastp | *Nam-Co_Pisuviricota_052* | 89.43 |
| LS_coassembly_k141_108479_1 | *Pisuviricota* | *c.0013* | *o.0025.base-Sobeli* | *f.0076* |  |  | blastp | *Nam-Co_Pisuviricota_053* | 92.31 |
| LS_0.5m_3um_k141_97861_1 | *Pisuviricota* | *c.0013* | *o.0025.base-Sobeli* | *f.0076* |  |  | blastp | *Nam-Co_Pisuviricota_054* | 92.42 |
| LC_coassembly_k141_133542_1 | *Pisuviricota* | *c.0013* | *o.0025.base-Sobeli* | *f.0076* | *Nam-Co_family_29* | *Nam-Co_genus_178* | blastp | *Nam-Co_Pisuviricota_055* | 89.06 |
| LS_coassembly_k141_1081653_1 | *Pisuviricota* | *c.0013* | *o.0025.base-Sobeli* | *f.0076* |  | *Nam-Co_genus_175* | blastp | *Nam-Co_Pisuviricota_056* | 91.94 |
| LS_coassembly_k141_873293_1 | *Pisuviricota* | *c.0013* | *o.0025.base-Sobeli* | *f.0076* |  | *Nam-Co_genus_171* | blastp | *Nam-Co_Pisuviricota_057* | 93.97 |
| LS_coassembly_k141_370863_1 | *Pisuviricota* | *c.0013* | *o.0025.base-Sobeli* | *f.0076* |  | *Nam-Co_genus_169* | blastp | *Nam-Co_Pisuviricota_058* | 89.72 |
| LC_coassembly_k141_137237_1 | *Pisuviricota* | *c.0013* | *o.0025.base-Sobeli* | *f.0076* |  | *Nam-Co_genus_179* | blastp | *Nam-Co_Pisuviricota_059* | 92.54 |
| LS_coassembly_k141_659713_1 | *Pisuviricota* | *c.0013* | *o.0025.base-Sobeli* | *f.0076* |  | *Nam-Co_genus_179* | blastp | *Nam-Co_Pisuviricota_060* | 91.22 |
| LC_coassembly_k141_1872912_1 | *Pisuviricota* | *c.0013* | *o.0025.base-Sobeli* | *f.0076* | *Nam-Co_family_29* | *Nam-Co_genus_178* | blastp | *Nam-Co_Pisuviricota_061* | 89.81 |
| LC_80m_0.2um_k141_138777_1 | *Pisuviricota* | *c.0013* | *o.0025.base-Sobeli* | *f.0076* |  | *Nam-Co_genus_180* | blastp | *Nam-Co_Pisuviricota_062* | 85.12 |
| LS_coassembly_k141_1174751_1 | *Pisuviricota* | *c.0013* | *o.0025.base-Sobeli* | *f.0076* |  | *Nam-Co_genus_170* | blastp | *Nam-Co_Pisuviricota_063* | 92.94 |
| LC_coassembly_k141_2612247_1 | *Pisuviricota* | *c.0013* | *o.0025.base-Sobeli* | *f.0076* |  | *Nam-Co_genus_181* | blastp | *Nam-Co_Pisuviricota_064* | 90.62 |
| LC_coassembly_k141_51051_1 | *Pisuviricota* | *c.0013* | *o.0025.base-Sobeli* | *f.0076* |  |  | blastp | *Nam-Co_Pisuviricota_065* | 92.4 |
| LS_coassembly_k141_357820_1 | *Pisuviricota* | *c.0013* | *o.0025.base-Sobeli* | *f.0076* |  |  | blastp | *Nam-Co_Pisuviricota_066* | 89.82 |
| LC_coassembly_k141_2450679_1 | *Pisuviricota* | *c.0013* | *o.0025.base-Sobeli* | *f.0076* |  | *Nam-Co_genus_182* | blastp | *Nam-Co_Pisuviricota_067* | 94.22 |
| LS_coassembly_k141_1002104_1 | *Pisuviricota* | *c.0013* | *o.0025.base-Sobeli* | *f.0076* |  | *Nam-Co_genus_174* | blastp | *Nam-Co_Pisuviricota_068* | 92.57 |
| LC_coassembly_k141_2084125_1 | *Pisuviricota* | *c.0013* | *o.0025.base-Sobeli* | *f.0076* |  | *Nam-Co_genus_173* | blastp | *Nam-Co_Pisuviricota_069* | 86.62 |
| LC_coassembly_k141_568262_1 | *Pisuviricota* | *c.0013* | *o.0025.base-Sobeli* | *f.0076* |  | *Nam-Co_genus_176* | blastp | *Nam-Co_Pisuviricota_070* | 92.94 |
| LS_coassembly_k141_126965_1 | *Pisuviricota* | *c.0013* | *o.0025.base-Sobeli* | *f.0076* |  |  | blastp | *Nam-Co_Pisuviricota_071* | 92.55 |
| LC_coassembly_k141_217665_1 | *Pisuviricota* | *c.0013* | *o.0025.base-Sobeli* | *f.0076* |  | *Nam-Co_genus_177* | blastp | *Nam-Co_Pisuviricota_072* | 85.76 |
| LC_50m_10um_k141_212090_2 | *Pisuviricota* | *Duplopiviricetes* | *Durnavirales* | *f.0102* |  | *Nam-Co_genus_188* | blastp | *Nam-Co_Pisuviricota_073* | 80.31 |
| LC_coassembly_k141_943468_1 | *Pisuviricota* | *Duplopiviricetes* | *Durnavirales* | *f.0102* |  |  | blastp | *Nam-Co_Pisuviricota_074* | 83.37 |
| LC_coassembly_k141_526161_2 | *Pisuviricota* | *Duplopiviricetes* | *Durnavirales* | *f.0102* |  |  | blastp | *Nam-Co_Pisuviricota_075* | 87.6 |
| LS_coassembly_k141_1054548_1 | *Pisuviricota* | *Duplopiviricetes* | *Durnavirales* | *f.0102* |  | *Nam-Co_genus_195* | blastp | *Nam-Co_Pisuviricota_076* | 85.31 |
| LC_50m_10um_k141_136548_1 | *Pisuviricota* | *Duplopiviricetes* | *Durnavirales* | *f.0102* |  |  | blastp | *Nam-Co_Pisuviricota_077* | 87 |
| LS_coassembly_k141_812161_1 | *Pisuviricota* | *Duplopiviricetes* | *Durnavirales* | *f.0102* |  |  | blastp | *Nam-Co_Pisuviricota_078* | 88.78 |
| LS_coassembly_k141_986866_2 | *Pisuviricota* | *Duplopiviricetes* | *Durnavirales* | *f.0102* | *Nam-Co_family_33* | *Nam-Co_genus_202* | blastp | *Nam-Co_Pisuviricota_079* | 62.17 |
| LC_20m_3um_k141_551912_1 | *Pisuviricota* | *Duplopiviricetes* | *Durnavirales* | *f.0102* |  |  | blastp | *Nam-Co_Pisuviricota_080* | 88.48 |
| LC_20m_10um_k141_70765_2 | *Pisuviricota* | *Duplopiviricetes* | *Durnavirales* | *f.0102* | *Nam-Co_family_32* | *Nam-Co_genus_190* | blastp | *Nam-Co_Pisuviricota_081* | 89.65 |
| LC_coassembly_k141_16449_2 | *Pisuviricota* | *Duplopiviricetes* | *Durnavirales* | *f.0102* |  |  | blastp | *Nam-Co_Pisuviricota_082* | 88.28 |
| LC_coassembly_k141_744534_1 | *Pisuviricota* | *Duplopiviricetes* | *Durnavirales* | *f.0102* |  |  | blastp | *Nam-Co_Pisuviricota_083* | 89.44 |
| LC_20m_3um_k141_236703_1 | *Pisuviricota* | *Duplopiviricetes* | *Durnavirales* | *f.0102* |  | *Nam-Co_genus_203* | blastp | *Nam-Co_Pisuviricota_084* | 86.18 |
| LC_80m_3um_k141_3455_1 | *Pisuviricota* | *Duplopiviricetes* | *Durnavirales* | *f.0102* |  | *Nam-Co_genus_203* | blastp | *Nam-Co_Pisuviricota_085* | 87.53 |
| LS_coassembly_k141_405215_1 | *Pisuviricota* | *Duplopiviricetes* | *Durnavirales* | *f.0102* |  | *Nam-Co_genus_185* | blastp | *Nam-Co_Pisuviricota_086* | 82.13 |
| LC_coassembly_k141_947088_2 | *Pisuviricota* | *Duplopiviricetes* | *Durnavirales* | *f.0102* |  |  | blastp | *Nam-Co_Pisuviricota_087* | 87.44 |
| LS_coassembly_k141_153349_1 | *Pisuviricota* | *Duplopiviricetes* | *Durnavirales* | *f.0102* |  | *Nam-Co_genus_197* | blastp | *Nam-Co_Pisuviricota_088* | 83.46 |
| LC_coassembly_k141_700243_1 | *Pisuviricota* | *Duplopiviricetes* | *Durnavirales* | *f.0102* |  | *Nam-Co_genus_203* | blastp | *Nam-Co_Pisuviricota_089* | 87.24 |
| LC_coassembly_k141_570098_1 | *Pisuviricota* | *Duplopiviricetes* | *Durnavirales* | *f.0102* |  | *Nam-Co_genus_189* | blastp | *Nam-Co_Pisuviricota_090* | 88.02 |
| LC_coassembly_k141_367052_1 | *Pisuviricota* | *Duplopiviricetes* | *Durnavirales* | *f.0102* |  | *Nam-Co_genus_204* | blastp | *Nam-Co_Pisuviricota_091* | 84.93 |
| LC_coassembly_k141_619132_2 | *Pisuviricota* | *Duplopiviricetes* | *Durnavirales* | *f.0102* | *Nam-Co_family_40* | *Nam-Co_genus_187* | blastp | *Nam-Co_Pisuviricota_092* | 85.55 |
| LS_coassembly_k141_947059_1 | *Pisuviricota* | *Duplopiviricetes* | *Durnavirales* | *f.0102* |  | *Nam-Co_genus_193* | blastp | *Nam-Co_Pisuviricota_093* | 84.04 |
| LS_coassembly_k141_165614_2 | *Pisuviricota* | *Duplopiviricetes* | *Durnavirales* | *f.0102* |  |  | blastp | *Nam-Co_Pisuviricota_094* | 86.6 |
| LS_coassembly_k141_969233_1 | *Pisuviricota* | *Duplopiviricetes* | *Durnavirales* | *f.0102* |  | *Nam-Co_genus_191* | blastp | *Nam-Co_Pisuviricota_095* | 76.4 |
| LC_coassembly_k141_727479_2 | *Pisuviricota* | *Duplopiviricetes* | *Durnavirales* | *f.0102* |  | *Nam-Co_genus_192* | blastp | *Nam-Co_Pisuviricota_096* | 88.54 |
| LC_coassembly_k141_2464698_2 | *Pisuviricota* | *Duplopiviricetes* | *Durnavirales* | *f.0102* |  | *Nam-Co_genus_188* | blastp | *Nam-Co_Pisuviricota_097* | 83.37 |
| LC_coassembly_k141_1739317_1 | *Pisuviricota* | *Duplopiviricetes* | *Durnavirales* | *f.0102* |  | *Nam-Co_genus_186* | blastp | *Nam-Co_Pisuviricota_098* | 85.73 |
| LC_0.5m_10um_k141_201724_1 | *Pisuviricota* | *Duplopiviricetes* | *Durnavirales* | *f.0102* |  | *Nam-Co_genus_186* | blastp | *Nam-Co_Pisuviricota_099* | 85.52 |
| LC_coassembly_k141_1186528_1 | *Pisuviricota* | *Duplopiviricetes* | *Durnavirales* | *f.0102* |  | *Nam-Co_genus_201* | blastp | *Nam-Co_Pisuviricota_100* | 86.36 |
| LC_0.5m_10um_k141_138773_1 | *Pisuviricota* | *Duplopiviricetes* | *Durnavirales* | *f.0102* |  | *Nam-Co_genus_201* | blastp | *Nam-Co_Pisuviricota_101* | 86.5 |
| LS_coassembly_k141_1002961_1 | *Pisuviricota* | *Duplopiviricetes* | *Durnavirales* | *f.0102* |  | *Nam-Co_genus_199* | blastp | *Nam-Co_Pisuviricota_102* | 88.75 |
| LS_0.5m_10um_k141_209189_1 | *Pisuviricota* | *Duplopiviricetes* | *Durnavirales* | *f.0102* |  | *Nam-Co_genus_191* | blastp | *Nam-Co_Pisuviricota_103* | 82.27 |
| LS_coassembly_k141_1274756_1 | *Pisuviricota* | *Duplopiviricetes* | *Durnavirales* | *f.0102* |  | *Nam-Co_genus_200* | blastp | *Nam-Co_Pisuviricota_104* | 89.64 |
| LS_0.5m_10um_k141_96668_1 | *Pisuviricota* | *Duplopiviricetes* | *Durnavirales* | *f.0102* | *Nam-Co_family_34* | *Nam-Co_genus_196* | blastp | *Nam-Co_Pisuviricota_105* | 85.89 |
| LC_coassembly_k141_101176_3 | *Pisuviricota* | *Duplopiviricetes* | *Durnavirales* | *f.0102* |  | *Nam-Co_genus_203* | blastp | *Nam-Co_Pisuviricota_106* | 86.71 |
| LC_0.5m_10um_k141_30436_2 | *Pisuviricota* | *Duplopiviricetes* | *Durnavirales* | *f.0102* |  | *Nam-Co_genus_203* | blastp | *Nam-Co_Pisuviricota_107* | 87.32 |
| LC_50m_3um_k141_385349_4 | *Pisuviricota* | *Duplopiviricetes* | *Durnavirales* | *f.0102* |  | *Nam-Co_genus_203* | blastp | *Nam-Co_Pisuviricota_108* | 86.78 |
| LS_coassembly_k141_292708_1 | *Pisuviricota* | *Duplopiviricetes* | *Durnavirales* | *f.0102* |  | *Nam-Co_genus_194* | blastp | *Nam-Co_Pisuviricota_109* | 80.14 |
| LS_0.5m_10um_k141_53807_1 | *Pisuviricota* | *Duplopiviricetes* | *Durnavirales* | *f.0102* |  | *Nam-Co_genus_194* | blastp | *Nam-Co_Pisuviricota_110* | 80.89 |
| LS_coassembly_k141_455458_1 | *Pisuviricota* | *Duplopiviricetes* | *Durnavirales* | *f.0102* | *Nam-Co_family_30* | *Nam-Co_genus_198* | blastp | *Nam-Co_Pisuviricota_111* | 88.96 |
| LC_coassembly_k141_2501884_2 | *Pisuviricota* | *Duplopiviricetes* | *Durnavirales* | *f.0102* |  |  | blastp | *Nam-Co_Pisuviricota_112* | 86.17 |
| LC_coassembly_k141_592818_2 | *Pisuviricota* | *Duplopiviricetes* | *Durnavirales* | *f.0102* |  | *Nam-Co_genus_188* | blastp | *Nam-Co_Pisuviricota_113* | 79.27 |
| LS_coassembly_k141_747216_1 | *Pisuviricota* | *Duplopiviricetes* | *Durnavirales* | *f.0102* |  |  | blastp | *Nam-Co_Pisuviricota_114* | 86.4 |
| LC_coassembly_k141_1421816_1 | *Pisuviricota* | *Duplopiviricetes* | *Durnavirales* | *f.0102* |  | *Nam-Co_genus_201* | blastp | *Nam-Co_Pisuviricota_115* | 88.31 |
| LC_coassembly_k141_589296_1 | *Pisuviricota* | *Duplopiviricetes* | *Durnavirales* | *f.0102* |  | *Nam-Co_genus_201* | blastp | *Nam-Co_Pisuviricota_116* | 87.68 |
| LC_coassembly_k141_1675587_1 | *Pisuviricota* | *Duplopiviricetes* | *Durnavirales* | *f.0102* |  | *Nam-Co_genus_201* | blastp | *Nam-Co_Pisuviricota_117* | 86.11 |
| LS_coassembly_k141_232625_1 | *Pisuviricota* | *Duplopiviricetes* | *Durnavirales* | *f.0102* |  |  | blastp | *Nam-Co_Pisuviricota_118* | 88.27 |
| LC_0.5m_3um_k141_167927_1 | *Pisuviricota* | *Duplopiviricetes* | *Durnavirales* | *f.0111* | *Nam-Co_family_39* | *Nam-Co_genus_212* | blastp | *Nam-Co_Pisuviricota_119* | 89.92 |
| LC_coassembly_k141_686723_1 | *Pisuviricota* | *Duplopiviricetes* | *Durnavirales* | *f.0111* |  | *Nam-Co_genus_207* | blastp | *Nam-Co_Pisuviricota_120* | 87.21 |
| LC_coassembly_k141_2759094_1 | *Pisuviricota* | *Duplopiviricetes* | *Durnavirales* | *f.0111* |  | *Nam-Co_genus_207* | blastp | *Nam-Co_Pisuviricota_121* | 87.94 |
| LC_50m_10um_k141_273261_1 | *Pisuviricota* | *Duplopiviricetes* | *Durnavirales* | *f.0111* |  | *Nam-Co_genus_207* | blastp | *Nam-Co_Pisuviricota_122* | 87.15 |
| LC_50m_3um_k141_450784_1 | *Pisuviricota* | *Duplopiviricetes* | *Durnavirales* | *f.0111* |  | *Nam-Co_genus_207* | blastp | *Nam-Co_Pisuviricota_123* | 88.13 |
| LC_coassembly_k141_1831437_1 | *Pisuviricota* | *Duplopiviricetes* | *Durnavirales* | *f.0111* |  |  | blastp | *Nam-Co_Pisuviricota_124* | 88.92 |
| LC_coassembly_k141_1578211_1 | *Pisuviricota* | *Duplopiviricetes* | *Durnavirales* | *f.0111* | *Nam-Co_family_35* | *Nam-Co_genus_205* | blastp | *Nam-Co_Pisuviricota_125* | 84.7 |
| LC_coassembly_k141_633441_1 | *Pisuviricota* | *Duplopiviricetes* | *Durnavirales* | *f.0111* | *Nam-Co_family_39* | *Nam-Co_genus_212* | blastp | *Nam-Co_Pisuviricota_126* | 89.21 |
| LC_50m_3um_k141_109718_1 | *Pisuviricota* | *Duplopiviricetes* | *Durnavirales* | *f.0111* | *Nam-Co_family_39* | *Nam-Co_genus_212* | blastp | *Nam-Co_Pisuviricota_127* | 89.52 |
| LS_coassembly_k141_370511_1 | *Pisuviricota* | *Duplopiviricetes* | *Durnavirales* | *f.0111* | *Nam-Co_family_39* | *Nam-Co_genus_212* | blastp | *Nam-Co_Pisuviricota_128* | 90.68 |
| LC_80m_3um_k141_98302_1 | *Pisuviricota* | *Duplopiviricetes* | *Durnavirales* | *f.0111* |  |  | blastp | *Nam-Co_Pisuviricota_129* | 84.07 |
| LC_coassembly_k141_601624_1 | *Pisuviricota* | *Duplopiviricetes* | *Durnavirales* | *f.0111* |  | *Nam-Co_genus_213* | blastp | *Nam-Co_Pisuviricota_130* | 89.44 |
| LC_0.5m_0.2um_k141_166103_1 | *Pisuviricota* | *Duplopiviricetes* | *Durnavirales* | *f.0111* |  |  | blastp | *Nam-Co_Pisuviricota_131* | 92.48 |
| LC_0.5m_10um_k141_147631_1 | *Pisuviricota* | *Duplopiviricetes* | *Durnavirales* | *f.0111* |  | *Nam-Co_genus_211* | blastp | *Nam-Co_Pisuviricota_132* | 91.08 |
| LC_0.5m_3um_k141_194855_1 | *Pisuviricota* | *Duplopiviricetes* | *Durnavirales* | *f.0111* |  | *Nam-Co_genus_211* | blastp | *Nam-Co_Pisuviricota_133* | 91.04 |
| LC_80m_3um_k141_184180_1 | *Pisuviricota* | *Duplopiviricetes* | *Durnavirales* | *f.0111* |  | *Nam-Co_genus_211* | blastp | *Nam-Co_Pisuviricota_134* | 91.23 |
| LS_coassembly_k141_406676_1 | *Pisuviricota* | *Duplopiviricetes* | *Durnavirales* | *f.0111* | *Nam-Co_family_38* | *Nam-Co_genus_209* | blastp | *Nam-Co_Pisuviricota_135* | 87 |
| LS_0.5m_3um_k141_161033_1 | *Pisuviricota* | *Duplopiviricetes* | *Durnavirales* | *f.0111* |  | *Nam-Co_genus_211* | blastp | *Nam-Co_Pisuviricota_136* | 91.31 |
| LC_coassembly_k141_1484123_1 | *Pisuviricota* | *Duplopiviricetes* | *Durnavirales* | *f.0111* |  | *Nam-Co_genus_206* | blastp | *Nam-Co_Pisuviricota_137* | 86.51 |
| LC_coassembly_k141_672667_1 | *Pisuviricota* | *Duplopiviricetes* | *Durnavirales* | *f.0111* |  | *Nam-Co_genus_213* | blastp | *Nam-Co_Pisuviricota_138* | 90.02 |
| LC_coassembly_k141_1185643_1 | *Pisuviricota* | *Duplopiviricetes* | *Durnavirales* | *f.0111* |  | *Nam-Co_genus_213* | blastp | *Nam-Co_Pisuviricota_139* | 89.77 |
| LC_50m_10um_k141_242393_1 | *Pisuviricota* | *Duplopiviricetes* | *Durnavirales* | *f.0111* |  | *Nam-Co_genus_213* | blastp | *Nam-Co_Pisuviricota_140* | 88.74 |
| LS_0.5m_3um_k141_94350_1 | *Pisuviricota* | *Duplopiviricetes* | *Durnavirales* | *f.0111* |  | *Nam-Co_genus_213* | blastp | *Nam-Co_Pisuviricota_141* | 90.83 |
| LC_coassembly_k141_1198831_1 | *Pisuviricota* | *Duplopiviricetes* | *Durnavirales* | *f.0111* |  |  | blastp | *Nam-Co_Pisuviricota_142* | 92.17 |
| LC_50m_3um_k141_71475_1 | *Pisuviricota* | *Duplopiviricetes* | *Durnavirales* | *f.0111* |  | *Nam-Co_genus_213* | blastp | *Nam-Co_Pisuviricota_143* | 84.57 |
| LC_coassembly_k141_2771384_1 | *Pisuviricota* | *Duplopiviricetes* | *Durnavirales* | *f.0111* |  |  | blastp | *Nam-Co_Pisuviricota_144* | 93.06 |
| LC_coassembly_k141_995284_1 | *Pisuviricota* | *Duplopiviricetes* | *Durnavirales* | *f.0111* |  | *Nam-Co_genus_211* | blastp | *Nam-Co_Pisuviricota_145* | 92.01 |
| LC_coassembly_k141_1148484_1 | *Pisuviricota* | *Duplopiviricetes* | *Durnavirales* | *f.0111* |  | *Nam-Co_genus_211* | blastp | *Nam-Co_Pisuviricota_146* | 91.03 |
| LC_coassembly_k141_497225_1 | *Pisuviricota* | *Duplopiviricetes* | *Durnavirales* | *f.0111* |  | *Nam-Co_genus_211* | blastp | *Nam-Co_Pisuviricota_147* | 91.64 |
| LC_50m_3um_k141_295552_1 | *Pisuviricota* | *Duplopiviricetes* | *Durnavirales* | *f.0111* |  | *Nam-Co_genus_211* | blastp | *Nam-Co_Pisuviricota_148* | 91.31 |
| LC_coassembly_k141_2642329_1 | *Pisuviricota* | *Duplopiviricetes* | *Durnavirales* | *f.0111* |  | *Nam-Co_genus_211* | blastp | *Nam-Co_Pisuviricota_149* | 90.35 |
| LC_50m_10um_k141_103028_1 | *Pisuviricota* | *Duplopiviricetes* | *Durnavirales* | *f.0111* |  | *Nam-Co_genus_213* | blastp | *Nam-Co_Pisuviricota_150* | 84.41 |
| LS_coassembly_k141_420880_1 | *Pisuviricota* | *Duplopiviricetes* | *Durnavirales* | *f.0111* | *Nam-Co_family_37* | *Nam-Co_genus_208* | blastp | *Nam-Co_Pisuviricota_151* | 87.23 |
| LC_coassembly_k141_629875_1 | *Pisuviricota* | *Duplopiviricetes* | *Durnavirales* | *f.0111* |  | *Nam-Co_genus_211* | blastp | *Nam-Co_Pisuviricota_152* | 90.81 |
| LC_50m_10um_k141_244857_1 | *Pisuviricota* | *Duplopiviricetes* | *Durnavirales* | *f.0111* |  | *Nam-Co_genus_211* | blastp | *Nam-Co_Pisuviricota_153* | 89.05 |
| LC_coassembly_k141_893532_1 | *Pisuviricota* | *Duplopiviricetes* | *Durnavirales* | *f.0111* | *Nam-Co_family_38* | *Nam-Co_genus_210* | blastp | *Nam-Co_Pisuviricota_154* | 79.55 |
| LC_50m_10um_k141_298364_1 | *Pisuviricota* | *Duplopiviricetes* | *Durnavirales* | *f.0111* |  | *Nam-Co_genus_213* | blastp | *Nam-Co_Pisuviricota_155* | 89.67 |
| LS_coassembly_k141_689206_1 | *Pisuviricota* | *Duplopiviricetes* | *Durnavirales* | *f.0111* | *Nam-Co_family_39* | *Nam-Co_genus_212* | blastp | *Nam-Co_Pisuviricota_156* | 93.38 |
| LC_coassembly_k141_1689211_1 | *Pisuviricota* | *Duplopiviricetes* | *Durnavirales* | *f.0111* | *Nam-Co_family_39* | *Nam-Co_genus_212* | blastp | *Nam-Co_Pisuviricota_157* | 90.73 |
| LC_coassembly_k141_1369234_1 | *Pisuviricota* | *Duplopiviricetes* | *Durnavirales* | *f.0111* | *Nam-Co_family_39* | *Nam-Co_genus_212* | blastp | *Nam-Co_Pisuviricota_158* | 91.39 |
| LC_20m_3um_k141_508020_1 | *Pisuviricota* | *Duplopiviricetes* | *Durnavirales* | *f.0111* | *Nam-Co_family_39* | *Nam-Co_genus_212* | blastp | *Nam-Co_Pisuviricota_159* | 91.2 |
| LS_coassembly_k141_35229_1 | *Pisuviricota* | *Duplopiviricetes* | *Durnavirales* | *f.0111* | *Nam-Co_family_39* | *Nam-Co_genus_212* | blastp | *Nam-Co_Pisuviricota_160* | 90.48 |
| LC_coassembly_k141_2545102_1 | *Pisuviricota* | *Duplopiviricetes* | *Durnavirales* | *f.0111* |  |  | blastp | *Nam-Co_Pisuviricota_161* | 84.77 |
| LS_coassembly_k141_481010_1 | *Pisuviricota* | *Duplopiviricetes* | *Durnavirales* | *f.0111* |  | *Nam-Co_genus_211* | blastp | *Nam-Co_Pisuviricota_162* | 90.83 |
| LC_coassembly_k141_1811975_1 | *Pisuviricota* | *Duplopiviricetes* | *Durnavirales* | *f.0120* |  | *Nam-Co_genus_214* | blastp | *Nam-Co_Pisuviricota_163* | 90.57 |
| LS_coassembly_k141_408106_1 | *Pisuviricota* | *Flasuviricetes* | *Amarillovirales* | *Flaviviridae* |  | *Nam-Co_genus_224* | blastp | *Nam-Co_Pisuviricota_164* | 94.23 |
| LC_coassembly_k141_1296706_1 | *Pisuviricota* | *c.0005.base-Stelpa* | *o.0012* | *Hypoviridae* |  | *Nam-Co_genus_160* | blastp | *Nam-Co_Pisuviricota_165* | 91.93 |
| LC_coassembly_k141_138224_1 | *Pisuviricota* | *c.0005.base-Stelpa* | *o.0012* | *Hypoviridae* |  | *Nam-Co_genus_159* | blastp | *Nam-Co_Pisuviricota_166* | 88.7 |
| LS_coassembly_k141_187532_1 | *Pisuviricota* | *Pisoniviricetes* | *Picornavirales* | *Iflaviridae* | *Nam-Co_family_41* | *Nam-Co_genus_238* | blastp | *Nam-Co_Pisuviricota_167* | 93.86 |
| LS_coassembly_k141_77005_1 | *Pisuviricota* | *Pisoniviricetes* | *Picornavirales* | *Marnaviridae* |  | *Nam-Co_genus_241* | blastp | *Nam-Co_Pisuviricota_168* | 93.45 |
| LC_coassembly_k141_2733097_1 | *Pisuviricota* | *Pisoniviricetes* | *Picornavirales* | *Marnaviridae* |  |  | blastp | *Nam-Co_Pisuviricota_169* | 95.02 |
| LS_coassembly_k141_666330_1 | *Pisuviricota* | *Pisoniviricetes* | *Picornavirales* | *Marnaviridae* | *Nam-Co_family_46* | *Nam-Co_genus_250* | blastp | *Nam-Co_Pisuviricota_170* | 90 |
| LS_0.5m_1.2um_k141_174384_1 | *Pisuviricota* | *Pisoniviricetes* | *Picornavirales* | *Marnaviridae* |  | *Nam-Co_genus_243* | blastp | *Nam-Co_Pisuviricota_171* | 92.99 |
| LC_coassembly_k141_2733507_1 | *Pisuviricota* | *Pisoniviricetes* | *Picornavirales* | *Marnaviridae* |  | *Nam-Co_genus_248* | blastp | *Nam-Co_Pisuviricota_172* | 93.25 |
| LC_coassembly_k141_1335197_1 | *Pisuviricota* | *Pisoniviricetes* | *Picornavirales* | *Marnaviridae* |  |  | blastp | *Nam-Co_Pisuviricota_173* | 95.28 |
| LS_coassembly_k141_1097885_1 | *Pisuviricota* | *Pisoniviricetes* | *Picornavirales* | *Marnaviridae* |  |  | blastp | *Nam-Co_Pisuviricota_174* | 93.49 |
| LS_coassembly_k141_741592_1 | *Pisuviricota* | *Pisoniviricetes* | *Picornavirales* | *Marnaviridae* |  |  | blastp | *Nam-Co_Pisuviricota_175* | 94.52 |
| LC_coassembly_k141_2042949_1 | *Pisuviricota* | *Pisoniviricetes* | *Picornavirales* | *Marnaviridae* |  |  | blastp | *Nam-Co_Pisuviricota_176* | 94.23 |
| LS_0.5m_3um_k141_29737_1 | *Pisuviricota* | *Pisoniviricetes* | *Picornavirales* | *Marnaviridae* |  | *Nam-Co_genus_247* | blastp | *Nam-Co_Pisuviricota_177* | 92.91 |
| LC_coassembly_k141_473258_1 | *Pisuviricota* | *Pisoniviricetes* | *Picornavirales* | *Marnaviridae* |  | *Nam-Co_genus_245* | blastp | *Nam-Co_Pisuviricota_178* | 93.95 |
| LC_coassembly_k141_1231733_1 | *Pisuviricota* | *Pisoniviricetes* | *Picornavirales* | *Marnaviridae* |  | *Nam-Co_genus_244* | blastp | *Nam-Co_Pisuviricota_179* | 94.94 |
| LS_coassembly_k141_1194284_1 | *Pisuviricota* | *Pisoniviricetes* | *Picornavirales* | *Marnaviridae* |  | *Nam-Co_genus_239* | blastp | *Nam-Co_Pisuviricota_180* | 91.33 |
| LC_20m_0.2um_k141_315972_1 | *Pisuviricota* | *Pisoniviricetes* | *Picornavirales* | *Marnaviridae* |  |  | blastp | *Nam-Co_Pisuviricota_181* | 93.54 |
| LS_coassembly_k141_178385_1 | *Pisuviricota* | *Pisoniviricetes* | *Picornavirales* | *Marnaviridae* |  |  | blastp | *Nam-Co_Pisuviricota_182* | 91.91 |
| LC_coassembly_k141_1474085_1 | *Pisuviricota* | *Pisoniviricetes* | *Picornavirales* | *Marnaviridae* |  |  | blastp | *Nam-Co_Pisuviricota_183* | 93.36 |
| LC_coassembly_k141_2285013_1 | *Pisuviricota* | *Pisoniviricetes* | *Picornavirales* | *Marnaviridae* | *Nam-Co_family_43* | *Nam-Co_genus_242* | blastp | *Nam-Co_Pisuviricota_184* | 91.03 |
| LC_coassembly_k141_2239335_1 | *Pisuviricota* | *Pisoniviricetes* | *Picornavirales* | *Marnaviridae* |  |  | blastp | *Nam-Co_Pisuviricota_185* | 95.82 |
| LC_20m_0.2um_k141_308140_2 | *Pisuviricota* | *Pisoniviricetes* | *Picornavirales* | *Marnaviridae* |  | *Nam-Co_genus_245* | blastp | *Nam-Co_Pisuviricota_186* | 94.86 |
| LC_20m_0.2um_k141_250544_1 | *Pisuviricota* | *Pisoniviricetes* | *Picornavirales* | *Marnaviridae* |  | *Nam-Co_genus_249* | blastp | *Nam-Co_Pisuviricota_187* | 93.45 |
| LS_coassembly_k141_897578_1 | *Pisuviricota* | *Pisoniviricetes* | *Picornavirales* | *Marnaviridae* |  | *Nam-Co_genus_246* | blastp | *Nam-Co_Pisuviricota_188* | 92.7 |
| LS_coassembly_k141_190706_1 | *Pisuviricota* | *Pisoniviricetes* | *Picornavirales* | *Marnaviridae* |  | *Nam-Co_genus_247* | blastp | *Nam-Co_Pisuviricota_189* | 92.33 |
| LC_coassembly_k141_916411_1 | *Pisuviricota* | *Pisoniviricetes* | *Picornavirales* | *Marnaviridae* |  | *Nam-Co_genus_240* | blastp | *Nam-Co_Pisuviricota_190* | 86.74 |
| LC_coassembly_k141_2145116_1 | *Pisuviricota* | *Pisoniviricetes* | *Picornavirales* | *Marnaviridae* |  |  | blastp | *Nam-Co_Pisuviricota_191* | 95.21 |
| LC_coassembly_k141_1432413_1 | *Pisuviricota* | *Pisoniviricetes* | *Picornavirales* | *Marnaviridae* |  |  | blastp | *Nam-Co_Pisuviricota_192* | 92.98 |
| LS_coassembly_k141_18378_1 | *Pisuviricota* | *Duplopiviricetes* | *Durnavirales* | *Partitiviridae* |  | *Nam-Co_genus_215* | blastp | *Nam-Co_Pisuviricota_193* | 87.49 |
| LC_coassembly_k141_689124_1 | *Pisuviricota* | *Duplopiviricetes* | *Durnavirales* | *Partitiviridae* |  | *Nam-Co_genus_217* | blastp | *Nam-Co_Pisuviricota_194* | 87.64 |
| LS_coassembly_k141_125234_1 | *Pisuviricota* | *Duplopiviricetes* | *Durnavirales* | *Partitiviridae* |  |  | blastp | *Nam-Co_Pisuviricota_195* | 90.76 |
| LS_coassembly_k141_126227_1 | *Pisuviricota* | *Duplopiviricetes* | *Durnavirales* | *Partitiviridae* |  | *Nam-Co_genus_219* | blastp | *Nam-Co_Pisuviricota_196* | 93.63 |
| LC_coassembly_k141_2580908_1 | *Pisuviricota* | *Duplopiviricetes* | *Durnavirales* | *Partitiviridae* |  | *Nam-Co_genus_222* | blastp | *Nam-Co_Pisuviricota_197* | 92.84 |
| LS_coassembly_k141_161594_1 | *Pisuviricota* | *Duplopiviricetes* | *Durnavirales* | *Partitiviridae* |  | *Nam-Co_genus_216* | blastp | *Nam-Co_Pisuviricota_198* | 90.42 |
| LS_coassembly_k141_774987_1 | *Pisuviricota* | *Duplopiviricetes* | *Durnavirales* | *Partitiviridae* |  | *Nam-Co_genus_218* | blastp | *Nam-Co_Pisuviricota_199* | 92.2 |
| LS_coassembly_k141_740060_1 | *Pisuviricota* | *Duplopiviricetes* | *Durnavirales* | *Partitiviridae* |  | *Nam-Co_genus_221* | blastp | *Nam-Co_Pisuviricota_200* | 87.92 |
| LC_coassembly_k141_2755829_1 | *Pisuviricota* | *Duplopiviricetes* | *Durnavirales* | *Partitiviridae* | *Nam-Co_family_31* | *Nam-Co_genus_220* | blastp | *Nam-Co_Pisuviricota_201* | 87.4 |
| LC_80m_3um_k141_197063_1 | *Pisuviricota* | *Duplopiviricetes* | *Durnavirales* | *Partitiviridae* | *Nam-Co_family_31* | *Nam-Co_genus_220* | blastp | *Nam-Co_Pisuviricota_202* | 87.35 |
| LC_coassembly_k141_868598_1 | *Pisuviricota* | *Duplopiviricetes* | *Durnavirales* | *Picobirnaviridae* |  | *Nam-Co_genus_223* | blastp | *Nam-Co_Pisuviricota_203* | 94.21 |
| LC_coassembly_k141_1394068_1 | *Pisuviricota* | *Pisoniviricetes* | *Picornavirales* | *Picornaviridae* |  |  | blastp | *Nam-Co_Pisuviricota_204* | 96.92 |
| LS_coassembly_k141_818259_5 | *Pisuviricota* | *Pisoniviricetes* | *Picornavirales* | *Polycipiviridae* |  |  | blastp | *Nam-Co_Pisuviricota_205* | 94.15 |
| LS_coassembly_k141_226167_1 | *Pisuviricota* | *Pisoniviricetes* | *Picornavirales* | *Polycipiviridae* |  |  | blastp | *Nam-Co_Pisuviricota_206* | 93.88 |
| LC_coassembly_k141_2634150_1 | *Pisuviricota* | *Pisoniviricetes* | *Picornavirales* | *Polycipiviridae* |  | *Nam-Co_genus_251* | blastp | *Nam-Co_Pisuviricota_207* | 94.48 |
| LS_coassembly_k141_409312_3 | *Pisuviricota* | *Pisoniviricetes* | *Picornavirales* | *Polycipiviridae* | *Nam-Co_family_42* | *Nam-Co_genus_252* | blastp | *Nam-Co_Pisuviricota_208* | 88.59 |
| LS_coassembly_k141_795198_1 | *Pisuviricota* | *Stelpaviricetes* | *Patatavirales* | *Potyviridae* |  |  | blastp | *Nam-Co_Pisuviricota_209* | 92.14 |
| LC_coassembly_k141_2159845_2 | *Pisuviricota* | *c.0012* | *Sobelivirales* | *Solemoviridae* |  |  | blastp | *Nam-Co_Pisuviricota_210* | 94.63 |
| LS_coassembly_k141_389387_2 | *Pisuviricota* | *c.0012* | *Sobelivirales* | *Solemoviridae* |  |  | blastp | *Nam-Co_Pisuviricota_211* | 94.19 |
| LC_coassembly_k141_86982_2 | *Pisuviricota* | *c.0012* | *Sobelivirales* | *Solemoviridae* |  | *Nam-Co_genus_164* | blastp | *Nam-Co_Pisuviricota_212* | 91.69 |
| LS_coassembly_k141_260675_1 | *Pisuviricota* | *c.0012* | *Sobelivirales* | *Solemoviridae* |  | *Nam-Co_genus_162* | blastp | *Nam-Co_Pisuviricota_213* | 94.01 |
| LS_coassembly_k141_283877_1 | *Pisuviricota* | *c.0012* | *Sobelivirales* | *Solemoviridae* |  | *Nam-Co_genus_161* | blastp | *Nam-Co_Pisuviricota_214* | 94.98 |
| LS_coassembly_k141_1223511_1 | *Pisuviricota* | *c.0012* | *Sobelivirales* | *Solemoviridae* |  | *Nam-Co_genus_163* | blastp | *Nam-Co_Pisuviricota_215* | 94.12 |
| LS_coassembly_k141_1130307_2 | *Pisuviricota* | *c.0012* | *Sobelivirales* | *Solemoviridae* | *Nam-Co_family_28* | *Nam-Co_genus_165* | blastp | *Nam-Co_Pisuviricota_216* | 91.9 |
| LC_80m_0.2um_k141_542511_1 | *Kitrinoviricota* |  |  |  | *Nam-Co_family_48* | *Nam-Co_genus_257* | structure | *Nam-Co_Kitrinoviricota_001* | 67.35 |
| LC_coassembly_k141_1596549_1 | *Kitrinoviricota* |  |  |  | *Nam-Co_family_47* | *Nam-Co_genus_256* | structure | *Nam-Co_Kitrinoviricota_002* | 82.65 |
| LC_coassembly_k141_1264191_1 | *Kitrinoviricota* | *Alsuviricetes* | *Hepelivirales* | *Benyviridae* |  |  | blastp | *Nam-Co_Kitrinoviricota_003* | 85.73 |
| LC_coassembly_k141_1830496_1 | *Kitrinoviricota* | *Alsuviricetes* | *Hepelivirales* | *Benyviridae* |  |  | blastp | *Nam-Co_Kitrinoviricota_004* | 88.94 |
| LC_0.5m_3um_k141_269472_1 | *Kitrinoviricota* | *Alsuviricetes* | *Hepelivirales* | *Benyviridae* |  |  | blastp | *Nam-Co_Kitrinoviricota_005* | 86.43 |
| LS_coassembly_k141_614602_1 | *Kitrinoviricota* | *Alsuviricetes* | *Hepelivirales* | *Benyviridae* |  |  | blastp | *Nam-Co_Kitrinoviricota_006* | 88.87 |
| LC_0.5m_0.2um_k141_202252_1 | *Kitrinoviricota* | *Alsuviricetes* | *Hepelivirales* | *Benyviridae* | *Nam-Co_family_50* | *Nam-Co_genus_258* | blastp | *Nam-Co_Kitrinoviricota_007* | 83.25 |
| LC_80m_3um_k141_47471_1 | *Kitrinoviricota* | *Alsuviricetes* | *Hepelivirales* | *Benyviridae* |  |  | blastp | *Nam-Co_Kitrinoviricota_008* | 90.38 |
| LC_20m_10um_k141_1321_1 | *Kitrinoviricota* | *Alsuviricetes* | *o.0055* | *Endornaviridae* |  | *Nam-Co_genus_265* | blastp | *Nam-Co_Kitrinoviricota_009* | 93.34 |
| LC_80m_10um_k141_20695_2 | *Kitrinoviricota* | *Alsuviricetes* | *o.0055* | *Endornaviridae* |  | *Nam-Co_genus_268* | blastp | *Nam-Co_Kitrinoviricota_010* | 93.16 |
| LC_coassembly_k141_2476813_1 | *Kitrinoviricota* | *Alsuviricetes* | *o.0055* | *Endornaviridae* |  | *Nam-Co_genus_266* | blastp | *Nam-Co_Kitrinoviricota_011* | 91.49 |
| LS_coassembly_k141_607677_1 | *Kitrinoviricota* | *Alsuviricetes* | *o.0055* | *Endornaviridae* |  | *Nam-Co_genus_264* | blastp | *Nam-Co_Kitrinoviricota_012* | 91.88 |
| LC_coassembly_k141_1576953_1 | *Kitrinoviricota* | *Alsuviricetes* | *o.0055* | *Endornaviridae* |  |  | blastp | *Nam-Co_Kitrinoviricota_013* | 93.1 |
| LS_coassembly_k141_210683_1 | *Kitrinoviricota* | *Alsuviricetes* | *o.0055* | *Endornaviridae* |  | *Nam-Co_genus_263* | blastp | *Nam-Co_Kitrinoviricota_014* | 90.71 |
| LC_coassembly_k141_515640_1 | *Kitrinoviricota* | *Alsuviricetes* | *o.0055* | *Endornaviridae* |  | *Nam-Co_genus_262* | blastp | *Nam-Co_Kitrinoviricota_015* | 91.7 |
| LC_0.5m_10um_k141_109854_2 | *Kitrinoviricota* | *Alsuviricetes* | *o.0055* | *Endornaviridae* |  | *Nam-Co_genus_265* | blastp | *Nam-Co_Kitrinoviricota_016* | 93.4 |
| LC_coassembly_k141_2772377_2 | *Kitrinoviricota* | *Alsuviricetes* | *o.0055* | *Endornaviridae* |  | *Nam-Co_genus_267* | blastp | *Nam-Co_Kitrinoviricota_017* | 92.19 |
| LS_coassembly_k141_332531_2 | *Kitrinoviricota* | *c.0021.base-Toluca* | *o.0032* | *f.0125* |  | *Nam-Co_genus_270* | blastp | *Nam-Co_Kitrinoviricota_018* | 91.85 |
| LC_coassembly_k141_2551561_2 | *Kitrinoviricota* | *c.0021.base-Toluca* | *o.0032* | *f.0125* |  | *Nam-Co_genus_269* | blastp | *Nam-Co_Kitrinoviricota_019* | 93.15 |
| LC_coassembly_k141_1385123_1 | *Kitrinoviricota* | *c.0021.base-Toluca* | *o.0032* | *f.0125* |  | *Nam-Co_genus_271* | blastp | *Nam-Co_Kitrinoviricota_020* | 90.14 |
| LS_coassembly_k141_981726_2 | *Kitrinoviricota* | *c.0024* | *o.0035* | *f.0144* |  | *Nam-Co_genus_274* | blastp | *Nam-Co_Kitrinoviricota_021* | 91.13 |
| LS_coassembly_k141_808201_2 | *Kitrinoviricota* | *c.0024* | *o.0035* | *f.0144* |  | *Nam-Co_genus_273* | blastp | *Nam-Co_Kitrinoviricota_022* | 93.65 |
| LS_coassembly_k141_660091_1 | *Kitrinoviricota* | *c.0024* | *o.0035* | *f.0144* |  |  | blastp | *Nam-Co_Kitrinoviricota_023* | 94.18 |
| LS_0.5m_1.2um_k141_304513_1 | *Kitrinoviricota* | *c.0024* | *o.0035* | *f.0144* |  | *Nam-Co_genus_272* | blastp | *Nam-Co_Kitrinoviricota_024* | 91.65 |
| LS_coassembly_k141_1029418_2 | *Kitrinoviricota* | *c.0024* | *o.0035* | *f.0144* |  |  | blastp | *Nam-Co_Kitrinoviricota_025* | 94.54 |
| LS_coassembly_k141_1225259_2 | *Kitrinoviricota* | *c.0025* | *o.0036* | *f.0145* | *Nam-Co_family_52* | *Nam-Co_genus_278* | blastp | *Nam-Co_Kitrinoviricota_026* | 83.73 |
| LC_coassembly_k141_945938_1 | *Kitrinoviricota* | *c.0025* | *o.0036* | *f.0145* | *Nam-Co_family_51* | *Nam-Co_genus_275* | blastp | *Nam-Co_Kitrinoviricota_027* | 84.13 |
| LC_coassembly_k141_779785_2 | *Kitrinoviricota* | *c.0025* | *o.0036* | *f.0145* |  | *Nam-Co_genus_277* | blastp | *Nam-Co_Kitrinoviricota_028* | 89.67 |
| LC_coassembly_k141_2737082_1 | *Kitrinoviricota* | *c.0025* | *o.0036* | *f.0145* |  | *Nam-Co_genus_276* | blastp | *Nam-Co_Kitrinoviricota_029* | 90.85 |
| LC_coassembly_k141_1148680_3 | *Kitrinoviricota* | *c.0028* | *o.0039* | *f.0150* |  | *Nam-Co_genus_279* | blastp | *Nam-Co_Kitrinoviricota_030* | 90.9 |
| LC_coassembly_k141_2619831_4 | *Kitrinoviricota* | *c.0028* | *o.0039* | *f.0150* |  | *Nam-Co_genus_280* | blastp | *Nam-Co_Kitrinoviricota_031* | 89.13 |
| LC_coassembly_k141_2173116_1 | *Kitrinoviricota* | *c.0031* | *o.0042* | *f.0153* |  | *Nam-Co_genus_281* | blastp | *Nam-Co_Kitrinoviricota_032* | 90.52 |
| LC_50m_1.2um_k141_61970_1 | *Kitrinoviricota* | *c.0031* | *o.0042* | *f.0153* |  | *Nam-Co_genus_282* | blastp | *Nam-Co_Kitrinoviricota_033* | 85.67 |
| LC_coassembly_k141_1913062_1 | *Kitrinoviricota* | *c.0031* | *o.0042* | *f.0153* |  | *Nam-Co_genus_284* | blastp | *Nam-Co_Kitrinoviricota_034* | 92.51 |
| LC_80m_0.2um_k141_512934_1 | *Kitrinoviricota* | *c.0031* | *o.0042* | *f.0153* |  | *Nam-Co_genus_286* | blastp | *Nam-Co_Kitrinoviricota_035* | 89.33 |
| LC_80m_1.2um_k141_431363_1 | *Kitrinoviricota* | *c.0031* | *o.0042* | *f.0153* |  | *Nam-Co_genus_286* | blastp | *Nam-Co_Kitrinoviricota_036* | 84.25 |
| LC_coassembly_k141_776867_2 | *Kitrinoviricota* | *c.0031* | *o.0042* | *f.0153* | *Nam-Co_family_53* | *Nam-Co_genus_285* | blastp | *Nam-Co_Kitrinoviricota_037* |  |
| LC_coassembly_k141_204657_2 | *Kitrinoviricota* | *c.0031* | *o.0042* | *f.0153* |  | *Nam-Co_genus_283* | blastp | *Nam-Co_Kitrinoviricota_038* | 93.18 |
| LS_coassembly_k141_1166858_1 | *Kitrinoviricota* | *c.0031* | *o.0042* | *f.0156* |  | *Nam-Co_genus_287* | blastp | *Nam-Co_Kitrinoviricota_039* | 91.06 |
| LS_coassembly_k141_1250841_2 | *Kitrinoviricota* | *c.0031* | *o.0042* | *f.0156* |  |  | blastp | *Nam-Co_Kitrinoviricota_040* | 89.92 |
| LC_coassembly_k141_1476401_5 | *Kitrinoviricota* | *c.0032* | *o.0043* | *f.0167* |  | *Nam-Co_genus_288* | blastp | *Nam-Co_Kitrinoviricota_041* | 89.14 |
| LS_coassembly_k141_738808_4 | *Kitrinoviricota* | *c.0032* | *o.0043* | *f.0167* | *Nam-Co_family_55* | *Nam-Co_genus_289* | blastp | *Nam-Co_Kitrinoviricota_042* | 85.87 |
| LC_coassembly_k141_2331475_4 | *Kitrinoviricota* | *c.0032* | *o.0043* | *f.0167* |  |  | blastp | *Nam-Co_Kitrinoviricota_043* | 90.47 |
| LC_20m_3um_k141_448366_1 | *Kitrinoviricota* | *c.0032* | *o.0043* | *f.0167* |  |  | blastp | *Nam-Co_Kitrinoviricota_044* | 90.75 |
| LC_coassembly_k141_1960344_7 | *Kitrinoviricota* | *c.0032* | *o.0043* | *f.0167* | *Nam-Co_family_54* | *Nam-Co_genus_290* | blastp | *Nam-Co_Kitrinoviricota_045* | 88.74 |
| LC_coassembly_k141_899203_3 | *Kitrinoviricota* | *c.0032* | *o.0043* | *f.0167* | *Nam-Co_family_54* | *Nam-Co_genus_290* | blastp | *Nam-Co_Kitrinoviricota_046* | 86.45 |
| LS_coassembly_k141_223527_1 | *Kitrinoviricota* | *c.0032* | *o.0043* | *f.0169* | *Nam-Co_family_56* | *Nam-Co_genus_291* | blastp | *Nam-Co_Kitrinoviricota_047* | 88.6 |
| LC_coassembly_k141_537252_5 | *Kitrinoviricota* | *c.0034* | *o.0045* | *f.0176* | *Nam-Co_family_57* | *Nam-Co_genus_296* | blastp | *Nam-Co_Kitrinoviricota_048* | 83.89 |
| LC_coassembly_k141_1638127_4 | *Kitrinoviricota* | *c.0034* | *o.0045* | *f.0176* |  | *Nam-Co_genus_295* | blastp | *Nam-Co_Kitrinoviricota_049* | 88.63 |
| LC_50m_0.2um_k141_320664_3 | *Kitrinoviricota* | *c.0034* | *o.0045* | *f.0176* |  | *Nam-Co_genus_295* | blastp | *Nam-Co_Kitrinoviricota_050* | 89.13 |
| LC_80m_3um_k141_96140_1 | *Kitrinoviricota* | *c.0034* | *o.0045* | *f.0176* |  | *Nam-Co_genus_295* | blastp | *Nam-Co_Kitrinoviricota_051* | 88.93 |
| LC_0.5m_1.2um_k141_135095_2 | *Kitrinoviricota* | *c.0034* | *o.0045* | *f.0176* |  | *Nam-Co_genus_295* | blastp | *Nam-Co_Kitrinoviricota_052* | 87.25 |
| LC_80m_10um_k141_122255_1 | *Kitrinoviricota* | *c.0034* | *o.0045* | *f.0176* |  | *Nam-Co_genus_294* | blastp | *Nam-Co_Kitrinoviricota_053* | 89.98 |
| LC_coassembly_k141_843059_2 | *Kitrinoviricota* | *c.0034* | *o.0045* | *f.0176* |  | *Nam-Co_genus_293* | blastp | *Nam-Co_Kitrinoviricota_054* | 93.36 |
| LC_50m_0.2um_k141_58378_2 | *Kitrinoviricota* | *c.0034* | *o.0045* | *f.0176* |  | *Nam-Co_genus_295* | blastp | *Nam-Co_Kitrinoviricota_055* | 88.76 |
| LC_0.5m_3um_k141_382100_1 | *Kitrinoviricota* | *c.0034* | *o.0045* | *f.0176* |  | *Nam-Co_genus_295* | blastp | *Nam-Co_Kitrinoviricota_056* | 93.06 |
| LC_50m_1.2um_k141_255003_1 | *Kitrinoviricota* | *c.0034* | *o.0045* | *f.0176* |  | *Nam-Co_genus_295* | blastp | *Nam-Co_Kitrinoviricota_057* | 92.44 |
| LC_80m_10um_k141_219831_1 | *Kitrinoviricota* | *c.0034* | *o.0045* | *f.0176* |  | *Nam-Co_genus_295* | blastp | *Nam-Co_Kitrinoviricota_058* | 91.75 |
| LC_coassembly_k141_1747163_4 | *Kitrinoviricota* | *c.0034* | *o.0045* | *f.0176* |  | *Nam-Co_genus_294* | blastp | *Nam-Co_Kitrinoviricota_059* | 90.84 |
| LC_0.5m_10um_k141_14159_3 | *Kitrinoviricota* | *c.0034* | *o.0045* | *f.0176* |  | *Nam-Co_genus_294* | blastp | *Nam-Co_Kitrinoviricota_060* | 90.22 |
| LC_20m_0.2um_k141_394151_1 | *Kitrinoviricota* | *c.0034* | *o.0045* | *f.0176* |  | *Nam-Co_genus_294* | blastp | *Nam-Co_Kitrinoviricota_061* | 90.39 |
| LC_20m_0.2um_k141_268307_4 | *Kitrinoviricota* | *c.0034* | *o.0045* | *f.0176* |  | *Nam-Co_genus_293* | blastp | *Nam-Co_Kitrinoviricota_062* | 91.27 |
| LS_coassembly_k141_1254060_1 | *Kitrinoviricota* | *c.0034* | *o.0045* | *f.0176* | *Nam-Co_family_58* | *Nam-Co_genus_292* | blastp | *Nam-Co_Kitrinoviricota_063* | 86.29 |
| LS_0.5m_1.2um_k141_335154_1 | *Kitrinoviricota* | *c.0034* | *o.0045* | *f.0176* |  | *Nam-Co_genus_294* | blastp | *Nam-Co_Kitrinoviricota_064* | 88.31 |
| LC_coassembly_k141_1687493_1 | *Kitrinoviricota* | *c.0034* | *o.0045* | *f.0176* |  | *Nam-Co_genus_294* | blastp | *Nam-Co_Kitrinoviricota_065* | 91.22 |
| LC_coassembly_k141_1418113_3 | *Kitrinoviricota* | *c.0034* | *o.0045* | *f.0176* |  | *Nam-Co_genus_294* | blastp | *Nam-Co_Kitrinoviricota_066* | 91.08 |
| LC_0.5m_10um_k141_200615_2 | *Kitrinoviricota* | *c.0034* | *o.0045* | *f.0176* |  | *Nam-Co_genus_294* | blastp | *Nam-Co_Kitrinoviricota_067* | 90.86 |
| LC_coassembly_k141_1021654_7 | *Kitrinoviricota* | *c.0034* | *o.0045* | *f.0176* |  | *Nam-Co_genus_293* | blastp | *Nam-Co_Kitrinoviricota_068* | 94 |
| LC_20m_1.2um_k141_139004_1 | *Kitrinoviricota* | *c.0034* | *o.0045* | *f.0176* |  | *Nam-Co_genus_293* | blastp | *Nam-Co_Kitrinoviricota_069* | 93.63 |
| LC_coassembly_k141_1111088_3 | *Kitrinoviricota* | *c.0034* | *o.0045* | *f.0176* |  | *Nam-Co_genus_295* | blastp | *Nam-Co_Kitrinoviricota_070* | 93.72 |
| LC_coassembly_k141_1653401_2 | *Kitrinoviricota* | *c.0034* | *o.0045* | *f.0176* |  | *Nam-Co_genus_295* | blastp | *Nam-Co_Kitrinoviricota_071* | 93.11 |
| LC_coassembly_k141_1295637_3 | *Kitrinoviricota* | *c.0034* | *o.0045* | *f.0176* |  | *Nam-Co_genus_295* | blastp | *Nam-Co_Kitrinoviricota_072* | 91.74 |
| LC_coassembly_k141_1296109_4 | *Kitrinoviricota* | *c.0034* | *o.0045* | *f.0176* |  | *Nam-Co_genus_295* | blastp | *Nam-Co_Kitrinoviricota_073* | 92.83 |
| LC_coassembly_k141_1896863_5 | *Kitrinoviricota* | *c.0034* | *o.0045* | *f.0176* |  | *Nam-Co_genus_295* | blastp | *Nam-Co_Kitrinoviricota_074* | 92.31 |
| LC_coassembly_k141_2258078_1 | *Kitrinoviricota* | *c.0034* | *o.0045* | *f.0176* |  | *Nam-Co_genus_295* | blastp | *Nam-Co_Kitrinoviricota_075* | 93.28 |
| LC_coassembly_k141_581270_3 | *Kitrinoviricota* | *c.0034* | *o.0045* | *f.0176* |  | *Nam-Co_genus_295* | blastp | *Nam-Co_Kitrinoviricota_076* | 92.42 |
| LC_coassembly_k141_75697_3 | *Kitrinoviricota* | *c.0034* | *o.0045* | *f.0176* |  | *Nam-Co_genus_295* | blastp | *Nam-Co_Kitrinoviricota_077* | 93.3 |
| LC_coassembly_k141_22087_2 | *Kitrinoviricota* | *c.0034* | *o.0045* | *f.0176* |  | *Nam-Co_genus_295* | blastp | *Nam-Co_Kitrinoviricota_078* | 93.36 |
| LC_coassembly_k141_1259623_2 | *Kitrinoviricota* | *c.0034* | *o.0045* | *f.0176* |  | *Nam-Co_genus_295* | blastp | *Nam-Co_Kitrinoviricota_079* | 92.97 |
| LC_0.5m_0.2um_k141_102965_1 | *Kitrinoviricota* | *c.0034* | *o.0045* | *f.0176* |  | *Nam-Co_genus_295* | blastp | *Nam-Co_Kitrinoviricota_080* | 93.11 |
| LC_0.5m_10um_k141_25814_1 | *Kitrinoviricota* | *c.0034* | *o.0045* | *f.0176* |  | *Nam-Co_genus_295* | blastp | *Nam-Co_Kitrinoviricota_081* | 92.12 |
| LC_0.5m_1.2um_k141_34913_1 | *Kitrinoviricota* | *c.0034* | *o.0045* | *f.0176* |  | *Nam-Co_genus_295* | blastp | *Nam-Co_Kitrinoviricota_082* | 93.07 |
| LC_20m_3um_k141_164113_3 | *Kitrinoviricota* | *c.0034* | *o.0045* | *f.0176* |  | *Nam-Co_genus_295* | blastp | *Nam-Co_Kitrinoviricota_083* | 93.66 |
| LC_50m_1.2um_k141_154296_1 | *Kitrinoviricota* | *c.0034* | *o.0045* | *f.0176* |  | *Nam-Co_genus_295* | blastp | *Nam-Co_Kitrinoviricota_084* | 93.75 |
| LC_50m_3um_k141_403292_3 | *Kitrinoviricota* | *c.0034* | *o.0045* | *f.0176* |  | *Nam-Co_genus_295* | blastp | *Nam-Co_Kitrinoviricota_085* | 93.59 |
| LC_coassembly_k141_1021355_5 | *Kitrinoviricota* | *c.0034* | *o.0045* | *f.0176* |  | *Nam-Co_genus_293* | blastp | *Nam-Co_Kitrinoviricota_086* | 94.34 |
| LC_coassembly_k141_854051_1 | *Kitrinoviricota* | *c.0034* | *o.0045* | *f.0176* |  | *Nam-Co_genus_295* | blastp | *Nam-Co_Kitrinoviricota_087* | 89.87 |
| LC_coassembly_k141_2300750_1 | *Kitrinoviricota* | *c.0034* | *o.0045* | *f.0176* |  | *Nam-Co_genus_295* | blastp | *Nam-Co_Kitrinoviricota_088* | 87.59 |
| LC_coassembly_k141_1350294_5 | *Kitrinoviricota* | *c.0034* | *o.0045* | *f.0176* |  | *Nam-Co_genus_295* | blastp | *Nam-Co_Kitrinoviricota_089* | 88.2 |
| LC_0.5m_1.2um_k141_85285_1 | *Kitrinoviricota* | *c.0034* | *o.0045* | *f.0176* |  | *Nam-Co_genus_295* | blastp | *Nam-Co_Kitrinoviricota_090* | 89.02 |
| LC_20m_0.2um_k141_406732_2 | *Kitrinoviricota* | *c.0034* | *o.0045* | *f.0176* |  | *Nam-Co_genus_295* | blastp | *Nam-Co_Kitrinoviricota_091* | 89.95 |
| LC_80m_10um_k141_242013_1 | *Kitrinoviricota* | *c.0034* | *o.0045* | *f.0176* |  | *Nam-Co_genus_295* | blastp | *Nam-Co_Kitrinoviricota_092* | 89.56 |
| LC_0.5m_3um_k141_163067_2 | *Kitrinoviricota* | *c.0034* | *o.0045* | *f.0176* |  | *Nam-Co_genus_295* | blastp | *Nam-Co_Kitrinoviricota_093* | 86.21 |
| LS_0.5m_3um_k141_311742_1 | *Kitrinoviricota* | *c.0034* | *o.0045* | *f.0176* |  | *Nam-Co_genus_294* | blastp | *Nam-Co_Kitrinoviricota_094* | 91.5 |
| LS_coassembly_k141_512850_2 | *Kitrinoviricota* | *c.0034* | *o.0045* | *f.0181* | *Nam-Co_family_59* | *Nam-Co_genus_298* | blastp | *Nam-Co_Kitrinoviricota_095* | 82.13 |
| LS_coassembly_k141_1072213_1 | *Kitrinoviricota* | *c.0034* | *o.0045* | *f.0181* |  | *Nam-Co_genus_297* | blastp | *Nam-Co_Kitrinoviricota_096* | 91.73 |
| LS_coassembly_k141_399972_1 | *Kitrinoviricota* | *Alsuviricetes* | *Hepelivirales* | *f.0221* |  | *Nam-Co_genus_259* | blastp | *Nam-Co_Kitrinoviricota_097* | 88.89 |
| LC_coassembly_k141_488786_1 | *Kitrinoviricota* | *Alsuviricetes* | *Hepelivirales* | *f.0221* |  |  | blastp | *Nam-Co_Kitrinoviricota_098* | 88.69 |
| LC_coassembly_k141_524640_2 | *Kitrinoviricota* | *Alsuviricetes* | *Hepelivirales* | *f.0234* | *Nam-Co_family_49* | *Nam-Co_genus_260* | blastp | *Nam-Co_Kitrinoviricota_099* |  |
| LS_coassembly_k141_507967_1 | *Kitrinoviricota* | *Magsaviricetes* | *Nodamuvirales* | *Nodaviridae* |  | *Nam-Co_genus_300* | blastp | *Nam-Co_Kitrinoviricota_100* | 93.6 |
| LS_coassembly_k141_1074570_1 | *Kitrinoviricota* | *Magsaviricetes* | *Nodamuvirales* | *Nodaviridae* |  |  | blastp | *Nam-Co_Kitrinoviricota_101* | 92.38 |
| LC_50m_3um_k141_164483_1 | *Kitrinoviricota* | *Magsaviricetes* | *Nodamuvirales* | *Nodaviridae* |  | *Nam-Co_genus_301* | blastp | *Nam-Co_Kitrinoviricota_102* | 89.32 |
| LS_coassembly_k141_952038_2 | *Kitrinoviricota* | *Magsaviricetes* | *Nodamuvirales* | *Nodaviridae* |  | *Nam-Co_genus_303* | blastp | *Nam-Co_Kitrinoviricota_103* | 68.71 |
| LC_coassembly_k141_2580714_1 | *Kitrinoviricota* | *Magsaviricetes* | *Nodamuvirales* | *Nodaviridae* |  | *Nam-Co_genus_301* | blastp | *Nam-Co_Kitrinoviricota_104* | 93.77 |
| LC_coassembly_k141_500165_1 | *Kitrinoviricota* | *Magsaviricetes* | *Nodamuvirales* | *Nodaviridae* |  | *Nam-Co_genus_302* | blastp | *Nam-Co_Kitrinoviricota_105* | 93.37 |
| LC_coassembly_k141_1841077_1 | *Kitrinoviricota* | *Magsaviricetes* | *Nodamuvirales* | *Nodaviridae* |  | *Nam-Co_genus_299* | blastp | *Nam-Co_Kitrinoviricota_106* | 94.6 |
| LC_20m_0.2um_k141_317770_1 | *Kitrinoviricota* | *Magsaviricetes* | *Nodamuvirales* | *Nodaviridae* |  |  | blastp | *Nam-Co_Kitrinoviricota_107* | 94.6 |
| LS_coassembly_k141_1311787_3 | *Kitrinoviricota* | *Tolucaviricetes* | *Tolivirales* | *Tombusviridae* | *Nam-Co_family_60* | *Nam-Co_genus_309* | blastp | *Nam-Co_Kitrinoviricota_108* | 90.94 |
| LS_coassembly_k141_708466_1 | *Kitrinoviricota* | *Tolucaviricetes* | *Tolivirales* | *Tombusviridae* |  | *Nam-Co_genus_306* | blastp | *Nam-Co_Kitrinoviricota_109* | 94.87 |
| LC_coassembly_k141_1773142_1 | *Kitrinoviricota* | *Tolucaviricetes* | *Tolivirales* | *Tombusviridae* |  |  | blastp | *Nam-Co_Kitrinoviricota_110* | 94.71 |
| LC_coassembly_k141_1075936_1 | *Kitrinoviricota* | *Tolucaviricetes* | *Tolivirales* | *Tombusviridae* |  |  | blastp | *Nam-Co_Kitrinoviricota_111* | 94.7 |
| LC_coassembly_k141_2449152_1 | *Kitrinoviricota* | *Tolucaviricetes* | *Tolivirales* | *Tombusviridae* |  |  | blastp | *Nam-Co_Kitrinoviricota_112* | 94.62 |
| LC_coassembly_k141_952072_1 | *Kitrinoviricota* | *Tolucaviricetes* | *Tolivirales* | *Tombusviridae* |  |  | blastp | *Nam-Co_Kitrinoviricota_113* | 94.98 |
| LS_coassembly_k141_1028965_2 | *Kitrinoviricota* | *Tolucaviricetes* | *Tolivirales* | *Tombusviridae* |  | *Nam-Co_genus_304* | blastp | *Nam-Co_Kitrinoviricota_114* | 92.3 |
| LS_coassembly_k141_220020_1 | *Kitrinoviricota* | *Tolucaviricetes* | *Tolivirales* | *Tombusviridae* |  |  | blastp | *Nam-Co_Kitrinoviricota_115* | 94.82 |
| LC_coassembly_k141_757367_1 | *Kitrinoviricota* | *Tolucaviricetes* | *Tolivirales* | *Tombusviridae* |  | *Nam-Co_genus_308* | blastp | *Nam-Co_Kitrinoviricota_116* | 93.8 |
| LC_coassembly_k141_76989_2 | *Kitrinoviricota* | *Tolucaviricetes* | *Tolivirales* | *Tombusviridae* |  |  | blastp | *Nam-Co_Kitrinoviricota_117* | 94.26 |
| LS_coassembly_k141_241270_2 | *Kitrinoviricota* | *Tolucaviricetes* | *Tolivirales* | *Tombusviridae* |  | *Nam-Co_genus_305* | blastp | *Nam-Co_Kitrinoviricota_118* | 94.79 |
| LS_coassembly_k141_1069082_1 | *Kitrinoviricota* | *Tolucaviricetes* | *Tolivirales* | *Tombusviridae* |  |  | blastp | *Nam-Co_Kitrinoviricota_119* | 94.85 |
| LS_0.5m_1.2um_k141_244393_1 | *Kitrinoviricota* | *Tolucaviricetes* | *Tolivirales* | *Tombusviridae* |  |  | blastp | *Nam-Co_Kitrinoviricota_120* | 93.41 |
| LS_coassembly_k141_57787_4 | *Kitrinoviricota* | *Tolucaviricetes* | *Tolivirales* | *Tombusviridae* |  | *Nam-Co_genus_307* | blastp | *Nam-Co_Kitrinoviricota_121* | 95.18 |
| LS_coassembly_k141_565475_1 | *Kitrinoviricota* | *Tolucaviricetes* | *Tolivirales* | *Tombusviridae* |  |  | blastp | *Nam-Co_Kitrinoviricota_122* | 95.35 |
| LS_coassembly_k141_783029_1 | *Kitrinoviricota* | *Tolucaviricetes* | *Tolivirales* | *Tombusviridae* |  |  | blastp | *Nam-Co_Kitrinoviricota_123* | 95.04 |
| LS_coassembly_k141_765865_1 | *Kitrinoviricota* | *Tolucaviricetes* | *Tolivirales* | *Tombusviridae* |  |  | blastp | *Nam-Co_Kitrinoviricota_124* | 93.09 |
| LC_coassembly_k141_1486019_1 | *Kitrinoviricota* | *Alsuviricetes* | *Martellivirales* | *Virgaviridae* |  | *Nam-Co_genus_261* | blastp | *Nam-Co_Kitrinoviricota_125* | 92.54 |
| LC_20m_3um_k141_43900_1 | *Kitrinoviricota* | *Alsuviricetes* | *Martellivirales* | *Virgaviridae* |  | *Nam-Co_genus_261* | blastp | *Nam-Co_Kitrinoviricota_126* | 92.51 |
| LC_coassembly_k141_2104764_1 | *Kitrinoviricota* | *Alsuviricetes* | *Martellivirales* | *Virgaviridae* |  |  | blastp | *Nam-Co_Kitrinoviricota_127* | 91.12 |
| LC_coassembly_k141_196466_2 | *Kitrinoviricota* | *Alsuviricetes* | *Martellivirales* | *Virgaviridae* |  |  | blastp | *Nam-Co_Kitrinoviricota_128* | 92.4 |
| LC_20m_3um_k141_304944_1 | *Duplornaviricota* |  |  |  | *Nam-Co_family_61* | *Nam-Co_genus_310* | structure | *Nam-Co_Duplornaviricota_001* | 61.36 |
| LC_coassembly_k141_1528827_1 | *Duplornaviricota* | *Chrymotiviricetes* | *Ghabrivirales* | *f.0282* |  | *Nam-Co_genus_311* | blastp | *Nam-Co_Duplornaviricota_002* | 90.71 |
| LC_coassembly_k141_2537970_1 | *Duplornaviricota* | *Chrymotiviricetes* | *Ghabrivirales* | *f.0282* |  | *Nam-Co_genus_312* | blastp | *Nam-Co_Duplornaviricota_003* | 87.38 |
| LC_50m_3um_k141_208177_2 | *Duplornaviricota* | *Chrymotiviricetes* | *Ghabrivirales* | *f.0282* |  | *Nam-Co_genus_313* | blastp | *Nam-Co_Duplornaviricota_004* | 90.91 |
| LC_coassembly_k141_276839_1 | *Duplornaviricota* | *Chrymotiviricetes* | *Ghabrivirales* | *f.0282* |  | *Nam-Co_genus_313* | blastp | *Nam-Co_Duplornaviricota_005* | 90.98 |
| LC_80m_3um_k141_257744_1 | *Duplornaviricota* | *Chrymotiviricetes* | *Ghabrivirales* | *f.0282* |  | *Nam-Co_genus_313* | blastp | *Nam-Co_Duplornaviricota_006* | 90.92 |
| LC_coassembly_k141_908918_1 | *Duplornaviricota* | *Chrymotiviricetes* | *Ghabrivirales* | *f.0283* |  | *Nam-Co_genus_318* | blastp | *Nam-Co_Duplornaviricota_007* | 89.02 |
| LC_coassembly_k141_40647_2 | *Duplornaviricota* | *Chrymotiviricetes* | *Ghabrivirales* | *f.0283* | *Nam-Co_family_65* | *Nam-Co_genus_315* | blastp | *Nam-Co_Duplornaviricota_008* | 89.6 |
| LC_coassembly_k141_1642998_2 | *Duplornaviricota* | *Chrymotiviricetes* | *Ghabrivirales* | *f.0283* |  |  | blastp | *Nam-Co_Duplornaviricota_009* | 85.43 |
| LC_coassembly_k141_1445686_1 | *Duplornaviricota* | *Chrymotiviricetes* | *Ghabrivirales* | *f.0283* |  | *Nam-Co_genus_318* | blastp | *Nam-Co_Duplornaviricota_010* | 88.76 |
| LC_20m_10um_k141_36758_1 | *Duplornaviricota* | *Chrymotiviricetes* | *Ghabrivirales* | *f.0283* |  | *Nam-Co_genus_316* | blastp | *Nam-Co_Duplornaviricota_011* | 89.88 |
| LC_coassembly_k141_1614393_1 | *Duplornaviricota* | *Chrymotiviricetes* | *Ghabrivirales* | *f.0283* |  | *Nam-Co_genus_317* | blastp | *Nam-Co_Duplornaviricota_012* | 89.82 |
| LC_20m_10um_k141_12836_1 | *Duplornaviricota* | *Chrymotiviricetes* | *Ghabrivirales* | *f.0283* |  | *Nam-Co_genus_318* | blastp | *Nam-Co_Duplornaviricota_013* | 89.13 |
| LC_20m_10um_k141_27694_1 | *Duplornaviricota* | *Chrymotiviricetes* | *Ghabrivirales* | *f.0283* |  | *Nam-Co_genus_318* | blastp | *Nam-Co_Duplornaviricota_014* | 91.03 |
| LC_20m_3um_k141_171088_1 | *Duplornaviricota* | *Chrymotiviricetes* | *Ghabrivirales* | *f.0283* |  | *Nam-Co_genus_319* | blastp | *Nam-Co_Duplornaviricota_015* | 89.12 |
| LC_coassembly_k141_1136066_1 | *Duplornaviricota* | *Chrymotiviricetes* | *Ghabrivirales* | *f.0283* |  | *Nam-Co_genus_318* | blastp | *Nam-Co_Duplornaviricota_016* | 89.97 |
| LC_20m_10um_k141_2297_1 | *Duplornaviricota* | *Chrymotiviricetes* | *Ghabrivirales* | *f.0283* |  | *Nam-Co_genus_318* | blastp | *Nam-Co_Duplornaviricota_017* | 89.52 |
| LC_20m_10um_k141_57615_1 | *Duplornaviricota* | *Chrymotiviricetes* | *Ghabrivirales* | *f.0283* |  | *Nam-Co_genus_318* | blastp | *Nam-Co_Duplornaviricota_018* | 88.1 |
| LC_20m_10um_k141_6666_1 | *Duplornaviricota* | *Chrymotiviricetes* | *Ghabrivirales* | *f.0283* |  | *Nam-Co_genus_316* | blastp | *Nam-Co_Duplornaviricota_019* | 88.49 |
| LC_20m_10um_k141_55953_1 | *Duplornaviricota* | *Chrymotiviricetes* | *Ghabrivirales* | *f.0283* |  | *Nam-Co_genus_318* | blastp | *Nam-Co_Duplornaviricota_020* | 89.16 |
| LC_20m_10um_k141_42987_1 | *Duplornaviricota* | *Chrymotiviricetes* | *Ghabrivirales* | *f.0283* |  | *Nam-Co_genus_319* | blastp | *Nam-Co_Duplornaviricota_021* | 88.77 |
| LC_coassembly_k141_1810732_1 | *Duplornaviricota* | *Chrymotiviricetes* | *Ghabrivirales* | *f.0283* |  | *Nam-Co_genus_319* | blastp | *Nam-Co_Duplornaviricota_022* | 87.49 |
| LC_coassembly_k141_483085_1 | *Duplornaviricota* | *Chrymotiviricetes* | *Ghabrivirales* | *f.0283* |  | *Nam-Co_genus_319* | blastp | *Nam-Co_Duplornaviricota_023* | 91.15 |
| LC_coassembly_k141_1763940_1 | *Duplornaviricota* | *Chrymotiviricetes* | *Ghabrivirales* | *f.0283* |  | *Nam-Co_genus_319* | blastp | *Nam-Co_Duplornaviricota_024* | 89.7 |
| LC_coassembly_k141_2314916_1 | *Duplornaviricota* | *Chrymotiviricetes* | *Ghabrivirales* | *f.0283* |  | *Nam-Co_genus_314* | blastp | *Nam-Co_Duplornaviricota_025* | 87.71 |
| LC_20m_10um_k141_70830_1 | *Duplornaviricota* | *Chrymotiviricetes* | *Ghabrivirales* | *f.0283* |  | *Nam-Co_genus_316* | blastp | *Nam-Co_Duplornaviricota_026* | 88.17 |
| LC_coassembly_k141_2477916_1 | *Duplornaviricota* | *Chrymotiviricetes* | *Ghabrivirales* | *f.0283* |  | *Nam-Co_genus_319* | blastp | *Nam-Co_Duplornaviricota_027* | 87.31 |
| LC_coassembly_k141_1135645_1 | *Duplornaviricota* | *Chrymotiviricetes* | *Ghabrivirales* | *f.0283* |  | *Nam-Co_genus_319* | blastp | *Nam-Co_Duplornaviricota_028* | 88.02 |
| LC_20m_10um_k141_12100_1 | *Duplornaviricota* | *Chrymotiviricetes* | *Ghabrivirales* | *f.0283* |  | *Nam-Co_genus_319* | blastp | *Nam-Co_Duplornaviricota_029* | 87.88 |
| LC_coassembly_k141_1779233_1 | *Duplornaviricota* | *Chrymotiviricetes* | *Ghabrivirales* | *f.0283* |  | *Nam-Co_genus_319* | blastp | *Nam-Co_Duplornaviricota_030* | 90.81 |
| LC_coassembly_k141_1840495_1 | *Duplornaviricota* | *Chrymotiviricetes* | *Ghabrivirales* | *f.0283* |  | *Nam-Co_genus_319* | blastp | *Nam-Co_Duplornaviricota_031* | 92.03 |
| LC_20m_10um_k141_45308_1 | *Duplornaviricota* | *Chrymotiviricetes* | *Ghabrivirales* | *f.0283* |  | *Nam-Co_genus_319* | blastp | *Nam-Co_Duplornaviricota_032* | 91.92 |
| LC_20m_3um_k141_328198_1 | *Duplornaviricota* | *Chrymotiviricetes* | *Ghabrivirales* | *f.0283* |  | *Nam-Co_genus_319* | blastp | *Nam-Co_Duplornaviricota_033* | 89.98 |
| LC_coassembly_k141_1485030_1 | *Duplornaviricota* | *Chrymotiviricetes* | *Ghabrivirales* | *f.0283* | *Nam-Co_family_65* | *Nam-Co_genus_315* | blastp | *Nam-Co_Duplornaviricota_034* | 89.73 |
| LC_coassembly_k141_2800168_1 | *Duplornaviricota* | *Chrymotiviricetes* | *Ghabrivirales* | *f.0283* |  | *Nam-Co_genus_319* | blastp | *Nam-Co_Duplornaviricota_035* | 91.61 |
| LC_20m_0.2um_k141_16816_1 | *Duplornaviricota* | *Chrymotiviricetes* | *Ghabrivirales* | *f.0283* |  | *Nam-Co_genus_319* | blastp | *Nam-Co_Duplornaviricota_036* | 90.85 |
| LC_20m_10um_k141_63442_1 | *Duplornaviricota* | *Chrymotiviricetes* | *Ghabrivirales* | *f.0283* |  | *Nam-Co_genus_319* | blastp | *Nam-Co_Duplornaviricota_037* | 90.77 |
| LC_20m_10um_k141_22158_1 | *Duplornaviricota* | *Chrymotiviricetes* | *Ghabrivirales* | *f.0283* |  | *Nam-Co_genus_319* | blastp | *Nam-Co_Duplornaviricota_038* | 91.08 |
| LC_coassembly_k141_2172855_1 | *Duplornaviricota* | *Chrymotiviricetes* | *Ghabrivirales* | *f.0283* |  | *Nam-Co_genus_314* | blastp | *Nam-Co_Duplornaviricota_039* | 89.65 |
| LC_20m_10um_k141_47367_1 | *Duplornaviricota* | *Chrymotiviricetes* | *Ghabrivirales* | *f.0283* |  | *Nam-Co_genus_319* | blastp | *Nam-Co_Duplornaviricota_040* | 84.74 |
| LC_20m_10um_k141_46604_1 | *Duplornaviricota* | *Chrymotiviricetes* | *Ghabrivirales* | *f.0283* |  | *Nam-Co_genus_319* | blastp | *Nam-Co_Duplornaviricota_041* | 86.26 |
| LC_coassembly_k141_721053_2 | *Duplornaviricota* | *Chrymotiviricetes* | *Ghabrivirales* | *f.0284* | *Nam-Co_family_62* | *Nam-Co_genus_320* | blastp | *Nam-Co_Duplornaviricota_042* | 84.15 |
| LC_coassembly_k141_2175510_1 | *Duplornaviricota* | *Chrymotiviricetes* | *Ghabrivirales* | *f.0285* | *Nam-Co_family_72* | *Nam-Co_genus_324* | blastp | *Nam-Co_Duplornaviricota_043* | 91.34 |
| LC_coassembly_k141_1868495_1 | *Duplornaviricota* | *Chrymotiviricetes* | *Ghabrivirales* | *f.0285* |  | *Nam-Co_genus_325* | blastp | *Nam-Co_Duplornaviricota_044* | 82.89 |
| LC_coassembly_k141_1666324_1 | *Duplornaviricota* | *Chrymotiviricetes* | *Ghabrivirales* | *f.0285* |  | *Nam-Co_genus_321* | blastp | *Nam-Co_Duplornaviricota_045* | 90.85 |
| LC_coassembly_k141_972273_1 | *Duplornaviricota* | *Chrymotiviricetes* | *Ghabrivirales* | *f.0285* | *Nam-Co_family_69* | *Nam-Co_genus_322* | blastp | *Nam-Co_Duplornaviricota_046* | 84.47 |
| LC_coassembly_k141_801488_3 | *Duplornaviricota* | *Chrymotiviricetes* | *Ghabrivirales* | *f.0285* | *Nam-Co_family_63* | *Nam-Co_genus_323* | blastp | *Nam-Co_Duplornaviricota_047* | 85.78 |
| LS_coassembly_k141_419215_1 | *Duplornaviricota* | *Chrymotiviricetes* | *Ghabrivirales* | *f.0285* | *Nam-Co_family_72* | *Nam-Co_genus_324* | blastp | *Nam-Co_Duplornaviricota_048* | 90.89 |
| LC_0.5m_3um_k141_25203_1 | *Duplornaviricota* | *Chrymotiviricetes* | *Ghabrivirales* | *f.0286* | *Nam-Co_family_66* | *Nam-Co_genus_326* | blastp | *Nam-Co_Duplornaviricota_049* | 85.81 |
| LC_coassembly_k141_2048707_1 | *Duplornaviricota* | *Chrymotiviricetes* | *Ghabrivirales* | *f.0286* | *Nam-Co_family_66* | *Nam-Co_genus_326* | blastp | *Nam-Co_Duplornaviricota_050* | 85.3 |
| LC_0.5m_3um_k141_396598_1 | *Duplornaviricota* | *Chrymotiviricetes* | *Ghabrivirales* | *f.0286* | *Nam-Co_family_66* | *Nam-Co_genus_329* | blastp | *Nam-Co_Duplornaviricota_051* | 85.45 |
| LC_20m_3um_k141_132526_2 | *Duplornaviricota* | *Chrymotiviricetes* | *Ghabrivirales* | *f.0286* | *Nam-Co_family_66* | *Nam-Co_genus_329* | blastp | *Nam-Co_Duplornaviricota_052* | 85.17 |
| LC_20m_3um_k141_11485_1 | *Duplornaviricota* | *Chrymotiviricetes* | *Ghabrivirales* | *f.0286* | *Nam-Co_family_66* | *Nam-Co_genus_329* | blastp | *Nam-Co_Duplornaviricota_053* | 84.97 |
| LC_coassembly_k141_2266943_1 | *Duplornaviricota* | *Chrymotiviricetes* | *Ghabrivirales* | *f.0286* | *Nam-Co_family_66* | *Nam-Co_genus_329* | blastp | *Nam-Co_Duplornaviricota_054* | 84.95 |
| LC_0.5m_10um_k141_146743_1 | *Duplornaviricota* | *Chrymotiviricetes* | *Ghabrivirales* | *f.0286* | *Nam-Co_family_64* | *Nam-Co_genus_327* | blastp | *Nam-Co_Duplornaviricota_055* | 88.57 |
| LC_coassembly_k141_1759012_3 | *Duplornaviricota* | *Chrymotiviricetes* | *Ghabrivirales* | *f.0286* | *Nam-Co_family_71* | *Nam-Co_genus_328* | blastp | *Nam-Co_Duplornaviricota_056* | 88.67 |
| LC_coassembly_k141_2599729_1 | *Duplornaviricota* | *Chrymotiviricetes* | *Ghabrivirales* | *f.0287* | *Nam-Co_family_74* | *Nam-Co_genus_331* | blastp | *Nam-Co_Duplornaviricota_057* | 89.41 |
| LC_coassembly_k141_698489_1 | *Duplornaviricota* | *Chrymotiviricetes* | *Ghabrivirales* | *f.0287* |  | *Nam-Co_genus_335* | blastp | *Nam-Co_Duplornaviricota_058* | 90.71 |
| LC_coassembly_k141_2813839_2 | *Duplornaviricota* | *Chrymotiviricetes* | *Ghabrivirales* | *f.0287* |  | *Nam-Co_genus_335* | blastp | *Nam-Co_Duplornaviricota_059* | 90.65 |
| LC_0.5m_10um_k141_217533_1 | *Duplornaviricota* | *Chrymotiviricetes* | *Ghabrivirales* | *f.0287* |  | *Nam-Co_genus_333* | blastp | *Nam-Co_Duplornaviricota_060* | 90.13 |
| LS_0.5m_3um_k141_179493_1 | *Duplornaviricota* | *Chrymotiviricetes* | *Ghabrivirales* | *f.0287* |  |  | blastp | *Nam-Co_Duplornaviricota_061* | 91.77 |
| LC_coassembly_k141_2418593_2 | *Duplornaviricota* | *Chrymotiviricetes* | *Ghabrivirales* | *f.0287* | *Nam-Co_family_74* | *Nam-Co_genus_332* | blastp | *Nam-Co_Duplornaviricota_062* | 92.67 |
| LC_coassembly_k141_2332014_1 | *Duplornaviricota* | *Chrymotiviricetes* | *Ghabrivirales* | *f.0287* |  | *Nam-Co_genus_330* | blastp | *Nam-Co_Duplornaviricota_063* | 90.63 |
| LC_coassembly_k141_709398_1 | *Duplornaviricota* | *Chrymotiviricetes* | *Ghabrivirales* | *f.0287* | *Nam-Co_family_68* | *Nam-Co_genus_334* | blastp | *Nam-Co_Duplornaviricota_064* | 90.22 |
| LC_coassembly_k141_219128_3 | *Duplornaviricota* | *Chrymotiviricetes* | *Ghabrivirales* | *f.0287* | *Nam-Co_family_73* | *Nam-Co_genus_336* | blastp | *Nam-Co_Duplornaviricota_065* | 80.75 |
| LC_20m_3um_k141_60113_1 | *Duplornaviricota* | *Chrymotiviricetes* | *Ghabrivirales* | *f.0287* | *Nam-Co_family_74* | *Nam-Co_genus_332* | blastp | *Nam-Co_Duplornaviricota_066* | 87.58 |
| LS_coassembly_k141_16911_2 | *Duplornaviricota* | *Chrymotiviricetes* | *Ghabrivirales* | *f.0287* |  |  | blastp | *Nam-Co_Duplornaviricota_067* | 91.59 |
| LC_coassembly_k141_2401611_3 | *Duplornaviricota* | *Chrymotiviricetes* | *Ghabrivirales* | *f.0287* | *Nam-Co_family_74* | *Nam-Co_genus_332* | blastp | *Nam-Co_Duplornaviricota_068* | 91.39 |
| LC_coassembly_k141_2269750_2 | *Duplornaviricota* | *Chrymotiviricetes* | *Ghabrivirales* | *f.0292* |  | *Nam-Co_genus_341* | blastp | *Nam-Co_Duplornaviricota_069* | 86.15 |
| LC_coassembly_k141_2732588_2 | *Duplornaviricota* | *Chrymotiviricetes* | *Ghabrivirales* | *f.0292* |  | *Nam-Co_genus_337* | blastp | *Nam-Co_Duplornaviricota_070* | 89.6 |
| LC_coassembly_k141_977866_1 | *Duplornaviricota* | *Chrymotiviricetes* | *Ghabrivirales* | *f.0292* |  | *Nam-Co_genus_338* | blastp | *Nam-Co_Duplornaviricota_071* | 88.31 |
| LS_coassembly_k141_1288836_2 | *Duplornaviricota* | *Chrymotiviricetes* | *Ghabrivirales* | *f.0292* |  | *Nam-Co_genus_339* | blastp | *Nam-Co_Duplornaviricota_072* | 89.13 |
| LC_80m_0.2um_k141_321522_2 | *Duplornaviricota* | *Chrymotiviricetes* | *Ghabrivirales* | *f.0292* |  | *Nam-Co_genus_341* | blastp | *Nam-Co_Duplornaviricota_073* | 87.45 |
| LS_coassembly_k141_542559_1 | *Duplornaviricota* | *Chrymotiviricetes* | *Ghabrivirales* | *f.0292* |  | *Nam-Co_genus_340* | blastp | *Nam-Co_Duplornaviricota_074* | 92.06 |
| LS_0.5m_3um_k141_361849_1 | *Duplornaviricota* | *Chrymotiviricetes* | *Ghabrivirales* | *f.0295* |  | *Nam-Co_genus_342* | blastp | *Nam-Co_Duplornaviricota_075* | 81.31 |
| LC_coassembly_k141_1046725_2 | *Duplornaviricota* | *Chrymotiviricetes* | *Ghabrivirales* | *f.0295* | *Nam-Co_family_70* | *Nam-Co_genus_343* | blastp | *Nam-Co_Duplornaviricota_076* | 83.66 |
| LC_20m_3um_k141_445022_3 | *Duplornaviricota* | *Chrymotiviricetes* | *Ghabrivirales* | *f.0295* |  | *Nam-Co_genus_342* | blastp | *Nam-Co_Duplornaviricota_077* | 83.37 |
| LC_coassembly_k141_1526117_3 | *Duplornaviricota* | *Chrymotiviricetes* | *Ghabrivirales* | *f.0295* |  | *Nam-Co_genus_342* | blastp | *Nam-Co_Duplornaviricota_078* | 82.72 |
| LC_coassembly_k141_1652841_1 | *Duplornaviricota* | *Chrymotiviricetes* | *Ghabrivirales* | *f.0296* |  | *Nam-Co_genus_350* | blastp | *Nam-Co_Duplornaviricota_079* | 92.04 |
| LC_coassembly_k141_1894690_1 | *Duplornaviricota* | *Chrymotiviricetes* | *Ghabrivirales* | *f.0296* |  | *Nam-Co_genus_350* | blastp | *Nam-Co_Duplornaviricota_080* | 92 |
| LC_20m_1.2um_k141_99042_1 | *Duplornaviricota* | *Chrymotiviricetes* | *Ghabrivirales* | *f.0296* |  | *Nam-Co_genus_350* | blastp | *Nam-Co_Duplornaviricota_081* | 90.66 |
| LC_20m_1.2um_k141_21785_1 | *Duplornaviricota* | *Chrymotiviricetes* | *Ghabrivirales* | *f.0296* |  | *Nam-Co_genus_350* | blastp | *Nam-Co_Duplornaviricota_082* | 92.01 |
| LS_coassembly_k141_665176_1 | *Duplornaviricota* | *Chrymotiviricetes* | *Ghabrivirales* | *f.0296* |  | *Nam-Co_genus_350* | blastp | *Nam-Co_Duplornaviricota_083* | 92.21 |
| LC_20m_3um_k141_163697_2 | *Duplornaviricota* | *Chrymotiviricetes* | *Ghabrivirales* | *f.0296* |  | *Nam-Co_genus_346* | blastp | *Nam-Co_Duplornaviricota_084* | 87.02 |
| LC_coassembly_k141_774642_1 | *Duplornaviricota* | *Chrymotiviricetes* | *Ghabrivirales* | *f.0296* | *Nam-Co_family_67* | *Nam-Co_genus_347* | blastp | *Nam-Co_Duplornaviricota_085* | 91.22 |
| LC_20m_10um_k141_54152_1 | *Duplornaviricota* | *Chrymotiviricetes* | *Ghabrivirales* | *f.0296* | *Nam-Co_family_67* | *Nam-Co_genus_347* | blastp | *Nam-Co_Duplornaviricota_086* | 87.69 |
| LC_20m_10um_k141_7746_1 | *Duplornaviricota* | *Chrymotiviricetes* | *Ghabrivirales* | *f.0296* |  | *Nam-Co_genus_350* | blastp | *Nam-Co_Duplornaviricota_087* | 90.61 |
| LC_20m_1.2um_k141_371464_1 | *Duplornaviricota* | *Chrymotiviricetes* | *Ghabrivirales* | *f.0296* | *Nam-Co_family_67* | *Nam-Co_genus_347* | blastp | *Nam-Co_Duplornaviricota_088* | 91.15 |
| LC_0.5m_10um_k141_130680_1 | *Duplornaviricota* | *Chrymotiviricetes* | *Ghabrivirales* | *f.0296* |  | *Nam-Co_genus_346* | blastp | *Nam-Co_Duplornaviricota_089* | 86.35 |
| LC_coassembly_k141_1892891_1 | *Duplornaviricota* | *Chrymotiviricetes* | *Ghabrivirales* | *f.0296* |  | *Nam-Co_genus_346* | blastp | *Nam-Co_Duplornaviricota_090* | 88.81 |
| LC_20m_3um_k141_526662_1 | *Duplornaviricota* | *Chrymotiviricetes* | *Ghabrivirales* | *f.0296* |  | *Nam-Co_genus_350* | blastp | *Nam-Co_Duplornaviricota_091* | 94.8 |
| LC_coassembly_k141_1421666_1 | *Duplornaviricota* | *Chrymotiviricetes* | *Ghabrivirales* | *f.0296* |  | *Nam-Co_genus_346* | blastp | *Nam-Co_Duplornaviricota_092* | 82.92 |
| LC_20m_10um_k141_72899_1 | *Duplornaviricota* | *Chrymotiviricetes* | *Ghabrivirales* | *f.0296* |  | *Nam-Co_genus_350* | blastp | *Nam-Co_Duplornaviricota_093* | 86.48 |
| LC_20m_10um_k141_65967_1 | *Duplornaviricota* | *Chrymotiviricetes* | *Ghabrivirales* | *f.0296* | *Nam-Co_family_67* | *Nam-Co_genus_344* | blastp | *Nam-Co_Duplornaviricota_094* | 91.25 |
| LC_20m_3um_k141_284224_1 | *Duplornaviricota* | *Chrymotiviricetes* | *Ghabrivirales* | *f.0296* |  | *Nam-Co_genus_352* | blastp | *Nam-Co_Duplornaviricota_095* | 92.29 |
| LC_20m_10um_k141_20727_1 | *Duplornaviricota* | *Chrymotiviricetes* | *Ghabrivirales* | *f.0296* |  | *Nam-Co_genus_352* | blastp | *Nam-Co_Duplornaviricota_096* | 91.09 |
| LC_20m_10um_k141_91954_1 | *Duplornaviricota* | *Chrymotiviricetes* | *Ghabrivirales* | *f.0296* |  | *Nam-Co_genus_352* | blastp | *Nam-Co_Duplornaviricota_097* | 91.26 |
| LC_20m_10um_k141_4823_1 | *Duplornaviricota* | *Chrymotiviricetes* | *Ghabrivirales* | *f.0296* |  | *Nam-Co_genus_352* | blastp | *Nam-Co_Duplornaviricota_098* | 91.04 |
| LC_coassembly_k141_2187462_8 | *Duplornaviricota* | *Chrymotiviricetes* | *Ghabrivirales* | *f.0296* |  | *Nam-Co_genus_353* | blastp | *Nam-Co_Duplornaviricota_099* | 88.82 |
| LC_50m_0.2um_k141_256122_2 | *Duplornaviricota* | *Chrymotiviricetes* | *Ghabrivirales* | *f.0296* |  | *Nam-Co_genus_349* | blastp | *Nam-Co_Duplornaviricota_100* | 87.94 |
| LC_coassembly_k141_210205_1 | *Duplornaviricota* | *Chrymotiviricetes* | *Ghabrivirales* | *f.0296* |  | *Nam-Co_genus_350* | blastp | *Nam-Co_Duplornaviricota_101* | 90.31 |
| LC_coassembly_k141_2452354_1 | *Duplornaviricota* | *Chrymotiviricetes* | *Ghabrivirales* | *f.0296* |  | *Nam-Co_genus_350* | blastp | *Nam-Co_Duplornaviricota_102* | 90.74 |
| LC_20m_10um_k141_15219_1 | *Duplornaviricota* | *Chrymotiviricetes* | *Ghabrivirales* | *f.0296* |  | *Nam-Co_genus_350* | blastp | *Nam-Co_Duplornaviricota_103* | 90.69 |
| LC_20m_10um_k141_65367_1 | *Duplornaviricota* | *Chrymotiviricetes* | *Ghabrivirales* | *f.0296* |  | *Nam-Co_genus_350* | blastp | *Nam-Co_Duplornaviricota_104* | 90.67 |
| LC_20m_10um_k141_14302_1 | *Duplornaviricota* | *Chrymotiviricetes* | *Ghabrivirales* | *f.0296* |  | *Nam-Co_genus_350* | blastp | *Nam-Co_Duplornaviricota_105* | 90.59 |
| LC_20m_1.2um_k141_50683_1 | *Duplornaviricota* | *Chrymotiviricetes* | *Ghabrivirales* | *f.0296* | *Nam-Co_family_67* | *Nam-Co_genus_345* | blastp | *Nam-Co_Duplornaviricota_106* | 89.04 |
| LC_coassembly_k141_2762379_1 | *Duplornaviricota* | *Chrymotiviricetes* | *Ghabrivirales* | *f.0296* |  | *Nam-Co_genus_350* | blastp | *Nam-Co_Duplornaviricota_107* | 90.36 |
| LC_coassembly_k141_1533203_1 | *Duplornaviricota* | *Chrymotiviricetes* | *Ghabrivirales* | *f.0296* |  | *Nam-Co_genus_350* | blastp | *Nam-Co_Duplornaviricota_108* | 90.7 |
| LC_coassembly_k141_969841_2 | *Duplornaviricota* | *Chrymotiviricetes* | *Ghabrivirales* | *f.0296* |  | *Nam-Co_genus_348* | blastp | *Nam-Co_Duplornaviricota_109* | 89.71 |
| LC_coassembly_k141_642659_2 | *Duplornaviricota* | *Chrymotiviricetes* | *Ghabrivirales* | *f.0296* |  | *Nam-Co_genus_351* | blastp | *Nam-Co_Duplornaviricota_110* | 85.92 |
| LC_20m_10um_k141_15810_1 | *Duplornaviricota* | *Chrymotiviricetes* | *Ghabrivirales* | *f.0296* |  | *Nam-Co_genus_350* | blastp | *Nam-Co_Duplornaviricota_111* | 90.65 |
| LC_coassembly_k141_420336_1 | *Duplornaviricota* | *Resentoviricetes* | *Reovirales* | *Reoviridae* | *Nam-Co_family_77* | *Nam-Co_genus_361* | blastp | *Nam-Co_Duplornaviricota_112* |  |
| LS_coassembly_k141_436759_1 | *Duplornaviricota* | *Resentoviricetes* | *Reovirales* | *Reoviridae* | *Nam-Co_family_76* | *Nam-Co_genus_358* | blastp | *Nam-Co_Duplornaviricota_113* | 87.37 |
| LS_coassembly_k141_356040_1 | *Duplornaviricota* | *Resentoviricetes* | *Reovirales* | *Reoviridae* | *Nam-Co_family_75* | *Nam-Co_genus_362* | blastp | *Nam-Co_Duplornaviricota_114* | 81.55 |
| LS_coassembly_k141_380972_1 | *Duplornaviricota* | *Resentoviricetes* | *Reovirales* | *Reoviridae* |  | *Nam-Co_genus_363* | blastp | *Nam-Co_Duplornaviricota_115* | 87 |
| LC_coassembly_k141_105435_1 | *Duplornaviricota* | *Resentoviricetes* | *Reovirales* | *Reoviridae* |  | *Nam-Co_genus_360* | blastp | *Nam-Co_Duplornaviricota_116* | 83.16 |
| LS_coassembly_k141_522337_1 | *Duplornaviricota* | *Resentoviricetes* | *Reovirales* | *Reoviridae* |  | *Nam-Co_genus_359* | blastp | *Nam-Co_Duplornaviricota_117* | 86.69 |
| LC_coassembly_k141_2171829_1 | *Duplornaviricota* | *Chrymotiviricetes* | *Ghabrivirales* | *Totiviridae* |  | *Nam-Co_genus_354* | blastp | *Nam-Co_Duplornaviricota_118* | 93.26 |
| LS_coassembly_k141_138809_1 | *Duplornaviricota* | *Chrymotiviricetes* | *Ghabrivirales* | *Totiviridae* |  | *Nam-Co_genus_355* | blastp | *Nam-Co_Duplornaviricota_119* | 86.01 |
| LC_coassembly_k141_1836560_1 | *Duplornaviricota* | *Chrymotiviricetes* | *Ghabrivirales* | *Totiviridae* |  | *Nam-Co_genus_356* | blastp | *Nam-Co_Duplornaviricota_120* | 91.74 |
| LC_coassembly_k141_2783551_3 | *Duplornaviricota* | *Chrymotiviricetes* | *Ghabrivirales* | *Totiviridae* |  | *Nam-Co_genus_356* | blastp | *Nam-Co_Duplornaviricota_121* | 90.85 |
| LC_coassembly_k141_1767469_1 | *Duplornaviricota* | *Chrymotiviricetes* | *Ghabrivirales* | *Totiviridae* |  | *Nam-Co_genus_357* | blastp | *Nam-Co_Duplornaviricota_122* | 89.27 |
| LC_80m_10um_k141_265723_1 | *Duplornaviricota* | *Chrymotiviricetes* | *Ghabrivirales* | *Totiviridae* |  |  | blastp | *Nam-Co_Duplornaviricota_123* | 90.89 |
| LC_coassembly_k141_2570745_1 | *Negarnaviricota* |  |  |  | *Nam-Co_family_79* | *Nam-Co_genus_364* | structure | *Nam-Co_Negarnaviricota_001* | 87.96 |
| LC_50m_3um_k141_453236_1 | *Negarnaviricota* |  |  |  | *Nam-Co_family_79* | *Nam-Co_genus_364* | structure | *Nam-Co_Negarnaviricota_002* | 80.98 |
| LS_coassembly_k141_516141_1 | *Negarnaviricota* |  |  |  | *Nam-Co_family_78* | *Nam-Co_genus_365* | structure | *Nam-Co_Negarnaviricota_003* | 80.18 |
| LC_coassembly_k141_2820748_1 | *Negarnaviricota* | *Milneviricetes* | *Serpentovirales* | *Aspiviridae* | *Nam-Co_family_81* | *Nam-Co_genus_378* | blastp | *Nam-Co_Negarnaviricota_004* | 81.26 |
| LC_coassembly_k141_339916_1 | *Negarnaviricota* | *Ellioviricetes* | *Bunyavirales* | *f.0300.base-Phenui* | *Nam-Co_family_80* | *Nam-Co_genus_366* | blastp | *Nam-Co_Negarnaviricota_005* | 81.01 |
| LC_coassembly_k141_2312099_1 | *Negarnaviricota* | *Ellioviricetes* | *Bunyavirales* | *f.0305.base-Leishbu* |  | *Nam-Co_genus_367* | blastp | *Nam-Co_Negarnaviricota_006* | 90.22 |
| LC_50m_0.2um_k141_322623_1 | *Negarnaviricota* | *Ellioviricetes* | *Bunyavirales* | *f.0313* |  |  | blastp | *Nam-Co_Negarnaviricota_007* | 86.43 |
| LC_coassembly_k141_769811_1 | *Negarnaviricota* | *Ellioviricetes* | *Bunyavirales* | *f.0315* |  | *Nam-Co_genus_368* | blastp | *Nam-Co_Negarnaviricota_008* | 91.64 |
| LC_coassembly_k141_1916082_1 | *Negarnaviricota* | *Ellioviricetes* | *Bunyavirales* | *f.0315* |  | *Nam-Co_genus_370* | blastp | *Nam-Co_Negarnaviricota_009* | 88.13 |
| LC_80m_0.2um_k141_601816_1 | *Negarnaviricota* | *Ellioviricetes* | *Bunyavirales* | *f.0315* |  | *Nam-Co_genus_370* | blastp | *Nam-Co_Negarnaviricota_010* | 86.95 |
| LC_coassembly_k141_662643_2 | *Negarnaviricota* | *Ellioviricetes* | *Bunyavirales* | *f.0315* |  | *Nam-Co_genus_369* | blastp | *Nam-Co_Negarnaviricota_011* | 91.91 |
| LC_coassembly_k141_1407661_1 | *Negarnaviricota* | *Ellioviricetes* | *Bunyavirales* | *f.0315* |  | *Nam-Co_genus_371* | blastp | *Nam-Co_Negarnaviricota_012* | 85.65 |
| LC_coassembly_k141_1863880_1 | *Negarnaviricota* | *Ellioviricetes* | *Bunyavirales* | *f.0327* |  | *Nam-Co_genus_372* | blastp | *Nam-Co_Negarnaviricota_013* | 91.44 |
| LC_coassembly_k141_732085_1 | *Negarnaviricota* | *Ellioviricetes* | *Bunyavirales* | *f.0327* |  | *Nam-Co_genus_374* | blastp | *Nam-Co_Negarnaviricota_014* | 92.34 |
| LS_coassembly_k141_1172872_1 | *Negarnaviricota* | *Ellioviricetes* | *Bunyavirales* | *f.0327* |  | *Nam-Co_genus_373* | blastp | *Nam-Co_Negarnaviricota_015* | 92.68 |
| LC_coassembly_k141_716515_1 | *Negarnaviricota* | *Ellioviricetes* | *Bunyavirales* | *f.0328* |  | *Nam-Co_genus_377* | blastp | *Nam-Co_Negarnaviricota_016* | 90.25 |
| LC_coassembly_k141_455553_1 | *Negarnaviricota* | *Ellioviricetes* | *Bunyavirales* | *f.0328* |  | *Nam-Co_genus_376* | blastp | *Nam-Co_Negarnaviricota_017* | 90.72 |
| LS_coassembly_k141_11576_1 | *Negarnaviricota* | *Ellioviricetes* | *Bunyavirales* | *f.0328* |  | *Nam-Co_genus_375* | blastp | *Nam-Co_Negarnaviricota_018* | 90.59 |
| LS_coassembly_k141_407824_1 | *Negarnaviricota* | *c.0054.base-Yunchang* | *o.0072* | *f.0354* |  |  | blastp | *Nam-Co_Negarnaviricota_019* | 87.66 |
| LS_coassembly_k141_950304_7 | *Negarnaviricota* | *Monjiviricetes* | *Mononegavirales* | *Rhabdoviridae* |  | *Nam-Co_genus_379* | blastp | *Nam-Co_Negarnaviricota_020* | 84.57 |
| LS_coassembly_k141_287825_1 | *Negarnaviricota* | *Monjiviricetes* | *Mononegavirales* | *Rhabdoviridae* |  | *Nam-Co_genus_380* | blastp | *Nam-Co_Negarnaviricota_021* | 89.91 |
| LS_0.5m_0.2um_k141_373946_1 | *unsure* |  |  |  | *Nam-Co_family_83* | *Nam-Co_genus_381* | structure | *Nam-Co_unknown_RdRP_001* | 87.49 |
| LS_coassembly_k141_1167127_2 | *unsure* |  |  |  | *Nam-Co_family_82* | *Nam-Co_genus_382* | structure | *Nam-Co_unknown_RdRP_002* | 85.67 |
| LS_coassembly_k141_755496_2 | *unsure* |  |  |  | *Nam-Co_family_84* | *Nam-Co_genus_383* | structure | *Nam-Co_unknown_RdRP_003* | 61.6 |

| **Table S4. Identification of novel RdRP clades in Lake Nam Co.** | | | | |
| --- | --- | --- | --- | --- |
| **Clade** | **Figure** | **Closest Family** | **Sequence number** | **^1^Pident** |
| *Lenarviricota*_Clade_Ⅰ | S5 | *Narnaviridae* | 4 | 31.5-40.2 |
| *Lenarviricota*_Clade_Ⅱ | S8 | *f.0388* | 4 | 36-39.7 |
| *Pisuviricota*_Clade_Ⅰ | S9 | *f.0025* | 4 | 35.6-40.2 |
| *Pisuviricota*_Clade_Ⅱ | S12 | *f.0102* | 3 | 34.3-38.9 |
| *Pisuviricota*_Clade_Ⅲ | S12 | *f.0102* | 5 | 42.4-44.2 |
| *Pisuviricota*_Clade_Ⅳ | S13 | *f.0111* | 22 | 35.3-44.9 |
| *Pisuviricota*_Clade_Ⅴ | S13 | *f.0111* | 5 | 40.1-47.6 |
| *Pisuviricota*_Clade_Ⅵ | S13 | *f.0111* | 9 | 32.6-38.9 |
| *Ktrinoviricota*_Clade_Ⅰ | S14 | *f.0176* | 45 | 29.4-36.3 |
| *Duplornaviricota*_Clade_Ⅰ | S18 | *f.0296* | 24 | 35.5-41.2 |
| *Duplornaviricota*_Clade_Ⅱ | S18 | *f.0296* | 5 | 40.9-44.6 |
| *Duplornaviricota*_Clade_Ⅲ | S18 | *f.0295* | 3 | 36.1-39 |
| *Duplornaviricota*_Clade_Ⅳ | S19 | *f.0286* | 7 | 25.9-32.5 |
| *Duplornaviricota*_Clade_Ⅴ | S20 | *f.0287* | 4 | 27.5-37.9 |
| 1.Pident: Percentage of identity range with references (%) | | |  |  |

| **Table S5. Basic statistics of RNA vOTUs identified in the Lake Nam Co.** | | | | | | | | |
| --- | --- | --- | --- | --- | --- | --- | --- | --- |
| **RNA vOTU** | **Length** | **Whether encoded complete  RdRp domain** | **Genome composition** | **Phylum** | **Class** | **Order** | **Family** | **Encoded ORFs** |
| LC_0.5m_0.2um_k141_202252 | 1039 | Yes | ssRNA(+) | *Kitrinoviricota* | *Alsuviricetes* | *Hepelivirales* | *Nam-Co_family_50* | 1 |
| LC_0.5m_0.2um_k141_268352 | 1442 | Yes | ssRNA(+) | *Kitrinoviricota* | *Alsuviricetes* | *Hepelivirales* | *Benyviridae* | 2 |
| LC_0.5m_0.2um_k141_300989 | 3380 | Yes | Unknown | *Kitrinoviricota* | *c.0034* | *o.0045* | *f.0176* | 2 |
| LC_0.5m_0.2um_k141_74024 | 1995 | Yes | Unknown | *Kitrinoviricota* | *c.0034* | *o.0045* | *f.0176* | 2 |
| LC_0.5m_0.2um_k141_86628 | 7108 | Yes | ssRNA(+) | *Pisuviricota* | *Pisoniviricetes* | *Picornavirales* | *Dicistroviridae* | 2 |
| LC_0.5m_1.2um_k141_119548 | 916 | Yes | Unknown | *Kitrinoviricota* | *c.0034* | *o.0045* | *f.0176* | 1 |
| LC_0.5m_1.2um_k141_131426 | 677 | No | NA | *NA* | *NA* | *NA* | *NA* | 1 |
| LC_0.5m_1.2um_k141_132419 | 2179 | Yes | Unknown | *Lenarviricota* | *c.0069.base-Howelto* | *o.0090* | *f.0386* | 1 |
| LC_0.5m_1.2um_k141_158692 | 3478 | Yes | ssRNA(+) | *Lenarviricota* | *Amabiliviricetes* | *Wolframvirales* | *Narnaviridae* | 1 |
| LC_0.5m_1.2um_k141_19237 | 1030 | Yes | ssRNA(+) | *Lenarviricota* | *Amabiliviricetes* | *Wolframvirales* | *Narnaviridae* | 1 |
| LC_0.5m_1.2um_k141_20885 | 6955 | Yes | Unknown | *Kitrinoviricota* | *c.0034* | *o.0045* | *f.0176* | 6 |
| LC_0.5m_1.2um_k141_226143 | 9771 | Yes | Unknown | *Duplornaviricota* | *Chrymotiviricetes* | *Ghabrivirales* | *f.0292* | 2 |
| LC_0.5m_1.2um_k141_244369 | 873 | Yes | Unknown | *Kitrinoviricota* | *c.0034* | *o.0045* | *f.0176* | 1 |
| LC_0.5m_1.2um_k141_251615 | 1812 | Yes | ssRNA(+) | *Lenarviricota* | *Amabiliviricetes* | *Wolframvirales* | *Narnaviridae* | 1 |
| LC_0.5m_1.2um_k141_253032 | 2666 | Yes | ssRNA(+) | *Lenarviricota* | *Amabiliviricetes* | *Wolframvirales* | *Narnaviridae* | 1 |
| LC_0.5m_1.2um_k141_276576 | 4973 | Yes | Unknown | *Kitrinoviricota* | *c.0025* | *o.0036* | *f.0145* | 3 |
| LC_0.5m_1.2um_k141_278295 | 1578 | Yes | dsRNA | *Duplornaviricota* | *Chrymotiviricetes* | *Ghabrivirales* | *Totiviridae* | 2 |
| LC_0.5m_1.2um_k141_281657 | 2680 | Yes | Unknown | *Kitrinoviricota* | *c.0034* | *o.0045* | *f.0176* | 2 |
| LC_0.5m_1.2um_k141_28664 | 1210 | Yes | Unknown | *Pisuviricota* | *Duplopiviricetes* | *Durnavirales* | *f.0102.base-Curvula* | 1 |
| LC_0.5m_1.2um_k141_300503 | 837 | No | NA | *NA* | *NA* | *NA* | *NA* | 1 |
| LC_0.5m_1.2um_k141_304317 | 1512 | Yes | ssRNA(+) | *Lenarviricota* | *Amabiliviricetes* | *Wolframvirales* | *Narnaviridae* | 1 |
| LC_0.5m_1.2um_k141_316701 | 1214 | Yes | ssRNA(+) | *Lenarviricota* | *Amabiliviricetes* | *Wolframvirales* | *Narnaviridae* | 1 |
| LC_0.5m_1.2um_k141_34913 | 2030 | Yes | Unknown | *Kitrinoviricota* | *c.0034* | *o.0045* | *f.0176* | 2 |
| LC_0.5m_1.2um_k141_36111 | 7221 | Yes | Unknown | *Kitrinoviricota* | *c.0034* | *o.0045* | *f.0176* | 6 |
| LC_0.5m_1.2um_k141_379089 | 1257 | Yes | ssRNA(+) | *Lenarviricota* | *Amabiliviricetes* | *Wolframvirales* | *Narnaviridae* | 1 |
| LC_0.5m_1.2um_k141_379937 | 1255 | Yes | ssRNA(+) | *Lenarviricota* | *Howeltoviricetes* | *Cryppavirales* | *Mitoviridae* | 1 |
| LC_0.5m_1.2um_k141_396381 | 9854 | Yes | Unknown | *Kitrinoviricota* | *c.0034* | *o.0045* | *f.0176* | 7 |
| LC_0.5m_1.2um_k141_403526 | 3551 | Yes | Unknown | *Kitrinoviricota* | *c.0034* | *o.0045* | *f.0176* | 3 |
| LC_0.5m_1.2um_k141_98739 | 1000 | Yes | ssRNA(+) | *Lenarviricota* | *Amabiliviricetes* | *Wolframvirales* | *Narnaviridae* | 1 |
| LC_0.5m_10um_k141_109854 | 4624 | Yes | ssRNA(+) | *Kitrinoviricota* | *Alsuviricetes* | *o.0055* | *Endornaviridae* | 2 |
| LC_0.5m_10um_k141_124243 | 4974 | Yes | ssRNA(+) | *Kitrinoviricota* | *Alsuviricetes* | *Martellivirales* | *Virgaviridae* | 5 |
| LC_0.5m_10um_k141_124797 | 926 | Yes | Unknown | *Duplornaviricota* | *Chrymotiviricetes* | *Ghabrivirales* | *f.0296* | 1 |
| LC_0.5m_10um_k141_129489 | 4763 | Yes | Unknown | *Kitrinoviricota* | *c.0034* | *o.0045* | *f.0176* | 3 |
| LC_0.5m_10um_k141_140412 | 3484 | Yes | Unknown | *Lenarviricota* | *c.0064.base-Howelto* | *o.0085* | *f.0381* | 2 |
| LC_0.5m_10um_k141_14159 | 8167 | Yes | Unknown | *Kitrinoviricota* | *c.0034* | *o.0045* | *f.0176* | 7 |
| LC_0.5m_10um_k141_146743 | 1974 | Yes | Unknown | *Duplornaviricota* | *Chrymotiviricetes* | *Ghabrivirales* | *Nam-Co_family_64* | 1 |
| LC_0.5m_10um_k141_153080 | 2261 | Yes | ssRNA(+) | *Lenarviricota* | *Amabiliviricetes* | *Wolframvirales* | *Narnaviridae* | 1 |
| LC_0.5m_10um_k141_153862 | 3073 | Yes | Unknown | *Duplornaviricota* | *Chrymotiviricetes* | *Ghabrivirales* | *f.0287* | 4 |
| LC_0.5m_10um_k141_170355 | 697 | Yes | Unknown | *Duplornaviricota* | *Chrymotiviricetes* | *Ghabrivirales* | *f.0296* | 1 |
| LC_0.5m_10um_k141_1755 | 2034 | Yes | ssRNA(+) | *Lenarviricota* | *Howeltoviricetes* | *Cryppavirales* | *Mitoviridae* | 1 |
| LC_0.5m_10um_k141_184308 | 810 | Yes | Unknown | *Duplornaviricota* | *Chrymotiviricetes* | *Ghabrivirales* | *f.0283.base-Megabirna* | 1 |
| LC_0.5m_10um_k141_193132 | 2152 | Yes | ssRNA(+) | *Lenarviricota* | *Howeltoviricetes* | *Cryppavirales* | *Mitoviridae* | 2 |
| LC_0.5m_10um_k141_200615 | 8001 | Yes | Unknown | *Kitrinoviricota* | *c.0034* | *o.0045* | *f.0176* | 3 |
| LC_0.5m_10um_k141_217533 | 7269 | Yes | Unknown | *Duplornaviricota* | *Chrymotiviricetes* | *Ghabrivirales* | *f.0287* | 1 |
| LC_0.5m_10um_k141_28114 | 1989 | Yes | ssRNA(+) | *Lenarviricota* | *Howeltoviricetes* | *Cryppavirales* | *Mitoviridae* | 1 |
| LC_0.5m_10um_k141_31696 | 796 | Yes | Unknown | *Duplornaviricota* | *Chrymotiviricetes* | *Ghabrivirales* | *f.0296* | 1 |
| LC_0.5m_10um_k141_4899 | 940 | Yes | ssRNA(+) | *Lenarviricota* | *Howeltoviricetes* | *Cryppavirales* | *Mitoviridae* | 1 |
| LC_0.5m_10um_k141_74703 | 1342 | Yes | Unknown | *Duplornaviricota* | *Chrymotiviricetes* | *Ghabrivirales* | *f.0296* | 1 |
| LC_0.5m_10um_k141_74781 | 918 | Yes | Unknown | *Negarnaviricota* | *Ellioviricetes* | *Bunyavirales* | *f.0328* | 1 |
| LC_0.5m_3um_k141_107358 | 841 | No | NA | *NA* | *NA* | *NA* | *NA* | 1 |
| LC_0.5m_3um_k141_126591 | 1762 | Yes | Unknown | *Pisuviricota* | *Duplopiviricetes* | *Durnavirales* | *f.0111* | 1 |
| LC_0.5m_3um_k141_130993 | 1291 | Yes | ssRNA(+) | *Lenarviricota* | *Howeltoviricetes* | *Cryppavirales* | *Mitoviridae* | 2 |
| LC_0.5m_3um_k141_162910 | 949 | Yes | Unknown | *Kitrinoviricota* | *c.0034* | *o.0045* | *f.0176* | 1 |
| LC_0.5m_3um_k141_163067 | 2217 | Yes | Unknown | *Kitrinoviricota* | *c.0034* | *o.0045* | *f.0176* | 3 |
| LC_0.5m_3um_k141_167927 | 1311 | Yes | Unknown | *Pisuviricota* | *Duplopiviricetes* | *Durnavirales* | *Nam-Co_family_39* | 2 |
| LC_0.5m_3um_k141_193515 | 1547 | Yes | ssRNA(+) | *Lenarviricota* | *Howeltoviricetes* | *Cryppavirales* | *Mitoviridae* | 1 |
| LC_0.5m_3um_k141_25203 | 980 | Yes | Unknown | *Duplornaviricota* | *Chrymotiviricetes* | *Ghabrivirales* | *Nam-Co_family_66* | 1 |
| LC_0.5m_3um_k141_255266 | 2193 | Yes | Unknown | *Pisuviricota* | *Duplopiviricetes* | *Durnavirales* | *f.0102.base-Curvula* | 3 |
| LC_0.5m_3um_k141_257773 | 1966 | Yes | ssRNA(+) | *Lenarviricota* | *Amabiliviricetes* | *Wolframvirales* | *Narnaviridae* | 1 |
| LC_0.5m_3um_k141_258279 | 8580 | Yes | Unknown | *Kitrinoviricota* | *c.0034* | *o.0045* | *f.0176* | 7 |
| LC_0.5m_3um_k141_265610 | 8531 | Yes | Unknown | *Kitrinoviricota* | *c.0034* | *o.0045* | *f.0176* | 6 |
| LC_0.5m_3um_k141_309296 | 787 | Yes | ssRNA(+) | *Lenarviricota* | *Howeltoviricetes* | *Cryppavirales* | *Mitoviridae* | 1 |
| LC_0.5m_3um_k141_311310 | 3300 | Yes | Unknown | *Kitrinoviricota* | *c.0034* | *o.0045* | *f.0176* | 3 |
| LC_0.5m_3um_k141_350484 | 7373 | Yes | Unknown | *Kitrinoviricota* | *c.0034* | *o.0045* | *f.0176* | 6 |
| LC_0.5m_3um_k141_36449 | 8374 | Yes | Unknown | *Kitrinoviricota* | *c.0034* | *o.0045* | *f.0176* | 6 |
| LC_0.5m_3um_k141_375717 | 870 | Yes | ssRNA(+) | *Lenarviricota* | *Howeltoviricetes* | *Cryppavirales* | *Mitoviridae* | 1 |
| LC_0.5m_3um_k141_383662 | 1451 | Yes | ssRNA(+) | *Lenarviricota* | *Amabiliviricetes* | *Wolframvirales* | *Narnaviridae* | 1 |
| LC_0.5m_3um_k141_392533 | 3376 | No | NA | *NA* | *NA* | *NA* | *NA* | 2 |
| LC_0.5m_3um_k141_396598 | 2649 | Yes | Unknown | *Duplornaviricota* | *Chrymotiviricetes* | *Ghabrivirales* | *Nam-Co_family_66* | 1 |
| LC_0.5m_3um_k141_398529 | 1682 | Yes | Unknown | *Kitrinoviricota* | *c.0034* | *o.0045* | *f.0176* | 2 |
| LC_0.5m_3um_k141_431029 | 7226 | Yes | Unknown | *Duplornaviricota* | *Chrymotiviricetes* | *Ghabrivirales* | *f.0287* | 2 |
| LC_0.5m_3um_k141_440210 | 1676 | Yes | ssRNA(+) | *Lenarviricota* | *Miaviricetes* | *Ourlivirales* | *Botourmiaviridae* | 1 |
| LC_0.5m_3um_k141_81056 | 759 | Yes | ssRNA(+) | *Lenarviricota* | *Howeltoviricetes* | *Cryppavirales* | *Mitoviridae* | 1 |
| LC_0.5m_3um_k141_95027 | 1298 | Yes | Unknown | *Kitrinoviricota* | *c.0034* | *o.0045* | *f.0176* | 2 |
| LC_20m_0.2um_k141_109192 | 830 | Yes | Unknown | *Duplornaviricota* | *Chrymotiviricetes* | *Ghabrivirales* | *f.0296* | 1 |
| LC_20m_0.2um_k141_127815 | 1432 | Yes | Unknown | *Kitrinoviricota* | *c.0034* | *o.0045* | *f.0176* | 2 |
| LC_20m_0.2um_k141_147742 | 1870 | Yes | Unknown | *Kitrinoviricota* | *c.0034* | *o.0045* | *f.0176* | 1 |
| LC_20m_0.2um_k141_158269 | 1676 | Yes | Unknown | *Pisuviricota* | *Duplopiviricetes* | *Durnavirales* | *f.0111* | 2 |
| LC_20m_0.2um_k141_162039 | 777 | Yes | ssRNA(+) | *Pisuviricota* | *Pisoniviricetes* | *Picornavirales* | *Marnaviridae* | 1 |
| LC_20m_0.2um_k141_16816 | 1081 | Yes | Unknown | *Duplornaviricota* | *Chrymotiviricetes* | *Ghabrivirales* | *f.0283.base-Megabirna* | 1 |
| LC_20m_0.2um_k141_169023 | 832 | Yes | Unknown | *Duplornaviricota* | *Chrymotiviricetes* | *Ghabrivirales* | *f.0283.base-Megabirna* | 1 |
| LC_20m_0.2um_k141_177655 | 3737 | Yes | ssRNA(+) | *Pisuviricota* | *Pisoniviricetes* | *Picornavirales* | *Dicistroviridae* | 2 |
| LC_20m_0.2um_k141_207851 | 1415 | No | NA | *NA* | *NA* | *NA* | *NA* | 2 |
| LC_20m_0.2um_k141_243185 | 2007 | Yes | Unknown | *Pisuviricota* | *Pisoniviricetes* | *Picornavirales* | *f.0008.base-Polycipi* | 1 |
| LC_20m_0.2um_k141_250544 | 4297 | Yes | ssRNA(+) | *Pisuviricota* | *Pisoniviricetes* | *Picornavirales* | *Marnaviridae* | 2 |
| LC_20m_0.2um_k141_260174 | 3477 | Yes | Unknown | *Pisuviricota* | *Pisoniviricetes* | *Picornavirales* | *f.0028.base-Dicistro* | 1 |
| LC_20m_0.2um_k141_261093 | 1743 | Yes | Unknown | *Duplornaviricota* | *Chrymotiviricetes* | *Ghabrivirales* | *f.0295* | 1 |
| LC_20m_0.2um_k141_263465 | 958 | Yes | Unknown | *Negarnaviricota* | *Ellioviricetes* | *Bunyavirales* | *f.0315* | 1 |
| LC_20m_0.2um_k141_268307 | 9856 | Yes | Unknown | *Kitrinoviricota* | *c.0034* | *o.0045* | *f.0176* | 5 |
| LC_20m_0.2um_k141_269092 | 11883 | Yes | ssRNA(+) | *Kitrinoviricota* | *Alsuviricetes* | *Hepelivirales* | *Benyviridae* | 6 |
| LC_20m_0.2um_k141_301076 | 3985 | Yes | Unknown | *Kitrinoviricota* | *c.0034* | *o.0045* | *f.0176* | 3 |
| LC_20m_0.2um_k141_305507 | 4371 | No | NA | *NA* | *NA* | *NA* | *NA* | 1 |
| LC_20m_0.2um_k141_308140 | 7001 | Yes | ssRNA(+) | *Pisuviricota* | *Pisoniviricetes* | *Picornavirales* | *Marnaviridae* | 2 |
| LC_20m_0.2um_k141_31373 | 2434 | Yes | Unknown | *Kitrinoviricota* | *c.0034* | *o.0045* | *f.0176* | 1 |
| LC_20m_0.2um_k141_315972 | 3867 | Yes | ssRNA(+) | *Pisuviricota* | *Pisoniviricetes* | *Picornavirales* | *Marnaviridae* | 1 |
| LC_20m_0.2um_k141_316934 | 4629 | Yes | Unknown | *Kitrinoviricota* | *c.0034* | *o.0045* | *f.0176* | 5 |
| LC_20m_0.2um_k141_317770 | 2207 | Yes | ssRNA(+) | *Kitrinoviricota* | *Magsaviricetes* | *Nodamuvirales* | *Nodaviridae* | 1 |
| LC_20m_0.2um_k141_332038 | 775 | Yes | Unknown | *Duplornaviricota* | *Chrymotiviricetes* | *Ghabrivirales* | *f.0296* | 1 |
| LC_20m_0.2um_k141_332893 | 898 | Yes | Unknown | *Pisuviricota* | *c.0013* | *o.0025.base-Sobeli* | *f.0076* | 1 |
| LC_20m_0.2um_k141_334030 | 2946 | Yes | ssRNA(+) | *Kitrinoviricota* | *Tolucaviricetes* | *Tolivirales* | *Tombusviridae* | 3 |
| LC_20m_0.2um_k141_362396 | 832 | Yes | Unknown | *Pisuviricota* | *c.0013* | *o.0025.base-Sobeli* | *f.0076* | 1 |
| LC_20m_0.2um_k141_366279 | 1463 | Yes | Unknown | *Kitrinoviricota* | *c.0034* | *o.0045* | *f.0176* | 2 |
| LC_20m_0.2um_k141_367775 | 3447 | Yes | ssRNA(+) | *Lenarviricota* | *Amabiliviricetes* | *Wolframvirales* | *Narnaviridae* | 1 |
| LC_20m_0.2um_k141_394709 | 876 | Yes | Unknown | *Kitrinoviricota* | *c.0032* | *o.0043* | *f.0167* | 1 |
| LC_20m_0.2um_k141_39978 | 2542 | Yes | ssRNA(+) | *Kitrinoviricota* | *Tolucaviricetes* | *Tolivirales* | *Tombusviridae* | 3 |
| LC_20m_0.2um_k141_403545 | 2420 | Yes | Unknown | *Pisuviricota* | *c.0013* | *o.0025.base-Sobeli* | *f.0076* | 1 |
| LC_20m_0.2um_k141_416520 | 926 | Yes | ssRNA(+) | *Lenarviricota* | *Amabiliviricetes* | *Wolframvirales* | *Narnaviridae* | 1 |
| LC_20m_0.2um_k141_419721 | 696 | Yes | ssRNA(+) | *Lenarviricota* | *Miaviricetes* | *Ourlivirales* | *Botourmiaviridae* | 1 |
| LC_20m_0.2um_k141_60959 | 2561 | Yes | Unknown | *Kitrinoviricota* | *c.0028* | *o.0039* | *f.0150* | 3 |
| LC_20m_0.2um_k141_70484 | 699 | No | NA | *NA* | *NA* | *NA* | *NA* | 1 |
| LC_20m_0.2um_k141_71229 | 946 | Yes | Unknown | *Pisuviricota* | *c.0013* | *o.0025.base-Sobeli* | *f.0076* | 1 |
| LC_20m_0.2um_k141_72846 | 2038 | Yes | Unknown | *Lenarviricota* | *c.0069.base-Howelto* | *o.0090* | *f.0386* | 1 |
| LC_20m_0.2um_k141_76919 | 5406 | Yes | Unknown | *Kitrinoviricota* | *c.0034* | *o.0045* | *f.0176* | 7 |
| LC_20m_0.2um_k141_79900 | 1652 | Yes | ssRNA(+) | *Lenarviricota* | *Amabiliviricetes* | *Wolframvirales* | *Narnaviridae* | 1 |
| LC_20m_1.2um_k141_139004 | 4768 | Yes | Unknown | *Kitrinoviricota* | *c.0034* | *o.0045* | *f.0176* | 2 |
| LC_20m_1.2um_k141_143448 | 991 | Yes | ssRNA(+) | *Lenarviricota* | *Howeltoviricetes* | *Cryppavirales* | *Mitoviridae* | 1 |
| LC_20m_1.2um_k141_147958 | 3938 | Yes | Unknown | *Kitrinoviricota* | *c.0034* | *o.0045* | *f.0176* | 3 |
| LC_20m_1.2um_k141_15803 | 2387 | Yes | ssRNA(+) | *Lenarviricota* | *Howeltoviricetes* | *Cryppavirales* | *Mitoviridae* | 1 |
| LC_20m_1.2um_k141_21785 | 792 | Yes | Unknown | *Duplornaviricota* | *Chrymotiviricetes* | *Ghabrivirales* | *f.0296* | 1 |
| LC_20m_1.2um_k141_246695 | 2523 | Yes | ssRNA(+) | *Lenarviricota* | *Amabiliviricetes* | *Wolframvirales* | *Narnaviridae* | 1 |
| LC_20m_1.2um_k141_257976 | 7218 | Yes | Unknown | *Kitrinoviricota* | *c.0034* | *o.0045* | *f.0176* | 6 |
| LC_20m_1.2um_k141_266503 | 8457 | Yes | Unknown | *Kitrinoviricota* | *c.0034* | *o.0045* | *f.0176* | 5 |
| LC_20m_1.2um_k141_292435 | 907 | Yes | Unknown | *Duplornaviricota* | *Chrymotiviricetes* | *Ghabrivirales* | *f.0296* | 2 |
| LC_20m_1.2um_k141_302326 | 8570 | Yes | Unknown | *Kitrinoviricota* | *c.0034* | *o.0045* | *f.0176* | 6 |
| LC_20m_1.2um_k141_315533 | 2879 | Yes | Unknown | *Pisuviricota* | *Duplopiviricetes* | *Durnavirales* | *f.0102.base-Curvula* | 2 |
| LC_20m_1.2um_k141_365480 | 1120 | Yes | Unknown | *Kitrinoviricota* | *c.0034* | *o.0045* | *f.0176* | 2 |
| LC_20m_1.2um_k141_371464 | 1324 | Yes | Unknown | *Duplornaviricota* | *Chrymotiviricetes* | *Ghabrivirales* | *Nam-Co_family_67* | 2 |
| LC_20m_1.2um_k141_39093 | 1083 | Yes | Unknown | *Duplornaviricota* | *Chrymotiviricetes* | *Ghabrivirales* | *f.0296* | 1 |
| LC_20m_1.2um_k141_397998 | 7188 | Yes | Unknown | *Kitrinoviricota* | *c.0034* | *o.0045* | *f.0176* | 6 |
| LC_20m_1.2um_k141_408408 | 5829 | Yes | Unknown | *Kitrinoviricota* | *c.0034* | *o.0045* | *f.0176* | 5 |
| LC_20m_1.2um_k141_5914 | 3870 | Yes | Unknown | *Kitrinoviricota* | *c.0034* | *o.0045* | *f.0176* | 4 |
| LC_20m_1.2um_k141_61849 | 952 | Yes | Unknown | *Duplornaviricota* | *Chrymotiviricetes* | *Ghabrivirales* | *f.0296* | 1 |
| LC_20m_1.2um_k141_94657 | 2575 | Yes | Unknown | *Duplornaviricota* | *Chrymotiviricetes* | *Ghabrivirales* | *f.0295* | 1 |
| LC_20m_1.2um_k141_95619 | 664 | No | NA | *NA* | *NA* | *NA* | *NA* | 1 |
| LC_20m_1.2um_k141_99042 | 1165 | Yes | Unknown | *Duplornaviricota* | *Chrymotiviricetes* | *Ghabrivirales* | *f.0296* | 1 |
| LC_20m_10um_k141_12100 | 1769 | Yes | Unknown | *Duplornaviricota* | *Chrymotiviricetes* | *Ghabrivirales* | *f.0283.base-Megabirna* | 1 |
| LC_20m_10um_k141_12836 | 785 | Yes | Unknown | *Duplornaviricota* | *Chrymotiviricetes* | *Ghabrivirales* | *f.0283.base-Megabirna* | 1 |
| LC_20m_10um_k141_14302 | 2023 | Yes | Unknown | *Duplornaviricota* | *Chrymotiviricetes* | *Ghabrivirales* | *f.0296* | 1 |
| LC_20m_10um_k141_1489 | 939 | Yes | Unknown | *Duplornaviricota* | *Chrymotiviricetes* | *Ghabrivirales* | *f.0283.base-Megabirna* | 1 |
| LC_20m_10um_k141_15219 | 1471 | Yes | Unknown | *Duplornaviricota* | *Chrymotiviricetes* | *Ghabrivirales* | *f.0296* | 1 |
| LC_20m_10um_k141_15810 | 1371 | Yes | Unknown | *Duplornaviricota* | *Chrymotiviricetes* | *Ghabrivirales* | *f.0296* | 1 |
| LC_20m_10um_k141_19732 | 2225 | Yes | ssRNA(+) | *Lenarviricota* | *Howeltoviricetes* | *Cryppavirales* | *Mitoviridae* | 2 |
| LC_20m_10um_k141_20727 | 2196 | Yes | Unknown | *Duplornaviricota* | *Chrymotiviricetes* | *Ghabrivirales* | *f.0296* | 1 |
| LC_20m_10um_k141_21145 | 701 | Yes | Unknown | *Duplornaviricota* | *Chrymotiviricetes* | *Ghabrivirales* | *f.0283.base-Megabirna* | 1 |
| LC_20m_10um_k141_21534 | 1069 | Yes | Unknown | *Duplornaviricota* | *Chrymotiviricetes* | *Ghabrivirales* | *f.0283.base-Megabirna* | 1 |
| LC_20m_10um_k141_22158 | 1267 | Yes | Unknown | *Duplornaviricota* | *Chrymotiviricetes* | *Ghabrivirales* | *f.0283.base-Megabirna* | 1 |
| LC_20m_10um_k141_2297 | 1013 | Yes | Unknown | *Duplornaviricota* | *Chrymotiviricetes* | *Ghabrivirales* | *f.0283.base-Megabirna* | 1 |
| LC_20m_10um_k141_27694 | 909 | Yes | Unknown | *Duplornaviricota* | *Chrymotiviricetes* | *Ghabrivirales* | *f.0283.base-Megabirna* | 1 |
| LC_20m_10um_k141_27699 | 774 | Yes | Unknown | *Duplornaviricota* | *Chrymotiviricetes* | *Ghabrivirales* | *f.0296* | 1 |
| LC_20m_10um_k141_33231 | 1455 | Yes | Unknown | *Duplornaviricota* | *Chrymotiviricetes* | *Ghabrivirales* | *f.0296* | 1 |
| LC_20m_10um_k141_33798 | 956 | Yes | Unknown | *Duplornaviricota* | *Chrymotiviricetes* | *Ghabrivirales* | *f.0283.base-Megabirna* | 2 |
| LC_20m_10um_k141_36758 | 1100 | Yes | Unknown | *Duplornaviricota* | *Chrymotiviricetes* | *Ghabrivirales* | *f.0283.base-Megabirna* | 1 |
| LC_20m_10um_k141_38328 | 944 | Yes | Unknown | *Duplornaviricota* | *Chrymotiviricetes* | *Ghabrivirales* | *f.0283.base-Megabirna* | 1 |
| LC_20m_10um_k141_39384 | 634 | Yes | Unknown | *Duplornaviricota* | *Chrymotiviricetes* | *Ghabrivirales* | *f.0296* | 1 |
| LC_20m_10um_k141_42987 | 1404 | Yes | Unknown | *Duplornaviricota* | *Chrymotiviricetes* | *Ghabrivirales* | *f.0283.base-Megabirna* | 1 |
| LC_20m_10um_k141_45308 | 1179 | Yes | Unknown | *Duplornaviricota* | *Chrymotiviricetes* | *Ghabrivirales* | *f.0283.base-Megabirna* | 1 |
| LC_20m_10um_k141_46604 | 1247 | Yes | Unknown | *Duplornaviricota* | *Chrymotiviricetes* | *Ghabrivirales* | *f.0283.base-Megabirna* | 1 |
| LC_20m_10um_k141_47367 | 1450 | Yes | Unknown | *Duplornaviricota* | *Chrymotiviricetes* | *Ghabrivirales* | *f.0283.base-Megabirna* | 1 |
| LC_20m_10um_k141_4823 | 3462 | Yes | Unknown | *Duplornaviricota* | *Chrymotiviricetes* | *Ghabrivirales* | *f.0296* | 1 |
| LC_20m_10um_k141_48657 | 680 | Yes | Unknown | *Duplornaviricota* | *Chrymotiviricetes* | *Ghabrivirales* | *f.0283.base-Megabirna* | 1 |
| LC_20m_10um_k141_49719 | 702 | Yes | Unknown | *Kitrinoviricota* | *c.0034* | *o.0045* | *f.0176* | 2 |
| LC_20m_10um_k141_52727 | 8403 | Yes | Unknown | *Duplornaviricota* | *Chrymotiviricetes* | *Ghabrivirales* | *f.0295* | 2 |
| LC_20m_10um_k141_54152 | 991 | Yes | Unknown | *Duplornaviricota* | *Chrymotiviricetes* | *Ghabrivirales* | *Nam-Co_family_67* | 1 |
| LC_20m_10um_k141_55953 | 1040 | Yes | Unknown | *Duplornaviricota* | *Chrymotiviricetes* | *Ghabrivirales* | *f.0283.base-Megabirna* | 1 |
| LC_20m_10um_k141_57615 | 1264 | Yes | Unknown | *Duplornaviricota* | *Chrymotiviricetes* | *Ghabrivirales* | *f.0283.base-Megabirna* | 1 |
| LC_20m_10um_k141_60255 | 3148 | Yes | Unknown | *Duplornaviricota* | *Chrymotiviricetes* | *Ghabrivirales* | *f.0296* | 2 |
| LC_20m_10um_k141_63441 | 2582 | Yes | Unknown | *Duplornaviricota* | *Chrymotiviricetes* | *Ghabrivirales* | *f.0296* | 2 |
| LC_20m_10um_k141_63442 | 1281 | Yes | Unknown | *Duplornaviricota* | *Chrymotiviricetes* | *Ghabrivirales* | *f.0283.base-Megabirna* | 1 |
| LC_20m_10um_k141_63754 | 651 | Yes | Unknown | *Duplornaviricota* | *Chrymotiviricetes* | *Ghabrivirales* | *f.0283.base-Megabirna* | 1 |
| LC_20m_10um_k141_65367 | 1578 | Yes | Unknown | *Duplornaviricota* | *Chrymotiviricetes* | *Ghabrivirales* | *f.0296* | 1 |
| LC_20m_10um_k141_65967 | 1488 | Yes | Unknown | *Duplornaviricota* | *Chrymotiviricetes* | *Ghabrivirales* | *Nam-Co_family_67* | 1 |
| LC_20m_10um_k141_6666 | 913 | Yes | Unknown | *Duplornaviricota* | *Chrymotiviricetes* | *Ghabrivirales* | *f.0283.base-Megabirna* | 1 |
| LC_20m_10um_k141_68216 | 844 | Yes | Unknown | *Duplornaviricota* | *Chrymotiviricetes* | *Ghabrivirales* | *f.0296* | 1 |
| LC_20m_10um_k141_70765 | 2100 | Yes | Unknown | *Pisuviricota* | *Duplopiviricetes* | *Durnavirales* | *Nam-Co_family_32* | 2 |
| LC_20m_10um_k141_70830 | 1080 | Yes | Unknown | *Duplornaviricota* | *Chrymotiviricetes* | *Ghabrivirales* | *f.0283.base-Megabirna* | 1 |
| LC_20m_10um_k141_72899 | 834 | Yes | Unknown | *Duplornaviricota* | *Chrymotiviricetes* | *Ghabrivirales* | *f.0296* | 2 |
| LC_20m_10um_k141_76613 | 1849 | Yes | ssRNA(+) | *Lenarviricota* | *Amabiliviricetes* | *Wolframvirales* | *Nam-Co_family_07* | 1 |
| LC_20m_10um_k141_7746 | 1455 | Yes | Unknown | *Duplornaviricota* | *Chrymotiviricetes* | *Ghabrivirales* | *f.0296* | 1 |
| LC_20m_10um_k141_8094 | 1076 | Yes | Unknown | *Duplornaviricota* | *Chrymotiviricetes* | *Ghabrivirales* | *f.0296* | 1 |
| LC_20m_10um_k141_90094 | 810 | Yes | Unknown | *Duplornaviricota* | *Chrymotiviricetes* | *Ghabrivirales* | *f.0283.base-Megabirna* | 1 |
| LC_20m_10um_k141_92587 | 699 | Yes | Unknown | *Duplornaviricota* | *Chrymotiviricetes* | *Ghabrivirales* | *f.0283.base-Megabirna* | 1 |
| LC_20m_10um_k141_92634 | 1416 | Yes | Unknown | *Duplornaviricota* | *Chrymotiviricetes* | *Ghabrivirales* | *f.0283.base-Megabirna* | 1 |
| LC_20m_3um_k141_102044 | 2051 | Yes | Unknown | *Pisuviricota* | *Duplopiviricetes* | *Durnavirales* | *f.0111* | 2 |
| LC_20m_3um_k141_11485 | 3577 | Yes | Unknown | *Duplornaviricota* | *Chrymotiviricetes* | *Ghabrivirales* | *Nam-Co_family_66* | 1 |
| LC_20m_3um_k141_115819 | 5096 | Yes | Unknown | *Kitrinoviricota* | *c.0034* | *o.0045* | *f.0176* | 5 |
| LC_20m_3um_k141_118459 | 2220 | Yes | ssRNA(+) | *Lenarviricota* | *Howeltoviricetes* | *Cryppavirales* | *Mitoviridae* | 2 |
| LC_20m_3um_k141_120178 | 1772 | Yes | Unknown | *Kitrinoviricota* | *c.0034* | *o.0045* | *f.0176* | 1 |
| LC_20m_3um_k141_122481 | 3607 | Yes | ssRNA(+) | *Lenarviricota* | *Howeltoviricetes* | *Cryppavirales* | *Mitoviridae* | 1 |
| LC_20m_3um_k141_132526 | 2843 | Yes | Unknown | *Duplornaviricota* | *Chrymotiviricetes* | *Ghabrivirales* | *Nam-Co_family_66* | 1 |
| LC_20m_3um_k141_142581 | 1277 | Yes | ssRNA(+) | *Lenarviricota* | *Amabiliviricetes* | *Wolframvirales* | *Narnaviridae* | 1 |
| LC_20m_3um_k141_14922 | 7063 | Yes | Unknown | *Kitrinoviricota* | *c.0034* | *o.0045* | *f.0176* | 4 |
| LC_20m_3um_k141_163697 | 2573 | Yes | Unknown | *Duplornaviricota* | *Chrymotiviricetes* | *Ghabrivirales* | *f.0296* | 3 |
| LC_20m_3um_k141_171088 | 868 | Yes | Unknown | *Duplornaviricota* | *Chrymotiviricetes* | *Ghabrivirales* | *f.0283.base-Megabirna* | 1 |
| LC_20m_3um_k141_197116 | 2019 | No | NA | *NA* | *NA* | *NA* | *NA* | 1 |
| LC_20m_3um_k141_197713 | 1271 | Yes | ssRNA(+) | *Lenarviricota* | *Amabiliviricetes* | *Wolframvirales* | *Narnaviridae* | 1 |
| LC_20m_3um_k141_227703 | 4152 | Yes | Unknown | *Pisuviricota* | *Duplopiviricetes* | *Durnavirales* | *f.0102.base-Curvula* | 3 |
| LC_20m_3um_k141_251991 | 2265 | Yes | ssRNA(+) | *Lenarviricota* | *Amabiliviricetes* | *Wolframvirales* | *Narnaviridae* | 2 |
| LC_20m_3um_k141_254313 | 1468 | Yes | ssRNA(+) | *Lenarviricota* | *Howeltoviricetes* | *Cryppavirales* | *Mitoviridae* | 1 |
| LC_20m_3um_k141_261003 | 644 | Yes | ssRNA(+) | *Lenarviricota* | *Howeltoviricetes* | *Cryppavirales* | *Mitoviridae* | 1 |
| LC_20m_3um_k141_267098 | 1960 | No | NA | *NA* | *NA* | *NA* | *Nam-Co_family_24* | 2 |
| LC_20m_3um_k141_276645 | 1101 | Yes | Unknown | *Duplornaviricota* | *Chrymotiviricetes* | *Ghabrivirales* | *f.0287* | 1 |
| LC_20m_3um_k141_283152 | 1497 | No | NA | *NA* | *NA* | *NA* | *NA* | 2 |
| LC_20m_3um_k141_284224 | 1691 | Yes | Unknown | *Duplornaviricota* | *Chrymotiviricetes* | *Ghabrivirales* | *f.0296* | 2 |
| LC_20m_3um_k141_285138 | 2158 | Yes | Unknown | *Kitrinoviricota* | *c.0028* | *o.0039* | *f.0150* | 2 |
| LC_20m_3um_k141_298856 | 6686 | Yes | Unknown | *Kitrinoviricota* | *c.0034* | *o.0045* | *f.0176* | 5 |
| LC_20m_3um_k141_299800 | 6685 | Yes | Unknown | *Kitrinoviricota* | *c.0034* | *o.0045* | *f.0176* | 6 |
| LC_20m_3um_k141_300840 | 3614 | Yes | Unknown | *Kitrinoviricota* | *c.0034* | *o.0045* | *f.0176* | 4 |
| LC_20m_3um_k141_302602 | 621 | Yes | Unknown | *Kitrinoviricota* | *c.0034* | *o.0045* | *f.0176* | 1 |
| LC_20m_3um_k141_304944 | 1262 | No | NA | *NA* | *NA* | *NA* | *Nam-Co_family_61* | 1 |
| LC_20m_3um_k141_306266 | 640 | No | NA | *NA* | *NA* | *NA* | *NA* | 1 |
| LC_20m_3um_k141_328198 | 1120 | Yes | Unknown | *Duplornaviricota* | *Chrymotiviricetes* | *Ghabrivirales* | *f.0283.base-Megabirna* | 1 |
| LC_20m_3um_k141_340784 | 1307 | Yes | Unknown | *Duplornaviricota* | *Chrymotiviricetes* | *Ghabrivirales* | *f.0296* | 1 |
| LC_20m_3um_k141_343813 | 3011 | Yes | ssRNA(+) | *Lenarviricota* | *Howeltoviricetes* | *Cryppavirales* | *Mitoviridae* | 2 |
| LC_20m_3um_k141_352940 | 702 | Yes | ssRNA(+) | *Lenarviricota* | *Amabiliviricetes* | *Wolframvirales* | *Narnaviridae* | 1 |
| LC_20m_3um_k141_357943 | 7305 | Yes | Unknown | *Kitrinoviricota* | *c.0032* | *o.0043* | *f.0167* | 5 |
| LC_20m_3um_k141_366443 | 916 | Yes | Unknown | *Negarnaviricota* | *Ellioviricetes* | *Bunyavirales* | *f.0300.base-Phenui* | 1 |
| LC_20m_3um_k141_389751 | 8020 | Yes | Unknown | *Kitrinoviricota* | *c.0034* | *o.0045* | *f.0176* | 4 |
| LC_20m_3um_k141_408031 | 655 | Yes | Unknown | *Pisuviricota* | *Duplopiviricetes* | *Durnavirales* | *f.0111* | 1 |
| LC_20m_3um_k141_423756 | 2580 | Yes | ssRNA(+) | *Lenarviricota* | *Amabiliviricetes* | *Wolframvirales* | *Narnaviridae* | 1 |
| LC_20m_3um_k141_43900 | 749 | Yes | ssRNA(+) | *Kitrinoviricota* | *Alsuviricetes* | *Martellivirales* | *Virgaviridae* | 1 |
| LC_20m_3um_k141_442215 | 1833 | Yes | ssRNA(+) | *Lenarviricota* | *Amabiliviricetes* | *Wolframvirales* | *Narnaviridae* | 1 |
| LC_20m_3um_k141_445022 | 4206 | Yes | Unknown | *Duplornaviricota* | *Chrymotiviricetes* | *Ghabrivirales* | *f.0295* | 4 |
| LC_20m_3um_k141_45593 | 885 | Yes | Unknown | *Pisuviricota* | *Duplopiviricetes* | *Durnavirales* | *f.0111* | 1 |
| LC_20m_3um_k141_459176 | 2230 | Yes | Unknown | *Pisuviricota* | *Duplopiviricetes* | *Durnavirales* | *f.0102.base-Curvula* | 2 |
| LC_20m_3um_k141_460466 | 2004 | Yes | Unknown | *Kitrinoviricota* | *c.0034* | *o.0045* | *f.0176* | 3 |
| LC_20m_3um_k141_460575 | 3254 | Yes | ssRNA(+) | *Lenarviricota* | *Amabiliviricetes* | *Wolframvirales* | *Narnaviridae* | 1 |
| LC_20m_3um_k141_469319 | 1191 | Yes | ssRNA(+) | *Lenarviricota* | *Amabiliviricetes* | *Wolframvirales* | *Narnaviridae* | 1 |
| LC_20m_3um_k141_492567 | 656 | Yes | ssRNA(+) | *Lenarviricota* | *Howeltoviricetes* | *Cryppavirales* | *Mitoviridae* | 1 |
| LC_20m_3um_k141_492967 | 2690 | Yes | Unknown | *Pisuviricota* | *Duplopiviricetes* | *Durnavirales* | *f.0102.base-Curvula* | 3 |
| LC_20m_3um_k141_494413 | 1339 | Yes | ssRNA(+) | *Lenarviricota* | *Howeltoviricetes* | *Cryppavirales* | *Mitoviridae* | 1 |
| LC_20m_3um_k141_522525 | 6510 | Yes | Unknown | *Kitrinoviricota* | *c.0034* | *o.0045* | *f.0176* | 5 |
| LC_20m_3um_k141_526662 | 647 | Yes | Unknown | *Duplornaviricota* | *Chrymotiviricetes* | *Ghabrivirales* | *f.0296* | 1 |
| LC_20m_3um_k141_531564 | 938 | Yes | Unknown | *Duplornaviricota* | *Chrymotiviricetes* | *Ghabrivirales* | *f.0285* | 1 |
| LC_20m_3um_k141_538898 | 1700 | Yes | Unknown | *Duplornaviricota* | *Chrymotiviricetes* | *Ghabrivirales* | *f.0287* | 3 |
| LC_20m_3um_k141_54880 | 1235 | No | NA | *NA* | *NA* | *NA* | *NA* | 1 |
| LC_20m_3um_k141_551912 | 1085 | Yes | Unknown | *Pisuviricota* | *Duplopiviricetes* | *Durnavirales* | *f.0102.base-Curvula* | 1 |
| LC_20m_3um_k141_555582 | 1753 | Yes | dsRNA | *Pisuviricota* | *Duplopiviricetes* | *Durnavirales* | *Picobirnaviridae* | 1 |
| LC_20m_3um_k141_585136 | 4375 | Yes | Unknown | *Kitrinoviricota* | *c.0034* | *o.0045* | *f.0176* | 4 |
| LC_20m_3um_k141_587133 | 8042 | Yes | Unknown | *Kitrinoviricota* | *c.0034* | *o.0045* | *f.0176* | 6 |
| LC_20m_3um_k141_93254 | 8464 | Yes | Unknown | *Kitrinoviricota* | *c.0034* | *o.0045* | *f.0176* | 6 |
| LC_50m_0.2um_k141_113601 | 5115 | Yes | Unknown | *Negarnaviricota* | *Ellioviricetes* | *Bunyavirales* | *f.0305.base-Leishbu* | 1 |
| LC_50m_0.2um_k141_137129 | 2510 | Yes | Unknown | *Pisuviricota* | *c.0013* | *o.0025.base-Sobeli* | *f.0076* | 1 |
| LC_50m_0.2um_k141_144468 | 3456 | Yes | ssRNA(+) | *Lenarviricota* | *Amabiliviricetes* | *Wolframvirales* | *Narnaviridae* | 2 |
| LC_50m_0.2um_k141_182361 | 8432 | Yes | Unknown | *Kitrinoviricota* | *c.0034* | *o.0045* | *f.0176* | 5 |
| LC_50m_0.2um_k141_187776 | 6973 | Yes | Unknown | *Kitrinoviricota* | *c.0034* | *o.0045* | *f.0176* | 6 |
| LC_50m_0.2um_k141_198770 | 5878 | Yes | Unknown | *Pisuviricota* | *c.0003.base-Pisoni* | *o.0003* | *Nam-Co_family_27* | 2 |
| LC_50m_0.2um_k141_22609 | 2366 | Yes | Unknown | *Pisuviricota* | *c.0013* | *o.0025.base-Sobeli* | *f.0076* | 1 |
| LC_50m_0.2um_k141_26077 | 916 | Yes | Unknown | *Kitrinoviricota* | *c.0034* | *o.0045* | *f.0176* | 1 |
| LC_50m_0.2um_k141_292980 | 8485 | Yes | Unknown | *Kitrinoviricota* | *c.0034* | *o.0045* | *f.0176* | 7 |
| LC_50m_0.2um_k141_293183 | 842 | Yes | Unknown | *Pisuviricota* | *c.0013* | *o.0025.base-Sobeli* | *f.0076* | 1 |
| LC_50m_0.2um_k141_318047 | 7923 | Yes | Unknown | *Kitrinoviricota* | *c.0034* | *o.0045* | *f.0176* | 6 |
| LC_50m_0.2um_k141_349625 | 4469 | Yes | Unknown | *Kitrinoviricota* | *c.0034* | *o.0045* | *f.0176* | 2 |
| LC_50m_0.2um_k141_35592 | 1680 | Yes | Unknown | *Kitrinoviricota* | *c.0034* | *o.0045* | *f.0176* | 1 |
| LC_50m_0.2um_k141_370905 | 5053 | Yes | Unknown | *Kitrinoviricota* | *c.0025* | *o.0036* | *f.0145* | 2 |
| LC_50m_0.2um_k141_373844 | 3546 | Yes | ssRNA(+) | *Lenarviricota* | *Amabiliviricetes* | *Wolframvirales* | *Narnaviridae* | 3 |
| LC_50m_0.2um_k141_412717 | 4524 | Yes | Unknown | *Kitrinoviricota* | *c.0031* | *o.0042* | *f.0153* | 1 |
| LC_50m_0.2um_k141_442366 | 7048 | Yes | Unknown | *Kitrinoviricota* | *c.0034* | *o.0045* | *f.0176* | 5 |
| LC_50m_0.2um_k141_447527 | 2739 | Yes | ssRNA(+) | *Lenarviricota* | *Amabiliviricetes* | *Wolframvirales* | *Narnaviridae* | 1 |
| LC_50m_0.2um_k141_489354 | 960 | Yes | Unknown | *Duplornaviricota* | *Chrymotiviricetes* | *Ghabrivirales* | *f.0296* | 1 |
| LC_50m_0.2um_k141_4949 | 1546 | Yes | Unknown | *Lenarviricota* | *c.0069.base-Howelto* | *o.0090* | *f.0386* | 1 |
| LC_50m_0.2um_k141_58378 | 8223 | Yes | Unknown | *Kitrinoviricota* | *c.0034* | *o.0045* | *f.0176* | 4 |
| LC_50m_0.2um_k141_65382 | 2014 | Yes | Unknown | *Pisuviricota* | *Duplopiviricetes* | *Durnavirales* | *f.0111* | 1 |
| LC_50m_0.2um_k141_68686 | 2259 | Yes | Unknown | *Lenarviricota* | *c.0071.base-Howelto* | *o.0092* | *f.0388* | 3 |
| LC_50m_0.2um_k141_80321 | 4940 | Yes | Unknown | *Kitrinoviricota* | *c.0034* | *o.0045* | *f.0176* | 4 |
| LC_50m_0.2um_k141_84230 | 6634 | Yes | Unknown | *Kitrinoviricota* | *c.0034* | *o.0045* | *f.0176* | 3 |
| LC_50m_1.2um_k141_133709 | 2710 | Yes | Unknown | *Kitrinoviricota* | *c.0034* | *o.0045* | *f.0176* | 2 |
| LC_50m_1.2um_k141_14943 | 3583 | Yes | Unknown | *Kitrinoviricota* | *c.0034* | *o.0045* | *f.0176* | 2 |
| LC_50m_1.2um_k141_154296 | 2336 | Yes | Unknown | *Kitrinoviricota* | *c.0034* | *o.0045* | *f.0176* | 2 |
| LC_50m_1.2um_k141_160657 | 1015 | Yes | Unknown | *Kitrinoviricota* | *c.0034* | *o.0045* | *f.0176* | 1 |
| LC_50m_1.2um_k141_161152 | 5236 | Yes | dsRNA | *Duplornaviricota* | *Chrymotiviricetes* | *Ghabrivirales* | *Totiviridae* | 2 |
| LC_50m_1.2um_k141_194815 | 3427 | Yes | Unknown | *Kitrinoviricota* | *c.0034* | *o.0045* | *f.0176* | 3 |
| LC_50m_1.2um_k141_21225 | 7635 | Yes | Unknown | *Kitrinoviricota* | *c.0034* | *o.0045* | *f.0176* | 6 |
| LC_50m_1.2um_k141_213095 | 7983 | Yes | Unknown | *Kitrinoviricota* | *c.0034* | *o.0045* | *f.0176* | 5 |
| LC_50m_1.2um_k141_22302 | 4860 | Yes | Unknown | *Kitrinoviricota* | *c.0034* | *o.0045* | *f.0176* | 4 |
| LC_50m_1.2um_k141_235637 | 1968 | Yes | Unknown | *Kitrinoviricota* | *c.0034* | *o.0045* | *f.0176* | 2 |
| LC_50m_1.2um_k141_238221 | 2713 | Yes | Unknown | *Lenarviricota* | *c.0071.base-Howelto* | *o.0092* | *f.0388* | 2 |
| LC_50m_1.2um_k141_257994 | 5673 | Yes | Unknown | *Duplornaviricota* | *Chrymotiviricetes* | *Ghabrivirales* | *f.0284.base-Megabirna* | 2 |
| LC_50m_1.2um_k141_276503 | 4270 | Yes | dsRNA | *Duplornaviricota* | *Resentoviricetes* | *Reovirales* | *Reoviridae* | 1 |
| LC_50m_1.2um_k141_297650 | 9763 | Yes | Unknown | *Duplornaviricota* | *Chrymotiviricetes* | *Ghabrivirales* | *f.0292* | 3 |
| LC_50m_1.2um_k141_297662 | 828 | Yes | Unknown | *Pisuviricota* | *Duplopiviricetes* | *Durnavirales* | *f.0111* | 1 |
| LC_50m_1.2um_k141_298547 | 7299 | Yes | Unknown | *Kitrinoviricota* | *c.0034* | *o.0045* | *f.0176* | 4 |
| LC_50m_1.2um_k141_307887 | 2166 | Yes | ssRNA(+) | *Lenarviricota* | *Howeltoviricetes* | *Cryppavirales* | *Mitoviridae* | 3 |
| LC_50m_1.2um_k141_329405 | 7698 | Yes | Unknown | *Negarnaviricota* | *Ellioviricetes* | *Bunyavirales* | *f.0328* | 2 |
| LC_50m_1.2um_k141_394034 | 3671 | Yes | ssRNA(+) | *Lenarviricota* | *Amabiliviricetes* | *Wolframvirales* | *Narnaviridae* | 4 |
| LC_50m_1.2um_k141_61970 | 6608 | Yes | Unknown | *Kitrinoviricota* | *c.0031* | *o.0042* | *f.0153* | 1 |
| LC_50m_1.2um_k141_78854 | 8461 | Yes | Unknown | *Kitrinoviricota* | *c.0034* | *o.0045* | *f.0176* | 6 |
| LC_50m_10um_k141_103028 | 1235 | Yes | Unknown | *Pisuviricota* | *Duplopiviricetes* | *Durnavirales* | *f.0111* | 1 |
| LC_50m_10um_k141_105181 | 646 | Yes | Unknown | *Kitrinoviricota* | *c.0034* | *o.0045* | *f.0176* | 1 |
| LC_50m_10um_k141_106799 | 1217 | Yes | Unknown | *Pisuviricota* | *Duplopiviricetes* | *Durnavirales* | *f.0111* | 1 |
| LC_50m_10um_k141_107693 | 2027 | Yes | Unknown | *Pisuviricota* | *Duplopiviricetes* | *Durnavirales* | *f.0102.base-Curvula* | 2 |
| LC_50m_10um_k141_114455 | 4475 | Yes | ssRNA(+) | *Kitrinoviricota* | *Alsuviricetes* | *o.0055* | *Endornaviridae* | 1 |
| LC_50m_10um_k141_114996 | 6306 | Yes | Unknown | *Kitrinoviricota* | *c.0034* | *o.0045* | *f.0176* | 6 |
| LC_50m_10um_k141_123519 | 2461 | Yes | Unknown | *Duplornaviricota* | *Chrymotiviricetes* | *Ghabrivirales* | *f.0283.base-Megabirna* | 1 |
| LC_50m_10um_k141_136548 | 1459 | Yes | Unknown | *Pisuviricota* | *Duplopiviricetes* | *Durnavirales* | *f.0102.base-Curvula* | 2 |
| LC_50m_10um_k141_142930 | 1339 | Yes | Unknown | *Pisuviricota* | *Duplopiviricetes* | *Durnavirales* | *f.0102.base-Curvula* | 2 |
| LC_50m_10um_k141_147952 | 2015 | Yes | Unknown | *Pisuviricota* | *Duplopiviricetes* | *Durnavirales* | *f.0111* | 2 |
| LC_50m_10um_k141_154418 | 2657 | Yes | ssRNA(+) | *Lenarviricota* | *Amabiliviricetes* | *Wolframvirales* | *Nam-Co_family_07* | 1 |
| LC_50m_10um_k141_162379 | 1488 | Yes | Unknown | *Duplornaviricota* | *Chrymotiviricetes* | *Ghabrivirales* | *f.0282.base-Megabirna* | 1 |
| LC_50m_10um_k141_180346 | 903 | Yes | Unknown | *Kitrinoviricota* | *c.0034* | *o.0045* | *f.0176* | 1 |
| LC_50m_10um_k141_212090 | 3302 | Yes | Unknown | *Pisuviricota* | *Duplopiviricetes* | *Durnavirales* | *f.0102.base-Curvula* | 2 |
| LC_50m_10um_k141_216893 | 2945 | Yes | Unknown | *Duplornaviricota* | *Chrymotiviricetes* | *Ghabrivirales* | *f.0296* | 2 |
| LC_50m_10um_k141_221720 | 1375 | Yes | ssRNA(+) | *Lenarviricota* | *Howeltoviricetes* | *Cryppavirales* | *Mitoviridae* | 1 |
| LC_50m_10um_k141_22825 | 3467 | Yes | Unknown | *Kitrinoviricota* | *c.0034* | *o.0045* | *f.0176* | 4 |
| LC_50m_10um_k141_242393 | 1757 | Yes | Unknown | *Pisuviricota* | *Duplopiviricetes* | *Durnavirales* | *f.0111* | 1 |
| LC_50m_10um_k141_244857 | 1498 | Yes | Unknown | *Pisuviricota* | *Duplopiviricetes* | *Durnavirales* | *f.0111* | 1 |
| LC_50m_10um_k141_252423 | 3854 | Yes | ssRNA(+) | *Lenarviricota* | *Amabiliviricetes* | *Wolframvirales* | *Narnaviridae* | 2 |
| LC_50m_10um_k141_26560 | 3366 | Yes | ssRNA(+) | *Lenarviricota* | *Howeltoviricetes* | *Cryppavirales* | *Mitoviridae* | 1 |
| LC_50m_10um_k141_274640 | 1818 | Yes | ssRNA(+) | *Lenarviricota* | *Howeltoviricetes* | *Cryppavirales* | *Mitoviridae* | 2 |
| LC_50m_10um_k141_279227 | 1048 | Yes | Unknown | *Kitrinoviricota* | *c.0028* | *o.0039* | *f.0150* | 2 |
| LC_50m_10um_k141_280124 | 2400 | Yes | ssRNA(+) | *Lenarviricota* | *Amabiliviricetes* | *Wolframvirales* | *Narnaviridae* | 1 |
| LC_50m_10um_k141_292869 | 1053 | Yes | ssRNA(+) | *Lenarviricota* | *Howeltoviricetes* | *Cryppavirales* | *Mitoviridae* | 1 |
| LC_50m_10um_k141_295282 | 1516 | Yes | ssRNA(+) | *Lenarviricota* | *Amabiliviricetes* | *Wolframvirales* | *Narnaviridae* | 1 |
| LC_50m_10um_k141_298364 | 1150 | Yes | Unknown | *Pisuviricota* | *Duplopiviricetes* | *Durnavirales* | *f.0111* | 1 |
| LC_50m_10um_k141_303294 | 1655 | Yes | Unknown | *Pisuviricota* | *Duplopiviricetes* | *Durnavirales* | *f.0102.base-Curvula* | 2 |
| LC_50m_10um_k141_304517 | 1008 | Yes | ssRNA(+) | *Lenarviricota* | *Amabiliviricetes* | *Wolframvirales* | *Narnaviridae* | 1 |
| LC_50m_10um_k141_308889 | 1220 | Yes | ssRNA(+) | *Lenarviricota* | *Howeltoviricetes* | *Cryppavirales* | *Mitoviridae* | 2 |
| LC_50m_10um_k141_309854 | 1318 | Yes | Unknown | *Pisuviricota* | *Duplopiviricetes* | *Durnavirales* | *f.0111* | 1 |
| LC_50m_10um_k141_313706 | 1358 | Yes | ssRNA(+) | *Lenarviricota* | *Howeltoviricetes* | *Cryppavirales* | *Mitoviridae* | 1 |
| LC_50m_10um_k141_48043 | 1759 | Yes | Unknown | *Pisuviricota* | *Duplopiviricetes* | *Durnavirales* | *f.0111* | 1 |
| LC_50m_10um_k141_55109 | 2184 | Yes | ssRNA(+) | *Lenarviricota* | *Howeltoviricetes* | *Cryppavirales* | *Mitoviridae* | 1 |
| LC_50m_10um_k141_63841 | 945 | Yes | ssRNA(+) | *Lenarviricota* | *Howeltoviricetes* | *Cryppavirales* | *Mitoviridae* | 1 |
| LC_50m_10um_k141_65562 | 1498 | Yes | Unknown | *Duplornaviricota* | *Chrymotiviricetes* | *Ghabrivirales* | *f.0296* | 1 |
| LC_50m_10um_k141_68708 | 724 | Yes | ssRNA(+) | *Kitrinoviricota* | *Alsuviricetes* | *o.0055* | *Endornaviridae* | 1 |
| LC_50m_10um_k141_83167 | 1168 | Yes | Unknown | *Kitrinoviricota* | *c.0034* | *o.0045* | *f.0176* | 1 |
| LC_50m_10um_k141_96448 | 908 | Yes | ssRNA(+) | *Lenarviricota* | *Miaviricetes* | *Ourlivirales* | *Botourmiaviridae* | 1 |
| LC_50m_10um_k141_99806 | 768 | Yes | Unknown | *Duplornaviricota* | *Chrymotiviricetes* | *Ghabrivirales* | *f.0287* | 1 |
| LC_50m_3um_k141_109718 | 1112 | Yes | Unknown | *Pisuviricota* | *Duplopiviricetes* | *Durnavirales* | *Nam-Co_family_39* | 2 |
| LC_50m_3um_k141_114574 | 1442 | Yes | ssRNA(+) | *Lenarviricota* | *Howeltoviricetes* | *Cryppavirales* | *Mitoviridae* | 2 |
| LC_50m_3um_k141_120753 | 2690 | Yes | Unknown | *Pisuviricota* | *Duplopiviricetes* | *Durnavirales* | *f.0102.base-Curvula* | 3 |
| LC_50m_3um_k141_133114 | 9769 | Yes | ssRNA(+) | *Kitrinoviricota* | *Alsuviricetes* | *Hepelivirales* | *Benyviridae* | 4 |
| LC_50m_3um_k141_139834 | 657 | Yes | ssRNA(+) | *Lenarviricota* | *Amabiliviricetes* | *Wolframvirales* | *Narnaviridae* | 1 |
| LC_50m_3um_k141_145688 | 876 | Yes | ssRNA(+) | *Lenarviricota* | *Howeltoviricetes* | *Cryppavirales* | *Mitoviridae* | 1 |
| LC_50m_3um_k141_156190 | 6543 | Yes | Unknown | *Kitrinoviricota* | *c.0034* | *o.0045* | *f.0176* | 5 |
| LC_50m_3um_k141_189961 | 7050 | Yes | Unknown | *Kitrinoviricota* | *c.0034* | *o.0045* | *f.0176* | 6 |
| LC_50m_3um_k141_208177 | 3646 | Yes | Unknown | *Duplornaviricota* | *Chrymotiviricetes* | *Ghabrivirales* | *f.0282.base-Megabirna* | 2 |
| LC_50m_3um_k141_210126 | 2176 | Yes | dsRNA | *Pisuviricota* | *Duplopiviricetes* | *Durnavirales* | *Partitiviridae* | 2 |
| LC_50m_3um_k141_234150 | 2122 | Yes | Unknown | *Pisuviricota* | *Duplopiviricetes* | *Durnavirales* | *f.0111* | 1 |
| LC_50m_3um_k141_238018 | 6523 | Yes | Unknown | *Kitrinoviricota* | *c.0034* | *o.0045* | *f.0176* | 5 |
| LC_50m_3um_k141_239166 | 7069 | Yes | Unknown | *Kitrinoviricota* | *c.0034* | *o.0045* | *f.0176* | 4 |
| LC_50m_3um_k141_239751 | 2330 | Yes | ssRNA(+) | *Lenarviricota* | *Howeltoviricetes* | *Cryppavirales* | *Mitoviridae* | 1 |
| LC_50m_3um_k141_256245 | 2532 | Yes | Unknown | *Kitrinoviricota* | *c.0034* | *o.0045* | *f.0176* | 2 |
| LC_50m_3um_k141_273654 | 761 | Yes | Unknown | *Pisuviricota* | *Duplopiviricetes* | *Durnavirales* | *f.0102.base-Curvula* | 1 |
| LC_50m_3um_k141_295552 | 1767 | Yes | Unknown | *Pisuviricota* | *Duplopiviricetes* | *Durnavirales* | *f.0111* | 1 |
| LC_50m_3um_k141_325294 | 3425 | Yes | Unknown | *Duplornaviricota* | *Chrymotiviricetes* | *Ghabrivirales* | *f.0296* | 3 |
| LC_50m_3um_k141_32763 | 1605 | Yes | Unknown | *Pisuviricota* | *Duplopiviricetes* | *Durnavirales* | *f.0111* | 2 |
| LC_50m_3um_k141_335639 | 1598 | Yes | Unknown | *Kitrinoviricota* | *c.0034* | *o.0045* | *f.0176* | 2 |
| LC_50m_3um_k141_385349 | 6282 | Yes | Unknown | *Pisuviricota* | *Duplopiviricetes* | *Durnavirales* | *f.0102.base-Curvula* | 4 |
| LC_50m_3um_k141_426641 | 1149 | Yes | dsRNA | *Pisuviricota* | *Duplopiviricetes* | *Durnavirales* | *Partitiviridae* | 1 |
| LC_50m_3um_k141_431241 | 3866 | Yes | Unknown | *Kitrinoviricota* | *c.0034* | *o.0045* | *f.0176* | 2 |
| LC_50m_3um_k141_450784 | 1533 | Yes | Unknown | *Pisuviricota* | *Duplopiviricetes* | *Durnavirales* | *f.0111* | 1 |
| LC_50m_3um_k141_459293 | 2649 | Yes | ssRNA(+) | *Lenarviricota* | *Amabiliviricetes* | *Wolframvirales* | *Narnaviridae* | 2 |
| LC_50m_3um_k141_468226 | 7203 | Yes | Unknown | *Kitrinoviricota* | *c.0034* | *o.0045* | *f.0176* | 6 |
| LC_50m_3um_k141_469426 | 3397 | Yes | Unknown | *Kitrinoviricota* | *c.0034* | *o.0045* | *f.0176* | 3 |
| LC_50m_3um_k141_494636 | 6270 | Yes | Unknown | *Negarnaviricota* | *Ellioviricetes* | *Bunyavirales* | *f.0313* | 1 |
| LC_50m_3um_k141_506229 | 1198 | Yes | Unknown | *Pisuviricota* | *Duplopiviricetes* | *Durnavirales* | *f.0111* | 1 |
| LC_50m_3um_k141_506636 | 6345 | Yes | Unknown | *Duplornaviricota* | *Chrymotiviricetes* | *Ghabrivirales* | *f.0296* | 2 |
| LC_50m_3um_k141_511998 | 1991 | Yes | Unknown | *Duplornaviricota* | *Chrymotiviricetes* | *Ghabrivirales* | *f.0296* | 2 |
| LC_50m_3um_k141_545291 | 988 | Yes | ssRNA(+) | *Lenarviricota* | *Amabiliviricetes* | *Wolframvirales* | *Narnaviridae* | 1 |
| LC_50m_3um_k141_75755 | 775 | Yes | Unknown | *Kitrinoviricota* | *c.0025* | *o.0036* | *f.0145* | 1 |
| LC_50m_3um_k141_76750 | 973 | Yes | Unknown | *Kitrinoviricota* | *c.0034* | *o.0045* | *f.0176* | 1 |
| LC_80m_0.2um_k141_102393 | 5352 | Yes | Unknown | *Kitrinoviricota* | *c.0034* | *o.0045* | *f.0176* | 4 |
| LC_80m_0.2um_k141_108571 | 7348 | Yes | Unknown | *Negarnaviricota* | *Ellioviricetes* | *Bunyavirales* | *f.0315* | 2 |
| LC_80m_0.2um_k141_138777 | 2474 | Yes | Unknown | *Pisuviricota* | *c.0013* | *o.0025.base-Sobeli* | *f.0076* | 1 |
| LC_80m_0.2um_k141_151559 | 658 | Yes | Unknown | *Kitrinoviricota* | *c.0034* | *o.0045* | *f.0176* | 1 |
| LC_80m_0.2um_k141_159826 | 2472 | Yes | dsRNA | *Pisuviricota* | *Duplopiviricetes* | *Durnavirales* | *Amalgaviridae* | 2 |
| LC_80m_0.2um_k141_185079 | 2623 | Yes | ssRNA(+) | *Lenarviricota* | *Amabiliviricetes* | *Wolframvirales* | *Narnaviridae* | 2 |
| LC_80m_0.2um_k141_237974 | 663 | Yes | ssRNA(+) | *Lenarviricota* | *Miaviricetes* | *Ourlivirales* | *Botourmiaviridae* | 1 |
| LC_80m_0.2um_k141_245920 | 858 | Yes | Unknown | *Pisuviricota* | *c.0013* | *o.0025.base-Sobeli* | *f.0076* | 1 |
| LC_80m_0.2um_k141_301895 | 774 | Yes | Unknown | *Pisuviricota* | *c.0013* | *o.0025.base-Sobeli* | *f.0076* | 1 |
| LC_80m_0.2um_k141_329916 | 2482 | Yes | ssRNA(+) | *Lenarviricota* | *Amabiliviricetes* | *Wolframvirales* | *Narnaviridae* | 2 |
| LC_80m_0.2um_k141_34599 | 1525 | Yes | Unknown | *Pisuviricota* | *c.0013* | *o.0025.base-Sobeli* | *f.0076* | 2 |
| LC_80m_0.2um_k141_354981 | 660 | Yes | Unknown | *Pisuviricota* | *c.0013* | *o.0025.base-Sobeli* | *f.0076* | 1 |
| LC_80m_0.2um_k141_367785 | 888 | Yes | ssRNA(+) | *Kitrinoviricota* | *Tolucaviricetes* | *Tolivirales* | *Tombusviridae* | 1 |
| LC_80m_0.2um_k141_385333 | 4133 | Yes | Unknown | *Kitrinoviricota* | *c.0034* | *o.0045* | *f.0176* | 2 |
| LC_80m_0.2um_k141_400659 | 726 | Yes | ssRNA(+) | *Lenarviricota* | *Leviviricetes* | *Norzivirales* | *Atkinsviridae* | 1 |
| LC_80m_0.2um_k141_447705 | 1849 | Yes | Unknown | *Kitrinoviricota* | *c.0034* | *o.0045* | *f.0176* | 2 |
| LC_80m_0.2um_k141_456306 | 8220 | Yes | Unknown | *Kitrinoviricota* | *c.0034* | *o.0045* | *f.0176* | 7 |
| LC_80m_0.2um_k141_512934 | 2005 | Yes | Unknown | *Kitrinoviricota* | *c.0031* | *o.0042* | *f.0153* | 1 |
| LC_80m_0.2um_k141_514539 | 626 | Yes | Unknown | *Duplornaviricota* | *Chrymotiviricetes* | *Ghabrivirales* | *f.0282.base-Megabirna* | 1 |
| LC_80m_0.2um_k141_542511 | 1617 | No | NA | *NA* | *NA* | *NA* | *Nam-Co_family_48* | 1 |
| LC_80m_0.2um_k141_547677 | 3381 | Yes | Unknown | *Negarnaviricota* | *Ellioviricetes* | *Bunyavirales* | *f.0313* | 1 |
| LC_80m_0.2um_k141_548164 | 1155 | Yes | Unknown | *Kitrinoviricota* | *c.0032* | *o.0043* | *f.0167* | 1 |
| LC_80m_0.2um_k141_558302 | 2889 | Yes | Unknown | *Duplornaviricota* | *Chrymotiviricetes* | *Ghabrivirales* | *f.0296* | 2 |
| LC_80m_0.2um_k141_57395 | 4466 | Yes | Unknown | *Kitrinoviricota* | *c.0031* | *o.0042* | *f.0153* | 1 |
| LC_80m_0.2um_k141_597934 | 1240 | Yes | Unknown | *Pisuviricota* | *Duplopiviricetes* | *Durnavirales* | *f.0120* | 1 |
| LC_80m_0.2um_k141_601816 | 4176 | Yes | Unknown | *Negarnaviricota* | *Ellioviricetes* | *Bunyavirales* | *f.0315* | 2 |
| LC_80m_0.2um_k141_605149 | 1560 | Yes | Unknown | *Pisuviricota* | *c.0013* | *o.0025.base-Sobeli* | *f.0076* | 1 |
| LC_80m_0.2um_k141_63445 | 789 | Yes | Unknown | *Kitrinoviricota* | *c.0031* | *o.0042* | *f.0153* | 1 |
| LC_80m_0.2um_k141_81878 | 7422 | Yes | Unknown | *Kitrinoviricota* | *c.0034* | *o.0045* | *f.0176* | 4 |
| LC_80m_0.2um_k141_84894 | 2754 | Yes | Unknown | *Pisuviricota* | *Duplopiviricetes* | *Durnavirales* | *f.0102.base-Curvula* | 2 |
| LC_80m_0.2um_k141_99178 | 1822 | Yes | Unknown | *Kitrinoviricota* | *c.0034* | *o.0045* | *f.0176* | 1 |
| LC_80m_1.2um_k141_125684 | 1348 | Yes | Unknown | *Kitrinoviricota* | *c.0031* | *o.0042* | *f.0153* | 1 |
| LC_80m_1.2um_k141_147906 | 1624 | Yes | Unknown | *Pisuviricota* | *Pisoniviricetes* | *Picornavirales* | *f.0044.base-Marna* | 1 |
| LC_80m_1.2um_k141_151565 | 3060 | Yes | ssRNA(+) | *Lenarviricota* | *Amabiliviricetes* | *Wolframvirales* | *Narnaviridae* | 1 |
| LC_80m_1.2um_k141_207456 | 1144 | No | NA | *NA* | *NA* | *NA* | *NA* | 1 |
| LC_80m_1.2um_k141_226493 | 8709 | Yes | Unknown | *Kitrinoviricota* | *c.0034* | *o.0045* | *f.0176* | 7 |
| LC_80m_1.2um_k141_231116 | 2602 | Yes | ssRNA(+) | *Lenarviricota* | *Howeltoviricetes* | *Cryppavirales* | *Mitoviridae* | 3 |
| LC_80m_1.2um_k141_267137 | 2695 | Yes | ssRNA(+) | *Lenarviricota* | *Howeltoviricetes* | *Cryppavirales* | *Mitoviridae* | 1 |
| LC_80m_1.2um_k141_311721 | 4283 | No | NA | *NA* | *NA* | *NA* | *NA* | 3 |
| LC_80m_1.2um_k141_344716 | 3490 | Yes | Unknown | *Kitrinoviricota* | *c.0034* | *o.0045* | *f.0176* | 3 |
| LC_80m_1.2um_k141_403562 | 8545 | Yes | Unknown | *Kitrinoviricota* | *c.0034* | *o.0045* | *f.0176* | 6 |
| LC_80m_1.2um_k141_412250 | 1180 | Yes | ssRNA(+) | *Pisuviricota* | *c.0012* | *Sobelivirales* | *Solemoviridae* | 2 |
| LC_80m_1.2um_k141_431363 | 737 | Yes | Unknown | *Kitrinoviricota* | *c.0031* | *o.0042* | *f.0156* | 1 |
| LC_80m_1.2um_k141_435798 | 1930 | Yes | ssRNA(+) | *Lenarviricota* | *Amabiliviricetes* | *Wolframvirales* | *Narnaviridae* | 2 |
| LC_80m_1.2um_k141_94366 | 1148 | Yes | Unknown | *Kitrinoviricota* | *c.0034* | *o.0045* | *f.0176* | 1 |
| LC_80m_10um_k141_110104 | 708 | Yes | ssRNA(+) | *Lenarviricota* | *Howeltoviricetes* | *Cryppavirales* | *Mitoviridae* | 1 |
| LC_80m_10um_k141_122255 | 1199 | Yes | Unknown | *Kitrinoviricota* | *c.0034* | *o.0045* | *f.0176* | 1 |
| LC_80m_10um_k141_12242 | 5155 | Yes | Unknown | *Pisuviricota* | *c.0003.base-Pisoni* | *o.0003* | *f.0056* | 1 |
| LC_80m_10um_k141_15366 | 1690 | Yes | Unknown | *Pisuviricota* | *Duplopiviricetes* | *Durnavirales* | *f.0111* | 1 |
| LC_80m_10um_k141_158158 | 4735 | No | NA | *NA* | *NA* | *NA* | *NA* | 2 |
| LC_80m_10um_k141_161908 | 2279 | Yes | ssRNA(+) | *Lenarviricota* | *Howeltoviricetes* | *Cryppavirales* | *Mitoviridae* | 1 |
| LC_80m_10um_k141_171565 | 916 | Yes | Unknown | *Kitrinoviricota* | *c.0031* | *o.0042* | *f.0153* | 1 |
| LC_80m_10um_k141_183687 | 1940 | Yes | ssRNA(+) | *Lenarviricota* | *Amabiliviricetes* | *Wolframvirales* | *Narnaviridae* | 1 |
| LC_80m_10um_k141_188622 | 2701 | Yes | Unknown | *Pisuviricota* | *Pisoniviricetes* | *Picornavirales* | *f.0025* | 1 |
| LC_80m_10um_k141_20695 | 14241 | Yes | ssRNA(+) | *Kitrinoviricota* | *Alsuviricetes* | *o.0055* | *Endornaviridae* | 2 |
| LC_80m_10um_k141_207174 | 8193 | Yes | Unknown | *Negarnaviricota* | *Ellioviricetes* | *Bunyavirales* | *f.0315* | 1 |
| LC_80m_10um_k141_265723 | 2004 | Yes | dsRNA | *Duplornaviricota* | *Chrymotiviricetes* | *Ghabrivirales* | *Totiviridae* | 1 |
| LC_80m_10um_k141_267292 | 3428 | Yes | Unknown | *Pisuviricota* | *Duplopiviricetes* | *Durnavirales* | *f.0102.base-Curvula* | 3 |
| LC_80m_10um_k141_289478 | 1512 | Yes | Unknown | *Pisuviricota* | *Pisoniviricetes* | *Picornavirales* | *f.0025* | 1 |
| LC_80m_10um_k141_36309 | 2356 | Yes | Unknown | *Duplornaviricota* | *Chrymotiviricetes* | *Ghabrivirales* | *f.0282.base-Megabirna* | 1 |
| LC_80m_10um_k141_74353 | 3183 | Yes | ssRNA(+) | *Lenarviricota* | *Amabiliviricetes* | *Wolframvirales* | *Narnaviridae* | 1 |
| LC_80m_10um_k141_93320 | 2181 | Yes | ssRNA(+) | *Lenarviricota* | *Amabiliviricetes* | *Wolframvirales* | *Narnaviridae* | 1 |
| LC_80m_3um_k141_104397 | 817 | No | NA | *NA* | *NA* | *NA* | *NA* | 1 |
| LC_80m_3um_k141_124331 | 912 | Yes | Unknown | *Pisuviricota* | *c.0013* | *o.0025.base-Sobeli* | *f.0076* | 1 |
| LC_80m_3um_k141_127394 | 1495 | Yes | Unknown | *Kitrinoviricota* | *c.0034* | *o.0045* | *f.0176* | 2 |
| LC_80m_3um_k141_145709 | 2885 | Yes | ssRNA(+) | *Lenarviricota* | *Amabiliviricetes* | *Wolframvirales* | *Narnaviridae* | 1 |
| LC_80m_3um_k141_150182 | 1437 | No | NA | *NA* | *NA* | *NA* | *NA* | 1 |
| LC_80m_3um_k141_157262 | 3499 | Yes | ssRNA(+) | *Lenarviricota* | *Amabiliviricetes* | *Wolframvirales* | *Narnaviridae* | 2 |
| LC_80m_3um_k141_184180 | 946 | Yes | Unknown | *Pisuviricota* | *Duplopiviricetes* | *Durnavirales* | *f.0111* | 1 |
| LC_80m_3um_k141_192746 | 1780 | Yes | Unknown | *Kitrinoviricota* | *c.0034* | *o.0045* | *f.0176* | 2 |
| LC_80m_3um_k141_217658 | 1153 | Yes | Unknown | *Lenarviricota* | *c.0074.base-Howelto* | *o.0095* | *f.0391* | 1 |
| LC_80m_3um_k141_232182 | 4508 | Yes | Unknown | *Kitrinoviricota* | *c.0034* | *o.0045* | *f.0176* | 2 |
| LC_80m_3um_k141_257744 | 2814 | Yes | Unknown | *Duplornaviricota* | *Chrymotiviricetes* | *Ghabrivirales* | *f.0282.base-Megabirna* | 1 |
| LC_80m_3um_k141_40456 | 829 | Yes | ssRNA(+) | *Pisuviricota* | *Pisoniviricetes* | *Picornavirales* | *Marnaviridae* | 1 |
| LC_80m_3um_k141_41719 | 979 | Yes | Unknown | *Pisuviricota* | *Duplopiviricetes* | *Durnavirales* | *f.0102.base-Curvula* | 1 |
| LC_80m_3um_k141_59616 | 3014 | No | NA | *NA* | *NA* | *NA* | *NA* | 1 |
| LC_80m_3um_k141_63205 | 6137 | Yes | Unknown | *Kitrinoviricota* | *c.0034* | *o.0045* | *f.0176* | 5 |
| LC_80m_3um_k141_96140 | 3952 | Yes | Unknown | *Kitrinoviricota* | *c.0034* | *o.0045* | *f.0176* | 4 |
| LC_80m_3um_k141_96969 | 5444 | Yes | Unknown | *Kitrinoviricota* | *c.0034* | *o.0045* | *f.0176* | 4 |
| LC_coassembly_k141_1001983 | 2638 | Yes | ssRNA(+) | *Lenarviricota* | *Amabiliviricetes* | *Wolframvirales* | *Narnaviridae* | 1 |
| LC_coassembly_k141_1002275 | 1538 | Yes | Unknown | *Lenarviricota* | *c.0071.base-Howelto* | *o.0092* | *f.0388* | 1 |
| LC_coassembly_k141_101176 | 6638 | Yes | Unknown | *Pisuviricota* | *Duplopiviricetes* | *Durnavirales* | *f.0102.base-Curvula* | 5 |
| LC_coassembly_k141_1014591 | 2810 | Yes | Unknown | *Kitrinoviricota* | *c.0034* | *o.0045* | *f.0176* | 2 |
| LC_coassembly_k141_1021355 | 7782 | Yes | Unknown | *Kitrinoviricota* | *c.0034* | *o.0045* | *f.0176* | 6 |
| LC_coassembly_k141_1021654 | 14734 | Yes | Unknown | *Kitrinoviricota* | *c.0034* | *o.0045* | *f.0176* | 9 |
| LC_coassembly_k141_1030321 | 2893 | Yes | Unknown | *Pisuviricota* | *c.0013* | *o.0025.base-Sobeli* | *f.0076* | 1 |
| LC_coassembly_k141_1046725 | 8509 | Yes | Unknown | *Duplornaviricota* | *Chrymotiviricetes* | *Ghabrivirales* | *Nam-Co_family_70* | 2 |
| LC_coassembly_k141_105435 | 4302 | Yes | dsRNA | *Duplornaviricota* | *Resentoviricetes* | *Reovirales* | *Reoviridae* | 1 |
| LC_coassembly_k141_1064585 | 4786 | Yes | Unknown | *Kitrinoviricota* | *c.0034* | *o.0045* | *f.0176* | 4 |
| LC_coassembly_k141_1074524 | 2047 | Yes | ssRNA(+) | *Lenarviricota* | *Howeltoviricetes* | *Cryppavirales* | *Mitoviridae* | 2 |
| LC_coassembly_k141_1091995 | 1310 | Yes | Unknown | *Pisuviricota* | *Pisoniviricetes* | *Picornavirales* | *f.0040.base-Marna* | 1 |
| LC_coassembly_k141_1111088 | 8448 | Yes | Unknown | *Kitrinoviricota* | *c.0034* | *o.0045* | *f.0176* | 4 |
| LC_coassembly_k141_1114277 | 845 | Yes | ssRNA(+) | *Lenarviricota* | *Amabiliviricetes* | *Wolframvirales* | *Narnaviridae* | 1 |
| LC_coassembly_k141_1116952 | 5862 | Yes | Unknown | *Kitrinoviricota* | *c.0034* | *o.0045* | *f.0176* | 3 |
| LC_coassembly_k141_1120492 | 2670 | Yes | ssRNA(+) | *Lenarviricota* | *Amabiliviricetes* | *Wolframvirales* | *Narnaviridae* | 1 |
| LC_coassembly_k141_1135645 | 1727 | Yes | Unknown | *Duplornaviricota* | *Chrymotiviricetes* | *Ghabrivirales* | *f.0283.base-Megabirna* | 1 |
| LC_coassembly_k141_1136066 | 861 | Yes | Unknown | *Duplornaviricota* | *Chrymotiviricetes* | *Ghabrivirales* | *f.0283.base-Megabirna* | 1 |
| LC_coassembly_k141_1136571 | 1235 | Yes | Unknown | *Pisuviricota* | *Duplopiviricetes* | *Durnavirales* | *f.0102.base-Curvula* | 1 |
| LC_coassembly_k141_1148680 | 12081 | Yes | Unknown | *Kitrinoviricota* | *c.0028* | *o.0039* | *f.0150* | 19 |
| LC_coassembly_k141_1151886 | 629 | No | NA | *NA* | *NA* | *NA* | *NA* | 1 |
| LC_coassembly_k141_1170521 | 3217 | Yes | ssRNA(+) | *Lenarviricota* | *Amabiliviricetes* | *Wolframvirales* | *Narnaviridae* | 1 |
| LC_coassembly_k141_1176970 | 6733 | Yes | Unknown | *Pisuviricota* | *Duplopiviricetes* | *Durnavirales* | *Nam-Co_family_26* | 6 |
| LC_coassembly_k141_1177772 | 4205 | Yes | ssRNA(+) | *Lenarviricota* | *Leviviricetes* | *Norzivirales* | *Atkinsviridae* | 4 |
| LC_coassembly_k141_1185643 | 1801 | Yes | Unknown | *Pisuviricota* | *Duplopiviricetes* | *Durnavirales* | *f.0111* | 1 |
| LC_coassembly_k141_1186528 | 979 | Yes | Unknown | *Pisuviricota* | *Duplopiviricetes* | *Durnavirales* | *f.0102.base-Curvula* | 1 |
| LC_coassembly_k141_1187871 | 3361 | Yes | Unknown | *Duplornaviricota* | *Chrymotiviricetes* | *Ghabrivirales* | *f.0296* | 2 |
| LC_coassembly_k141_1198831 | 2652 | Yes | Unknown | *Pisuviricota* | *Duplopiviricetes* | *Durnavirales* | *f.0111* | 1 |
| LC_coassembly_k141_1209836 | 1273 | Yes | ssRNA(+) | *Lenarviricota* | *Amabiliviricetes* | *Wolframvirales* | *Narnaviridae* | 1 |
| LC_coassembly_k141_1231733 | 9248 | Yes | ssRNA(+) | *Pisuviricota* | *Pisoniviricetes* | *Picornavirales* | *Marnaviridae* | 2 |
| LC_coassembly_k141_1259623 | 6678 | Yes | Unknown | *Kitrinoviricota* | *c.0034* | *o.0045* | *f.0176* | 5 |
| LC_coassembly_k141_125988 | 1601 | Yes | ssRNA(+) | *Lenarviricota* | *Howeltoviricetes* | *Cryppavirales* | *Mitoviridae* | 2 |
| LC_coassembly_k141_1264256 | 1949 | Yes | Unknown | *Duplornaviricota* | *Chrymotiviricetes* | *Ghabrivirales* | *f.0296* | 1 |
| LC_coassembly_k141_1271871 | 957 | Yes | ssRNA(+) | *Lenarviricota* | *Leviviricetes* | *Timlovirales* | *Steitzviridae* | 1 |
| LC_coassembly_k141_1276178 | 969 | Yes | Unknown | *Duplornaviricota* | *Chrymotiviricetes* | *Ghabrivirales* | *f.0296* | 1 |
| LC_coassembly_k141_1296109 | 8098 | Yes | Unknown | *Kitrinoviricota* | *c.0034* | *o.0045* | *f.0176* | 5 |
| LC_coassembly_k141_1296706 | 620 | Yes | ssRNA | *Pisuviricota* | *c.0005.base-Stelpa* | *o.0012* | *Hypoviridae* | 1 |
| LC_coassembly_k141_1335197 | 2689 | Yes | ssRNA(+) | *Pisuviricota* | *Pisoniviricetes* | *Picornavirales* | *Marnaviridae* | 1 |
| LC_coassembly_k141_133542 | 2545 | Yes | Unknown | *Pisuviricota* | *c.0013* | *o.0025.base-Sobeli* | *Nam-Co_family_29* | 1 |
| LC_coassembly_k141_1350294 | 8828 | Yes | Unknown | *Kitrinoviricota* | *c.0034* | *o.0045* | *f.0176* | 7 |
| LC_coassembly_k141_1350414 | 2833 | Yes | ssRNA(+) | *Lenarviricota* | *Amabiliviricetes* | *Wolframvirales* | *Narnaviridae* | 1 |
| LC_coassembly_k141_1369234 | 3269 | Yes | Unknown | *Pisuviricota* | *Duplopiviricetes* | *Durnavirales* | *Nam-Co_family_39* | 2 |
| LC_coassembly_k141_137237 | 2417 | Yes | Unknown | *Pisuviricota* | *c.0013* | *o.0025.base-Sobeli* | *f.0076* | 1 |
| LC_coassembly_k141_138224 | 3390 | Yes | ssRNA | *Pisuviricota* | *c.0005.base-Stelpa* | *o.0012* | *Hypoviridae* | 1 |
| LC_coassembly_k141_1385123 | 708 | Yes | Unknown | *Kitrinoviricota* | *c.0021.base-Toluca* | *o.0032* | *f.0125* | 1 |
| LC_coassembly_k141_1394068 | 7293 | Yes | ssRNA(+) | *Pisuviricota* | *Pisoniviricetes* | *Picornavirales* | *Picornaviridae* | 1 |
| LC_coassembly_k141_1396795 | 2531 | Yes | ssRNA(+) | *Lenarviricota* | *Amabiliviricetes* | *Wolframvirales* | *Narnaviridae* | 1 |
| LC_coassembly_k141_1407661 | 9645 | Yes | Unknown | *Negarnaviricota* | *Ellioviricetes* | *Bunyavirales* | *f.0315* | 1 |
| LC_coassembly_k141_14177 | 1710 | Yes | ssRNA(+) | *Lenarviricota* | *Miaviricetes* | *Ourlivirales* | *Botourmiaviridae* | 1 |
| LC_coassembly_k141_1421666 | 1273 | Yes | Unknown | *Duplornaviricota* | *Chrymotiviricetes* | *Ghabrivirales* | *f.0296* | 1 |
| LC_coassembly_k141_1421816 | 1503 | Yes | Unknown | *Pisuviricota* | *Duplopiviricetes* | *Durnavirales* | *f.0102.base-Curvula* | 1 |
| LC_coassembly_k141_1432413 | 1450 | Yes | ssRNA(+) | *Pisuviricota* | *Pisoniviricetes* | *Picornavirales* | *Marnaviridae* | 2 |
| LC_coassembly_k141_1445686 | 1583 | Yes | Unknown | *Duplornaviricota* | *Chrymotiviricetes* | *Ghabrivirales* | *f.0283.base-Megabirna* | 1 |
| LC_coassembly_k141_147022 | 851 | Yes | ssRNA(+) | *Lenarviricota* | *Amabiliviricetes* | *Wolframvirales* | *Narnaviridae* | 1 |
| LC_coassembly_k141_1474085 | 650 | Yes | ssRNA(+) | *Pisuviricota* | *Pisoniviricetes* | *Picornavirales* | *Marnaviridae* | 1 |
| LC_coassembly_k141_1476401 | 6642 | Yes | Unknown | *Kitrinoviricota* | *c.0032* | *o.0043* | *f.0167* | 4 |
| LC_coassembly_k141_1484123 | 1077 | Yes | Unknown | *Pisuviricota* | *Duplopiviricetes* | *Durnavirales* | *f.0111* | 2 |
| LC_coassembly_k141_1485030 | 1821 | Yes | Unknown | *Duplornaviricota* | *Chrymotiviricetes* | *Ghabrivirales* | *Nam-Co_family_65* | 1 |
| LC_coassembly_k141_1486019 | 11716 | Yes | ssRNA(+) | *Kitrinoviricota* | *Alsuviricetes* | *Martellivirales* | *Virgaviridae* | 1 |
| LC_coassembly_k141_1486921 | 988 | Yes | ssRNA(+) | *Lenarviricota* | *Howeltoviricetes* | *Cryppavirales* | *Mitoviridae* | 2 |
| LC_coassembly_k141_1498540 | 840 | Yes | ssRNA(+) | *Lenarviricota* | *Miaviricetes* | *Ourlivirales* | *Botourmiaviridae* | 1 |
| LC_coassembly_k141_1498943 | 3330 | Yes | ssRNA(+) | *Lenarviricota* | *Amabiliviricetes* | *Wolframvirales* | *Narnaviridae* | 1 |
| LC_coassembly_k141_1526117 | 8801 | Yes | Unknown | *Duplornaviricota* | *Chrymotiviricetes* | *Ghabrivirales* | *f.0295* | 3 |
| LC_coassembly_k141_1528827 | 1725 | Yes | Unknown | *Duplornaviricota* | *Chrymotiviricetes* | *Ghabrivirales* | *f.0282.base-Megabirna* | 1 |
| LC_coassembly_k141_1533203 | 1909 | Yes | Unknown | *Duplornaviricota* | *Chrymotiviricetes* | *Ghabrivirales* | *f.0296* | 1 |
| LC_coassembly_k141_1538122 | 3879 | Yes | ssRNA(+) | *Lenarviricota* | *Amabiliviricetes* | *Wolframvirales* | *Narnaviridae* | 2 |
| LC_coassembly_k141_1539147 | 4342 | Yes | ssRNA(+) | *Pisuviricota* | *Pisoniviricetes* | *Picornavirales* | *Dicistroviridae* | 3 |
| LC_coassembly_k141_1544798 | 3475 | Yes | ssRNA(+) | *Lenarviricota* | *Amabiliviricetes* | *Wolframvirales* | *Nam-Co_family_12* | 1 |
| LC_coassembly_k141_1576953 | 874 | Yes | ssRNA(+) | *Kitrinoviricota* | *Alsuviricetes* | *o.0055* | *Endornaviridae* | 1 |
| LC_coassembly_k141_1578211 | 1027 | Yes | Unknown | *Pisuviricota* | *Duplopiviricetes* | *Durnavirales* | *Nam-Co_family_35* | 1 |
| LC_coassembly_k141_1594068 | 2207 | Yes | Unknown | *Pisuviricota* | *c.0013* | *o.0025.base-Sobeli* | *f.0076* | 1 |
| LC_coassembly_k141_159514 | 951 | No | NA | *NA* | *NA* | *NA* | *NA* | 1 |
| LC_coassembly_k141_1595254 | 775 | Yes | Unknown | *Lenarviricota* | *c.0069.base-Howelto* | *o.0090* | *f.0386* | 1 |
| LC_coassembly_k141_1596549 | 1261 | No | NA | *NA* | *NA* | *NA* | *Nam-Co_family_47* | 2 |
| LC_coassembly_k141_1606298 | 8258 | Yes | Unknown | *Pisuviricota* | *Pisoniviricetes* | *Picornavirales* | *Nam-Co_family_44* | 2 |
| LC_coassembly_k141_1614393 | 2033 | Yes | Unknown | *Duplornaviricota* | *Chrymotiviricetes* | *Ghabrivirales* | *f.0283.base-Megabirna* | 2 |
| LC_coassembly_k141_1642998 | 7239 | Yes | Unknown | *Duplornaviricota* | *Chrymotiviricetes* | *Ghabrivirales* | *f.0283.base-Megabirna* | 2 |
| LC_coassembly_k141_16449 | 2900 | Yes | Unknown | *Pisuviricota* | *Duplopiviricetes* | *Durnavirales* | *f.0102.base-Curvula* | 2 |
| LC_coassembly_k141_1666324 | 943 | Yes | Unknown | *Duplornaviricota* | *Chrymotiviricetes* | *Ghabrivirales* | *f.0285* | 1 |
| LC_coassembly_k141_1669236 | 1392 | Yes | Unknown | *Lenarviricota* | *c.0074.base-Howelto* | *o.0095* | *Nam-Co_family_16* | 1 |
| LC_coassembly_k141_1669284 | 984 | Yes | ssRNA(+) | *Lenarviricota* | *Howeltoviricetes* | *Cryppavirales* | *Mitoviridae* | 1 |
| LC_coassembly_k141_1675587 | 1195 | Yes | Unknown | *Pisuviricota* | *Duplopiviricetes* | *Durnavirales* | *f.0102.base-Curvula* | 1 |
| LC_coassembly_k141_1680250 | 646 | Yes | Unknown | *Duplornaviricota* | *Chrymotiviricetes* | *Ghabrivirales* | *f.0283.base-Megabirna* | 1 |
| LC_coassembly_k141_1686962 | 1347 | Yes | Unknown | *Pisuviricota* | *c.0003.base-Pisoni* | *o.0003* | *Nam-Co_family_27* | 1 |
| LC_coassembly_k141_1687493 | 3098 | Yes | Unknown | *Kitrinoviricota* | *c.0034* | *o.0045* | *f.0176* | 2 |
| LC_coassembly_k141_1689211 | 2073 | Yes | Unknown | *Pisuviricota* | *Duplopiviricetes* | *Durnavirales* | *Nam-Co_family_39* | 3 |
| LC_coassembly_k141_1727070 | 7901 | Yes | Unknown | *Duplornaviricota* | *Chrymotiviricetes* | *Ghabrivirales* | *f.0287* | 7 |
| LC_coassembly_k141_1727227 | 1282 | Yes | ssRNA(+) | *Lenarviricota* | *Howeltoviricetes* | *Cryppavirales* | *Mitoviridae* | 1 |
| LC_coassembly_k141_1739317 | 3063 | Yes | Unknown | *Pisuviricota* | *Duplopiviricetes* | *Durnavirales* | *f.0102.base-Curvula* | 2 |
| LC_coassembly_k141_1747163 | 6675 | Yes | Unknown | *Kitrinoviricota* | *c.0034* | *o.0045* | *f.0176* | 4 |
| LC_coassembly_k141_1754048 | 2727 | Yes | ssRNA(+) | *Lenarviricota* | *Amabiliviricetes* | *Wolframvirales* | *Narnaviridae* | 1 |
| LC_coassembly_k141_1759012 | 8445 | Yes | Unknown | *Duplornaviricota* | *Chrymotiviricetes* | *Ghabrivirales* | *Nam-Co_family_71* | 4 |
| LC_coassembly_k141_1763940 | 772 | Yes | Unknown | *Duplornaviricota* | *Chrymotiviricetes* | *Ghabrivirales* | *f.0283.base-Megabirna* | 1 |
| LC_coassembly_k141_1767469 | 2947 | Yes | dsRNA | *Duplornaviricota* | *Chrymotiviricetes* | *Ghabrivirales* | *Totiviridae* | 1 |
| LC_coassembly_k141_1773142 | 2931 | Yes | ssRNA(+) | *Kitrinoviricota* | *Tolucaviricetes* | *Tolivirales* | *Tombusviridae* | 3 |
| LC_coassembly_k141_1773347 | 1273 | Yes | ssRNA(+) | *Lenarviricota* | *Amabiliviricetes* | *Wolframvirales* | *Narnaviridae* | 1 |
| LC_coassembly_k141_1779233 | 1326 | Yes | Unknown | *Duplornaviricota* | *Chrymotiviricetes* | *Ghabrivirales* | *f.0283.base-Megabirna* | 1 |
| LC_coassembly_k141_1788793 | 849 | Yes | Unknown | *Duplornaviricota* | *Chrymotiviricetes* | *Ghabrivirales* | *f.0296* | 1 |
| LC_coassembly_k141_1805655 | 794 | Yes | ssRNA(+) | *Lenarviricota* | *Leviviricetes* | *Norzivirales* | *Fiersviridae* | 1 |
| LC_coassembly_k141_1810732 | 1216 | Yes | Unknown | *Duplornaviricota* | *Chrymotiviricetes* | *Ghabrivirales* | *f.0283.base-Megabirna* | 1 |
| LC_coassembly_k141_1811975 | 2298 | Yes | Unknown | *Pisuviricota* | *Duplopiviricetes* | *Durnavirales* | *f.0120* | 1 |
| LC_coassembly_k141_1814031 | 3068 | Yes | ssRNA(+) | *Lenarviricota* | *Amabiliviricetes* | *Wolframvirales* | *Narnaviridae* | 1 |
| LC_coassembly_k141_1817896 | 1258 | Yes | ssRNA(+) | *Lenarviricota* | *Leviviricetes* | *Norzivirales* | *Atkinsviridae* | 1 |
| LC_coassembly_k141_1827790 | 957 | Yes | ssRNA(+) | *Lenarviricota* | *Leviviricetes* | *Norzivirales* | *Fiersviridae* | 1 |
| LC_coassembly_k141_1828284 | 2814 | Yes | ssRNA(+) | *Lenarviricota* | *Amabiliviricetes* | *Wolframvirales* | *Narnaviridae* | 1 |
| LC_coassembly_k141_1831437 | 1510 | Yes | Unknown | *Pisuviricota* | *Duplopiviricetes* | *Durnavirales* | *f.0111* | 1 |
| LC_coassembly_k141_1836560 | 2376 | Yes | dsRNA | *Duplornaviricota* | *Chrymotiviricetes* | *Ghabrivirales* | *Totiviridae* | 1 |
| LC_coassembly_k141_1840495 | 1141 | Yes | Unknown | *Duplornaviricota* | *Chrymotiviricetes* | *Ghabrivirales* | *f.0283.base-Megabirna* | 2 |
| LC_coassembly_k141_1841077 | 1230 | Yes | ssRNA(+) | *Kitrinoviricota* | *Magsaviricetes* | *Nodamuvirales* | *Nodaviridae* | 1 |
| LC_coassembly_k141_1841635 | 703 | Yes | Unknown | *Pisuviricota* | *c.0013* | *o.0025.base-Sobeli* | *f.0076* | 1 |
| LC_coassembly_k141_1848390 | 2554 | Yes | ssRNA(+) | *Pisuviricota* | *Stelpaviricetes* | *Stellavirales* | *Astroviridae* | 2 |
| LC_coassembly_k141_1863880 | 3774 | Yes | Unknown | *Negarnaviricota* | *Ellioviricetes* | *Bunyavirales* | *f.0327* | 1 |
| LC_coassembly_k141_1868495 | 722 | Yes | Unknown | *Duplornaviricota* | *Chrymotiviricetes* | *Ghabrivirales* | *f.0285* | 1 |
| LC_coassembly_k141_1872912 | 1694 | Yes | Unknown | *Pisuviricota* | *c.0013* | *o.0025.base-Sobeli* | *Nam-Co_family_29* | 1 |
| LC_coassembly_k141_1873206 | 3100 | Yes | ssRNA(+) | *Lenarviricota* | *Howeltoviricetes* | *Cryppavirales* | *Nam-Co_family_17* | 2 |
| LC_coassembly_k141_1887030 | 5066 | Yes | ssRNA(+) | *Pisuviricota* | *Pisoniviricetes* | *Picornavirales* | *Marnaviridae* | 2 |
| LC_coassembly_k141_1892891 | 2616 | Yes | Unknown | *Duplornaviricota* | *Chrymotiviricetes* | *Ghabrivirales* | *f.0296* | 2 |
| LC_coassembly_k141_1897578 | 1457 | Yes | ssRNA(+) | *Lenarviricota* | *Howeltoviricetes* | *Cryppavirales* | *Mitoviridae* | 1 |
| LC_coassembly_k141_1900159 | 997 | Yes | ssRNA(+) | *Lenarviricota* | *Howeltoviricetes* | *Cryppavirales* | *Mitoviridae* | 1 |
| LC_coassembly_k141_1913062 | 4539 | Yes | Unknown | *Kitrinoviricota* | *c.0031* | *o.0042* | *f.0153* | 2 |
| LC_coassembly_k141_1938999 | 702 | Yes | ssRNA(+) | *Lenarviricota* | *Amabiliviricetes* | *Wolframvirales* | *Narnaviridae* | 1 |
| LC_coassembly_k141_1955953 | 738 | Yes | Unknown | *Duplornaviricota* | *Chrymotiviricetes* | *Ghabrivirales* | *f.0283.base-Megabirna* | 1 |
| LC_coassembly_k141_196466 | 6309 | Yes | ssRNA(+) | *Kitrinoviricota* | *Alsuviricetes* | *Martellivirales* | *Virgaviridae* | 4 |
| LC_coassembly_k141_1968509 | 663 | Yes | ssRNA(+) | *Lenarviricota* | *Leviviricetes* | *Norzivirales* | *Atkinsviridae* | 1 |
| LC_coassembly_k141_1984682 | 1503 | Yes | ssRNA(+) | *Lenarviricota* | *Amabiliviricetes* | *Wolframvirales* | *Narnaviridae* | 1 |
| LC_coassembly_k141_1995525 | 794 | Yes | Unknown | *Duplornaviricota* | *Chrymotiviricetes* | *Ghabrivirales* | *f.0283.base-Megabirna* | 1 |
| LC_coassembly_k141_1998157 | 1587 | Yes | Unknown | *Pisuviricota* | *c.0013* | *o.0025.base-Sobeli* | *f.0076* | 1 |
| LC_coassembly_k141_2019234 | 1307 | Yes | Unknown | *Duplornaviricota* | *Chrymotiviricetes* | *Ghabrivirales* | *f.0296* | 1 |
| LC_coassembly_k141_2042949 | 607 | Yes | ssRNA(+) | *Pisuviricota* | *Pisoniviricetes* | *Picornavirales* | *Marnaviridae* | 1 |
| LC_coassembly_k141_2045611 | 8433 | Yes | Unknown | *Kitrinoviricota* | *c.0034* | *o.0045* | *f.0176* | 6 |
| LC_coassembly_k141_204657 | 4414 | Yes | Unknown | *Kitrinoviricota* | *c.0031* | *o.0042* | *f.0153* | 3 |
| LC_coassembly_k141_2048581 | 1004 | Yes | Unknown | *Pisuviricota* | *Pisoniviricetes* | *Picornavirales* | *f.0030* | 1 |
| LC_coassembly_k141_2048707 | 859 | Yes | Unknown | *Duplornaviricota* | *Chrymotiviricetes* | *Ghabrivirales* | *Nam-Co_family_66* | 1 |
| LC_coassembly_k141_2084125 | 2756 | Yes | Unknown | *Pisuviricota* | *c.0013* | *o.0025.base-Sobeli* | *f.0076* | 1 |
| LC_coassembly_k141_210205 | 1635 | Yes | Unknown | *Duplornaviricota* | *Chrymotiviricetes* | *Ghabrivirales* | *f.0296* | 1 |
| LC_coassembly_k141_2104764 | 654 | Yes | ssRNA(+) | *Kitrinoviricota* | *Alsuviricetes* | *Martellivirales* | *Virgaviridae* | 1 |
| LC_coassembly_k141_2106280 | 1878 | Yes | ssRNA(+) | *Lenarviricota* | *Howeltoviricetes* | *Cryppavirales* | *Mitoviridae* | 2 |
| LC_coassembly_k141_2111917 | 1067 | Yes | ssRNA(+) | *Lenarviricota* | *Howeltoviricetes* | *Cryppavirales* | *Mitoviridae* | 1 |
| LC_coassembly_k141_2127533 | 2113 | Yes | Unknown | *Lenarviricota* | *c.0064.base-Howelto* | *o.0085* | *f.0381* | 1 |
| LC_coassembly_k141_213387 | 799 | Yes | ssRNA(+) | *Lenarviricota* | *Howeltoviricetes* | *Cryppavirales* | *Mitoviridae* | 1 |
| LC_coassembly_k141_2134336 | 4582 | Yes | Unknown | *Negarnaviricota* | *Ellioviricetes* | *Bunyavirales* | *f.0313* | 1 |
| LC_coassembly_k141_2139485 | 8251 | Yes | ssRNA(+) | *Pisuviricota* | *Pisoniviricetes* | *Picornavirales* | *Dicistroviridae* | 1 |
| LC_coassembly_k141_2145116 | 2141 | Yes | ssRNA(+) | *Pisuviricota* | *Pisoniviricetes* | *Picornavirales* | *Marnaviridae* | 2 |
| LC_coassembly_k141_2159845 | 4510 | Yes | ssRNA(+) | *Pisuviricota* | *c.0012* | *Sobelivirales* | *Solemoviridae* | 4 |
| LC_coassembly_k141_2160220 | 3377 | No | NA | *NA* | *NA* | *NA* | *NA* | 2 |
| LC_coassembly_k141_216290 | 917 | Yes | Unknown | *Kitrinoviricota* | *c.0034* | *o.0045* | *f.0176* | 1 |
| LC_coassembly_k141_2164532 | 5565 | Yes | Unknown | *Kitrinoviricota* | *c.0034* | *o.0045* | *f.0176* | 4 |
| LC_coassembly_k141_2171829 | 1779 | Yes | dsRNA | *Duplornaviricota* | *Chrymotiviricetes* | *Ghabrivirales* | *Totiviridae* | 2 |
| LC_coassembly_k141_2172855 | 1352 | Yes | Unknown | *Duplornaviricota* | *Chrymotiviricetes* | *Ghabrivirales* | *f.0283.base-Megabirna* | 1 |
| LC_coassembly_k141_2173116 | 1198 | Yes | Unknown | *Kitrinoviricota* | *c.0031* | *o.0042* | *f.0153* | 1 |
| LC_coassembly_k141_2173443 | 749 | No | NA | *NA* | *NA* | *NA* | *NA* | 1 |
| LC_coassembly_k141_2175510 | 1699 | Yes | Unknown | *Duplornaviricota* | *Chrymotiviricetes* | *Ghabrivirales* | *Nam-Co_family_72* | 1 |
| LC_coassembly_k141_217665 | 2527 | Yes | Unknown | *Pisuviricota* | *c.0013* | *o.0025.base-Sobeli* | *f.0076* | 1 |
| LC_coassembly_k141_2187462 | 6506 | Yes | Unknown | *Duplornaviricota* | *Chrymotiviricetes* | *Ghabrivirales* | *f.0296* | 3 |
| LC_coassembly_k141_219128 | 7840 | No | NA | *NA* | *NA* | *NA* | *Nam-Co_family_73* | 3 |
| LC_coassembly_k141_2206238 | 3992 | Yes | dsRNA | *Pisuviricota* | *Duplopiviricetes* | *Durnavirales* | *Nam-Co_family_36* | 3 |
| LC_coassembly_k141_2214956 | 1036 | Yes | Unknown | *Duplornaviricota* | *Chrymotiviricetes* | *Ghabrivirales* | *f.0296* | 1 |
| LC_coassembly_k141_221694 | 702 | Yes | ssRNA(+) | *Lenarviricota* | *Amabiliviricetes* | *Wolframvirales* | *Narnaviridae* | 1 |
| LC_coassembly_k141_2234358 | 2687 | Yes | ssRNA(+) | *Lenarviricota* | *Amabiliviricetes* | *Wolframvirales* | *Narnaviridae* | 1 |
| LC_coassembly_k141_2239335 | 2028 | Yes | ssRNA(+) | *Pisuviricota* | *Pisoniviricetes* | *Picornavirales* | *Marnaviridae* | 1 |
| LC_coassembly_k141_2258078 | 3348 | Yes | Unknown | *Kitrinoviricota* | *c.0034* | *o.0045* | *f.0176* | 2 |
| LC_coassembly_k141_2261207 | 1520 | Yes | ssRNA(+) | *Lenarviricota* | *Amabiliviricetes* | *Wolframvirales* | *Narnaviridae* | 1 |
| LC_coassembly_k141_2261759 | 2356 | Yes | ssRNA(+) | *Lenarviricota* | *Amabiliviricetes* | *Wolframvirales* | *Narnaviridae* | 1 |
| LC_coassembly_k141_2266943 | 2073 | Yes | Unknown | *Duplornaviricota* | *Chrymotiviricetes* | *Ghabrivirales* | *Nam-Co_family_66* | 1 |
| LC_coassembly_k141_2273513 | 1077 | Yes | ssRNA(+) | *Lenarviricota* | *Howeltoviricetes* | *Cryppavirales* | *Mitoviridae* | 1 |
| LC_coassembly_k141_227913 | 1049 | Yes | ssRNA(+) | *Lenarviricota* | *Miaviricetes* | *Ourlivirales* | *Botourmiaviridae* | 1 |
| LC_coassembly_k141_2285013 | 697 | Yes | ssRNA(+) | *Pisuviricota* | *Pisoniviricetes* | *Picornavirales* | *Nam-Co_family_43* | 1 |
| LC_coassembly_k141_2292250 | 1265 | Yes | ssRNA(+) | *Lenarviricota* | *Amabiliviricetes* | *Wolframvirales* | *Narnaviridae* | 1 |
| LC_coassembly_k141_2300750 | 2590 | Yes | Unknown | *Kitrinoviricota* | *c.0034* | *o.0045* | *f.0176* | 2 |
| LC_coassembly_k141_2312099 | 6172 | Yes | Unknown | *Negarnaviricota* | *Ellioviricetes* | *Bunyavirales* | *f.0305.base-Leishbu* | 1 |
| LC_coassembly_k141_2314916 | 2639 | Yes | Unknown | *Duplornaviricota* | *Chrymotiviricetes* | *Ghabrivirales* | *f.0283.base-Megabirna* | 1 |
| LC_coassembly_k141_2315390 | 682 | Yes | ssRNA(+) | *Lenarviricota* | *Leviviricetes* | *Norzivirales* | *Atkinsviridae* | 1 |
| LC_coassembly_k141_2331475 | 6310 | Yes | Unknown | *Kitrinoviricota* | *c.0032* | *o.0043* | *f.0167* | 3 |
| LC_coassembly_k141_2332014 | 1134 | Yes | Unknown | *Duplornaviricota* | *Chrymotiviricetes* | *Ghabrivirales* | *f.0287* | 1 |
| LC_coassembly_k141_2347465 | 707 | Yes | Unknown | *Pisuviricota* | *c.0013* | *o.0025.base-Sobeli* | *f.0076* | 1 |
| LC_coassembly_k141_2368185 | 1199 | Yes | ssRNA(+) | *Lenarviricota* | *Amabiliviricetes* | *Wolframvirales* | *Narnaviridae* | 1 |
| LC_coassembly_k141_2371561 | 2600 | Yes | ssRNA(+) | *Lenarviricota* | *Amabiliviricetes* | *Wolframvirales* | *Narnaviridae* | 1 |
| LC_coassembly_k141_2383019 | 2787 | Yes | ssRNA(+) | *Lenarviricota* | *Amabiliviricetes* | *Wolframvirales* | *Narnaviridae* | 1 |
| LC_coassembly_k141_2401611 | 7841 | Yes | Unknown | *Duplornaviricota* | *Chrymotiviricetes* | *Ghabrivirales* | *Nam-Co_family_74* | 5 |
| LC_coassembly_k141_2410216 | 1335 | Yes | ssRNA(+) | *Kitrinoviricota* | *Alsuviricetes* | *o.0055* | *Endornaviridae* | 1 |
| LC_coassembly_k141_2413734 | 2033 | Yes | Unknown | *Kitrinoviricota* | *c.0034* | *o.0045* | *f.0176* | 2 |
| LC_coassembly_k141_2414581 | 670 | Yes | Unknown | *Pisuviricota* | *Pisoniviricetes* | *Picornavirales* | *Nam-Co_family_45* | 1 |
| LC_coassembly_k141_2418593 | 3456 | Yes | Unknown | *Duplornaviricota* | *Chrymotiviricetes* | *Ghabrivirales* | *Nam-Co_family_74* | 4 |
| LC_coassembly_k141_242337 | 1615 | Yes | ssRNA(+) | *Lenarviricota* | *Howeltoviricetes* | *Cryppavirales* | *Mitoviridae* | 1 |
| LC_coassembly_k141_2427120 | 5955 | Yes | Unknown | *Pisuviricota* | *Pisoniviricetes* | *Picornavirales* | *f.0028.base-Dicistro* | 1 |
| LC_coassembly_k141_2428978 | 2659 | Yes | ssRNA(+) | *Lenarviricota* | *Howeltoviricetes* | *Cryppavirales* | *Mitoviridae* | 1 |
| LC_coassembly_k141_2449152 | 2006 | Yes | ssRNA(+) | *Kitrinoviricota* | *Tolucaviricetes* | *Tolivirales* | *Tombusviridae* | 2 |
| LC_coassembly_k141_2450679 | 1115 | Yes | Unknown | *Pisuviricota* | *c.0013* | *o.0025.base-Sobeli* | *f.0076* | 1 |
| LC_coassembly_k141_2452354 | 1413 | Yes | Unknown | *Duplornaviricota* | *Chrymotiviricetes* | *Ghabrivirales* | *f.0296* | 1 |
| LC_coassembly_k141_2459206 | 1851 | No | NA | *NA* | *NA* | *NA* | *NA* | 2 |
| LC_coassembly_k141_2464698 | 2898 | Yes | Unknown | *Pisuviricota* | *Duplopiviricetes* | *Durnavirales* | *f.0102.base-Curvula* | 3 |
| LC_coassembly_k141_2476813 | 823 | Yes | ssRNA(+) | *Kitrinoviricota* | *Alsuviricetes* | *o.0055* | *Endornaviridae* | 1 |
| LC_coassembly_k141_2477916 | 1882 | Yes | Unknown | *Duplornaviricota* | *Chrymotiviricetes* | *Ghabrivirales* | *f.0283.base-Megabirna* | 1 |
| LC_coassembly_k141_2485855 | 624 | Yes | ssRNA(+) | *Lenarviricota* | *Amabiliviricetes* | *Wolframvirales* | *Narnaviridae* | 1 |
| LC_coassembly_k141_2488396 | 706 | Yes | ssRNA(+) | *Lenarviricota* | *Howeltoviricetes* | *Cryppavirales* | *Mitoviridae* | 1 |
| LC_coassembly_k141_2501884 | 2963 | Yes | Unknown | *Pisuviricota* | *Duplopiviricetes* | *Durnavirales* | *f.0102.base-Curvula* | 2 |
| LC_coassembly_k141_2512221 | 2496 | Yes | Unknown | *Pisuviricota* | *c.0013* | *o.0025.base-Sobeli* | *f.0076* | 1 |
| LC_coassembly_k141_2537970 | 2697 | Yes | Unknown | *Duplornaviricota* | *Chrymotiviricetes* | *Ghabrivirales* | *f.0282.base-Megabirna* | 2 |
| LC_coassembly_k141_2541618 | 867 | Yes | ssRNA(+) | *Lenarviricota* | *Leviviricetes* | *Timlovirales* | *Steitzviridae* | 1 |
| LC_coassembly_k141_2545102 | 1034 | Yes | Unknown | *Pisuviricota* | *Duplopiviricetes* | *Durnavirales* | *f.0111* | 1 |
| LC_coassembly_k141_2551561 | 1707 | Yes | Unknown | *Kitrinoviricota* | *c.0021.base-Toluca* | *o.0032* | *f.0125* | 2 |
| LC_coassembly_k141_2570745 | 4935 | No | NA | *Negarnaviricota* | *NA* | *NA* | *Nam-Co_family_79* | 1 |
| LC_coassembly_k141_2580714 | 7373 | Yes | ssRNA(+) | *Kitrinoviricota* | *Magsaviricetes* | *Nodamuvirales* | *Nodaviridae* | 2 |
| LC_coassembly_k141_2580908 | 1363 | Yes | dsRNA | *Pisuviricota* | *Duplopiviricetes* | *Durnavirales* | *Partitiviridae* | 1 |
| LC_coassembly_k141_2586978 | 2905 | Yes | ssRNA(+) | *Lenarviricota* | *Amabiliviricetes* | *Wolframvirales* | *Narnaviridae* | 1 |
| LC_coassembly_k141_2599729 | 2538 | Yes | Unknown | *Duplornaviricota* | *Chrymotiviricetes* | *Ghabrivirales* | *Nam-Co_family_74* | 3 |
| LC_coassembly_k141_2612247 | 1042 | Yes | Unknown | *Pisuviricota* | *c.0013* | *o.0025.base-Sobeli* | *f.0076* | 1 |
| LC_coassembly_k141_2619831 | 11233 | Yes | Unknown | *Kitrinoviricota* | *c.0028* | *o.0039* | *f.0150* | 15 |
| LC_coassembly_k141_2622128 | 627 | Yes | Unknown | *Duplornaviricota* | *Chrymotiviricetes* | *Ghabrivirales* | *f.0283.base-Megabirna* | 1 |
| LC_coassembly_k141_2634150 | 987 | Yes | ssRNA(+) | *Pisuviricota* | *Pisoniviricetes* | *Picornavirales* | *Polycipiviridae* | 1 |
| LC_coassembly_k141_2638362 | 1759 | Yes | ssRNA(+) | *Lenarviricota* | *Amabiliviricetes* | *Wolframvirales* | *Narnaviridae* | 2 |
| LC_coassembly_k141_2642329 | 637 | Yes | Unknown | *Pisuviricota* | *Duplopiviricetes* | *Durnavirales* | *f.0111* | 1 |
| LC_coassembly_k141_2655355 | 718 | Yes | Unknown | *Duplornaviricota* | *Chrymotiviricetes* | *Ghabrivirales* | *f.0296* | 1 |
| LC_coassembly_k141_2670924 | 1387 | Yes | ssRNA(+) | *Pisuviricota* | *Pisoniviricetes* | *Picornavirales* | *Dicistroviridae* | 2 |
| LC_coassembly_k141_2686142 | 2085 | Yes | ssRNA(+) | *Lenarviricota* | *Howeltoviricetes* | *Cryppavirales* | *Mitoviridae* | 1 |
| LC_coassembly_k141_2695273 | 1180 | Yes | ssRNA(+) | *Lenarviricota* | *Howeltoviricetes* | *Cryppavirales* | *Mitoviridae* | 1 |
| LC_coassembly_k141_2698212 | 1791 | Yes | ssRNA(+) | *Lenarviricota* | *Miaviricetes* | *Ourlivirales* | *Botourmiaviridae* | 1 |
| LC_coassembly_k141_2721539 | 2120 | Yes | ssRNA(+) | *Lenarviricota* | *Howeltoviricetes* | *Cryppavirales* | *Mitoviridae* | 1 |
| LC_coassembly_k141_2723452 | 687 | Yes | ssRNA(+) | *Lenarviricota* | *Howeltoviricetes* | *Cryppavirales* | *Mitoviridae* | 1 |
| LC_coassembly_k141_2728316 | 1846 | Yes | ssRNA(+) | *Lenarviricota* | *Howeltoviricetes* | *Cryppavirales* | *Mitoviridae* | 1 |
| LC_coassembly_k141_2733097 | 8267 | Yes | ssRNA(+) | *Pisuviricota* | *Pisoniviricetes* | *Picornavirales* | *Marnaviridae* | 2 |
| LC_coassembly_k141_2733507 | 8477 | Yes | ssRNA(+) | *Pisuviricota* | *Pisoniviricetes* | *Picornavirales* | *Marnaviridae* | 2 |
| LC_coassembly_k141_2737082 | 4972 | Yes | Unknown | *Kitrinoviricota* | *c.0025* | *o.0036* | *f.0145* | 2 |
| LC_coassembly_k141_2752873 | 1819 | Yes | ssRNA(+) | *Lenarviricota* | *Howeltoviricetes* | *Cryppavirales* | *Mitoviridae* | 1 |
| LC_coassembly_k141_2755829 | 2200 | Yes | dsRNA | *Pisuviricota* | *Duplopiviricetes* | *Durnavirales* | *Nam-Co_family_31* | 2 |
| LC_coassembly_k141_2759094 | 1555 | Yes | Unknown | *Pisuviricota* | *Duplopiviricetes* | *Durnavirales* | *f.0111* | 1 |
| LC_coassembly_k141_2762379 | 2145 | Yes | Unknown | *Duplornaviricota* | *Chrymotiviricetes* | *Ghabrivirales* | *f.0296* | 1 |
| LC_coassembly_k141_276839 | 3083 | Yes | Unknown | *Duplornaviricota* | *Chrymotiviricetes* | *Ghabrivirales* | *f.0282.base-Megabirna* | 1 |
| LC_coassembly_k141_2770412 | 647 | Yes | ssRNA(+) | *Lenarviricota* | *Amabiliviricetes* | *Wolframvirales* | *Narnaviridae* | 1 |
| LC_coassembly_k141_2771384 | 2108 | Yes | Unknown | *Pisuviricota* | *Duplopiviricetes* | *Durnavirales* | *f.0111* | 1 |
| LC_coassembly_k141_2772377 | 2338 | Yes | ssRNA(+) | *Kitrinoviricota* | *Alsuviricetes* | *o.0055* | *Endornaviridae* | 2 |
| LC_coassembly_k141_2783551 | 5257 | Yes | dsRNA | *Duplornaviricota* | *Chrymotiviricetes* | *Ghabrivirales* | *Totiviridae* | 2 |
| LC_coassembly_k141_2800168 | 1240 | Yes | Unknown | *Duplornaviricota* | *Chrymotiviricetes* | *Ghabrivirales* | *f.0283.base-Megabirna* | 1 |
| LC_coassembly_k141_284008 | 4281 | Yes | ssRNA(+) | *Pisuviricota* | *Pisoniviricetes* | *Picornavirales* | *Marnaviridae* | 2 |
| LC_coassembly_k141_303976 | 1716 | Yes | ssRNA(+) | *Lenarviricota* | *Miaviricetes* | *Ourlivirales* | *Botourmiaviridae* | 1 |
| LC_coassembly_k141_309995 | 1425 | No | NA | *NA* | *NA* | *NA* | *Nam-Co_family_02* | 1 |
| LC_coassembly_k141_31205 | 2558 | Yes | Unknown | *Kitrinoviricota* | *c.0034* | *o.0045* | *f.0176* | 3 |
| LC_coassembly_k141_328812 | 1572 | Yes | ssRNA(+) | *Lenarviricota* | *Howeltoviricetes* | *Cryppavirales* | *Mitoviridae* | 1 |
| LC_coassembly_k141_330298 | 612 | Yes | dsRNA | *Pisuviricota* | *c.0017* | *o.0028* | *Nam-Co_family_03* | 1 |
| LC_coassembly_k141_339916 | 1590 | Yes | Unknown | *Negarnaviricota* | *Ellioviricetes* | *Bunyavirales* | *Nam-Co_family_80* | 1 |
| LC_coassembly_k141_365724 | 2153 | Yes | Unknown | *Pisuviricota* | *Pisoniviricetes* | *Picornavirales* | *f.0025* | 1 |
| LC_coassembly_k141_367052 | 3464 | Yes | Unknown | *Pisuviricota* | *Duplopiviricetes* | *Durnavirales* | *f.0102.base-Curvula* | 3 |
| LC_coassembly_k141_383156 | 981 | Yes | Unknown | *Duplornaviricota* | *Chrymotiviricetes* | *Ghabrivirales* | *f.0283.base-Megabirna* | 1 |
| LC_coassembly_k141_398455 | 1206 | Yes | ssRNA(+) | *Lenarviricota* | *Howeltoviricetes* | *Cryppavirales* | *Mitoviridae* | 2 |
| LC_coassembly_k141_40647 | 2604 | Yes | Unknown | *Duplornaviricota* | *Chrymotiviricetes* | *Ghabrivirales* | *Nam-Co_family_65* | 2 |
| LC_coassembly_k141_415235 | 2334 | Yes | Unknown | *Pisuviricota* | *Pisoniviricetes* | *Picornavirales* | *f.0008.base-Polycipi* | 1 |
| LC_coassembly_k141_41569 | 1798 | Yes | Unknown | *Lenarviricota* | *c.0071.base-Howelto* | *o.0092* | *f.0388* | 1 |
| LC_coassembly_k141_420336 | 1107 | Yes | dsRNA | *Duplornaviricota* | *Resentoviricetes* | *Reovirales* | *Nam-Co_family_77* | 1 |
| LC_coassembly_k141_421224 | 617 | Yes | Unknown | *Pisuviricota* | *c.0013* | *o.0025.base-Sobeli* | *f.0076* | 1 |
| LC_coassembly_k141_431617 | 2888 | Yes | ssRNA(+) | *Lenarviricota* | *Amabiliviricetes* | *Wolframvirales* | *Nam-Co_family_05* | 1 |
| LC_coassembly_k141_455553 | 1147 | Yes | Unknown | *Negarnaviricota* | *Ellioviricetes* | *Bunyavirales* | *f.0328* | 1 |
| LC_coassembly_k141_473258 | 1405 | Yes | ssRNA(+) | *Pisuviricota* | *Pisoniviricetes* | *Picornavirales* | *Marnaviridae* | 1 |
| LC_coassembly_k141_483085 | 735 | Yes | Unknown | *Duplornaviricota* | *Chrymotiviricetes* | *Ghabrivirales* | *f.0283.base-Megabirna* | 1 |
| LC_coassembly_k141_488786 | 1207 | Yes | Unknown | *Kitrinoviricota* | *Alsuviricetes* | *Hepelivirales* | *f.0221.base-Alphatetra* | 1 |
| LC_coassembly_k141_497225 | 1784 | Yes | Unknown | *Pisuviricota* | *Duplopiviricetes* | *Durnavirales* | *f.0111* | 1 |
| LC_coassembly_k141_500165 | 1592 | Yes | ssRNA(+) | *Kitrinoviricota* | *Magsaviricetes* | *Nodamuvirales* | *Nodaviridae* | 1 |
| LC_coassembly_k141_502336 | 3100 | Yes | ssRNA(+) | *Lenarviricota* | *Amabiliviricetes* | *Wolframvirales* | *Narnaviridae* | 1 |
| LC_coassembly_k141_51051 | 2496 | Yes | Unknown | *Pisuviricota* | *c.0013* | *o.0025.base-Sobeli* | *f.0076* | 1 |
| LC_coassembly_k141_51246 | 8463 | Yes | Unknown | *Pisuviricota* | *Pisoniviricetes* | *Picornavirales* | *f.0025* | 1 |
| LC_coassembly_k141_515640 | 6206 | Yes | ssRNA(+) | *Kitrinoviricota* | *Alsuviricetes* | *o.0055* | *Endornaviridae* | 1 |
| LC_coassembly_k141_518944 | 1137 | Yes | dsRNA | *Duplornaviricota* | *Chrymotiviricetes* | *Ghabrivirales* | *Totiviridae* | 1 |
| LC_coassembly_k141_524640 | 2629 | Yes | Unknown | *Kitrinoviricota* | *Alsuviricetes* | *Hepelivirales* | *Nam-Co_family_49* | 2 |
| LC_coassembly_k141_526161 | 3463 | Yes | Unknown | *Pisuviricota* | *Duplopiviricetes* | *Durnavirales* | *f.0102.base-Curvula* | 3 |
| LC_coassembly_k141_537252 | 11543 | Yes | Unknown | *Kitrinoviricota* | *c.0034* | *o.0045* | *Nam-Co_family_57* | 9 |
| LC_coassembly_k141_567048 | 1364 | Yes | Unknown | *Pisuviricota* | *Pisoniviricetes* | *Picornavirales* | *f.0042.base-Marna* | 1 |
| LC_coassembly_k141_568262 | 2423 | Yes | Unknown | *Pisuviricota* | *c.0013* | *o.0025.base-Sobeli* | *f.0076* | 2 |
| LC_coassembly_k141_570098 | 1392 | Yes | Unknown | *Pisuviricota* | *Duplopiviricetes* | *Durnavirales* | *f.0102.base-Curvula* | 1 |
| LC_coassembly_k141_581270 | 6753 | Yes | Unknown | *Kitrinoviricota* | *c.0034* | *o.0045* | *f.0176* | 5 |
| LC_coassembly_k141_585258 | 646 | Yes | dsRNA | *Duplornaviricota* | *Chrymotiviricetes* | *Ghabrivirales* | *Totiviridae* | 1 |
| LC_coassembly_k141_589296 | 1694 | Yes | Unknown | *Pisuviricota* | *Duplopiviricetes* | *Durnavirales* | *f.0102.base-Curvula* | 1 |
| LC_coassembly_k141_592818 | 3278 | Yes | Unknown | *Pisuviricota* | *Duplopiviricetes* | *Durnavirales* | *f.0102.base-Curvula* | 3 |
| LC_coassembly_k141_601624 | 1833 | Yes | Unknown | *Pisuviricota* | *Duplopiviricetes* | *Durnavirales* | *f.0111* | 1 |
| LC_coassembly_k141_619132 | 2930 | Yes | Unknown | *Pisuviricota* | *Duplopiviricetes* | *Durnavirales* | *Nam-Co_family_40* | 2 |
| LC_coassembly_k141_629875 | 1871 | Yes | Unknown | *Pisuviricota* | *Duplopiviricetes* | *Durnavirales* | *f.0111* | 1 |
| LC_coassembly_k141_633441 | 818 | Yes | Unknown | *Pisuviricota* | *Duplopiviricetes* | *Durnavirales* | *Nam-Co_family_39* | 1 |
| LC_coassembly_k141_635520 | 3262 | Yes | ssRNA(+) | *Lenarviricota* | *Amabiliviricetes* | *Wolframvirales* | *Nam-Co_family_13* | 1 |
| LC_coassembly_k141_635711 | 904 | Yes | ssRNA(+) | *Lenarviricota* | *Leviviricetes* | *Timlovirales* | *Steitzviridae* | 1 |
| LC_coassembly_k141_642659 | 3730 | Yes | Unknown | *Duplornaviricota* | *Chrymotiviricetes* | *Ghabrivirales* | *f.0296* | 2 |
| LC_coassembly_k141_662643 | 5789 | Yes | Unknown | *Negarnaviricota* | *Ellioviricetes* | *Bunyavirales* | *f.0315* | 2 |
| LC_coassembly_k141_672667 | 1751 | Yes | Unknown | *Pisuviricota* | *Duplopiviricetes* | *Durnavirales* | *f.0111* | 1 |
| LC_coassembly_k141_682946 | 4642 | Yes | Unknown | *Pisuviricota* | *Pisoniviricetes* | *Picornavirales* | *f.0030* | 2 |
| LC_coassembly_k141_686723 | 1307 | Yes | Unknown | *Pisuviricota* | *Duplopiviricetes* | *Durnavirales* | *f.0111* | 1 |
| LC_coassembly_k141_689124 | 1822 | Yes | dsRNA | *Pisuviricota* | *Duplopiviricetes* | *Durnavirales* | *Partitiviridae* | 2 |
| LC_coassembly_k141_690501 | 767 | Yes | ssRNA(+) | *Pisuviricota* | *Pisoniviricetes* | *Picornavirales* | *Dicistroviridae* | 1 |
| LC_coassembly_k141_698489 | 1893 | Yes | Unknown | *Duplornaviricota* | *Chrymotiviricetes* | *Ghabrivirales* | *f.0287* | 1 |
| LC_coassembly_k141_700243 | 1180 | Yes | Unknown | *Pisuviricota* | *Duplopiviricetes* | *Durnavirales* | *f.0102.base-Curvula* | 1 |
| LC_coassembly_k141_709398 | 9456 | Yes | Unknown | *Duplornaviricota* | *Chrymotiviricetes* | *Ghabrivirales* | *Nam-Co_family_68* | 1 |
| LC_coassembly_k141_70982 | 642 | Yes | dsRNA | *Duplornaviricota* | *Resentoviricetes* | *Reovirales* | *Reoviridae* | 1 |
| LC_coassembly_k141_709835 | 2820 | Yes | ssRNA(+) | *Lenarviricota* | *Howeltoviricetes* | *Cryppavirales* | *Mitoviridae* | 1 |
| LC_coassembly_k141_715064 | 978 | Yes | Unknown | *Duplornaviricota* | *Chrymotiviricetes* | *Ghabrivirales* | *f.0296* | 1 |
| LC_coassembly_k141_716515 | 7700 | Yes | Unknown | *Negarnaviricota* | *Ellioviricetes* | *Bunyavirales* | *f.0328* | 1 |
| LC_coassembly_k141_721053 | 5986 | Yes | Unknown | *Duplornaviricota* | *Chrymotiviricetes* | *Ghabrivirales* | *Nam-Co_family_62* | 2 |
| LC_coassembly_k141_727479 | 3625 | Yes | Unknown | *Pisuviricota* | *Duplopiviricetes* | *Durnavirales* | *f.0102.base-Curvula* | 2 |
| LC_coassembly_k141_732085 | 3613 | Yes | Unknown | *Negarnaviricota* | *Ellioviricetes* | *Bunyavirales* | *f.0327* | 1 |
| LC_coassembly_k141_737588 | 955 | Yes | Unknown | *Pisuviricota* | *Pisoniviricetes* | *Picornavirales* | *f.0029.base-Dicistro* | 1 |
| LC_coassembly_k141_744534 | 1110 | Yes | Unknown | *Pisuviricota* | *Duplopiviricetes* | *Durnavirales* | *f.0102.base-Curvula* | 1 |
| LC_coassembly_k141_75697 | 8592 | Yes | Unknown | *Kitrinoviricota* | *c.0034* | *o.0045* | *f.0176* | 6 |
| LC_coassembly_k141_757367 | 2161 | Yes | ssRNA(+) | *Kitrinoviricota* | *Tolucaviricetes* | *Tolivirales* | *Tombusviridae* | 2 |
| LC_coassembly_k141_76639 | 4744 | No | NA | *NA* | *NA* | *NA* | *Nam-Co_family_04* | 2 |
| LC_coassembly_k141_769811 | 3027 | Yes | Unknown | *Negarnaviricota* | *Ellioviricetes* | *Bunyavirales* | *f.0315* | 1 |
| LC_coassembly_k141_76989 | 3728 | Yes | ssRNA(+) | *Kitrinoviricota* | *Tolucaviricetes* | *Tolivirales* | *Tombusviridae* | 3 |
| LC_coassembly_k141_774642 | 1043 | Yes | Unknown | *Duplornaviricota* | *Chrymotiviricetes* | *Ghabrivirales* | *Nam-Co_family_67* | 1 |
| LC_coassembly_k141_775494 | 2851 | Yes | ssRNA(+) | *Lenarviricota* | *Amabiliviricetes* | *Wolframvirales* | *Narnaviridae* | 2 |
| LC_coassembly_k141_776867 | 2546 | Yes | Unknown | *Kitrinoviricota* | *c.0031* | *o.0042* | *Nam-Co_family_53* | 2 |
| LC_coassembly_k141_779785 | 4707 | Yes | Unknown | *Kitrinoviricota* | *c.0025* | *o.0036* | *f.0145* | 3 |
| LC_coassembly_k141_781199 | 856 | Yes | ssRNA(+) | *Lenarviricota* | *Amabiliviricetes* | *Wolframvirales* | *Narnaviridae* | 1 |
| LC_coassembly_k141_787612 | 2502 | Yes | ssRNA(+) | *Lenarviricota* | *Howeltoviricetes* | *Cryppavirales* | *Mitoviridae* | 2 |
| LC_coassembly_k141_788326 | 2077 | No | NA | *NA* | *NA* | *NA* | *Nam-Co_family_25* | 2 |
| LC_coassembly_k141_801488 | 8050 | Yes | Unknown | *Duplornaviricota* | *Chrymotiviricetes* | *Ghabrivirales* | *Nam-Co_family_63* | 2 |
| LC_coassembly_k141_843059 | 7504 | Yes | Unknown | *Kitrinoviricota* | *c.0034* | *o.0045* | *f.0176* | 3 |
| LC_coassembly_k141_860293 | 3094 | Yes | Unknown | *Lenarviricota* | *c.0071.base-Howelto* | *o.0092* | *f.0388* | 3 |
| LC_coassembly_k141_864888 | 753 | Yes | Unknown | *Lenarviricota* | *Leviviricetes* | *Norzivirales* | *f.0363* | 1 |
| LC_coassembly_k141_868598 | 1747 | Yes | dsRNA | *Pisuviricota* | *Duplopiviricetes* | *Durnavirales* | *Picobirnaviridae* | 1 |
| LC_coassembly_k141_86982 | 4216 | Yes | ssRNA(+) | *Pisuviricota* | *c.0012* | *Sobelivirales* | *Solemoviridae* | 3 |
| LC_coassembly_k141_893532 | 932 | Yes | Unknown | *Pisuviricota* | *Duplopiviricetes* | *Durnavirales* | *Nam-Co_family_38* | 1 |
| LC_coassembly_k141_908918 | 621 | Yes | Unknown | *Duplornaviricota* | *Chrymotiviricetes* | *Ghabrivirales* | *f.0283.base-Megabirna* | 1 |
| LC_coassembly_k141_914305 | 692 | Yes | Unknown | *Duplornaviricota* | *Chrymotiviricetes* | *Ghabrivirales* | *f.0296* | 1 |
| LC_coassembly_k141_916411 | 4822 | Yes | ssRNA(+) | *Pisuviricota* | *Pisoniviricetes* | *Picornavirales* | *Marnaviridae* | 1 |
| LC_coassembly_k141_943468 | 661 | Yes | Unknown | *Pisuviricota* | *Duplopiviricetes* | *Durnavirales* | *f.0102.base-Curvula* | 1 |
| LC_coassembly_k141_945938 | 5247 | Yes | Unknown | *Kitrinoviricota* | *c.0025* | *o.0036* | *Nam-Co_family_51* | 3 |
| LC_coassembly_k141_947088 | 3283 | Yes | Unknown | *Pisuviricota* | *Duplopiviricetes* | *Durnavirales* | *f.0102.base-Curvula* | 3 |
| LC_coassembly_k141_952072 | 944 | Yes | ssRNA(+) | *Kitrinoviricota* | *Tolucaviricetes* | *Tolivirales* | *Tombusviridae* | 1 |
| LC_coassembly_k141_961667 | 2332 | Yes | ssRNA(+) | *Lenarviricota* | *Howeltoviricetes* | *Cryppavirales* | *Mitoviridae* | 1 |
| LC_coassembly_k141_964946 | 1496 | Yes | Unknown | *Pisuviricota* | *c.0013* | *o.0025.base-Sobeli* | *f.0076* | 1 |
| LC_coassembly_k141_969841 | 4338 | Yes | Unknown | *Duplornaviricota* | *Chrymotiviricetes* | *Ghabrivirales* | *f.0296* | 2 |
| LC_coassembly_k141_972273 | 964 | Yes | Unknown | *Duplornaviricota* | *Chrymotiviricetes* | *Ghabrivirales* | *Nam-Co_family_69* | 1 |
| LC_coassembly_k141_97255 | 3137 | Yes | Unknown | *Duplornaviricota* | *Chrymotiviricetes* | *Ghabrivirales* | *f.0286* | 1 |
| LC_coassembly_k141_977866 | 664 | Yes | Unknown | *Duplornaviricota* | *Chrymotiviricetes* | *Ghabrivirales* | *f.0292* | 1 |
| LC_coassembly_k141_995284 | 1797 | Yes | Unknown | *Pisuviricota* | *Duplopiviricetes* | *Durnavirales* | *f.0111* | 1 |
| LS_0.5m_0.2um_k141_101583 | 2614 | Yes | Unknown | *Pisuviricota* | *c.0013* | *o.0025.base-Sobeli* | *f.0076* | 1 |
| LS_0.5m_0.2um_k141_120401 | 617 | Yes | ssRNA(+) | *Lenarviricota* | *Howeltoviricetes* | *Cryppavirales* | *Mitoviridae* | 1 |
| LS_0.5m_0.2um_k141_127309 | 3659 | Yes | ssRNA(+) | *Kitrinoviricota* | *Magsaviricetes* | *Nodamuvirales* | *Nodaviridae* | 2 |
| LS_0.5m_0.2um_k141_12995 | 2018 | Yes | ssRNA(+) | *Lenarviricota* | *Miaviricetes* | *Ourlivirales* | *Botourmiaviridae* | 1 |
| LS_0.5m_0.2um_k141_13024 | 1384 | Yes | ssRNA(+) | *Lenarviricota* | *Amabiliviricetes* | *Wolframvirales* | *Narnaviridae* | 2 |
| LS_0.5m_0.2um_k141_167028 | 1256 | Yes | Unknown | *Kitrinoviricota* | *c.0034* | *o.0045* | *f.0181* | 1 |
| LS_0.5m_0.2um_k141_180478 | 2016 | Yes | ssRNA(+) | *Lenarviricota* | *Leviviricetes* | *Norzivirales* | *Solspiviridae* | 2 |
| LS_0.5m_0.2um_k141_187456 | 723 | Yes | Unknown | *Negarnaviricota* | *c.0054.base-Yunchang* | *o.0072* | *f.0354* | 1 |
| LS_0.5m_0.2um_k141_202953 | 776 | Yes | ssRNA(+) | *Lenarviricota* | *Leviviricetes* | *Norzivirales* | *Fiersviridae* | 1 |
| LS_0.5m_0.2um_k141_211660 | 796 | No | NA | *NA* | *NA* | *NA* | *NA* | 1 |
| LS_0.5m_0.2um_k141_219883 | 852 | Yes | Unknown | *Lenarviricota* | *c.0069.base-Howelto* | *o.0090* | *f.0386* | 1 |
| LS_0.5m_0.2um_k141_227368 | 626 | Yes | ssRNA(+) | *Lenarviricota* | *Miaviricetes* | *Ourlivirales* | *Botourmiaviridae* | 1 |
| LS_0.5m_0.2um_k141_234728 | 2489 | Yes | ssRNA(+) | *Lenarviricota* | *Amabiliviricetes* | *Wolframvirales* | *Narnaviridae* | 1 |
| LS_0.5m_0.2um_k141_234956 | 611 | Yes | dsRNA | *Duplornaviricota* | *Resentoviricetes* | *Reovirales* | *Reoviridae* | 1 |
| LS_0.5m_0.2um_k141_272854 | 1701 | Yes | Unknown | *Pisuviricota* | *c.0013* | *o.0025.base-Sobeli* | *f.0076* | 1 |
| LS_0.5m_0.2um_k141_275484 | 935 | Yes | Unknown | *Pisuviricota* | *c.0013* | *o.0025.base-Sobeli* | *f.0076* | 1 |
| LS_0.5m_0.2um_k141_289194 | 686 | Yes | Unknown | *Lenarviricota* | *c.0069.base-Howelto* | *o.0090* | *f.0386* | 1 |
| LS_0.5m_0.2um_k141_296402 | 1162 | Yes | ssRNA(+) | *Lenarviricota* | *Miaviricetes* | *Ourlivirales* | *Botourmiaviridae* | 1 |
| LS_0.5m_0.2um_k141_30045 | 2438 | Yes | Unknown | *Kitrinoviricota* | *c.0034* | *o.0045* | *f.0176* | 3 |
| LS_0.5m_0.2um_k141_312953 | 3334 | Yes | ssRNA(+) | *Pisuviricota* | *Pisoniviricetes* | *Picornavirales* | *Dicistroviridae* | 2 |
| LS_0.5m_0.2um_k141_344707 | 1321 | Yes | Unknown | *Pisuviricota* | *c.0013* | *o.0025.base-Sobeli* | *f.0076* | 2 |
| LS_0.5m_0.2um_k141_392045 | 732 | Yes | Unknown | *Pisuviricota* | *c.0013* | *o.0025.base-Sobeli* | *f.0076* | 1 |
| LS_0.5m_0.2um_k141_42719 | 673 | Yes | Unknown | *Kitrinoviricota* | *c.0034* | *o.0045* | *f.0176* | 1 |
| LS_0.5m_0.2um_k141_46292 | 4887 | Yes | ssRNA(+) | *Kitrinoviricota* | *Tolucaviricetes* | *Tolivirales* | *Tombusviridae* | 4 |
| LS_0.5m_0.2um_k141_84063 | 2504 | Yes | ssRNA(+) | *Lenarviricota* | *Amabiliviricetes* | *Wolframvirales* | *Narnaviridae* | 1 |
| LS_0.5m_0.2um_k141_86899 | 1171 | No | NA | *NA* | *NA* | *NA* | *NA* | 1 |
| LS_0.5m_0.2um_k141_97488 | 857 | Yes | Unknown | *Lenarviricota* | *Leviviricetes* | *Norzivirales* | *f.0361.base-Solspi* | 2 |
| LS_0.5m_1.2um_k141_104181 | 1598 | Yes | ssRNA(+) | *Lenarviricota* | *Amabiliviricetes* | *Wolframvirales* | *Nam-Co_family_09* | 1 |
| LS_0.5m_1.2um_k141_105741 | 1006 | Yes | ssRNA(+) | *Lenarviricota* | *Howeltoviricetes* | *Cryppavirales* | *Mitoviridae* | 1 |
| LS_0.5m_1.2um_k141_110339 | 3407 | Yes | ssRNA(+) | *Lenarviricota* | *Miaviricetes* | *Ourlivirales* | *Botourmiaviridae* | 1 |
| LS_0.5m_1.2um_k141_112240 | 2489 | Yes | Unknown | *Pisuviricota* | *c.0013* | *o.0025.base-Sobeli* | *f.0076* | 1 |
| LS_0.5m_1.2um_k141_152394 | 1128 | Yes | Unknown | *Negarnaviricota* | *Ellioviricetes* | *Bunyavirales* | *f.0328* | 1 |
| LS_0.5m_1.2um_k141_16205 | 2060 | Yes | ssRNA(+) | *Lenarviricota* | *Leviviricetes* | *Timlovirales* | *Steitzviridae* | 2 |
| LS_0.5m_1.2um_k141_16215 | 687 | Yes | Unknown | *Kitrinoviricota* | *c.0025* | *o.0036* | *f.0145* | 1 |
| LS_0.5m_1.2um_k141_174384 | 667 | Yes | ssRNA(+) | *Pisuviricota* | *Pisoniviricetes* | *Picornavirales* | *Marnaviridae* | 1 |
| LS_0.5m_1.2um_k141_197594 | 704 | Yes | ssRNA(+) | *Lenarviricota* | *Leviviricetes* | *Norzivirales* | *Fiersviridae* | 1 |
| LS_0.5m_1.2um_k141_203498 | 1421 | Yes | ssRNA(+) | *Lenarviricota* | *Amabiliviricetes* | *Wolframvirales* | *Narnaviridae* | 1 |
| LS_0.5m_1.2um_k141_213974 | 5125 | Yes | ssRNA(+) | *Kitrinoviricota* | *Tolucaviricetes* | *Tolivirales* | *Tombusviridae* | 7 |
| LS_0.5m_1.2um_k141_221766 | 1000 | Yes | Unknown | *Lenarviricota* | *Leviviricetes* | *Norzivirales* | *f.0361.base-Solspi* | 1 |
| LS_0.5m_1.2um_k141_232702 | 3954 | Yes | Unknown | *Kitrinoviricota* | *c.0032* | *o.0043* | *f.0167* | 2 |
| LS_0.5m_1.2um_k141_244393 | 738 | Yes | ssRNA(+) | *Kitrinoviricota* | *Tolucaviricetes* | *Tolivirales* | *Tombusviridae* | 1 |
| LS_0.5m_1.2um_k141_249566 | 1249 | Yes | dsRNA | *Pisuviricota* | *Duplopiviricetes* | *Durnavirales* | *Partitiviridae* | 1 |
| LS_0.5m_1.2um_k141_285352 | 962 | Yes | ssRNA(+) | *Lenarviricota* | *Howeltoviricetes* | *Cryppavirales* | *Mitoviridae* | 1 |
| LS_0.5m_1.2um_k141_293620 | 708 | Yes | ssRNA(+) | *Pisuviricota* | *Pisoniviricetes* | *Picornavirales* | *Dicistroviridae* | 1 |
| LS_0.5m_1.2um_k141_296991 | 1222 | Yes | Unknown | *Negarnaviricota* | *c.0054.base-Yunchang* | *o.0072* | *f.0354* | 1 |
| LS_0.5m_1.2um_k141_297325 | 4578 | Yes | ssRNA(+) | *Pisuviricota* | *Pisoniviricetes* | *Picornavirales* | *Polycipiviridae* | 2 |
| LS_0.5m_1.2um_k141_302852 | 4142 | Yes | Unknown | *Kitrinoviricota* | *c.0024* | *o.0035* | *f.0144* | 2 |
| LS_0.5m_1.2um_k141_304513 | 2217 | Yes | Unknown | *Kitrinoviricota* | *c.0024* | *o.0035* | *f.0144* | 2 |
| LS_0.5m_1.2um_k141_315047 | 3920 | Yes | Unknown | *Kitrinoviricota* | *c.0034* | *o.0045* | *f.0176* | 3 |
| LS_0.5m_1.2um_k141_324076 | 1470 | Yes | ssRNA(+) | *Lenarviricota* | *Amabiliviricetes* | *Wolframvirales* | *Narnaviridae* | 1 |
| LS_0.5m_1.2um_k141_335154 | 849 | Yes | Unknown | *Kitrinoviricota* | *c.0034* | *o.0045* | *f.0176* | 1 |
| LS_0.5m_1.2um_k141_340418 | 1040 | Yes | Unknown | *Lenarviricota* | *c.0067.base-Howelto* | *o.0088* | *f.0384* | 1 |
| LS_0.5m_1.2um_k141_346045 | 2450 | Yes | ssRNA(+) | *Lenarviricota* | *Leviviricetes* | *Timlovirales* | *Steitzviridae* | 2 |
| LS_0.5m_1.2um_k141_350481 | 2036 | Yes | ssRNA(+) | *Lenarviricota* | *Howeltoviricetes* | *Cryppavirales* | *Mitoviridae* | 1 |
| LS_0.5m_1.2um_k141_380385 | 863 | No | NA | *NA* | *NA* | *NA* | *NA* | 1 |
| LS_0.5m_1.2um_k141_395063 | 838 | Yes | ssRNA(+) | *Lenarviricota* | *Howeltoviricetes* | *Cryppavirales* | *Mitoviridae* | 1 |
| LS_0.5m_1.2um_k141_441494 | 3395 | No | NA | *NA* | *NA* | *NA* | *NA* | 1 |
| LS_0.5m_1.2um_k141_450204 | 812 | Yes | ssRNA(+) | *Lenarviricota* | *Amabiliviricetes* | *Wolframvirales* | *Narnaviridae* | 1 |
| LS_0.5m_1.2um_k141_450766 | 998 | Yes | ssRNA(+) | *Lenarviricota* | *Amabiliviricetes* | *Wolframvirales* | *Narnaviridae* | 1 |
| LS_0.5m_1.2um_k141_463252 | 10091 | Yes | ssRNA(+) | *Kitrinoviricota* | *Alsuviricetes* | *Hepelivirales* | *Benyviridae* | 6 |
| LS_0.5m_1.2um_k141_487950 | 670 | Yes | Unknown | *Negarnaviricota* | *Ellioviricetes* | *Bunyavirales* | *f.0327* | 1 |
| LS_0.5m_1.2um_k141_519525 | 803 | Yes | Unknown | *Negarnaviricota* | *Ellioviricetes* | *Bunyavirales* | *f.0328* | 1 |
| LS_0.5m_1.2um_k141_563554 | 741 | Yes | ssRNA(+) | *Lenarviricota* | *Amabiliviricetes* | *Wolframvirales* | *Nam-Co_family_13* | 1 |
| LS_0.5m_1.2um_k141_567195 | 1036 | Yes | ssRNA(+) | *Lenarviricota* | *Amabiliviricetes* | *Wolframvirales* | *Narnaviridae* | 1 |
| LS_0.5m_1.2um_k141_585706 | 3450 | Yes | ssRNA(+) | *Lenarviricota* | *Leviviricetes* | *Norzivirales* | *Fiersviridae* | 3 |
| LS_0.5m_1.2um_k141_606453 | 4984 | Yes | ssRNA(+) | *Pisuviricota* | *Pisoniviricetes* | *Picornavirales* | *Dicistroviridae* | 1 |
| LS_0.5m_1.2um_k141_623158 | 1682 | Yes | Unknown | *Lenarviricota* | *Leviviricetes* | *Norzivirales* | *f.0361.base-Solspi* | 1 |
| LS_0.5m_1.2um_k141_627694 | 703 | Yes | ssRNA(+) | *Lenarviricota* | *Leviviricetes* | *Norzivirales* | *Fiersviridae* | 1 |
| LS_0.5m_1.2um_k141_663049 | 1198 | Yes | ssRNA(+) | *Lenarviricota* | *Miaviricetes* | *Ourlivirales* | *Botourmiaviridae* | 1 |
| LS_0.5m_1.2um_k141_670678 | 843 | Yes | Unknown | *Duplornaviricota* | *Chrymotiviricetes* | *Ghabrivirales* | *f.0287* | 1 |
| LS_0.5m_1.2um_k141_674011 | 923 | Yes | dsRNA | *Pisuviricota* | *c.0012* | *Sobelivirales* | *Barnaviridae* | 1 |
| LS_0.5m_1.2um_k141_706656 | 765 | Yes | ssRNA(+) | *Lenarviricota* | *Amabiliviricetes* | *Wolframvirales* | *Narnaviridae* | 1 |
| LS_0.5m_1.2um_k141_7672 | 1122 | Yes | ssRNA(+) | *Lenarviricota* | *Amabiliviricetes* | *Wolframvirales* | *Narnaviridae* | 1 |
| LS_0.5m_1.2um_k141_89262 | 1413 | Yes | ssRNA(+) | *Lenarviricota* | *Amabiliviricetes* | *Wolframvirales* | *Narnaviridae* | 1 |
| LS_0.5m_1.2um_k141_96485 | 605 | Yes | Unknown | *Lenarviricota* | *Leviviricetes* | *Norzivirales* | *f.0361.base-Solspi* | 1 |
| LS_0.5m_10um_k141_105168 | 712 | Yes | ssRNA(+) | *Lenarviricota* | *Howeltoviricetes* | *Cryppavirales* | *Mitoviridae* | 1 |
| LS_0.5m_10um_k141_124284 | 3292 | Yes | ssRNA(+) | *Kitrinoviricota* | *Alsuviricetes* | *Hepelivirales* | *Benyviridae* | 1 |
| LS_0.5m_10um_k141_12963 | 4468 | Yes | ssRNA(+) | *Pisuviricota* | *c.0012* | *Sobelivirales* | *Solemoviridae* | 4 |
| LS_0.5m_10um_k141_131043 | 625 | Yes | dsRNA | *Duplornaviricota* | *Resentoviricetes* | *Reovirales* | *Reoviridae* | 1 |
| LS_0.5m_10um_k141_134516 | 958 | Yes | ssRNA(+) | *Lenarviricota* | *Howeltoviricetes* | *Cryppavirales* | *Mitoviridae* | 1 |
| LS_0.5m_10um_k141_136572 | 3626 | No | NA | *NA* | *NA* | *NA* | *NA* | 1 |
| LS_0.5m_10um_k141_137938 | 995 | Yes | Unknown | *Pisuviricota* | *Duplopiviricetes* | *Durnavirales* | *f.0102.base-Curvula* | 1 |
| LS_0.5m_10um_k141_145047 | 2218 | No | NA | *NA* | *NA* | *NA* | *NA* | 1 |
| LS_0.5m_10um_k141_163294 | 673 | Yes | dsRNA | *Duplornaviricota* | *Resentoviricetes* | *Reovirales* | *Reoviridae* | 1 |
| LS_0.5m_10um_k141_165019 | 904 | Yes | ssRNA(+) | *Lenarviricota* | *Miaviricetes* | *Ourlivirales* | *Botourmiaviridae* | 1 |
| LS_0.5m_10um_k141_171521 | 1575 | Yes | ssRNA(+) | *Lenarviricota* | *Howeltoviricetes* | *Cryppavirales* | *Mitoviridae* | 1 |
| LS_0.5m_10um_k141_1769 | 1106 | Yes | Unknown | *Pisuviricota* | *Duplopiviricetes* | *Durnavirales* | *f.0111* | 2 |
| LS_0.5m_10um_k141_184862 | 781 | Yes | Unknown | *Duplornaviricota* | *Chrymotiviricetes* | *Ghabrivirales* | *f.0296* | 1 |
| LS_0.5m_10um_k141_196336 | 1542 | Yes | ssRNA(-) | *Negarnaviricota* | *Monjiviricetes* | *Mononegavirales* | *Rhabdoviridae* | 1 |
| LS_0.5m_10um_k141_209189 | 1253 | Yes | Unknown | *Pisuviricota* | *Duplopiviricetes* | *Durnavirales* | *f.0102.base-Curvula* | 1 |
| LS_0.5m_10um_k141_212226 | 5239 | Yes | ssRNA(+) | *Lenarviricota* | *Miaviricetes* | *Ourlivirales* | *Botourmiaviridae* | 2 |
| LS_0.5m_10um_k141_23405 | 888 | Yes | ssRNA(-) | *Negarnaviricota* | *Monjiviricetes* | *Mononegavirales* | *Rhabdoviridae* | 1 |
| LS_0.5m_10um_k141_3393 | 676 | Yes | ssRNA(+) | *Lenarviricota* | *Amabiliviricetes* | *Wolframvirales* | *Narnaviridae* | 1 |
| LS_0.5m_10um_k141_46079 | 2099 | Yes | Unknown | *Pisuviricota* | *Duplopiviricetes* | *Durnavirales* | *f.0102.base-Curvula* | 4 |
| LS_0.5m_10um_k141_53807 | 1383 | Yes | Unknown | *Pisuviricota* | *Duplopiviricetes* | *Durnavirales* | *f.0102.base-Curvula* | 1 |
| LS_0.5m_10um_k141_78851 | 9085 | Yes | ssRNA(+) | *Pisuviricota* | *Pisoniviricetes* | *Picornavirales* | *Marnaviridae* | 2 |
| LS_0.5m_10um_k141_84898 | 836 | Yes | Unknown | *Duplornaviricota* | *Chrymotiviricetes* | *Ghabrivirales* | *f.0296* | 1 |
| LS_0.5m_10um_k141_96668 | 675 | Yes | Unknown | *Pisuviricota* | *Duplopiviricetes* | *Durnavirales* | *Nam-Co_family_34* | 1 |
| LS_0.5m_10um_k141_96805 | 689 | Yes | ssRNA(+) | *Lenarviricota* | *Miaviricetes* | *Ourlivirales* | *Botourmiaviridae* | 1 |
| LS_0.5m_3um_k141_101718 | 784 | Yes | Unknown | *Duplornaviricota* | *Chrymotiviricetes* | *Ghabrivirales* | *f.0286* | 1 |
| LS_0.5m_3um_k141_122486 | 1508 | Yes | Unknown | *Lenarviricota* | *Leviviricetes* | *Norzivirales* | *f.0368.base-Atkins* | 1 |
| LS_0.5m_3um_k141_155241 | 1465 | Yes | Unknown | *Lenarviricota* | *c.0069.base-Howelto* | *o.0090* | *f.0386* | 1 |
| LS_0.5m_3um_k141_156627 | 788 | Yes | ssRNA(+) | *Lenarviricota* | *Amabiliviricetes* | *Wolframvirales* | *Narnaviridae* | 1 |
| LS_0.5m_3um_k141_157301 | 1596 | Yes | ssRNA(+) | *Lenarviricota* | *Howeltoviricetes* | *Cryppavirales* | *Mitoviridae* | 2 |
| LS_0.5m_3um_k141_161033 | 1005 | Yes | Unknown | *Pisuviricota* | *Duplopiviricetes* | *Durnavirales* | *f.0111* | 1 |
| LS_0.5m_3um_k141_161223 | 1448 | Yes | ssRNA(+) | *Lenarviricota* | *Howeltoviricetes* | *Cryppavirales* | *Mitoviridae* | 2 |
| LS_0.5m_3um_k141_164826 | 1293 | Yes | Unknown | *Pisuviricota* | *c.0013* | *o.0025.base-Sobeli* | *f.0076* | 1 |
| LS_0.5m_3um_k141_164912 | 874 | Yes | ssRNA(+) | *Pisuviricota* | *Pisoniviricetes* | *Picornavirales* | *Marnaviridae* | 1 |
| LS_0.5m_3um_k141_178749 | 1007 | Yes | ssRNA(+) | *Pisuviricota* | *Pisoniviricetes* | *Picornavirales* | *Dicistroviridae* | 1 |
| LS_0.5m_3um_k141_179493 | 933 | Yes | Unknown | *Duplornaviricota* | *Chrymotiviricetes* | *Ghabrivirales* | *f.0287* | 1 |
| LS_0.5m_3um_k141_188545 | 956 | Yes | Unknown | *Duplornaviricota* | *Chrymotiviricetes* | *Ghabrivirales* | *f.0287* | 1 |
| LS_0.5m_3um_k141_209056 | 1535 | Yes | Unknown | *Duplornaviricota* | *Chrymotiviricetes* | *Ghabrivirales* | *f.0287* | 1 |
| LS_0.5m_3um_k141_209520 | 1442 | Yes | Unknown | *Pisuviricota* | *c.0013* | *o.0025.base-Sobeli* | *f.0076* | 2 |
| LS_0.5m_3um_k141_238771 | 1677 | Yes | Unknown | *Pisuviricota* | *Duplopiviricetes* | *Durnavirales* | *f.0111* | 1 |
| LS_0.5m_3um_k141_246611 | 651 | Yes | dsRNA | *Pisuviricota* | *Duplopiviricetes* | *Durnavirales* | *Partitiviridae* | 1 |
| LS_0.5m_3um_k141_264243 | 621 | Yes | Unknown | *Lenarviricota* | *c.0069.base-Howelto* | *o.0090* | *f.0386* | 1 |
| LS_0.5m_3um_k141_270374 | 995 | Yes | Unknown | *Pisuviricota* | *Duplopiviricetes* | *Durnavirales* | *f.0111* | 1 |
| LS_0.5m_3um_k141_281258 | 2292 | Yes | Unknown | *Pisuviricota* | *c.0013* | *o.0025.base-Sobeli* | *f.0076* | 1 |
| LS_0.5m_3um_k141_29737 | 1318 | Yes | ssRNA(+) | *Pisuviricota* | *Pisoniviricetes* | *Picornavirales* | *Marnaviridae* | 1 |
| LS_0.5m_3um_k141_311742 | 1498 | Yes | Unknown | *Kitrinoviricota* | *c.0034* | *o.0045* | *f.0176* | 1 |
| LS_0.5m_3um_k141_320330 | 2504 | Yes | Unknown | *Pisuviricota* | *Duplopiviricetes* | *Durnavirales* | *f.0102.base-Curvula* | 2 |
| LS_0.5m_3um_k141_337163 | 1393 | Yes | ssRNA(+) | *Lenarviricota* | *Amabiliviricetes* | *Wolframvirales* | *Narnaviridae* | 1 |
| LS_0.5m_3um_k141_342628 | 1119 | Yes | ssRNA(+) | *Pisuviricota* | *Pisoniviricetes* | *Picornavirales* | *Polycipiviridae* | 1 |
| LS_0.5m_3um_k141_347973 | 2289 | Yes | Unknown | *Kitrinoviricota* | *c.0031* | *o.0042* | *f.0156* | 2 |
| LS_0.5m_3um_k141_350269 | 624 | Yes | Unknown | *Duplornaviricota* | *Chrymotiviricetes* | *Ghabrivirales* | *f.0286* | 1 |
| LS_0.5m_3um_k141_360239 | 947 | Yes | ssRNA(+) | *Lenarviricota* | *Leviviricetes* | *Norzivirales* | *Fiersviridae* | 1 |
| LS_0.5m_3um_k141_360401 | 1159 | Yes | ssRNA(+) | *Lenarviricota* | *Miaviricetes* | *Ourlivirales* | *Botourmiaviridae* | 1 |
| LS_0.5m_3um_k141_37077 | 612 | Yes | Unknown | *Lenarviricota* | *Leviviricetes* | *Norzivirales* | *f.0367.base-Atkins* | 1 |
| LS_0.5m_3um_k141_388117 | 10204 | Yes | ssRNA(+) | *Kitrinoviricota* | *Alsuviricetes* | *Hepelivirales* | *Benyviridae* | 4 |
| LS_0.5m_3um_k141_393240 | 3475 | Yes | Unknown | *Duplornaviricota* | *Chrymotiviricetes* | *Ghabrivirales* | *f.0287* | 1 |
| LS_0.5m_3um_k141_409801 | 1840 | Yes | ssRNA(+) | *Lenarviricota* | *Amabiliviricetes* | *Wolframvirales* | *Narnaviridae* | 1 |
| LS_0.5m_3um_k141_412695 | 2188 | Yes | ssRNA(+) | *Kitrinoviricota* | *Tolucaviricetes* | *Tolivirales* | *Tombusviridae* | 2 |
| LS_0.5m_3um_k141_415219 | 1960 | Yes | Unknown | *Kitrinoviricota* | *c.0034* | *o.0045* | *f.0176* | 3 |
| LS_0.5m_3um_k141_430588 | 786 | Yes | ssRNA(+) | *Pisuviricota* | *Pisoniviricetes* | *Picornavirales* | *Marnaviridae* | 1 |
| LS_0.5m_3um_k141_448451 | 779 | Yes | Unknown | *Lenarviricota* | *c.0068.base-Howelto* | *o.0089* | *f.0385* | 1 |
| LS_0.5m_3um_k141_63322 | 1604 | No | NA | *NA* | *NA* | *NA* | *NA* | 1 |
| LS_0.5m_3um_k141_76062 | 653 | Yes | ssRNA(+) | *Lenarviricota* | *Miaviricetes* | *Ourlivirales* | *Botourmiaviridae* | 1 |
| LS_0.5m_3um_k141_94350 | 1350 | Yes | Unknown | *Pisuviricota* | *Duplopiviricetes* | *Durnavirales* | *f.0111* | 1 |
| LS_coassembly_k141_1002104 | 2439 | Yes | Unknown | *Pisuviricota* | *c.0013* | *o.0025.base-Sobeli* | *f.0076* | 1 |
| LS_coassembly_k141_1002961 | 2008 | Yes | Unknown | *Pisuviricota* | *Duplopiviricetes* | *Durnavirales* | *f.0102.base-Curvula* | 2 |
| LS_coassembly_k141_1013649 | 2365 | Yes | ssRNA(+) | *Lenarviricota* | *Miaviricetes* | *Ourlivirales* | *Botourmiaviridae* | 2 |
| LS_coassembly_k141_1016243 | 1204 | Yes | Unknown | *Lenarviricota* | *c.0069.base-Howelto* | *o.0090* | *f.0386* | 1 |
| LS_coassembly_k141_1023795 | 2506 | Yes | ssRNA(+) | *Lenarviricota* | *Amabiliviricetes* | *Wolframvirales* | *Narnaviridae* | 1 |
| LS_coassembly_k141_1028965 | 2893 | Yes | ssRNA(+) | *Kitrinoviricota* | *Tolucaviricetes* | *Tolivirales* | *Tombusviridae* | 4 |
| LS_coassembly_k141_1029418 | 4411 | Yes | Unknown | *Kitrinoviricota* | *c.0024* | *o.0035* | *f.0144* | 2 |
| LS_coassembly_k141_1030339 | 2337 | Yes | ssRNA(+) | *Pisuviricota* | *Pisoniviricetes* | *Picornavirales* | *Marnaviridae* | 1 |
| LS_coassembly_k141_1032564 | 814 | Yes | ssRNA(+) | *Lenarviricota* | *Amabiliviricetes* | *Wolframvirales* | *Narnaviridae* | 1 |
| LS_coassembly_k141_1033411 | 9918 | Yes | ssRNA(+) | *Kitrinoviricota* | *Alsuviricetes* | *Hepelivirales* | *Benyviridae* | 4 |
| LS_coassembly_k141_1046808 | 1555 | Yes | Unknown | *Lenarviricota* | *c.0068.base-Howelto* | *o.0089* | *f.0385* | 1 |
| LS_coassembly_k141_1048877 | 1352 | Yes | Unknown | *Duplornaviricota* | *Chrymotiviricetes* | *Ghabrivirales* | *f.0296* | 1 |
| LS_coassembly_k141_1052634 | 786 | Yes | ssRNA(+) | *Lenarviricota* | *Howeltoviricetes* | *Cryppavirales* | *Mitoviridae* | 1 |
| LS_coassembly_k141_1054548 | 683 | Yes | Unknown | *Pisuviricota* | *Duplopiviricetes* | *Durnavirales* | *f.0102.base-Curvula* | 1 |
| LS_coassembly_k141_1069082 | 618 | Yes | ssRNA(+) | *Kitrinoviricota* | *Tolucaviricetes* | *Tolivirales* | *Tombusviridae* | 1 |
| LS_coassembly_k141_1070766 | 1509 | Yes | ssRNA(+) | *Lenarviricota* | *Amabiliviricetes* | *Wolframvirales* | *Narnaviridae* | 1 |
| LS_coassembly_k141_1072213 | 1987 | Yes | Unknown | *Kitrinoviricota* | *c.0034* | *o.0045* | *f.0181* | 1 |
| LS_coassembly_k141_1074570 | 4465 | Yes | ssRNA(+) | *Kitrinoviricota* | *Magsaviricetes* | *Nodamuvirales* | *Nodaviridae* | 2 |
| LS_coassembly_k141_1081653 | 2618 | Yes | Unknown | *Pisuviricota* | *c.0013* | *o.0025.base-Sobeli* | *f.0076* | 1 |
| LS_coassembly_k141_108362 | 1812 | Yes | ssRNA(+) | *Lenarviricota* | *Howeltoviricetes* | *Cryppavirales* | *Mitoviridae* | 1 |
| LS_coassembly_k141_108479 | 2555 | Yes | Unknown | *Pisuviricota* | *c.0013* | *o.0025.base-Sobeli* | *f.0076* | 1 |
| LS_coassembly_k141_1097885 | 1064 | Yes | ssRNA(+) | *Pisuviricota* | *Pisoniviricetes* | *Picornavirales* | *Marnaviridae* | 1 |
| LS_coassembly_k141_1106676 | 2118 | Yes | ssRNA(+) | *Lenarviricota* | *Howeltoviricetes* | *Cryppavirales* | *Mitoviridae* | 1 |
| LS_coassembly_k141_1109689 | 1722 | Yes | ssRNA(+) | *Lenarviricota* | *Amabiliviricetes* | *Wolframvirales* | *Narnaviridae* | 1 |
| LS_coassembly_k141_1113875 | 1310 | Yes | Unknown | *Lenarviricota* | *c.0069.base-Howelto* | *o.0090* | *f.0386* | 1 |
| LS_coassembly_k141_1115973 | 1447 | Yes | Unknown | *Lenarviricota* | *c.0069.base-Howelto* | *o.0090* | *f.0386* | 1 |
| LS_coassembly_k141_112006 | 694 | Yes | ssRNA(+) | *Lenarviricota* | *Amabiliviricetes* | *Wolframvirales* | *Narnaviridae* | 1 |
| LS_coassembly_k141_1121902 | 1070 | Yes | ssRNA(+) | *Lenarviricota* | *Leviviricetes* | *Norzivirales* | *Fiersviridae* | 2 |
| LS_coassembly_k141_1122728 | 1175 | Yes | Unknown | *Lenarviricota* | *c.0069.base-Howelto* | *o.0090* | *f.0386* | 1 |
| LS_coassembly_k141_1124569 | 1238 | Yes | ssRNA(+) | *Lenarviricota* | *Amabiliviricetes* | *Wolframvirales* | *Narnaviridae* | 1 |
| LS_coassembly_k141_112997 | 1420 | Yes | Unknown | *Lenarviricota* | *Leviviricetes* | *Norzivirales* | *f.0368.base-Atkins* | 1 |
| LS_coassembly_k141_1130307 | 1777 | Yes | ssRNA(+) | *Pisuviricota* | *c.0012* | *Sobelivirales* | *Nam-Co_family_28* | 1 |
| LS_coassembly_k141_1156203 | 841 | Yes | Unknown | *Negarnaviricota* | *Ellioviricetes* | *Bunyavirales* | *f.0327* | 1 |
| LS_coassembly_k141_1156677 | 974 | Yes | ssRNA(+) | *Lenarviricota* | *Amabiliviricetes* | *Wolframvirales* | *Narnaviridae* | 1 |
| LS_coassembly_k141_11576 | 4170 | Yes | Unknown | *Negarnaviricota* | *Ellioviricetes* | *Bunyavirales* | *f.0328* | 1 |
| LS_coassembly_k141_1166806 | 3650 | Yes | ssRNA(+) | *Lenarviricota* | *Leviviricetes* | *Timlovirales* | *Steitzviridae* | 3 |
| LS_coassembly_k141_1166858 | 952 | Yes | Unknown | *Kitrinoviricota* | *c.0031* | *o.0042* | *f.0156* | 1 |
| LS_coassembly_k141_1167127 | 3285 | No | NA | *NA* | *NA* | *NA* | *Nam-Co_family_82* | 2 |
| LS_coassembly_k141_1172872 | 2282 | Yes | Unknown | *Negarnaviricota* | *Ellioviricetes* | *Bunyavirales* | *f.0327* | 1 |
| LS_coassembly_k141_1174751 | 1948 | Yes | Unknown | *Pisuviricota* | *c.0013* | *o.0025.base-Sobeli* | *f.0076* | 1 |
| LS_coassembly_k141_1176845 | 740 | Yes | Unknown | *Lenarviricota* | *c.0069.base-Howelto* | *o.0090* | *f.0386* | 1 |
| LS_coassembly_k141_1180159 | 657 | Yes | ssRNA(+) | *Lenarviricota* | *Amabiliviricetes* | *Wolframvirales* | *Narnaviridae* | 1 |
| LS_coassembly_k141_118321 | 658 | Yes | ssRNA(+) | *Lenarviricota* | *Amabiliviricetes* | *Wolframvirales* | *Narnaviridae* | 1 |
| LS_coassembly_k141_1183900 | 1763 | Yes | Unknown | *Lenarviricota* | *Leviviricetes* | *Norzivirales* | *f.0361.base-Solspi* | 2 |
| LS_coassembly_k141_1186789 | 1359 | Yes | ssRNA(+) | *Lenarviricota* | *Amabiliviricetes* | *Wolframvirales* | *Narnaviridae* | 1 |
| LS_coassembly_k141_1194284 | 1824 | Yes | ssRNA(+) | *Pisuviricota* | *Pisoniviricetes* | *Picornavirales* | *Marnaviridae* | 1 |
| LS_coassembly_k141_1200633 | 612 | Yes | ssRNA(+) | *Lenarviricota* | *Howeltoviricetes* | *Cryppavirales* | *Mitoviridae* | 1 |
| LS_coassembly_k141_1216739 | 1222 | Yes | Unknown | *Lenarviricota* | *c.0069.base-Howelto* | *o.0090* | *f.0386* | 1 |
| LS_coassembly_k141_1219574 | 931 | Yes | Unknown | *Lenarviricota* | *c.0067.base-Howelto* | *o.0088* | *f.0384* | 1 |
| LS_coassembly_k141_1223511 | 1113 | Yes | ssRNA(+) | *Pisuviricota* | *c.0012* | *Sobelivirales* | *Solemoviridae* | 1 |
| LS_coassembly_k141_1225259 | 2602 | Yes | Unknown | *Kitrinoviricota* | *c.0025* | *o.0036* | *Nam-Co_family_52* | 2 |
| LS_coassembly_k141_122735 | 2816 | Yes | Unknown | *Lenarviricota* | *c.0071.base-Howelto* | *o.0092* | *f.0388* | 2 |
| LS_coassembly_k141_1250841 | 4717 | Yes | Unknown | *Kitrinoviricota* | *c.0031* | *o.0042* | *f.0156* | 3 |
| LS_coassembly_k141_125234 | 1263 | Yes | dsRNA | *Pisuviricota* | *Duplopiviricetes* | *Durnavirales* | *Partitiviridae* | 1 |
| LS_coassembly_k141_1254060 | 5952 | Yes | Unknown | *Kitrinoviricota* | *c.0034* | *o.0045* | *Nam-Co_family_58* | 2 |
| LS_coassembly_k141_126227 | 1382 | Yes | dsRNA | *Pisuviricota* | *Duplopiviricetes* | *Durnavirales* | *Partitiviridae* | 1 |
| LS_coassembly_k141_1266212 | 875 | Yes | Unknown | *Lenarviricota* | *c.0074.base-Howelto* | *o.0095* | *f.0391* | 1 |
| LS_coassembly_k141_126965 | 1497 | Yes | Unknown | *Pisuviricota* | *c.0013* | *o.0025.base-Sobeli* | *f.0076* | 1 |
| LS_coassembly_k141_127221 | 994 | Yes | Unknown | *Pisuviricota* | *c.0013* | *o.0025.base-Sobeli* | *f.0076* | 1 |
| LS_coassembly_k141_1274756 | 1304 | Yes | Unknown | *Pisuviricota* | *Duplopiviricetes* | *Durnavirales* | *f.0102.base-Curvula* | 1 |
| LS_coassembly_k141_1278143 | 2669 | Yes | ssRNA(+) | *Lenarviricota* | *Amabiliviricetes* | *Wolframvirales* | *Narnaviridae* | 1 |
| LS_coassembly_k141_1288836 | 4241 | Yes | Unknown | *Duplornaviricota* | *Chrymotiviricetes* | *Ghabrivirales* | *f.0292* | 2 |
| LS_coassembly_k141_1317493 | 1330 | Yes | Unknown | *Pisuviricota* | *Duplopiviricetes* | *Durnavirales* | *f.0102.base-Curvula* | 1 |
| LS_coassembly_k141_1328102 | 790 | Yes | ssRNA(+) | *Lenarviricota* | *Amabiliviricetes* | *Wolframvirales* | *Narnaviridae* | 1 |
| LS_coassembly_k141_1328847 | 737 | Yes | ssRNA(+) | *Lenarviricota* | *Amabiliviricetes* | *Wolframvirales* | *Narnaviridae* | 1 |
| LS_coassembly_k141_1338665 | 1337 | Yes | Unknown | *Lenarviricota* | *Leviviricetes* | *Norzivirales* | *f.0368.base-Atkins* | 1 |
| LS_coassembly_k141_1340706 | 2599 | Yes | ssRNA(+) | *Lenarviricota* | *Amabiliviricetes* | *Wolframvirales* | *Narnaviridae* | 1 |
| LS_coassembly_k141_1348429 | 2024 | Yes | Unknown | *Lenarviricota* | *c.0069.base-Howelto* | *o.0090* | *f.0386* | 1 |
| LS_coassembly_k141_1350286 | 759 | Yes | ssRNA(+) | *Lenarviricota* | *Amabiliviricetes* | *Wolframvirales* | *Nam-Co_family_10* | 1 |
| LS_coassembly_k141_138809 | 722 | Yes | dsRNA | *Duplornaviricota* | *Chrymotiviricetes* | *Ghabrivirales* | *Totiviridae* | 2 |
| LS_coassembly_k141_150032 | 1461 | Yes | Unknown | *Lenarviricota* | *c.0061.base-Levi* | *o.0082* | *Nam-Co_family_14* | 1 |
| LS_coassembly_k141_152525 | 882 | Yes | ssRNA(+) | *Lenarviricota* | *Leviviricetes* | *Timlovirales* | *Steitzviridae* | 1 |
| LS_coassembly_k141_153349 | 2219 | Yes | Unknown | *Pisuviricota* | *Duplopiviricetes* | *Durnavirales* | *f.0102.base-Curvula* | 1 |
| LS_coassembly_k141_161594 | 785 | Yes | dsRNA | *Pisuviricota* | *Duplopiviricetes* | *Durnavirales* | *Partitiviridae* | 1 |
| LS_coassembly_k141_165614 | 3727 | Yes | Unknown | *Pisuviricota* | *Duplopiviricetes* | *Durnavirales* | *f.0102.base-Curvula* | 4 |
| LS_coassembly_k141_16911 | 3332 | Yes | Unknown | *Duplornaviricota* | *Chrymotiviricetes* | *Ghabrivirales* | *f.0287* | 2 |
| LS_coassembly_k141_169245 | 1005 | Yes | Unknown | *Lenarviricota* | *Leviviricetes* | *Norzivirales* | *f.0367.base-Atkins* | 1 |
| LS_coassembly_k141_17420 | 822 | Yes | dsRNA | *Pisuviricota* | *c.0012* | *Sobelivirales* | *Barnaviridae* | 1 |
| LS_coassembly_k141_174926 | 1589 | Yes | ssRNA(+) | *Lenarviricota* | *Amabiliviricetes* | *Wolframvirales* | *Narnaviridae* | 1 |
| LS_coassembly_k141_178385 | 9475 | Yes | ssRNA(+) | *Pisuviricota* | *Pisoniviricetes* | *Picornavirales* | *Marnaviridae* | 2 |
| LS_coassembly_k141_182501 | 1988 | Yes | ssRNA(+) | *Lenarviricota* | *Howeltoviricetes* | *Cryppavirales* | *Mitoviridae* | 1 |
| LS_coassembly_k141_18251 | 1079 | Yes | Unknown | *Lenarviricota* | *c.0069.base-Howelto* | *o.0090* | *Nam-Co_family_15* | 1 |
| LS_coassembly_k141_18378 | 1285 | Yes | dsRNA | *Pisuviricota* | *Duplopiviricetes* | *Durnavirales* | *Partitiviridae* | 1 |
| LS_coassembly_k141_186086 | 1072 | Yes | ssRNA(+) | *Lenarviricota* | *Amabiliviricetes* | *Wolframvirales* | *Narnaviridae* | 1 |
| LS_coassembly_k141_186661 | 821 | Yes | ssRNA(+) | *Lenarviricota* | *Amabiliviricetes* | *Wolframvirales* | *Narnaviridae* | 1 |
| LS_coassembly_k141_187532 | 1074 | Yes | ssRNA(+) | *Pisuviricota* | *Pisoniviricetes* | *Picornavirales* | *Nam-Co_family_41* | 1 |
| LS_coassembly_k141_190706 | 1159 | Yes | ssRNA(+) | *Pisuviricota* | *Pisoniviricetes* | *Picornavirales* | *Marnaviridae* | 1 |
| LS_coassembly_k141_191710 | 2061 | Yes | Unknown | *Pisuviricota* | *Duplopiviricetes* | *Durnavirales* | *f.0111* | 2 |
| LS_coassembly_k141_192514 | 1170 | Yes | ssRNA(+) | *Lenarviricota* | *Amabiliviricetes* | *Wolframvirales* | *Narnaviridae* | 1 |
| LS_coassembly_k141_19643 | 2038 | Yes | ssRNA(+) | *Lenarviricota* | *Amabiliviricetes* | *Wolframvirales* | *Narnaviridae* | 1 |
| LS_coassembly_k141_199685 | 1852 | Yes | ssRNA(+) | *Lenarviricota* | *Miaviricetes* | *Ourlivirales* | *Botourmiaviridae* | 1 |
| LS_coassembly_k141_210683 | 869 | Yes | ssRNA(+) | *Kitrinoviricota* | *Alsuviricetes* | *o.0055* | *Endornaviridae* | 1 |
| LS_coassembly_k141_223527 | 761 | Yes | Unknown | *Kitrinoviricota* | *c.0032* | *o.0043* | *Nam-Co_family_56* | 1 |
| LS_coassembly_k141_226167 | 3794 | Yes | ssRNA(+) | *Pisuviricota* | *Pisoniviricetes* | *Picornavirales* | *Polycipiviridae* | 1 |
| LS_coassembly_k141_232625 | 1922 | Yes | Unknown | *Pisuviricota* | *Duplopiviricetes* | *Durnavirales* | *f.0102.base-Curvula* | 2 |
| LS_coassembly_k141_241270 | 2425 | Yes | ssRNA(+) | *Kitrinoviricota* | *Tolucaviricetes* | *Tolivirales* | *Tombusviridae* | 2 |
| LS_coassembly_k141_244743 | 701 | Yes | Unknown | *Lenarviricota* | *c.0069.base-Howelto* | *o.0090* | *f.0386* | 1 |
| LS_coassembly_k141_259450 | 1074 | Yes | ssRNA(+) | *Lenarviricota* | *Leviviricetes* | *Norzivirales* | *Fiersviridae* | 1 |
| LS_coassembly_k141_260675 | 873 | Yes | ssRNA(+) | *Pisuviricota* | *c.0012* | *Sobelivirales* | *Solemoviridae* | 1 |
| LS_coassembly_k141_268503 | 10051 | Yes | ssRNA(+) | *Pisuviricota* | *Pisoniviricetes* | *Picornavirales* | *Dicistroviridae* | 4 |
| LS_coassembly_k141_280312 | 1037 | Yes | Unknown | *Lenarviricota* | *c.0068.base-Howelto* | *o.0089* | *f.0385* | 1 |
| LS_coassembly_k141_283877 | 1605 | Yes | ssRNA(+) | *Pisuviricota* | *c.0012* | *Sobelivirales* | *Solemoviridae* | 2 |
| LS_coassembly_k141_287825 | 776 | Yes | ssRNA(-) | *Negarnaviricota* | *Monjiviricetes* | *Mononegavirales* | *Rhabdoviridae* | 1 |
| LS_coassembly_k141_292708 | 1716 | Yes | Unknown | *Pisuviricota* | *Duplopiviricetes* | *Durnavirales* | *f.0102.base-Curvula* | 1 |
| LS_coassembly_k141_296363 | 2357 | Yes | Unknown | *Pisuviricota* | *c.0013* | *o.0025.base-Sobeli* | *f.0076* | 1 |
| LS_coassembly_k141_300460 | 2665 | Yes | ssRNA(+) | *Lenarviricota* | *Amabiliviricetes* | *Wolframvirales* | *Narnaviridae* | 1 |
| LS_coassembly_k141_303746 | 1420 | Yes | Unknown | *Pisuviricota* | *Pisoniviricetes* | *Picornavirales* | *f.0048.base-Marna* | 1 |
| LS_coassembly_k141_30722 | 3124 | Yes | ssRNA(+) | *Lenarviricota* | *Miaviricetes* | *Ourlivirales* | *Botourmiaviridae* | 2 |
| LS_coassembly_k141_327859 | 1963 | Yes | ssRNA(+) | *Lenarviricota* | *Amabiliviricetes* | *Wolframvirales* | *Narnaviridae* | 1 |
| LS_coassembly_k141_332531 | 2999 | Yes | Unknown | *Kitrinoviricota* | *c.0021.base-Toluca* | *o.0032* | *f.0125* | 2 |
| LS_coassembly_k141_335016 | 837 | Yes | ssRNA(+) | *Lenarviricota* | *Howeltoviricetes* | *Cryppavirales* | *Mitoviridae* | 1 |
| LS_coassembly_k141_338604 | 2669 | No | NA | *NA* | *NA* | *NA* | *NA* | 1 |
| LS_coassembly_k141_345578 | 773 | Yes | ssRNA(+) | *Lenarviricota* | *Amabiliviricetes* | *Wolframvirales* | *Narnaviridae* | 1 |
| LS_coassembly_k141_347998 | 3546 | Yes | ssRNA(+) | *Lenarviricota* | *Miaviricetes* | *Ourlivirales* | *Nam-Co_family_19* | 2 |
| LS_coassembly_k141_35229 | 1398 | Yes | Unknown | *Pisuviricota* | *Duplopiviricetes* | *Durnavirales* | *Nam-Co_family_39* | 1 |
| LS_coassembly_k141_353392 | 1567 | Yes | Unknown | *Pisuviricota* | *Duplopiviricetes* | *Durnavirales* | *f.0111* | 1 |
| LS_coassembly_k141_356040 | 1694 | Yes | dsRNA | *Duplornaviricota* | *Resentoviricetes* | *Reovirales* | *Nam-Co_family_75* | 1 |
| LS_coassembly_k141_356991 | 1233 | Yes | Unknown | *Kitrinoviricota* | *c.0034* | *o.0045* | *f.0176* | 1 |
| LS_coassembly_k141_357820 | 1581 | Yes | Unknown | *Pisuviricota* | *c.0013* | *o.0025.base-Sobeli* | *f.0076* | 2 |
| LS_coassembly_k141_368163 | 731 | Yes | ssRNA(+) | *Lenarviricota* | *Amabiliviricetes* | *Wolframvirales* | *Narnaviridae* | 1 |
| LS_coassembly_k141_370211 | 731 | Yes | ssRNA(+) | *Lenarviricota* | *Leviviricetes* | *Timlovirales* | *Steitzviridae* | 1 |
| LS_coassembly_k141_370511 | 849 | Yes | Unknown | *Pisuviricota* | *Duplopiviricetes* | *Durnavirales* | *Nam-Co_family_39* | 1 |
| LS_coassembly_k141_370863 | 2549 | Yes | Unknown | *Pisuviricota* | *c.0013* | *o.0025.base-Sobeli* | *f.0076* | 2 |
| LS_coassembly_k141_374349 | 2273 | No | NA | *NA* | *NA* | *NA* | *Nam-Co_family_20* | 1 |
| LS_coassembly_k141_379730 | 4264 | Yes | Unknown | *Lenarviricota* | *Leviviricetes* | *Norzivirales* | *Nam-Co_family_18* | 3 |
| LS_coassembly_k141_380972 | 3285 | Yes | dsRNA | *Duplornaviricota* | *Resentoviricetes* | *Reovirales* | *Reoviridae* | 1 |
| LS_coassembly_k141_389387 | 2762 | Yes | ssRNA(+) | *Pisuviricota* | *c.0012* | *Sobelivirales* | *Solemoviridae* | 2 |
| LS_coassembly_k141_399847 | 1840 | Yes | Unknown | *Lenarviricota* | *c.0067.base-Howelto* | *o.0088* | *f.0384* | 1 |
| LS_coassembly_k141_399972 | 662 | Yes | Unknown | *Kitrinoviricota* | *Alsuviricetes* | *Hepelivirales* | *f.0221.base-Alphatetra* | 1 |
| LS_coassembly_k141_401458 | 629 | Yes | ssRNA(+) | *Lenarviricota* | *Howeltoviricetes* | *Cryppavirales* | *Mitoviridae* | 1 |
| LS_coassembly_k141_405215 | 775 | Yes | Unknown | *Pisuviricota* | *Duplopiviricetes* | *Durnavirales* | *f.0102.base-Curvula* | 1 |
| LS_coassembly_k141_407824 | 6744 | Yes | Unknown | *Negarnaviricota* | *c.0054.base-Yunchang* | *o.0072* | *f.0354* | 2 |
| LS_coassembly_k141_408106 | 1389 | Yes | ssRNA(+) | *Pisuviricota* | *Flasuviricetes* | *Amarillovirales* | *Flaviviridae* | 1 |
| LS_coassembly_k141_409312 | 6491 | Yes | ssRNA(+) | *Pisuviricota* | *Pisoniviricetes* | *Picornavirales* | *Nam-Co_family_42* | 2 |
| LS_coassembly_k141_417327 | 640 | Yes | ssRNA(+) | *Lenarviricota* | *Howeltoviricetes* | *Cryppavirales* | *Mitoviridae* | 1 |
| LS_coassembly_k141_419215 | 1017 | Yes | Unknown | *Duplornaviricota* | *Chrymotiviricetes* | *Ghabrivirales* | *Nam-Co_family_72* | 1 |
| LS_coassembly_k141_420880 | 1260 | Yes | Unknown | *Pisuviricota* | *Duplopiviricetes* | *Durnavirales* | *Nam-Co_family_37* | 1 |
| LS_coassembly_k141_426567 | 652 | No | NA | *NA* | *NA* | *NA* | *NA* | 1 |
| LS_coassembly_k141_434311 | 732 | Yes | Unknown | *Pisuviricota* | *Pisoniviricetes* | *Picornavirales* | *f.0008.base-Polycipi* | 1 |
| LS_coassembly_k141_436759 | 1176 | Yes | dsRNA | *Duplornaviricota* | *Resentoviricetes* | *Reovirales* | *Nam-Co_family_76* | 1 |
| LS_coassembly_k141_447615 | 1277 | Yes | ssRNA(+) | *Lenarviricota* | *Leviviricetes* | *Norzivirales* | *Fiersviridae* | 1 |
| LS_coassembly_k141_4481 | 2263 | Yes | ssRNA(+) | *Lenarviricota* | *Leviviricetes* | *Norzivirales* | *Solspiviridae* | 2 |
| LS_coassembly_k141_460169 | 2281 | Yes | ssRNA(+) | *Lenarviricota* | *Howeltoviricetes* | *Cryppavirales* | *Mitoviridae* | 1 |
| LS_coassembly_k141_481010 | 1266 | Yes | Unknown | *Pisuviricota* | *Duplopiviricetes* | *Durnavirales* | *f.0111* | 1 |
| LS_coassembly_k141_483032 | 1419 | Yes | ssRNA(+) | *Lenarviricota* | *Amabiliviricetes* | *Wolframvirales* | *Nam-Co_family_08* | 1 |
| LS_coassembly_k141_496413 | 2091 | No | NA | *NA* | *NA* | *NA* | *NA* | 1 |
| LS_coassembly_k141_502318 | 1146 | Yes | ssRNA(+) | *Lenarviricota* | *Howeltoviricetes* | *Cryppavirales* | *Mitoviridae* | 1 |
| LS_coassembly_k141_507967 | 2237 | Yes | ssRNA(+) | *Kitrinoviricota* | *Magsaviricetes* | *Nodamuvirales* | *Nodaviridae* | 1 |
| LS_coassembly_k141_512850 | 6868 | Yes | Unknown | *Kitrinoviricota* | *c.0034* | *o.0045* | *Nam-Co_family_59* | 2 |
| LS_coassembly_k141_516141 | 5071 | No | NA | *NA* | *NA* | *NA* | *Nam-Co_family_78* | 1 |
| LS_coassembly_k141_522337 | 2365 | Yes | dsRNA | *Duplornaviricota* | *Resentoviricetes* | *Reovirales* | *Reoviridae* | 1 |
| LS_coassembly_k141_532808 | 778 | Yes | Unknown | *Lenarviricota* | *c.0069.base-Howelto* | *o.0090* | *f.0386* | 1 |
| LS_coassembly_k141_542386 | 690 | Yes | ssRNA(+) | *Lenarviricota* | *Leviviricetes* | *Timlovirales* | *Steitzviridae* | 1 |
| LS_coassembly_k141_542559 | 903 | Yes | Unknown | *Duplornaviricota* | *Chrymotiviricetes* | *Ghabrivirales* | *f.0292* | 1 |
| LS_coassembly_k141_543237 | 3710 | Yes | ssRNA(+) | *Lenarviricota* | *Leviviricetes* | *Norzivirales* | *Fiersviridae* | 3 |
| LS_coassembly_k141_54629 | 2059 | Yes | ssRNA(+) | *Lenarviricota* | *Miaviricetes* | *Ourlivirales* | *Botourmiaviridae* | 1 |
| LS_coassembly_k141_547291 | 3438 | No | NA | *NA* | *NA* | *NA* | *Nam-Co_family_21* | 1 |
| LS_coassembly_k141_559789 | 1653 | Yes | Unknown | *Pisuviricota* | *Pisoniviricetes* | *Picornavirales* | *f.0030* | 2 |
| LS_coassembly_k141_562058 | 2093 | Yes | Unknown | *Lenarviricota* | *c.0069.base-Howelto* | *o.0090* | *f.0386* | 1 |
| LS_coassembly_k141_565475 | 1598 | Yes | ssRNA(+) | *Kitrinoviricota* | *Tolucaviricetes* | *Tolivirales* | *Tombusviridae* | 2 |
| LS_coassembly_k141_575813 | 783 | Yes | ssRNA(+) | *Lenarviricota* | *Amabiliviricetes* | *Wolframvirales* | *Narnaviridae* | 1 |
| LS_coassembly_k141_576688 | 1930 | Yes | Unknown | *Duplornaviricota* | *Chrymotiviricetes* | *Ghabrivirales* | *f.0296* | 1 |
| LS_coassembly_k141_57787 | 5137 | Yes | ssRNA(+) | *Kitrinoviricota* | *Tolucaviricetes* | *Tolivirales* | *Tombusviridae* | 7 |
| LS_coassembly_k141_582323 | 2518 | Yes | ssRNA(+) | *Lenarviricota* | *Howeltoviricetes* | *Cryppavirales* | *Mitoviridae* | 2 |
| LS_coassembly_k141_583536 | 1110 | Yes | ssRNA(+) | *Kitrinoviricota* | *Alsuviricetes* | *o.0055* | *Endornaviridae* | 1 |
| LS_coassembly_k141_588663 | 1798 | Yes | Unknown | *Lenarviricota* | *c.0069.base-Howelto* | *o.0090* | *f.0386* | 1 |
| LS_coassembly_k141_604047 | 656 | Yes | ssRNA(+) | *Lenarviricota* | *Miaviricetes* | *Ourlivirales* | *Botourmiaviridae* | 1 |
| LS_coassembly_k141_604560 | 2658 | Yes | ssRNA(+) | *Lenarviricota* | *Amabiliviricetes* | *Wolframvirales* | *Narnaviridae* | 1 |
| LS_coassembly_k141_607677 | 952 | Yes | ssRNA(+) | *Kitrinoviricota* | *Alsuviricetes* | *o.0055* | *Endornaviridae* | 1 |
| LS_coassembly_k141_611362 | 8000 | Yes | ssRNA(+) | *Pisuviricota* | *Pisoniviricetes* | *Picornavirales* | *Dicistroviridae* | 2 |
| LS_coassembly_k141_613763 | 1226 | Yes | Unknown | *Pisuviricota* | *c.0013* | *o.0025.base-Sobeli* | *f.0076* | 1 |
| LS_coassembly_k141_625058 | 954 | Yes | ssRNA(+) | *Pisuviricota* | *Pisoniviricetes* | *Picornavirales* | *Dicistroviridae* | 1 |
| LS_coassembly_k141_631699 | 627 | Yes | ssRNA(+) | *Lenarviricota* | *Leviviricetes* | *Timlovirales* | *Steitzviridae* | 1 |
| LS_coassembly_k141_63722 | 3453 | Yes | ssRNA(+) | *Lenarviricota* | *Leviviricetes* | *Norzivirales* | *Fiersviridae* | 3 |
| LS_coassembly_k141_638690 | 4034 | Yes | ssRNA(+) | *Lenarviricota* | *Leviviricetes* | *Timlovirales* | *Steitzviridae* | 3 |
| LS_coassembly_k141_640025 | 825 | Yes | Unknown | *Lenarviricota* | *c.0068.base-Howelto* | *o.0089* | *f.0385* | 1 |
| LS_coassembly_k141_647081 | 705 | Yes | ssRNA(+) | *Lenarviricota* | *Amabiliviricetes* | *Wolframvirales* | *Narnaviridae* | 1 |
| LS_coassembly_k141_650154 | 1705 | Yes | ssRNA(+) | *Lenarviricota* | *Amabiliviricetes* | *Wolframvirales* | *Narnaviridae* | 1 |
| LS_coassembly_k141_656443 | 3806 | Yes | ssRNA(+) | *Lenarviricota* | *Leviviricetes* | *Norzivirales* | *Fiersviridae* | 4 |
| LS_coassembly_k141_659713 | 1973 | Yes | Unknown | *Pisuviricota* | *c.0013* | *o.0025.base-Sobeli* | *f.0076* | 1 |
| LS_coassembly_k141_660091 | 1043 | Yes | Unknown | *Kitrinoviricota* | *c.0024* | *o.0035* | *f.0144* | 2 |
| LS_coassembly_k141_665176 | 960 | Yes | Unknown | *Duplornaviricota* | *Chrymotiviricetes* | *Ghabrivirales* | *f.0296* | 1 |
| LS_coassembly_k141_666330 | 677 | Yes | ssRNA(+) | *Pisuviricota* | *Pisoniviricetes* | *Picornavirales* | *Nam-Co_family_46* | 1 |
| LS_coassembly_k141_668838 | 1665 | No | NA | *NA* | *NA* | *NA* | *NA* | 2 |
| LS_coassembly_k141_673941 | 2617 | Yes | Unknown | *Pisuviricota* | *Pisoniviricetes* | *Picornavirales* | *f.0024.base-Solinvi* | 1 |
| LS_coassembly_k141_677142 | 611 | Yes | ssRNA(+) | *Pisuviricota* | *Pisoniviricetes* | *Picornavirales* | *Dicistroviridae* | 1 |
| LS_coassembly_k141_677412 | 672 | Yes | ssRNA(+) | *Lenarviricota* | *Amabiliviricetes* | *Wolframvirales* | *Narnaviridae* | 1 |
| LS_coassembly_k141_689206 | 1737 | Yes | Unknown | *Pisuviricota* | *Duplopiviricetes* | *Durnavirales* | *Nam-Co_family_39* | 1 |
| LS_coassembly_k141_690095 | 655 | Yes | ssRNA(+) | *Lenarviricota* | *Howeltoviricetes* | *Cryppavirales* | *Mitoviridae* | 1 |
| LS_coassembly_k141_70220 | 1685 | Yes | ssRNA(+) | *Lenarviricota* | *Amabiliviricetes* | *Wolframvirales* | *Nam-Co_family_13* | 1 |
| LS_coassembly_k141_705223 | 2525 | Yes | ssRNA(+) | *Pisuviricota* | *Stelpaviricetes* | *Stellavirales* | *Astroviridae* | 2 |
| LS_coassembly_k141_708466 | 647 | Yes | ssRNA(+) | *Kitrinoviricota* | *Tolucaviricetes* | *Tolivirales* | *Tombusviridae* | 1 |
| LS_coassembly_k141_709523 | 1777 | Yes | ssRNA(+) | *Lenarviricota* | *Leviviricetes* | *Norzivirales* | *Fiersviridae* | 2 |
| LS_coassembly_k141_718255 | 649 | Yes | ssRNA(+) | *Lenarviricota* | *Howeltoviricetes* | *Cryppavirales* | *Mitoviridae* | 1 |
| LS_coassembly_k141_723190 | 816 | Yes | Unknown | *Lenarviricota* | *c.0066.base-Howelto* | *o.0087* | *f.0383* | 1 |
| LS_coassembly_k141_734239 | 2600 | Yes | ssRNA(+) | *Pisuviricota* | *Pisoniviricetes* | *Picornavirales* | *Dicistroviridae* | 1 |
| LS_coassembly_k141_73853 | 1462 | Yes | ssRNA(+) | *Lenarviricota* | *Amabiliviricetes* | *Wolframvirales* | *Narnaviridae* | 1 |
| LS_coassembly_k141_738808 | 6229 | Yes | Unknown | *Kitrinoviricota* | *c.0032* | *o.0043* | *Nam-Co_family_55* | 3 |
| LS_coassembly_k141_740060 | 706 | Yes | dsRNA | *Pisuviricota* | *Duplopiviricetes* | *Durnavirales* | *Partitiviridae* | 1 |
| LS_coassembly_k141_741592 | 2381 | Yes | ssRNA(+) | *Pisuviricota* | *Pisoniviricetes* | *Picornavirales* | *Marnaviridae* | 2 |
| LS_coassembly_k141_747216 | 949 | Yes | Unknown | *Pisuviricota* | *Duplopiviricetes* | *Durnavirales* | *f.0102.base-Curvula* | 1 |
| LS_coassembly_k141_755496 | 3856 | No | NA | *NA* | *NA* | *NA* | *Nam-Co_family_84* | 2 |
| LS_coassembly_k141_765865 | 2893 | Yes | ssRNA(+) | *Kitrinoviricota* | *Tolucaviricetes* | *Tolivirales* | *Tombusviridae* | 3 |
| LS_coassembly_k141_77005 | 932 | Yes | ssRNA(+) | *Pisuviricota* | *Pisoniviricetes* | *Picornavirales* | *Marnaviridae* | 1 |
| LS_coassembly_k141_774987 | 1010 | Yes | dsRNA | *Pisuviricota* | *Duplopiviricetes* | *Durnavirales* | *Partitiviridae* | 1 |
| LS_coassembly_k141_777368 | 1406 | Yes | ssRNA(+) | *Lenarviricota* | *Howeltoviricetes* | *Cryppavirales* | *Mitoviridae* | 1 |
| LS_coassembly_k141_783029 | 1463 | Yes | ssRNA(+) | *Kitrinoviricota* | *Tolucaviricetes* | *Tolivirales* | *Tombusviridae* | 2 |
| LS_coassembly_k141_786484 | 1947 | Yes | Unknown | *Lenarviricota* | *c.0069.base-Howelto* | *o.0090* | *f.0386* | 1 |
| LS_coassembly_k141_790503 | 897 | Yes | Unknown | *Pisuviricota* | *Pisoniviricetes* | *Picornavirales* | *f.0033* | 1 |
| LS_coassembly_k141_795198 | 9975 | Yes | ssRNA(+) | *Pisuviricota* | *Stelpaviricetes* | *Patatavirales* | *Potyviridae* | 1 |
| LS_coassembly_k141_808201 | 4145 | Yes | Unknown | *Kitrinoviricota* | *c.0024* | *o.0035* | *f.0144* | 2 |
| LS_coassembly_k141_808804 | 1056 | Yes | ssRNA(+) | *Lenarviricota* | *Leviviricetes* | *Norzivirales* | *Fiersviridae* | 1 |
| LS_coassembly_k141_812161 | 1479 | Yes | Unknown | *Pisuviricota* | *Duplopiviricetes* | *Durnavirales* | *f.0102.base-Curvula* | 1 |
| LS_coassembly_k141_814387 | 1486 | Yes | ssRNA(+) | *Lenarviricota* | *Amabiliviricetes* | *Wolframvirales* | *Narnaviridae* | 1 |
| LS_coassembly_k141_818259 | 11439 | Yes | ssRNA(+) | *Pisuviricota* | *Pisoniviricetes* | *Picornavirales* | *Polycipiviridae* | 6 |
| LS_coassembly_k141_824362 | 2872 | Yes | ssRNA(+) | *Lenarviricota* | *Leviviricetes* | *Timlovirales* | *Steitzviridae* | 3 |
| LS_coassembly_k141_839853 | 4032 | Yes | ssRNA(+) | *Lenarviricota* | *Leviviricetes* | *Timlovirales* | *Steitzviridae* | 3 |
| LS_coassembly_k141_839961 | 1328 | Yes | ssRNA(+) | *Lenarviricota* | *Leviviricetes* | *Norzivirales* | *Fiersviridae* | 1 |
| LS_coassembly_k141_841983 | 613 | Yes | Unknown | *Pisuviricota* | *Duplopiviricetes* | *Durnavirales* | *f.0111* | 1 |
| LS_coassembly_k141_84489 | 739 | Yes | Unknown | *Pisuviricota* | *Duplopiviricetes* | *Durnavirales* | *f.0111* | 1 |
| LS_coassembly_k141_84850 | 8411 | Yes | ssRNA(+) | *Pisuviricota* | *Pisoniviricetes* | *Picornavirales* | *Dicistroviridae* | 3 |
| LS_coassembly_k141_851065 | 1283 | Yes | ssRNA(+) | *Lenarviricota* | *Howeltoviricetes* | *Cryppavirales* | *Mitoviridae* | 1 |
| LS_coassembly_k141_856822 | 2777 | Yes | ssRNA(+) | *Lenarviricota* | *Miaviricetes* | *Ourlivirales* | *Botourmiaviridae* | 2 |
| LS_coassembly_k141_873293 | 1049 | Yes | Unknown | *Pisuviricota* | *c.0013* | *o.0025.base-Sobeli* | *f.0076* | 1 |
| LS_coassembly_k141_880328 | 2217 | Yes | ssRNA(+) | *Lenarviricota* | *Howeltoviricetes* | *Cryppavirales* | *Mitoviridae* | 1 |
| LS_coassembly_k141_881219 | 1003 | Yes | ssRNA(+) | *Lenarviricota* | *Amabiliviricetes* | *Wolframvirales* | *Narnaviridae* | 1 |
| LS_coassembly_k141_882900 | 714 | Yes | Unknown | *Pisuviricota* | *Pisoniviricetes* | *Picornavirales* | *f.0008.base-Polycipi* | 1 |
| LS_coassembly_k141_882929 | 789 | Yes | Unknown | *Lenarviricota* | *c.0069.base-Howelto* | *o.0090* | *f.0386* | 1 |
| LS_coassembly_k141_886170 | 1659 | Yes | Unknown | *Lenarviricota* | *c.0069.base-Howelto* | *o.0090* | *f.0386* | 2 |
| LS_coassembly_k141_8864 | 3379 | Yes | ssRNA(+) | *Lenarviricota* | *Leviviricetes* | *Norzivirales* | *Fiersviridae* | 3 |
| LS_coassembly_k141_891738 | 2470 | Yes | ssRNA(+) | *Lenarviricota* | *Howeltoviricetes* | *Cryppavirales* | *Mitoviridae* | 5 |
| LS_coassembly_k141_897578 | 625 | Yes | ssRNA(+) | *Pisuviricota* | *Pisoniviricetes* | *Picornavirales* | *Marnaviridae* | 1 |
| LS_coassembly_k141_908476 | 1149 | Yes | ssRNA(+) | *Pisuviricota* | *Stelpaviricetes* | *Stellavirales* | *Astroviridae* | 1 |
| LS_coassembly_k141_910916 | 3604 | No | NA | *NA* | *NA* | *NA* | *Nam-Co_family_01* | 1 |
| LS_coassembly_k141_919836 | 2151 | Yes | ssRNA(+) | *Lenarviricota* | *Howeltoviricetes* | *Cryppavirales* | *Mitoviridae* | 1 |
| LS_coassembly_k141_928946 | 703 | Yes | ssRNA(+) | *Lenarviricota* | *Amabiliviricetes* | *Wolframvirales* | *Narnaviridae* | 1 |
| LS_coassembly_k141_947059 | 1042 | Yes | Unknown | *Pisuviricota* | *Duplopiviricetes* | *Durnavirales* | *f.0102.base-Curvula* | 1 |
| LS_coassembly_k141_947790 | 3040 | Yes | ssRNA(+) | *Lenarviricota* | *Leviviricetes* | *Norzivirales* | *Fiersviridae* | 3 |
| LS_coassembly_k141_950304 | 16031 | Yes | ssRNA(-) | *Negarnaviricota* | *Monjiviricetes* | *Mononegavirales* | *Rhabdoviridae* | 11 |
| LS_coassembly_k141_952038 | 2096 | Yes | ssRNA(+) | *Kitrinoviricota* | *Magsaviricetes* | *Nodamuvirales* | *Nodaviridae* | 1 |
| LS_coassembly_k141_95774 | 870 | Yes | Unknown | *Lenarviricota* | *c.0069.base-Howelto* | *o.0090* | *f.0386* | 1 |
| LS_coassembly_k141_965508 | 1454 | Yes | ssRNA(+) | *Lenarviricota* | *Amabiliviricetes* | *Wolframvirales* | *Narnaviridae* | 1 |
| LS_coassembly_k141_966457 | 637 | Yes | Unknown | *Lenarviricota* | *c.0071.base-Howelto* | *o.0092* | *f.0388* | 1 |
| LS_coassembly_k141_967088 | 864 | Yes | Unknown | *Lenarviricota* | *c.0069.base-Howelto* | *o.0090* | *f.0386* | 1 |
| LS_coassembly_k141_969233 | 1246 | Yes | Unknown | *Pisuviricota* | *Duplopiviricetes* | *Durnavirales* | *f.0102.base-Curvula* | 1 |
| LS_coassembly_k141_969611 | 818 | Yes | ssRNA(+) | *Lenarviricota* | *Amabiliviricetes* | *Wolframvirales* | *Narnaviridae* | 1 |
| LS_coassembly_k141_980808 | 3301 | Yes | ssRNA(+) | *Lenarviricota* | *Miaviricetes* | *Ourlivirales* | *Botourmiaviridae* | 1 |
| LS_coassembly_k141_981221 | 3805 | Yes | Unknown | *Lenarviricota* | *Leviviricetes* | *Norzivirales* | *f.0361.base-Solspi* | 3 |
| LS_coassembly_k141_981726 | 744 | Yes | Unknown | *Kitrinoviricota* | *c.0024* | *o.0035* | *f.0144* | 1 |
| LS_coassembly_k141_985680 | 636 | No | NA | *NA* | *NA* | *NA* | *NA* | 1 |
| LS_coassembly_k141_986866 | 3533 | Yes | Unknown | *Pisuviricota* | *Duplopiviricetes* | *Durnavirales* | *Nam-Co_family_33* | 2 |
| LS_coassembly_k141_9898 | 2572 | Yes | Unknown | *Duplornaviricota* | *Chrymotiviricetes* | *Ghabrivirales* | *f.0286* | 2 |
| LS_coassembly_k141_995692 | 2186 | Yes | ssRNA(+) | *Lenarviricota* | *Amabiliviricetes* | *Wolframvirales* | *Narnaviridae* | 1 |

| **Table S6. Gene annotations for RdRP-containing contigs (over 3 ORFs) in Lake Nam Co.** | | | | | | |
| --- | --- | --- | --- | --- | --- | --- |
| **Gene** | **Contig** | **Start** | **End** | **Strand** | **Annotation** | **Type** |
| LC_80m_1.2um_k141_403562_1 | LC_80m_1.2um_k141_403562 | 530 | 1540 | - | hypothetical protein | hypothetical protein |
| LC_80m_1.2um_k141_403562_2 | LC_80m_1.2um_k141_403562 | 1622 | 3610 | - | RDRP | RDRP |
| LC_80m_1.2um_k141_403562_3 | LC_80m_1.2um_k141_403562 | 3641 | 4033 | - | hypothetical protein | hypothetical protein |
| LC_80m_1.2um_k141_403562_4 | LC_80m_1.2um_k141_403562 | 4454 | 4723 | - | hypothetical protein | hypothetical protein |
| LC_80m_1.2um_k141_403562_5 | LC_80m_1.2um_k141_403562 | 5021 | 7762 | - | hypothetical protein | hypothetical protein |
| LC_80m_1.2um_k141_403562_6 | LC_80m_1.2um_k141_403562 | 7796 | 8545 | - | hypothetical protein | hypothetical protein |
| LC_80m_1.2um_k141_226493_1 | LC_80m_1.2um_k141_226493 | 1 | 630 | + | hypothetical protein | hypothetical protein |
| LC_80m_1.2um_k141_226493_2 | LC_80m_1.2um_k141_226493 | 871 | 1599 | + | hypothetical protein | hypothetical protein |
| LC_80m_1.2um_k141_226493_3 | LC_80m_1.2um_k141_226493 | 1735 | 4005 | + | hypothetical protein | hypothetical protein |
| LC_80m_1.2um_k141_226493_4 | LC_80m_1.2um_k141_226493 | 4453 | 5130 | + | hypothetical protein | hypothetical protein |
| LC_80m_1.2um_k141_226493_5 | LC_80m_1.2um_k141_226493 | 5155 | 5865 | + | RDRP | RDRP |
| LC_80m_1.2um_k141_226493_6 | LC_80m_1.2um_k141_226493 | 5987 | 7015 | + | hypothetical protein | hypothetical protein |
| LC_80m_1.2um_k141_226493_7 | LC_80m_1.2um_k141_226493 | 7160 | 8170 | + | hypothetical protein | hypothetical protein |
| LC_80m_1.2um_k141_344716_1 | LC_80m_1.2um_k141_344716 | 133 | 714 | + | hypothetical protein | hypothetical protein |
| LC_80m_1.2um_k141_344716_2 | LC_80m_1.2um_k141_344716 | 1099 | 1392 | + | hypothetical protein | hypothetical protein |
| LC_80m_1.2um_k141_344716_3 | LC_80m_1.2um_k141_344716 | 1423 | 3411 | + | RDRP | RDRP |
| LC_80m_3um_k141_96140_1 | LC_80m_3um_k141_96140 | 3 | 2462 | - | RDRP | RDRP |
| LC_80m_3um_k141_96140_2 | LC_80m_3um_k141_96140 | 2553 | 2795 | + | hypothetical protein | hypothetical protein |
| LC_80m_3um_k141_96140_3 | LC_80m_3um_k141_96140 | 3003 | 3467 | - | hypothetical protein | hypothetical protein |
| LC_80m_3um_k141_96140_4 | LC_80m_3um_k141_96140 | 3486 | 3950 | - | hypothetical protein | hypothetical protein |
| LC_80m_3um_k141_63205_1 | LC_80m_3um_k141_63205 | 251 | 2476 | - | hypothetical protein | hypothetical protein |
| LC_80m_3um_k141_63205_2 | LC_80m_3um_k141_63205 | 2480 | 2995 | - | RDRP | RDRP |
| LC_80m_3um_k141_63205_3 | LC_80m_3um_k141_63205 | 3029 | 3781 | - | hypothetical protein | hypothetical protein |
| LC_80m_3um_k141_63205_4 | LC_80m_3um_k141_63205 | 3938 | 4744 | - | hypothetical protein | hypothetical protein |
| LC_80m_3um_k141_63205_5 | LC_80m_3um_k141_63205 | 4751 | 6136 | - | hypothetical protein | hypothetical protein |
| LC_80m_3um_k141_96969_1 | LC_80m_3um_k141_96969 | 42 | 1337 | - | hypothetical protein | hypothetical protein |
| LC_80m_3um_k141_96969_2 | LC_80m_3um_k141_96969 | 1404 | 2702 | - | RDRP | RDRP |
| LC_80m_3um_k141_96969_3 | LC_80m_3um_k141_96969 | 3210 | 3467 | - | hypothetical protein | hypothetical protein |
| LC_80m_3um_k141_96969_4 | LC_80m_3um_k141_96969 | 3486 | 5444 | - | hypothetical protein | hypothetical protein |
| LS_0.5m_0.2um_k141_46292_1 | LS_0.5m_0.2um_k141_46292 | 2 | 148 | + | hypothetical protein | hypothetical protein |
| LS_0.5m_0.2um_k141_46292_2 | LS_0.5m_0.2um_k141_46292 | 417 | 1994 | + | hypothetical protein | hypothetical protein |
| LS_0.5m_0.2um_k141_46292_3 | LS_0.5m_0.2um_k141_46292 | 2039 | 3544 | + | RDRP | RDRP |
| LS_0.5m_0.2um_k141_46292_4 | LS_0.5m_0.2um_k141_46292 | 3826 | 4692 | + | hypothetical protein | hypothetical protein |
| LS_0.5m_0.2um_k141_30045_1 | LS_0.5m_0.2um_k141_30045 | 2 | 193 | - | hypothetical protein | hypothetical protein |
| LS_0.5m_0.2um_k141_30045_2 | LS_0.5m_0.2um_k141_30045 | 332 | 1390 | - | RDRP | RDRP |
| LS_0.5m_0.2um_k141_30045_3 | LS_0.5m_0.2um_k141_30045 | 2033 | 2437 | - | hypothetical protein | hypothetical protein |
| LS_0.5m_10um_k141_46079_1 | LS_0.5m_10um_k141_46079 | 3 | 578 | - | RDRP | RDRP |
| LS_0.5m_10um_k141_46079_2 | LS_0.5m_10um_k141_46079 | 803 | 1294 | - | hypothetical protein | hypothetical protein |
| LS_0.5m_10um_k141_46079_3 | LS_0.5m_10um_k141_46079 | 1245 | 1898 | - | hypothetical protein | hypothetical protein |
| LS_0.5m_10um_k141_46079_4 | LS_0.5m_10um_k141_46079 | 1944 | 2099 | - | hypothetical protein | hypothetical protein |
| LC_coassembly_k141_219128_1 | LC_coassembly_k141_219128 | 2 | 2899 | + | hypothetical protein | hypothetical protein |
| LC_coassembly_k141_219128_2 | LC_coassembly_k141_219128 | 2896 | 3993 | + | hypothetical protein | hypothetical protein |
| LC_coassembly_k141_219128_3 | LC_coassembly_k141_219128 | 4116 | 7790 | + | RDRP | RDRP |
| LS_0.5m_1.2um_k141_213974_1 | LS_0.5m_1.2um_k141_213974 | 159 | 521 | - | hypothetical protein | hypothetical protein |
| LS_0.5m_1.2um_k141_213974_2 | LS_0.5m_1.2um_k141_213974 | 485 | 1048 | - | hypothetical protein | hypothetical protein |
| LS_0.5m_1.2um_k141_213974_3 | LS_0.5m_1.2um_k141_213974 | 1153 | 1272 | + | hypothetical protein | hypothetical protein |
| LS_0.5m_1.2um_k141_213974_4 | LS_0.5m_1.2um_k141_213974 | 1253 | 1948 | - | capsid/coat protein | capsid/coat protein |
| LS_0.5m_1.2um_k141_213974_5 | LS_0.5m_1.2um_k141_213974 | 2064 | 2375 | + | hypothetical protein | hypothetical protein |
| LS_0.5m_1.2um_k141_213974_6 | LS_0.5m_1.2um_k141_213974 | 2372 | 3958 | - | RDRP | RDRP |
| LS_0.5m_1.2um_k141_213974_7 | LS_0.5m_1.2um_k141_213974 | 3968 | 4963 | - | hypothetical protein | hypothetical protein |
| LS_0.5m_1.2um_k141_315047_1 | LS_0.5m_1.2um_k141_315047 | 1 | 2133 | + | hypothetical protein | hypothetical protein |
| LS_0.5m_1.2um_k141_315047_2 | LS_0.5m_1.2um_k141_315047 | 2152 | 3570 | + | RDRP | RDRP |
| LS_0.5m_1.2um_k141_315047_3 | LS_0.5m_1.2um_k141_315047 | 3637 | 3918 | + | hypothetical protein | hypothetical protein |
| LS_0.5m_1.2um_k141_585706_1 | LS_0.5m_1.2um_k141_585706 | 50 | 1687 | - | RDRP | RDRP |
| LS_0.5m_1.2um_k141_585706_2 | LS_0.5m_1.2um_k141_585706 | 1726 | 2118 | - | hypothetical protein | hypothetical protein |
| LS_0.5m_1.2um_k141_585706_3 | LS_0.5m_1.2um_k141_585706 | 2235 | 3305 | - | capsid/coat protein | capsid/coat protein |
| LS_0.5m_1.2um_k141_585706_4 | LS_0.5m_1.2um_k141_585706 | 3293 | 3415 | - | hypothetical protein | hypothetical protein |
| LS_0.5m_3um_k141_415219_1 | LS_0.5m_3um_k141_415219 | 1 | 612 | + | RDRP | RDRP |
| LS_0.5m_3um_k141_415219_2 | LS_0.5m_3um_k141_415219 | 616 | 1131 | + | hypothetical protein | hypothetical protein |
| LS_0.5m_3um_k141_415219_3 | LS_0.5m_3um_k141_415219 | 1198 | 1959 | + | hypothetical protein | hypothetical protein |
| LC_coassembly_k141_1021654_1 | LC_coassembly_k141_1021654 | 2 | 2347 | - | RDRP | RDRP |
| LC_coassembly_k141_1021654_2 | LC_coassembly_k141_1021654 | 2459 | 2818 | - | hypothetical protein | hypothetical protein |
| LC_coassembly_k141_1021654_3 | LC_coassembly_k141_1021654 | 2885 | 4600 | - | hypothetical protein | hypothetical protein |
| LC_coassembly_k141_1021654_4 | LC_coassembly_k141_1021654 | 4646 | 6862 | - | hypothetical protein | hypothetical protein |
| LC_coassembly_k141_1021654_5 | LC_coassembly_k141_1021654 | 7079 | 8650 | + | hypothetical protein | hypothetical protein |
| LC_coassembly_k141_1021654_6 | LC_coassembly_k141_1021654 | 8768 | 11623 | + | hypothetical protein | hypothetical protein |
| LC_coassembly_k141_1021654_7 | LC_coassembly_k141_1021654 | 11756 | 13177 | + | hypothetical protein | hypothetical protein |
| LC_coassembly_k141_1021654_8 | LC_coassembly_k141_1021654 | 13199 | 13423 | + | hypothetical protein | hypothetical protein |
| LC_coassembly_k141_1021654_9 | LC_coassembly_k141_1021654 | 13424 | 14734 | + | RDRP | RDRP |
| LC_coassembly_k141_581270_1 | LC_coassembly_k141_581270 | 1 | 2001 | - | hypothetical protein | hypothetical protein |
| LC_coassembly_k141_581270_2 | LC_coassembly_k141_581270 | 2002 | 3288 | - | RDRP | RDRP |
| LC_coassembly_k141_581270_3 | LC_coassembly_k141_581270 | 3415 | 4050 | - | hypothetical protein | hypothetical protein |
| LC_coassembly_k141_581270_4 | LC_coassembly_k141_581270 | 4273 | 6498 | - | hypothetical protein | hypothetical protein |
| LC_coassembly_k141_581270_5 | LC_coassembly_k141_581270 | 6499 | 6753 | - | hypothetical protein | hypothetical protein |
| LC_coassembly_k141_101176_1 | LC_coassembly_k141_101176 | 2 | 790 | - | hypothetical protein | hypothetical protein |
| LC_coassembly_k141_101176_2 | LC_coassembly_k141_101176 | 871 | 1692 | - | hypothetical protein | hypothetical protein |
| LC_coassembly_k141_101176_3 | LC_coassembly_k141_101176 | 2008 | 3933 | - | RDRP | RDRP |
| LC_coassembly_k141_101176_4 | LC_coassembly_k141_101176 | 4014 | 4835 | - | hypothetical protein | hypothetical protein |
| LC_coassembly_k141_101176_5 | LC_coassembly_k141_101176 | 5151 | 6638 | - | RDRP | RDRP |
| LC_coassembly_k141_1064585_1 | LC_coassembly_k141_1064585 | 2 | 484 | + | hypothetical protein | hypothetical protein |
| LC_coassembly_k141_1064585_2 | LC_coassembly_k141_1064585 | 869 | 1162 | + | hypothetical protein | hypothetical protein |
| LC_coassembly_k141_1064585_3 | LC_coassembly_k141_1064585 | 1193 | 3181 | + | RDRP | RDRP |
| LC_coassembly_k141_1064585_4 | LC_coassembly_k141_1064585 | 3263 | 4273 | + | hypothetical protein | hypothetical protein |
| LC_coassembly_k141_2206238_1 | LC_coassembly_k141_2206238 | 3 | 2255 | - | RDRP | RDRP |
| LC_coassembly_k141_2206238_2 | LC_coassembly_k141_2206238 | 2239 | 3450 | - | hypothetical protein | hypothetical protein |
| LC_coassembly_k141_2206238_3 | LC_coassembly_k141_2206238 | 3447 | 3992 | - | hypothetical protein | hypothetical protein |
| LC_coassembly_k141_31205_1 | LC_coassembly_k141_31205 | 232 | 507 | - | hypothetical protein | hypothetical protein |
| LC_coassembly_k141_31205_2 | LC_coassembly_k141_31205 | 523 | 735 | + | hypothetical protein | hypothetical protein |
| LC_coassembly_k141_31205_3 | LC_coassembly_k141_31205 | 757 | 2556 | + | RDRP | RDRP |
| LC_coassembly_k141_1727070_1 | LC_coassembly_k141_1727070 | 3 | 260 | + | hypothetical protein | hypothetical protein |
| LC_coassembly_k141_1727070_2 | LC_coassembly_k141_1727070 | 303 | 4283 | + | hypothetical protein | hypothetical protein |
| LC_coassembly_k141_1727070_3 | LC_coassembly_k141_1727070 | 4302 | 5102 | + | hypothetical protein | hypothetical protein |
| LC_coassembly_k141_1727070_4 | LC_coassembly_k141_1727070 | 5069 | 5392 | + | hypothetical protein | hypothetical protein |
| LC_coassembly_k141_1727070_5 | LC_coassembly_k141_1727070 | 5744 | 6232 | + | hypothetical protein | hypothetical protein |
| LC_coassembly_k141_1727070_6 | LC_coassembly_k141_1727070 | 6239 | 6871 | + | RDRP | RDRP |
| LC_coassembly_k141_1727070_7 | LC_coassembly_k141_1727070 | 7169 | 7624 | + | hypothetical protein | hypothetical protein |
| LC_coassembly_k141_526161_1 | LC_coassembly_k141_526161 | 165 | 545 | + | hypothetical protein | hypothetical protein |
| LC_coassembly_k141_526161_2 | LC_coassembly_k141_526161 | 595 | 1602 | + | hypothetical protein | hypothetical protein |
| LC_coassembly_k141_526161_3 | LC_coassembly_k141_526161 | 1653 | 3461 | + | RDRP | RDRP |
| LC_coassembly_k141_75697_1 | LC_coassembly_k141_75697 | 556 | 1566 | - | hypothetical protein | hypothetical protein |
| LC_coassembly_k141_75697_2 | LC_coassembly_k141_75697 | 1648 | 2640 | - | hypothetical protein | hypothetical protein |
| LC_coassembly_k141_75697_3 | LC_coassembly_k141_75697 | 2779 | 3960 | - | RDRP | RDRP |
| LC_coassembly_k141_75697_4 | LC_coassembly_k141_75697 | 4480 | 4842 | - | hypothetical protein | hypothetical protein |
| LC_coassembly_k141_75697_5 | LC_coassembly_k141_75697 | 5047 | 7764 | - | hypothetical protein | hypothetical protein |
| LC_coassembly_k141_75697_6 | LC_coassembly_k141_75697 | 7942 | 8592 | - | hypothetical protein | hypothetical protein |
| LC_coassembly_k141_947088_1 | LC_coassembly_k141_947088 | 418 | 1080 | + | hypothetical protein | hypothetical protein |
| LC_coassembly_k141_947088_2 | LC_coassembly_k141_947088 | 1128 | 2600 | + | RDRP | RDRP |
| LC_coassembly_k141_947088_3 | LC_coassembly_k141_947088 | 2546 | 3280 | + | hypothetical protein | hypothetical protein |
| LC_coassembly_k141_2418593_1 | LC_coassembly_k141_2418593 | 3 | 1007 | + | hypothetical protein | hypothetical protein |
| LC_coassembly_k141_2418593_2 | LC_coassembly_k141_2418593 | 1359 | 1763 | + | hypothetical protein | hypothetical protein |
| LC_coassembly_k141_2418593_3 | LC_coassembly_k141_2418593 | 1815 | 2552 | + | RDRP | RDRP |
| LC_coassembly_k141_2418593_4 | LC_coassembly_k141_2418593 | 2580 | 3239 | + | hypothetical protein | hypothetical protein |
| LC_coassembly_k141_1759012_1 | LC_coassembly_k141_1759012 | 433 | 3756 | + | hypothetical protein | hypothetical protein |
| LC_coassembly_k141_1759012_2 | LC_coassembly_k141_1759012 | 4059 | 4244 | - | hypothetical protein | hypothetical protein |
| LC_coassembly_k141_1759012_3 | LC_coassembly_k141_1759012 | 4312 | 4806 | + | hypothetical protein | hypothetical protein |
| LC_coassembly_k141_1759012_4 | LC_coassembly_k141_1759012 | 5044 | 8271 | + | RDRP | RDRP |
| LC_coassembly_k141_2599729_1 | LC_coassembly_k141_2599729 | 1 | 123 | + | hypothetical protein | hypothetical protein |
| LC_coassembly_k141_2599729_2 | LC_coassembly_k141_2599729 | 475 | 1476 | + | RDRP | RDRP |
| LC_coassembly_k141_2599729_3 | LC_coassembly_k141_2599729 | 1780 | 2355 | + | hypothetical protein | hypothetical protein |
| LC_coassembly_k141_76989_1 | LC_coassembly_k141_76989 | 1 | 1041 | - | capsid/coat protein | capsid/coat protein |
| LC_coassembly_k141_76989_2 | LC_coassembly_k141_76989 | 1038 | 2546 | - | RDRP | RDRP |
| LC_coassembly_k141_76989_3 | LC_coassembly_k141_76989 | 2643 | 3728 | - | hypothetical protein | hypothetical protein |
| LC_coassembly_k141_1773142_1 | LC_coassembly_k141_1773142 | 213 | 878 | - | capsid/coat protein | capsid/coat protein |
| LC_coassembly_k141_1773142_2 | LC_coassembly_k141_1773142 | 894 | 2393 | - | RDRP | RDRP |
| LC_coassembly_k141_1773142_3 | LC_coassembly_k141_1773142 | 2412 | 2930 | - | RDRP | RDRP |
| LC_coassembly_k141_2331475_1 | LC_coassembly_k141_2331475 | 278 | 1918 | - | RDRP | RDRP |
| LC_coassembly_k141_2331475_2 | LC_coassembly_k141_2331475 | 1936 | 4041 | - | hypothetical protein | hypothetical protein |
| LC_coassembly_k141_2331475_3 | LC_coassembly_k141_2331475 | 4913 | 6055 | - | capsid/coat protein | capsid/coat protein |
| LC_coassembly_k141_860293_1 | LC_coassembly_k141_860293 | 41 | 625 | - | hypothetical protein | hypothetical protein |
| LC_coassembly_k141_860293_2 | LC_coassembly_k141_860293 | 663 | 899 | - | hypothetical protein | hypothetical protein |
| LC_coassembly_k141_860293_3 | LC_coassembly_k141_860293 | 901 | 3000 | - | RDRP | RDRP |
| LC_coassembly_k141_592818_1 | LC_coassembly_k141_592818 | 59 | 2233 | - | RDRP | RDRP |
| LC_coassembly_k141_592818_2 | LC_coassembly_k141_592818 | 2248 | 3078 | - | hypothetical protein | hypothetical protein |
| LC_coassembly_k141_592818_3 | LC_coassembly_k141_592818 | 3093 | 3278 | - | hypothetical protein | hypothetical protein |
| LC_coassembly_k141_1526117_1 | LC_coassembly_k141_1526117 | 259 | 2832 | - | RDRP | RDRP |
| LC_coassembly_k141_1526117_2 | LC_coassembly_k141_1526117 | 2854 | 7509 | - | hypothetical protein | hypothetical protein |
| LC_coassembly_k141_1526117_3 | LC_coassembly_k141_1526117 | 7574 | 8710 | - | hypothetical protein | hypothetical protein |
| LC_coassembly_k141_204657_1 | LC_coassembly_k141_204657 | 45 | 1526 | + | hypothetical protein | hypothetical protein |
| LC_coassembly_k141_204657_2 | LC_coassembly_k141_204657 | 1478 | 2986 | + | RDRP | RDRP |
| LC_coassembly_k141_204657_3 | LC_coassembly_k141_204657 | 3216 | 4274 | + | capsid/coat protein | capsid/coat protein |
| LC_coassembly_k141_843059_1 | LC_coassembly_k141_843059 | 2 | 661 | - | RDRP | RDRP |
| LC_coassembly_k141_843059_2 | LC_coassembly_k141_843059 | 695 | 4339 | - | hypothetical protein | hypothetical protein |
| LC_coassembly_k141_843059_3 | LC_coassembly_k141_843059 | 4355 | 7504 | - | hypothetical protein | hypothetical protein |
| LC_coassembly_k141_2187462_1 | LC_coassembly_k141_2187462 | 279 | 518 | - | hypothetical protein | hypothetical protein |
| LC_coassembly_k141_2187462_2 | LC_coassembly_k141_2187462 | 457 | 3231 | - | RDRP | RDRP |
| LC_coassembly_k141_2187462_3 | LC_coassembly_k141_2187462 | 3231 | 5378 | - | hypothetical protein | hypothetical protein |
| LC_coassembly_k141_537252_1 | LC_coassembly_k141_537252 | 1000 | 1983 | - | hypothetical protein | hypothetical protein |
| LC_coassembly_k141_537252_2 | LC_coassembly_k141_537252 | 2146 | 2760 | - | hypothetical protein | hypothetical protein |
| LC_coassembly_k141_537252_3 | LC_coassembly_k141_537252 | 2767 | 3327 | - | RDRP | RDRP |
| LC_coassembly_k141_537252_4 | LC_coassembly_k141_537252 | 3460 | 3750 | - | RDRP | RDRP |
| LC_coassembly_k141_537252_5 | LC_coassembly_k141_537252 | 3832 | 6912 | - | hypothetical protein | hypothetical protein |
| LC_coassembly_k141_537252_6 | LC_coassembly_k141_537252 | 6958 | 8019 | - | hypothetical protein | hypothetical protein |
| LC_coassembly_k141_537252_7 | LC_coassembly_k141_537252 | 8257 | 8763 | - | hypothetical protein | hypothetical protein |
| LC_coassembly_k141_537252_8 | LC_coassembly_k141_537252 | 8764 | 9579 | - | hypothetical protein | hypothetical protein |
| LC_coassembly_k141_537252_9 | LC_coassembly_k141_537252 | 9703 | 11355 | - | hypothetical protein | hypothetical protein |
| LC_coassembly_k141_2164532_1 | LC_coassembly_k141_2164532 | 1 | 744 | + | hypothetical protein | hypothetical protein |
| LC_coassembly_k141_2164532_2 | LC_coassembly_k141_2164532 | 760 | 1509 | + | hypothetical protein | hypothetical protein |
| LC_coassembly_k141_2164532_3 | LC_coassembly_k141_2164532 | 1576 | 4404 | + | RDRP | RDRP |
| LC_coassembly_k141_2164532_4 | LC_coassembly_k141_2164532 | 4438 | 5565 | + | hypothetical protein | hypothetical protein |
| LC_coassembly_k141_1259623_1 | LC_coassembly_k141_1259623 | 1 | 804 | - | hypothetical protein | hypothetical protein |
| LC_coassembly_k141_1259623_2 | LC_coassembly_k141_1259623 | 871 | 2169 | - | RDRP | RDRP |
| LC_coassembly_k141_1259623_3 | LC_coassembly_k141_1259623 | 2633 | 2791 | + | hypothetical protein | hypothetical protein |
| LC_coassembly_k141_1259623_4 | LC_coassembly_k141_1259623 | 3139 | 6003 | - | hypothetical protein | hypothetical protein |
| LC_coassembly_k141_1259623_5 | LC_coassembly_k141_1259623 | 6142 | 6678 | - | hypothetical protein | hypothetical protein |
| LC_coassembly_k141_1350294_1 | LC_coassembly_k141_1350294 | 529 | 1539 | - | hypothetical protein | hypothetical protein |
| LC_coassembly_k141_1350294_2 | LC_coassembly_k141_1350294 | 1684 | 3546 | - | RDRP | RDRP |
| LC_coassembly_k141_1350294_3 | LC_coassembly_k141_1350294 | 3571 | 4248 | - | hypothetical protein | hypothetical protein |
| LC_coassembly_k141_1350294_4 | LC_coassembly_k141_1350294 | 4405 | 4683 | - | hypothetical protein | hypothetical protein |
| LC_coassembly_k141_1350294_5 | LC_coassembly_k141_1350294 | 4696 | 6966 | - | hypothetical protein | hypothetical protein |
| LC_coassembly_k141_1350294_6 | LC_coassembly_k141_1350294 | 7102 | 7830 | - | hypothetical protein | hypothetical protein |
| LC_coassembly_k141_1350294_7 | LC_coassembly_k141_1350294 | 8071 | 8742 | - | hypothetical protein | hypothetical protein |
| LC_coassembly_k141_779785_1 | LC_coassembly_k141_779785 | 379 | 597 | - | hypothetical protein | hypothetical protein |
| LC_coassembly_k141_779785_2 | LC_coassembly_k141_779785 | 652 | 1557 | - | hypothetical protein | hypothetical protein |
| LC_coassembly_k141_779785_3 | LC_coassembly_k141_779785 | 2560 | 4707 | - | RDRP | RDRP |
| LC_coassembly_k141_2401611_1 | LC_coassembly_k141_2401611 | 3 | 3044 | + | hypothetical protein | hypothetical protein |
| LC_coassembly_k141_2401611_2 | LC_coassembly_k141_2401611 | 3062 | 5389 | + | hypothetical protein | hypothetical protein |
| LC_coassembly_k141_2401611_3 | LC_coassembly_k141_2401611 | 5480 | 5596 | + | hypothetical protein | hypothetical protein |
| LC_coassembly_k141_2401611_4 | LC_coassembly_k141_2401611 | 5624 | 6694 | + | RDRP | RDRP |
| LC_coassembly_k141_2401611_5 | LC_coassembly_k141_2401611 | 7055 | 7621 | + | hypothetical protein | hypothetical protein |
| LS_coassembly_k141_656443_1 | LS_coassembly_k141_656443 | 131 | 1483 | + | capsid/coat protein | capsid/coat protein |
| LS_coassembly_k141_656443_2 | LS_coassembly_k141_656443 | 1597 | 1899 | + | hypothetical protein | hypothetical protein |
| LS_coassembly_k141_656443_3 | LS_coassembly_k141_656443 | 1984 | 3771 | + | RDRP | RDRP |
| LS_coassembly_k141_57787_1 | LS_coassembly_k141_57787 | 169 | 1164 | + | hypothetical protein | hypothetical protein |
| LS_coassembly_k141_57787_2 | LS_coassembly_k141_57787 | 1174 | 2760 | + | RDRP | RDRP |
| LS_coassembly_k141_57787_3 | LS_coassembly_k141_57787 | 2757 | 3068 | - | hypothetical protein | hypothetical protein |
| LS_coassembly_k141_57787_4 | LS_coassembly_k141_57787 | 3184 | 3879 | + | capsid/coat protein | capsid/coat protein |
| LS_coassembly_k141_57787_5 | LS_coassembly_k141_57787 | 3860 | 3979 | - | hypothetical protein | hypothetical protein |
| LS_coassembly_k141_57787_6 | LS_coassembly_k141_57787 | 4084 | 4647 | + | hypothetical protein | hypothetical protein |
| LS_coassembly_k141_57787_7 | LS_coassembly_k141_57787 | 4611 | 4973 | + | hypothetical protein | hypothetical protein |
| LS_coassembly_k141_543237_1 | LS_coassembly_k141_543237 | 3 | 1454 | + | capsid/coat protein | capsid/coat protein |
| LS_coassembly_k141_543237_2 | LS_coassembly_k141_543237 | 1454 | 1849 | + | hypothetical protein | hypothetical protein |
| LS_coassembly_k141_543237_3 | LS_coassembly_k141_543237 | 1960 | 3672 | + | RDRP | RDRP |
| LS_coassembly_k141_1028965_1 | LS_coassembly_k141_1028965 | 1 | 723 | + | hypothetical protein | hypothetical protein |
| LS_coassembly_k141_1028965_2 | LS_coassembly_k141_1028965 | 789 | 992 | + | hypothetical protein | hypothetical protein |
| LS_coassembly_k141_1028965_3 | LS_coassembly_k141_1028965 | 952 | 2388 | + | RDRP | RDRP |
| LS_coassembly_k141_1028965_4 | LS_coassembly_k141_1028965 | 2415 | 2594 | + | hypothetical protein | hypothetical protein |
| LS_coassembly_k141_818259_1 | LS_coassembly_k141_818259 | 185 | 895 | + | hypothetical protein | hypothetical protein |
| LS_coassembly_k141_818259_2 | LS_coassembly_k141_818259 | 892 | 1554 | + | hypothetical protein | hypothetical protein |
| LS_coassembly_k141_818259_3 | LS_coassembly_k141_818259 | 1555 | 2304 | + | hypothetical protein | hypothetical protein |
| LS_coassembly_k141_818259_4 | LS_coassembly_k141_818259 | 2301 | 3392 | + | hypothetical protein | hypothetical protein |
| LS_coassembly_k141_818259_5 | LS_coassembly_k141_818259 | 3331 | 3552 | + | hypothetical protein | hypothetical protein |
| LS_coassembly_k141_818259_6 | LS_coassembly_k141_818259 | 3986 | 11068 | + | RDRP | RDRP |
| LS_coassembly_k141_63722_1 | LS_coassembly_k141_63722 | 39 | 161 | + | hypothetical protein | hypothetical protein |
| LS_coassembly_k141_63722_2 | LS_coassembly_k141_63722 | 149 | 1219 | + | capsid/coat protein | capsid/coat protein |
| LS_coassembly_k141_63722_3 | LS_coassembly_k141_63722 | 1327 | 1728 | + | hypothetical protein | hypothetical protein |
| LS_coassembly_k141_63722_4 | LS_coassembly_k141_63722 | 1767 | 3404 | + | RDRP | RDRP |
| LS_coassembly_k141_891738_1 | LS_coassembly_k141_891738 | 1 | 324 | + | hypothetical protein | hypothetical protein |
| LS_coassembly_k141_891738_2 | LS_coassembly_k141_891738 | 385 | 696 | + | RDRP | RDRP |
| LS_coassembly_k141_891738_3 | LS_coassembly_k141_891738 | 946 | 1596 | + | RDRP | RDRP |
| LS_coassembly_k141_891738_4 | LS_coassembly_k141_891738 | 1612 | 1833 | + | hypothetical protein | hypothetical protein |
| LS_coassembly_k141_891738_5 | LS_coassembly_k141_891738 | 1846 | 2247 | + | hypothetical protein | hypothetical protein |
| LS_coassembly_k141_165614_1 | LS_coassembly_k141_165614 | 1 | 156 | + | hypothetical protein | hypothetical protein |
| LS_coassembly_k141_165614_2 | LS_coassembly_k141_165614 | 202 | 1296 | + | hypothetical protein | hypothetical protein |
| LS_coassembly_k141_165614_3 | LS_coassembly_k141_165614 | 1521 | 3191 | + | RDRP | RDRP |
| LS_coassembly_k141_165614_4 | LS_coassembly_k141_165614 | 3424 | 3726 | + | hypothetical protein | hypothetical protein |
| LS_coassembly_k141_379730_1 | LS_coassembly_k141_379730 | 72 | 2303 | - | RDRP | RDRP |
| LS_coassembly_k141_379730_2 | LS_coassembly_k141_379730 | 2362 | 2709 | - | hypothetical protein | hypothetical protein |
| LS_coassembly_k141_379730_3 | LS_coassembly_k141_379730 | 2745 | 4223 | - | capsid/coat protein | capsid/coat protein |
| LS_coassembly_k141_8864_1 | LS_coassembly_k141_8864 | 3 | 1835 | - | RDRP | RDRP |
| LS_coassembly_k141_8864_2 | LS_coassembly_k141_8864 | 1877 | 2320 | - | hypothetical protein | hypothetical protein |
| LS_coassembly_k141_8864_3 | LS_coassembly_k141_8864 | 2239 | 3378 | - | capsid/coat protein | capsid/coat protein |
| LC_coassembly_k141_2045611_1 | LC_coassembly_k141_2045611 | 1 | 3540 | + | hypothetical protein | hypothetical protein |
| LC_coassembly_k141_2045611_2 | LC_coassembly_k141_2045611 | 3541 | 4392 | + | hypothetical protein | hypothetical protein |
| LC_coassembly_k141_2045611_3 | LC_coassembly_k141_2045611 | 4519 | 4704 | + | hypothetical protein | hypothetical protein |
| LC_coassembly_k141_2045611_4 | LC_coassembly_k141_2045611 | 4750 | 5805 | + | RDRP | RDRP |
| LC_coassembly_k141_2045611_5 | LC_coassembly_k141_2045611 | 5806 | 6879 | + | hypothetical protein | hypothetical protein |
| LC_coassembly_k141_2045611_6 | LC_coassembly_k141_2045611 | 6934 | 8034 | + | hypothetical protein | hypothetical protein |
| LS_coassembly_k141_765865_1 | LS_coassembly_k141_765865 | 123 | 719 | - | capsid/coat protein | capsid/coat protein |
| LS_coassembly_k141_765865_2 | LS_coassembly_k141_765865 | 796 | 2304 | - | RDRP | RDRP |
| LS_coassembly_k141_765865_3 | LS_coassembly_k141_765865 | 2362 | 2892 | - | RDRP | RDRP |
| LS_coassembly_k141_1250841_1 | LS_coassembly_k141_1250841 | 3 | 1622 | - | RDRP | RDRP |
| LS_coassembly_k141_1250841_2 | LS_coassembly_k141_1250841 | 1619 | 2896 | - | hypothetical protein | hypothetical protein |
| LS_coassembly_k141_1250841_3 | LS_coassembly_k141_1250841 | 4446 | 4715 | - | hypothetical protein | hypothetical protein |
| LS_coassembly_k141_638690_1 | LS_coassembly_k141_638690 | 85 | 1965 | - | RDRP | RDRP |
| LS_coassembly_k141_638690_2 | LS_coassembly_k141_638690 | 2141 | 2521 | - | hypothetical protein | hypothetical protein |
| LS_coassembly_k141_638690_3 | LS_coassembly_k141_638690 | 2569 | 4032 | - | hypothetical protein | hypothetical protein |
| LS_coassembly_k141_738808_1 | LS_coassembly_k141_738808 | 59 | 1912 | - | RDRP | RDRP |
| LS_coassembly_k141_738808_2 | LS_coassembly_k141_738808 | 1925 | 4705 | - | hypothetical protein | hypothetical protein |
| LS_coassembly_k141_738808_3 | LS_coassembly_k141_738808 | 4876 | 6117 | - | hypothetical protein | hypothetical protein |
| LS_coassembly_k141_824362_1 | LS_coassembly_k141_824362 | 95 | 1864 | - | RDRP | RDRP |
| LS_coassembly_k141_824362_2 | LS_coassembly_k141_824362 | 1923 | 2306 | - | hypothetical protein | hypothetical protein |
| LS_coassembly_k141_824362_3 | LS_coassembly_k141_824362 | 2345 | 2872 | - | capsid/coat protein | capsid/coat protein |
| LS_coassembly_k141_268503_1 | LS_coassembly_k141_268503 | 373 | 6372 | + | RDRP | RDRP |
| LS_coassembly_k141_268503_2 | LS_coassembly_k141_268503 | 6520 | 9483 | + | capsid/coat protein | capsid/coat protein |
| LS_coassembly_k141_268503_3 | LS_coassembly_k141_268503 | 9519 | 9836 | + | capsid/coat protein | capsid/coat protein |
| LS_coassembly_k141_268503_4 | LS_coassembly_k141_268503 | 9833 | 10051 | - | hypothetical protein | hypothetical protein |
| LS_coassembly_k141_981221_1 | LS_coassembly_k141_981221 | 3 | 1487 | + | capsid/coat protein | capsid/coat protein |
| LS_coassembly_k141_981221_2 | LS_coassembly_k141_981221 | 1524 | 1904 | + | hypothetical protein | hypothetical protein |
| LS_coassembly_k141_981221_3 | LS_coassembly_k141_981221 | 1938 | 3761 | + | RDRP | RDRP |
| LS_coassembly_k141_1166806_1 | LS_coassembly_k141_1166806 | 14 | 1750 | - | RDRP | RDRP |
| LS_coassembly_k141_1166806_2 | LS_coassembly_k141_1166806 | 1806 | 2159 | - | hypothetical protein | hypothetical protein |
| LS_coassembly_k141_1166806_3 | LS_coassembly_k141_1166806 | 2231 | 3649 | - | capsid/coat protein | capsid/coat protein |
| LS_coassembly_k141_839853_1 | LS_coassembly_k141_839853 | 1 | 102 | - | hypothetical protein | hypothetical protein |
| LS_coassembly_k141_839853_2 | LS_coassembly_k141_839853 | 208 | 2136 | - | RDRP | RDRP |
| LS_coassembly_k141_839853_3 | LS_coassembly_k141_839853 | 2239 | 2571 | - | hypothetical protein | hypothetical protein |
| LS_coassembly_k141_839853_4 | LS_coassembly_k141_839853 | 2653 | 4032 | - | capsid/coat protein | capsid/coat protein |
| LS_coassembly_k141_84850_1 | LS_coassembly_k141_84850 | 429 | 2639 | - | capsid/coat protein | capsid/coat protein |
| LS_coassembly_k141_84850_2 | LS_coassembly_k141_84850 | 2677 | 2925 | + | hypothetical protein | hypothetical protein |
| LS_coassembly_k141_84850_3 | LS_coassembly_k141_84850 | 3098 | 7813 | - | RDRP | RDRP |
| LC_0.5m_10um_k141_200615_1 | LC_0.5m_10um_k141_200615 | 83 | 4333 | + | hypothetical protein | hypothetical protein |
| LC_0.5m_10um_k141_200615_2 | LC_0.5m_10um_k141_200615 | 4475 | 6805 | + | RDRP | RDRP |
| LC_0.5m_10um_k141_200615_3 | LC_0.5m_10um_k141_200615 | 6872 | 7894 | + | hypothetical protein | hypothetical protein |
| LC_coassembly_k141_1111088_1 | LC_coassembly_k141_1111088 | 440 | 2668 | - | hypothetical protein | hypothetical protein |
| LC_coassembly_k141_1111088_2 | LC_coassembly_k141_1111088 | 2669 | 3955 | - | RDRP | RDRP |
| LC_coassembly_k141_1111088_3 | LC_coassembly_k141_1111088 | 4082 | 4933 | - | hypothetical protein | hypothetical protein |
| LC_coassembly_k141_1111088_4 | LC_coassembly_k141_1111088 | 4934 | 8446 | - | hypothetical protein | hypothetical protein |
| LC_0.5m_10um_k141_14159_1 | LC_0.5m_10um_k141_14159 | 415 | 1437 | - | hypothetical protein | hypothetical protein |
| LC_0.5m_10um_k141_14159_2 | LC_0.5m_10um_k141_14159 | 1504 | 3366 | - | RDRP | RDRP |
| LC_0.5m_10um_k141_14159_3 | LC_0.5m_10um_k141_14159 | 3388 | 3600 | - | hypothetical protein | hypothetical protein |
| LC_0.5m_10um_k141_14159_4 | LC_0.5m_10um_k141_14159 | 3616 | 3891 | + | hypothetical protein | hypothetical protein |
| LC_0.5m_10um_k141_14159_5 | LC_0.5m_10um_k141_14159 | 3976 | 5946 | - | hypothetical protein | hypothetical protein |
| LC_0.5m_10um_k141_14159_6 | LC_0.5m_10um_k141_14159 | 5953 | 6429 | - | hypothetical protein | hypothetical protein |
| LC_0.5m_10um_k141_14159_7 | LC_0.5m_10um_k141_14159 | 6451 | 8166 | - | hypothetical protein | hypothetical protein |
| LC_0.5m_10um_k141_129489_1 | LC_0.5m_10um_k141_129489 | 2 | 790 | + | hypothetical protein | hypothetical protein |
| LC_0.5m_10um_k141_129489_2 | LC_0.5m_10um_k141_129489 | 932 | 3259 | + | RDRP | RDRP |
| LC_0.5m_10um_k141_129489_3 | LC_0.5m_10um_k141_129489 | 3326 | 4348 | + | hypothetical protein | hypothetical protein |
| LC_0.5m_10um_k141_153862_1 | LC_0.5m_10um_k141_153862 | 3 | 599 | - | hypothetical protein | hypothetical protein |
| LC_0.5m_10um_k141_153862_2 | LC_0.5m_10um_k141_153862 | 627 | 1364 | - | RDRP | RDRP |
| LC_0.5m_10um_k141_153862_3 | LC_0.5m_10um_k141_153862 | 1416 | 1820 | - | hypothetical protein | hypothetical protein |
| LC_0.5m_10um_k141_153862_4 | LC_0.5m_10um_k141_153862 | 2172 | 3071 | - | hypothetical protein | hypothetical protein |
| LC_0.5m_1.2um_k141_276576_1 | LC_0.5m_1.2um_k141_276576 | 344 | 550 | - | hypothetical protein | hypothetical protein |
| LC_0.5m_1.2um_k141_276576_2 | LC_0.5m_1.2um_k141_276576 | 556 | 1404 | - | hypothetical protein | hypothetical protein |
| LC_0.5m_1.2um_k141_276576_3 | LC_0.5m_1.2um_k141_276576 | 2429 | 4669 | - | RDRP | RDRP |
| LC_0.5m_1.2um_k141_36111_1 | LC_0.5m_1.2um_k141_36111 | 2 | 754 | + | hypothetical protein | hypothetical protein |
| LC_0.5m_1.2um_k141_36111_2 | LC_0.5m_1.2um_k141_36111 | 788 | 3529 | + | hypothetical protein | hypothetical protein |
| LC_0.5m_1.2um_k141_36111_3 | LC_0.5m_1.2um_k141_36111 | 3827 | 4096 | + | hypothetical protein | hypothetical protein |
| LC_0.5m_1.2um_k141_36111_4 | LC_0.5m_1.2um_k141_36111 | 4499 | 4909 | + | hypothetical protein | hypothetical protein |
| LC_0.5m_1.2um_k141_36111_5 | LC_0.5m_1.2um_k141_36111 | 4940 | 6928 | + | RDRP | RDRP |
| LC_0.5m_1.2um_k141_36111_6 | LC_0.5m_1.2um_k141_36111 | 7010 | 7219 | + | hypothetical protein | hypothetical protein |
| LC_0.5m_1.2um_k141_403526_1 | LC_0.5m_1.2um_k141_403526 | 3 | 527 | + | hypothetical protein | hypothetical protein |
| LC_0.5m_1.2um_k141_403526_2 | LC_0.5m_1.2um_k141_403526 | 795 | 2096 | + | RDRP | RDRP |
| LC_0.5m_1.2um_k141_403526_3 | LC_0.5m_1.2um_k141_403526 | 2163 | 3551 | + | hypothetical protein | hypothetical protein |
| LC_coassembly_k141_1296109_1 | LC_coassembly_k141_1296109 | 2 | 1858 | + | hypothetical protein | hypothetical protein |
| LC_coassembly_k141_1296109_2 | LC_coassembly_k141_1296109 | 2084 | 3550 | + | hypothetical protein | hypothetical protein |
| LC_coassembly_k141_1296109_3 | LC_coassembly_k141_1296109 | 3557 | 4252 | + | hypothetical protein | hypothetical protein |
| LC_coassembly_k141_1296109_4 | LC_coassembly_k141_1296109 | 4421 | 5821 | + | RDRP | RDRP |
| LC_coassembly_k141_1296109_5 | LC_coassembly_k141_1296109 | 5822 | 8050 | + | hypothetical protein | hypothetical protein |
| LC_0.5m_1.2um_k141_20885_1 | LC_0.5m_1.2um_k141_20885 | 1 | 1161 | - | hypothetical protein | hypothetical protein |
| LC_0.5m_1.2um_k141_20885_2 | LC_0.5m_1.2um_k141_20885 | 1162 | 2448 | - | RDRP | RDRP |
| LC_0.5m_1.2um_k141_20885_3 | LC_0.5m_1.2um_k141_20885 | 3006 | 3425 | - | hypothetical protein | hypothetical protein |
| LC_0.5m_1.2um_k141_20885_4 | LC_0.5m_1.2um_k141_20885 | 3432 | 4898 | - | hypothetical protein | hypothetical protein |
| LC_0.5m_1.2um_k141_20885_5 | LC_0.5m_1.2um_k141_20885 | 5124 | 6302 | - | hypothetical protein | hypothetical protein |
| LC_0.5m_1.2um_k141_20885_6 | LC_0.5m_1.2um_k141_20885 | 6429 | 6953 | - | hypothetical protein | hypothetical protein |
| LC_0.5m_3um_k141_255266_1 | LC_0.5m_3um_k141_255266 | 73 | 429 | + | hypothetical protein | hypothetical protein |
| LC_0.5m_3um_k141_255266_2 | LC_0.5m_3um_k141_255266 | 317 | 772 | - | hypothetical protein | hypothetical protein |
| LC_0.5m_3um_k141_255266_3 | LC_0.5m_3um_k141_255266 | 1088 | 2191 | - | RDRP | RDRP |
| LC_0.5m_3um_k141_311310_1 | LC_0.5m_3um_k141_311310 | 443 | 1453 | - | hypothetical protein | hypothetical protein |
| LC_0.5m_3um_k141_311310_2 | LC_0.5m_3um_k141_311310 | 1535 | 2527 | - | hypothetical protein | hypothetical protein |
| LC_0.5m_3um_k141_311310_3 | LC_0.5m_3um_k141_311310 | 2666 | 3298 | - | RDRP | RDRP |
| LC_0.5m_3um_k141_163067_1 | LC_0.5m_3um_k141_163067 | 3 | 797 | + | hypothetical protein | hypothetical protein |
| LC_0.5m_3um_k141_163067_2 | LC_0.5m_3um_k141_163067 | 801 | 2036 | + | RDRP | RDRP |
| LC_0.5m_3um_k141_163067_3 | LC_0.5m_3um_k141_163067 | 2089 | 2217 | + | hypothetical protein | hypothetical protein |
| LC_0.5m_3um_k141_265610_1 | LC_0.5m_3um_k141_265610 | 3 | 632 | + | hypothetical protein | hypothetical protein |
| LC_0.5m_3um_k141_265610_2 | LC_0.5m_3um_k141_265610 | 810 | 1709 | + | hypothetical protein | hypothetical protein |
| LC_0.5m_3um_k141_265610_3 | LC_0.5m_3um_k141_265610 | 1995 | 3527 | + | hypothetical protein | hypothetical protein |
| LC_0.5m_3um_k141_265610_4 | LC_0.5m_3um_k141_265610 | 3648 | 4196 | + | hypothetical protein | hypothetical protein |
| LC_0.5m_3um_k141_265610_5 | LC_0.5m_3um_k141_265610 | 4938 | 6926 | + | RDRP | RDRP |
| LC_0.5m_3um_k141_265610_6 | LC_0.5m_3um_k141_265610 | 7008 | 8018 | + | hypothetical protein | hypothetical protein |
| LC_0.5m_3um_k141_350484_1 | LC_0.5m_3um_k141_350484 | 1 | 360 | - | hypothetical protein | hypothetical protein |
| LC_0.5m_3um_k141_350484_2 | LC_0.5m_3um_k141_350484 | 442 | 2430 | - | RDRP | RDRP |
| LC_0.5m_3um_k141_350484_3 | LC_0.5m_3um_k141_350484 | 2461 | 2853 | - | hypothetical protein | hypothetical protein |
| LC_0.5m_3um_k141_350484_4 | LC_0.5m_3um_k141_350484 | 3139 | 3720 | - | hypothetical protein | hypothetical protein |
| LC_0.5m_3um_k141_350484_5 | LC_0.5m_3um_k141_350484 | 3841 | 6558 | - | hypothetical protein | hypothetical protein |
| LC_0.5m_3um_k141_350484_6 | LC_0.5m_3um_k141_350484 | 6616 | 7371 | - | hypothetical protein | hypothetical protein |
| LC_0.5m_3um_k141_258279_1 | LC_0.5m_3um_k141_258279 | 2 | 760 | + | hypothetical protein | hypothetical protein |
| LC_0.5m_3um_k141_258279_2 | LC_0.5m_3um_k141_258279 | 797 | 2038 | + | hypothetical protein | hypothetical protein |
| LC_0.5m_3um_k141_258279_3 | LC_0.5m_3um_k141_258279 | 2069 | 3535 | + | hypothetical protein | hypothetical protein |
| LC_0.5m_3um_k141_258279_4 | LC_0.5m_3um_k141_258279 | 3740 | 4102 | + | hypothetical protein | hypothetical protein |
| LC_0.5m_3um_k141_258279_5 | LC_0.5m_3um_k141_258279 | 4622 | 5803 | + | RDRP | RDRP |
| LC_0.5m_3um_k141_258279_6 | LC_0.5m_3um_k141_258279 | 5942 | 6934 | + | hypothetical protein | hypothetical protein |
| LC_0.5m_3um_k141_258279_7 | LC_0.5m_3um_k141_258279 | 7016 | 8026 | + | hypothetical protein | hypothetical protein |
| LC_coassembly_k141_1747163_1 | LC_coassembly_k141_1747163 | 1 | 1596 | - | RDRP | RDRP |
| LC_coassembly_k141_1747163_2 | LC_coassembly_k141_1747163 | 1660 | 2286 | - | hypothetical protein | hypothetical protein |
| LC_coassembly_k141_1747163_3 | LC_coassembly_k141_1747163 | 2428 | 4284 | - | hypothetical protein | hypothetical protein |
| LC_coassembly_k141_1747163_4 | LC_coassembly_k141_1747163 | 4405 | 6675 | - | hypothetical protein | hypothetical protein |
| LC_0.5m_3um_k141_36449_1 | LC_0.5m_3um_k141_36449 | 344 | 1492 | - | hypothetical protein | hypothetical protein |
| LC_0.5m_3um_k141_36449_2 | LC_0.5m_3um_k141_36449 | 1499 | 2569 | - | hypothetical protein | hypothetical protein |
| LC_0.5m_3um_k141_36449_3 | LC_0.5m_3um_k141_36449 | 2570 | 3856 | - | RDRP | RDRP |
| LC_0.5m_3um_k141_36449_4 | LC_0.5m_3um_k141_36449 | 3983 | 4606 | - | hypothetical protein | hypothetical protein |
| LC_0.5m_3um_k141_36449_5 | LC_0.5m_3um_k141_36449 | 4841 | 6307 | - | hypothetical protein | hypothetical protein |
| LC_0.5m_3um_k141_36449_6 | LC_0.5m_3um_k141_36449 | 6533 | 8374 | - | hypothetical protein | hypothetical protein |
| LC_20m_0.2um_k141_39978_1 | LC_20m_0.2um_k141_39978 | 3 | 512 | + | hypothetical protein | hypothetical protein |
| LC_20m_0.2um_k141_39978_2 | LC_20m_0.2um_k141_39978 | 531 | 2030 | + | RDRP | RDRP |
| LC_20m_0.2um_k141_39978_3 | LC_20m_0.2um_k141_39978 | 2046 | 2540 | + | capsid/coat protein | capsid/coat protein |
| LC_20m_0.2um_k141_301076_1 | LC_20m_0.2um_k141_301076 | 285 | 677 | + | hypothetical protein | hypothetical protein |
| LC_20m_0.2um_k141_301076_2 | LC_20m_0.2um_k141_301076 | 708 | 2696 | + | RDRP | RDRP |
| LC_20m_0.2um_k141_301076_3 | LC_20m_0.2um_k141_301076 | 2778 | 3788 | + | hypothetical protein | hypothetical protein |
| LC_20m_0.2um_k141_76919_1 | LC_20m_0.2um_k141_76919 | 1 | 339 | + | hypothetical protein | hypothetical protein |
| LC_20m_0.2um_k141_76919_2 | LC_20m_0.2um_k141_76919 | 358 | 729 | + | hypothetical protein | hypothetical protein |
| LC_20m_0.2um_k141_76919_3 | LC_20m_0.2um_k141_76919 | 751 | 1227 | + | hypothetical protein | hypothetical protein |
| LC_20m_0.2um_k141_76919_4 | LC_20m_0.2um_k141_76919 | 1234 | 3204 | + | hypothetical protein | hypothetical protein |
| LC_20m_0.2um_k141_76919_5 | LC_20m_0.2um_k141_76919 | 3289 | 3564 | - | hypothetical protein | hypothetical protein |
| LC_20m_0.2um_k141_76919_6 | LC_20m_0.2um_k141_76919 | 3580 | 3792 | + | hypothetical protein | hypothetical protein |
| LC_20m_0.2um_k141_76919_7 | LC_20m_0.2um_k141_76919 | 3814 | 5406 | + | RDRP | RDRP |
| LC_20m_0.2um_k141_334030_1 | LC_20m_0.2um_k141_334030 | 55 | 594 | + | hypothetical protein | hypothetical protein |
| LC_20m_0.2um_k141_334030_2 | LC_20m_0.2um_k141_334030 | 688 | 2112 | + | RDRP | RDRP |
| LC_20m_0.2um_k141_334030_3 | LC_20m_0.2um_k141_334030 | 2135 | 2800 | + | capsid/coat protein | capsid/coat protein |
| LC_20m_0.2um_k141_60959_1 | LC_20m_0.2um_k141_60959 | 1 | 558 | + | hypothetical protein | hypothetical protein |
| LC_20m_0.2um_k141_60959_2 | LC_20m_0.2um_k141_60959 | 940 | 1596 | + | hypothetical protein | hypothetical protein |
| LC_20m_0.2um_k141_60959_3 | LC_20m_0.2um_k141_60959 | 1723 | 2559 | + | RDRP | RDRP |
| LC_20m_0.2um_k141_268307_1 | LC_20m_0.2um_k141_268307 | 126 | 1697 | + | hypothetical protein | hypothetical protein |
| LC_20m_0.2um_k141_268307_2 | LC_20m_0.2um_k141_268307 | 1815 | 4670 | + | hypothetical protein | hypothetical protein |
| LC_20m_0.2um_k141_268307_3 | LC_20m_0.2um_k141_268307 | 4803 | 6224 | + | hypothetical protein | hypothetical protein |
| LC_20m_0.2um_k141_268307_4 | LC_20m_0.2um_k141_268307 | 6246 | 6470 | + | hypothetical protein | hypothetical protein |
| LC_20m_0.2um_k141_268307_5 | LC_20m_0.2um_k141_268307 | 6471 | 9251 | + | RDRP | RDRP |
| LC_20m_0.2um_k141_316934_1 | LC_20m_0.2um_k141_316934 | 3 | 665 | - | hypothetical protein | hypothetical protein |
| LC_20m_0.2um_k141_316934_2 | LC_20m_0.2um_k141_316934 | 771 | 1772 | - | hypothetical protein | hypothetical protein |
| LC_20m_0.2um_k141_316934_3 | LC_20m_0.2um_k141_316934 | 1839 | 3137 | - | RDRP | RDRP |
| LC_20m_0.2um_k141_316934_4 | LC_20m_0.2um_k141_316934 | 3601 | 3759 | + | hypothetical protein | hypothetical protein |
| LC_20m_0.2um_k141_316934_5 | LC_20m_0.2um_k141_316934 | 4107 | 4628 | - | hypothetical protein | hypothetical protein |
| LC_coassembly_k141_1116952_1 | LC_coassembly_k141_1116952 | 2 | 868 | + | hypothetical protein | hypothetical protein |
| LC_coassembly_k141_1116952_2 | LC_coassembly_k141_1116952 | 1037 | 5098 | + | hypothetical protein | hypothetical protein |
| LC_coassembly_k141_1116952_3 | LC_coassembly_k141_1116952 | 5120 | 5860 | + | RDRP | RDRP |
| LC_20m_1.2um_k141_408408_1 | LC_20m_1.2um_k141_408408 | 3 | 1256 | - | RDRP | RDRP |
| LC_20m_1.2um_k141_408408_2 | LC_20m_1.2um_k141_408408 | 1287 | 1679 | - | hypothetical protein | hypothetical protein |
| LC_20m_1.2um_k141_408408_3 | LC_20m_1.2um_k141_408408 | 2100 | 2369 | - | hypothetical protein | hypothetical protein |
| LC_20m_1.2um_k141_408408_4 | LC_20m_1.2um_k141_408408 | 2667 | 5384 | - | hypothetical protein | hypothetical protein |
| LC_20m_1.2um_k141_408408_5 | LC_20m_1.2um_k141_408408 | 5562 | 5828 | - | hypothetical protein | hypothetical protein |
| LC_20m_1.2um_k141_266503_1 | LC_20m_1.2um_k141_266503 | 438 | 2600 | - | hypothetical protein | hypothetical protein |
| LC_20m_1.2um_k141_266503_2 | LC_20m_1.2um_k141_266503 | 2667 | 4085 | - | RDRP | RDRP |
| LC_20m_1.2um_k141_266503_3 | LC_20m_1.2um_k141_266503 | 4104 | 6404 | - | hypothetical protein | hypothetical protein |
| LC_20m_1.2um_k141_266503_4 | LC_20m_1.2um_k141_266503 | 6423 | 7796 | - | hypothetical protein | hypothetical protein |
| LC_20m_1.2um_k141_266503_5 | LC_20m_1.2um_k141_266503 | 7830 | 8456 | - | hypothetical protein | hypothetical protein |
| LC_20m_1.2um_k141_147958_1 | LC_20m_1.2um_k141_147958 | 3 | 1562 | - | hypothetical protein | hypothetical protein |
| LC_20m_1.2um_k141_147958_2 | LC_20m_1.2um_k141_147958 | 1629 | 2915 | - | RDRP | RDRP |
| LC_20m_1.2um_k141_147958_3 | LC_20m_1.2um_k141_147958 | 3042 | 3677 | - | hypothetical protein | hypothetical protein |
| LC_20m_1.2um_k141_257976_1 | LC_20m_1.2um_k141_257976 | 3 | 209 | - | hypothetical protein | hypothetical protein |
| LC_20m_1.2um_k141_257976_2 | LC_20m_1.2um_k141_257976 | 291 | 2279 | - | RDRP | RDRP |
| LC_20m_1.2um_k141_257976_3 | LC_20m_1.2um_k141_257976 | 2310 | 2702 | - | hypothetical protein | hypothetical protein |
| LC_20m_1.2um_k141_257976_4 | LC_20m_1.2um_k141_257976 | 2988 | 3569 | - | hypothetical protein | hypothetical protein |
| LC_20m_1.2um_k141_257976_5 | LC_20m_1.2um_k141_257976 | 3690 | 6431 | - | hypothetical protein | hypothetical protein |
| LC_20m_1.2um_k141_257976_6 | LC_20m_1.2um_k141_257976 | 6465 | 7217 | - | hypothetical protein | hypothetical protein |
| LC_20m_1.2um_k141_5914_1 | LC_20m_1.2um_k141_5914 | 1 | 282 | - | hypothetical protein | hypothetical protein |
| LC_20m_1.2um_k141_5914_2 | LC_20m_1.2um_k141_5914 | 388 | 1389 | - | hypothetical protein | hypothetical protein |
| LC_20m_1.2um_k141_5914_3 | LC_20m_1.2um_k141_5914 | 1456 | 2781 | - | RDRP | RDRP |
| LC_20m_1.2um_k141_5914_4 | LC_20m_1.2um_k141_5914 | 3022 | 3519 | - | hypothetical protein | hypothetical protein |
| LC_20m_1.2um_k141_397998_1 | LC_20m_1.2um_k141_397998 | 1 | 645 | + | hypothetical protein | hypothetical protein |
| LC_20m_1.2um_k141_397998_2 | LC_20m_1.2um_k141_397998 | 679 | 1323 | + | hypothetical protein | hypothetical protein |
| LC_20m_1.2um_k141_397998_3 | LC_20m_1.2um_k141_397998 | 1324 | 3543 | + | hypothetical protein | hypothetical protein |
| LC_20m_1.2um_k141_397998_4 | LC_20m_1.2um_k141_397998 | 3712 | 4110 | + | hypothetical protein | hypothetical protein |
| LC_20m_1.2um_k141_397998_5 | LC_20m_1.2um_k141_397998 | 4513 | 5811 | + | RDRP | RDRP |
| LC_20m_1.2um_k141_397998_6 | LC_20m_1.2um_k141_397998 | 5950 | 7188 | + | hypothetical protein | hypothetical protein |
| LC_20m_1.2um_k141_302326_1 | LC_20m_1.2um_k141_302326 | 3 | 641 | + | hypothetical protein | hypothetical protein |
| LC_20m_1.2um_k141_302326_2 | LC_20m_1.2um_k141_302326 | 675 | 1319 | + | hypothetical protein | hypothetical protein |
| LC_20m_1.2um_k141_302326_3 | LC_20m_1.2um_k141_302326 | 1320 | 4106 | + | hypothetical protein | hypothetical protein |
| LC_20m_1.2um_k141_302326_4 | LC_20m_1.2um_k141_302326 | 4509 | 5288 | + | hypothetical protein | hypothetical protein |
| LC_20m_1.2um_k141_302326_5 | LC_20m_1.2um_k141_302326 | 5292 | 6884 | + | RDRP | RDRP |
| LC_20m_1.2um_k141_302326_6 | LC_20m_1.2um_k141_302326 | 6990 | 8039 | + | hypothetical protein | hypothetical protein |
| LC_20m_3um_k141_163697_1 | LC_20m_3um_k141_163697 | 1 | 108 | + | hypothetical protein | hypothetical protein |
| LC_20m_3um_k141_163697_2 | LC_20m_3um_k141_163697 | 158 | 964 | + | hypothetical protein | hypothetical protein |
| LC_20m_3um_k141_163697_3 | LC_20m_3um_k141_163697 | 990 | 2498 | + | RDRP | RDRP |
| LC_20m_3um_k141_522525_1 | LC_20m_3um_k141_522525 | 3 | 689 | - | hypothetical protein | hypothetical protein |
| LC_20m_3um_k141_522525_2 | LC_20m_3um_k141_522525 | 690 | 1745 | - | RDRP | RDRP |
| LC_20m_3um_k141_522525_3 | LC_20m_3um_k141_522525 | 1791 | 1976 | - | hypothetical protein | hypothetical protein |
| LC_20m_3um_k141_522525_4 | LC_20m_3um_k141_522525 | 2103 | 2954 | - | hypothetical protein | hypothetical protein |
| LC_20m_3um_k141_522525_5 | LC_20m_3um_k141_522525 | 2955 | 6509 | - | hypothetical protein | hypothetical protein |
| LC_20m_3um_k141_460466_1 | LC_20m_3um_k141_460466 | 2 | 604 | + | RDRP | RDRP |
| LC_20m_3um_k141_460466_2 | LC_20m_3um_k141_460466 | 743 | 1678 | + | hypothetical protein | hypothetical protein |
| LC_20m_3um_k141_460466_3 | LC_20m_3um_k141_460466 | 1733 | 2002 | + | hypothetical protein | hypothetical protein |
| LC_20m_3um_k141_227703_1 | LC_20m_3um_k141_227703 | 68 | 889 | - | hypothetical protein | hypothetical protein |
| LC_20m_3um_k141_227703_2 | LC_20m_3um_k141_227703 | 1196 | 3121 | - | RDRP | RDRP |
| LC_20m_3um_k141_227703_3 | LC_20m_3um_k141_227703 | 3202 | 4023 | - | hypothetical protein | hypothetical protein |
| LC_20m_3um_k141_587133_1 | LC_20m_3um_k141_587133 | 3 | 1793 | + | hypothetical protein | hypothetical protein |
| LC_20m_3um_k141_587133_2 | LC_20m_3um_k141_587133 | 2019 | 3485 | + | hypothetical protein | hypothetical protein |
| LC_20m_3um_k141_587133_3 | LC_20m_3um_k141_587133 | 3720 | 4343 | + | hypothetical protein | hypothetical protein |
| LC_20m_3um_k141_587133_4 | LC_20m_3um_k141_587133 | 4470 | 5756 | + | RDRP | RDRP |
| LC_20m_3um_k141_587133_5 | LC_20m_3um_k141_587133 | 5757 | 6827 | + | hypothetical protein | hypothetical protein |
| LC_20m_3um_k141_587133_6 | LC_20m_3um_k141_587133 | 6834 | 7982 | + | hypothetical protein | hypothetical protein |
| LC_20m_3um_k141_14922_1 | LC_20m_3um_k141_14922 | 2 | 3532 | + | hypothetical protein | hypothetical protein |
| LC_20m_3um_k141_14922_2 | LC_20m_3um_k141_14922 | 3533 | 4384 | + | hypothetical protein | hypothetical protein |
| LC_20m_3um_k141_14922_3 | LC_20m_3um_k141_14922 | 4511 | 5797 | + | RDRP | RDRP |
| LC_20m_3um_k141_14922_4 | LC_20m_3um_k141_14922 | 5798 | 7063 | + | hypothetical protein | hypothetical protein |
| LC_20m_3um_k141_115819_1 | LC_20m_3um_k141_115819 | 2 | 436 | + | hypothetical protein | hypothetical protein |
| LC_20m_3um_k141_115819_2 | LC_20m_3um_k141_115819 | 458 | 1435 | + | hypothetical protein | hypothetical protein |
| LC_20m_3um_k141_115819_3 | LC_20m_3um_k141_115819 | 1447 | 1890 | - | hypothetical protein | hypothetical protein |
| LC_20m_3um_k141_115819_4 | LC_20m_3um_k141_115819 | 2019 | 2243 | - | hypothetical protein | hypothetical protein |
| LC_20m_3um_k141_115819_5 | LC_20m_3um_k141_115819 | 2938 | 5094 | + | RDRP | RDRP |
| LC_20m_3um_k141_298856_1 | LC_20m_3um_k141_298856 | 3 | 536 | + | hypothetical protein | hypothetical protein |
| LC_20m_3um_k141_298856_2 | LC_20m_3um_k141_298856 | 675 | 3539 | + | hypothetical protein | hypothetical protein |
| LC_20m_3um_k141_298856_3 | LC_20m_3um_k141_298856 | 3744 | 4241 | + | hypothetical protein | hypothetical protein |
| LC_20m_3um_k141_298856_4 | LC_20m_3um_k141_298856 | 4509 | 5807 | + | RDRP | RDRP |
| LC_20m_3um_k141_298856_5 | LC_20m_3um_k141_298856 | 5874 | 6686 | + | hypothetical protein | hypothetical protein |
| LC_20m_3um_k141_538898_1 | LC_20m_3um_k141_538898 | 333 | 497 | - | hypothetical protein | hypothetical protein |
| LC_20m_3um_k141_538898_2 | LC_20m_3um_k141_538898 | 507 | 1577 | - | RDRP | RDRP |
| LC_20m_3um_k141_538898_3 | LC_20m_3um_k141_538898 | 1605 | 1700 | - | hypothetical protein | hypothetical protein |
| LC_20m_3um_k141_299800_1 | LC_20m_3um_k141_299800 | 3 | 536 | + | hypothetical protein | hypothetical protein |
| LC_20m_3um_k141_299800_2 | LC_20m_3um_k141_299800 | 675 | 1673 | + | hypothetical protein | hypothetical protein |
| LC_20m_3um_k141_299800_3 | LC_20m_3um_k141_299800 | 2006 | 3538 | + | hypothetical protein | hypothetical protein |
| LC_20m_3um_k141_299800_4 | LC_20m_3um_k141_299800 | 3743 | 4240 | + | hypothetical protein | hypothetical protein |
| LC_20m_3um_k141_299800_5 | LC_20m_3um_k141_299800 | 4481 | 5806 | + | RDRP | RDRP |
| LC_20m_3um_k141_299800_6 | LC_20m_3um_k141_299800 | 5873 | 6496 | + | hypothetical protein | hypothetical protein |
| LC_coassembly_k141_367052_1 | LC_coassembly_k141_367052 | 2 | 181 | + | hypothetical protein | hypothetical protein |
| LC_coassembly_k141_367052_2 | LC_coassembly_k141_367052 | 715 | 2193 | - | RDRP | RDRP |
| LC_coassembly_k141_367052_3 | LC_coassembly_k141_367052 | 2655 | 3224 | - | hypothetical protein | hypothetical protein |
| LC_20m_3um_k141_445022_1 | LC_20m_3um_k141_445022 | 3 | 674 | + | hypothetical protein | hypothetical protein |
| LC_20m_3um_k141_445022_2 | LC_20m_3um_k141_445022 | 622 | 2286 | + | hypothetical protein | hypothetical protein |
| LC_20m_3um_k141_445022_3 | LC_20m_3um_k141_445022 | 2308 | 2580 | + | hypothetical protein | hypothetical protein |
| LC_20m_3um_k141_445022_4 | LC_20m_3um_k141_445022 | 2618 | 4192 | + | RDRP | RDRP |
| LC_20m_3um_k141_300840_1 | LC_20m_3um_k141_300840 | 1 | 1158 | - | RDRP | RDRP |
| LC_20m_3um_k141_300840_2 | LC_20m_3um_k141_300840 | 1177 | 2196 | - | hypothetical protein | hypothetical protein |
| LC_20m_3um_k141_300840_3 | LC_20m_3um_k141_300840 | 2211 | 3476 | - | hypothetical protein | hypothetical protein |
| LC_20m_3um_k141_300840_4 | LC_20m_3um_k141_300840 | 3495 | 3614 | - | hypothetical protein | hypothetical protein |
| LC_20m_3um_k141_93254_1 | LC_20m_3um_k141_93254 | 2 | 532 | + | hypothetical protein | hypothetical protein |
| LC_20m_3um_k141_93254_2 | LC_20m_3um_k141_93254 | 671 | 3325 | + | hypothetical protein | hypothetical protein |
| LC_20m_3um_k141_93254_3 | LC_20m_3um_k141_93254 | 3362 | 3721 | + | hypothetical protein | hypothetical protein |
| LC_20m_3um_k141_93254_4 | LC_20m_3um_k141_93254 | 3883 | 4071 | - | hypothetical protein | hypothetical protein |
| LC_20m_3um_k141_93254_5 | LC_20m_3um_k141_93254 | 4505 | 5803 | + | RDRP | RDRP |
| LC_20m_3um_k141_93254_6 | LC_20m_3um_k141_93254 | 5870 | 8026 | + | hypothetical protein | hypothetical protein |
| LC_20m_3um_k141_389751_1 | LC_20m_3um_k141_389751 | 134 | 3148 | + | hypothetical protein | hypothetical protein |
| LC_20m_3um_k141_389751_2 | LC_20m_3um_k141_389751 | 3196 | 4536 | + | hypothetical protein | hypothetical protein |
| LC_20m_3um_k141_389751_3 | LC_20m_3um_k141_389751 | 4546 | 7005 | + | RDRP | RDRP |
| LC_20m_3um_k141_389751_4 | LC_20m_3um_k141_389751 | 7117 | 8019 | + | hypothetical protein | hypothetical protein |
| LC_20m_3um_k141_585136_1 | LC_20m_3um_k141_585136 | 2 | 937 | - | hypothetical protein | hypothetical protein |
| LC_20m_3um_k141_585136_2 | LC_20m_3um_k141_585136 | 1004 | 2644 | - | RDRP | RDRP |
| LC_20m_3um_k141_585136_3 | LC_20m_3um_k141_585136 | 2645 | 3394 | - | hypothetical protein | hypothetical protein |
| LC_20m_3um_k141_585136_4 | LC_20m_3um_k141_585136 | 3476 | 4375 | - | hypothetical protein | hypothetical protein |
| LC_50m_0.2um_k141_182361_1 | LC_50m_0.2um_k141_182361 | 1 | 1308 | + | hypothetical protein | hypothetical protein |
| LC_50m_0.2um_k141_182361_2 | LC_50m_0.2um_k141_182361 | 1309 | 3534 | + | hypothetical protein | hypothetical protein |
| LC_50m_0.2um_k141_182361_3 | LC_50m_0.2um_k141_182361 | 3757 | 4392 | + | hypothetical protein | hypothetical protein |
| LC_50m_0.2um_k141_182361_4 | LC_50m_0.2um_k141_182361 | 4519 | 5805 | + | RDRP | RDRP |
| LC_50m_0.2um_k141_182361_5 | LC_50m_0.2um_k141_182361 | 5872 | 8031 | + | hypothetical protein | hypothetical protein |
| LC_50m_0.2um_k141_187776_1 | LC_50m_0.2um_k141_187776 | 1 | 1008 | - | hypothetical protein | hypothetical protein |
| LC_50m_0.2um_k141_187776_2 | LC_50m_0.2um_k141_187776 | 1147 | 2445 | - | RDRP | RDRP |
| LC_50m_0.2um_k141_187776_3 | LC_50m_0.2um_k141_187776 | 2848 | 3246 | - | hypothetical protein | hypothetical protein |
| LC_50m_0.2um_k141_187776_4 | LC_50m_0.2um_k141_187776 | 3415 | 5634 | - | hypothetical protein | hypothetical protein |
| LC_50m_0.2um_k141_187776_5 | LC_50m_0.2um_k141_187776 | 5635 | 6279 | - | hypothetical protein | hypothetical protein |
| LC_50m_0.2um_k141_187776_6 | LC_50m_0.2um_k141_187776 | 6313 | 6957 | - | hypothetical protein | hypothetical protein |
| LC_coassembly_k141_1177772_1 | LC_coassembly_k141_1177772 | 32 | 1843 | - | RDRP | RDRP |
| LC_coassembly_k141_1177772_2 | LC_coassembly_k141_1177772 | 1901 | 2032 | - | hypothetical protein | hypothetical protein |
| LC_coassembly_k141_1177772_3 | LC_coassembly_k141_1177772 | 2046 | 2510 | - | hypothetical protein | hypothetical protein |
| LC_coassembly_k141_1177772_4 | LC_coassembly_k141_1177772 | 2533 | 3873 | - | capsid/coat protein | capsid/coat protein |
| LC_coassembly_k141_1177772_5 | LC_coassembly_k141_1177772 | 4086 | 4205 | - | hypothetical protein | hypothetical protein |
| LC_50m_0.2um_k141_442366_1 | LC_50m_0.2um_k141_442366 | 2 | 2974 | + | hypothetical protein | hypothetical protein |
| LC_50m_0.2um_k141_442366_2 | LC_50m_0.2um_k141_442366 | 3377 | 3478 | - | hypothetical protein | hypothetical protein |
| LC_50m_0.2um_k141_442366_3 | LC_50m_0.2um_k141_442366 | 3942 | 5240 | + | RDRP | RDRP |
| LC_50m_0.2um_k141_442366_4 | LC_50m_0.2um_k141_442366 | 5307 | 6308 | + | hypothetical protein | hypothetical protein |
| LC_50m_0.2um_k141_442366_5 | LC_50m_0.2um_k141_442366 | 6414 | 7046 | + | hypothetical protein | hypothetical protein |
| LC_50m_0.2um_k141_68686_1 | LC_50m_0.2um_k141_68686 | 2 | 1417 | + | RDRP | RDRP |
| LC_50m_0.2um_k141_68686_2 | LC_50m_0.2um_k141_68686 | 1419 | 1655 | + | hypothetical protein | hypothetical protein |
| LC_50m_0.2um_k141_68686_3 | LC_50m_0.2um_k141_68686 | 1693 | 2259 | + | hypothetical protein | hypothetical protein |
| LC_50m_0.2um_k141_84230_1 | LC_50m_0.2um_k141_84230 | 3 | 2366 | - | RDRP | RDRP |
| LC_50m_0.2um_k141_84230_2 | LC_50m_0.2um_k141_84230 | 2394 | 4847 | - | hypothetical protein | hypothetical protein |
| LC_50m_0.2um_k141_84230_3 | LC_50m_0.2um_k141_84230 | 4869 | 6632 | - | hypothetical protein | hypothetical protein |
| LC_50m_0.2um_k141_58378_1 | LC_50m_0.2um_k141_58378 | 3 | 494 | + | hypothetical protein | hypothetical protein |
| LC_50m_0.2um_k141_58378_2 | LC_50m_0.2um_k141_58378 | 633 | 3683 | + | hypothetical protein | hypothetical protein |
| LC_50m_0.2um_k141_58378_3 | LC_50m_0.2um_k141_58378 | 3702 | 4166 | + | hypothetical protein | hypothetical protein |
| LC_50m_0.2um_k141_58378_4 | LC_50m_0.2um_k141_58378 | 4416 | 7988 | + | RDRP | RDRP |
| LC_50m_0.2um_k141_318047_1 | LC_50m_0.2um_k141_318047 | 2 | 1381 | + | hypothetical protein | hypothetical protein |
| LC_50m_0.2um_k141_318047_2 | LC_50m_0.2um_k141_318047 | 1406 | 1777 | + | hypothetical protein | hypothetical protein |
| LC_50m_0.2um_k141_318047_3 | LC_50m_0.2um_k141_318047 | 1799 | 2275 | + | hypothetical protein | hypothetical protein |
| LC_50m_0.2um_k141_318047_4 | LC_50m_0.2um_k141_318047 | 2282 | 4252 | + | hypothetical protein | hypothetical protein |
| LC_50m_0.2um_k141_318047_5 | LC_50m_0.2um_k141_318047 | 4394 | 6724 | + | RDRP | RDRP |
| LC_50m_0.2um_k141_318047_6 | LC_50m_0.2um_k141_318047 | 6791 | 7813 | + | hypothetical protein | hypothetical protein |
| LC_50m_0.2um_k141_292980_1 | LC_50m_0.2um_k141_292980 | 2 | 535 | + | hypothetical protein | hypothetical protein |
| LC_50m_0.2um_k141_292980_2 | LC_50m_0.2um_k141_292980 | 659 | 1594 | + | hypothetical protein | hypothetical protein |
| LC_50m_0.2um_k141_292980_3 | LC_50m_0.2um_k141_292980 | 1895 | 3517 | + | hypothetical protein | hypothetical protein |
| LC_50m_0.2um_k141_292980_4 | LC_50m_0.2um_k141_292980 | 3524 | 3859 | + | hypothetical protein | hypothetical protein |
| LC_50m_0.2um_k141_292980_5 | LC_50m_0.2um_k141_292980 | 3920 | 4330 | + | hypothetical protein | hypothetical protein |
| LC_50m_0.2um_k141_292980_6 | LC_50m_0.2um_k141_292980 | 4487 | 5788 | + | RDRP | RDRP |
| LC_50m_0.2um_k141_292980_7 | LC_50m_0.2um_k141_292980 | 5855 | 8017 | + | hypothetical protein | hypothetical protein |
| LC_50m_0.2um_k141_80321_1 | LC_50m_0.2um_k141_80321 | 1 | 1086 | - | RDRP | RDRP |
| LC_50m_0.2um_k141_80321_2 | LC_50m_0.2um_k141_80321 | 1213 | 1848 | - | hypothetical protein | hypothetical protein |
| LC_50m_0.2um_k141_80321_3 | LC_50m_0.2um_k141_80321 | 2071 | 4296 | - | hypothetical protein | hypothetical protein |
| LC_50m_0.2um_k141_80321_4 | LC_50m_0.2um_k141_80321 | 4297 | 4938 | - | hypothetical protein | hypothetical protein |
| LC_50m_0.2um_k141_373844_1 | LC_50m_0.2um_k141_373844 | 442 | 894 | - | hypothetical protein | hypothetical protein |
| LC_50m_0.2um_k141_373844_2 | LC_50m_0.2um_k141_373844 | 1349 | 1801 | - | RDRP | RDRP |
| LC_50m_0.2um_k141_373844_3 | LC_50m_0.2um_k141_373844 | 2126 | 2233 | - | hypothetical protein | hypothetical protein |
| LC_50m_10um_k141_22825_1 | LC_50m_10um_k141_22825 | 1 | 387 | + | hypothetical protein | hypothetical protein |
| LC_50m_10um_k141_22825_2 | LC_50m_10um_k141_22825 | 506 | 1606 | - | hypothetical protein | hypothetical protein |
| LC_50m_10um_k141_22825_3 | LC_50m_10um_k141_22825 | 1661 | 2734 | - | hypothetical protein | hypothetical protein |
| LC_50m_10um_k141_22825_4 | LC_50m_10um_k141_22825 | 2735 | 3466 | - | RDRP | RDRP |
| LC_50m_10um_k141_114996_1 | LC_50m_10um_k141_114996 | 206 | 559 | + | hypothetical protein | hypothetical protein |
| LC_50m_10um_k141_114996_2 | LC_50m_10um_k141_114996 | 767 | 2389 | + | hypothetical protein | hypothetical protein |
| LC_50m_10um_k141_114996_3 | LC_50m_10um_k141_114996 | 2474 | 3334 | + | hypothetical protein | hypothetical protein |
| LC_50m_10um_k141_114996_4 | LC_50m_10um_k141_114996 | 3632 | 3901 | + | hypothetical protein | hypothetical protein |
| LC_50m_10um_k141_114996_5 | LC_50m_10um_k141_114996 | 4322 | 4714 | + | hypothetical protein | hypothetical protein |
| LC_50m_10um_k141_114996_6 | LC_50m_10um_k141_114996 | 4745 | 6304 | + | RDRP | RDRP |
| LC_50m_1.2um_k141_22302_1 | LC_50m_1.2um_k141_22302 | 1 | 804 | - | hypothetical protein | hypothetical protein |
| LC_50m_1.2um_k141_22302_2 | LC_50m_1.2um_k141_22302 | 871 | 2967 | - | RDRP | RDRP |
| LC_50m_1.2um_k141_22302_3 | LC_50m_1.2um_k141_22302 | 2983 | 3258 | + | hypothetical protein | hypothetical protein |
| LC_50m_1.2um_k141_22302_4 | LC_50m_1.2um_k141_22302 | 3343 | 4860 | - | hypothetical protein | hypothetical protein |
| LC_50m_1.2um_k141_213095_1 | LC_50m_1.2um_k141_213095 | 439 | 1449 | - | hypothetical protein | hypothetical protein |
| LC_50m_1.2um_k141_213095_2 | LC_50m_1.2um_k141_213095 | 1531 | 3519 | - | RDRP | RDRP |
| LC_50m_1.2um_k141_213095_3 | LC_50m_1.2um_k141_213095 | 4363 | 4632 | - | hypothetical protein | hypothetical protein |
| LC_50m_1.2um_k141_213095_4 | LC_50m_1.2um_k141_213095 | 4930 | 7647 | - | hypothetical protein | hypothetical protein |
| LC_50m_1.2um_k141_213095_5 | LC_50m_1.2um_k141_213095 | 7825 | 7983 | - | hypothetical protein | hypothetical protein |
| LC_50m_1.2um_k141_297650_1 | LC_50m_1.2um_k141_297650 | 2 | 208 | - | hypothetical protein | hypothetical protein |
| LC_50m_1.2um_k141_297650_2 | LC_50m_1.2um_k141_297650 | 488 | 6928 | + | capsid/coat protein | capsid/coat protein |
| LC_50m_1.2um_k141_297650_3 | LC_50m_1.2um_k141_297650 | 6994 | 9639 | + | RDRP | RDRP |
| LC_50m_1.2um_k141_298547_1 | LC_50m_1.2um_k141_298547 | 2 | 4252 | + | hypothetical protein | hypothetical protein |
| LC_50m_1.2um_k141_298547_2 | LC_50m_1.2um_k141_298547 | 4334 | 5083 | + | RDRP | RDRP |
| LC_50m_1.2um_k141_298547_3 | LC_50m_1.2um_k141_298547 | 5084 | 6724 | + | RDRP | RDRP |
| LC_50m_1.2um_k141_298547_4 | LC_50m_1.2um_k141_298547 | 6791 | 7297 | + | hypothetical protein | hypothetical protein |
| LC_50m_1.2um_k141_307887_1 | LC_50m_1.2um_k141_307887 | 2 | 73 | - | hypothetical protein | hypothetical protein |
| LC_50m_1.2um_k141_307887_2 | LC_50m_1.2um_k141_307887 | 948 | 1256 | - | RDRP | RDRP |
| LC_50m_1.2um_k141_307887_3 | LC_50m_1.2um_k141_307887 | 1588 | 2166 | + | hypothetical protein | hypothetical protein |
| LC_50m_1.2um_k141_78854_1 | LC_50m_1.2um_k141_78854 | 435 | 2591 | - | hypothetical protein | hypothetical protein |
| LC_50m_1.2um_k141_78854_2 | LC_50m_1.2um_k141_78854 | 2658 | 3956 | - | RDRP | RDRP |
| LC_50m_1.2um_k141_78854_3 | LC_50m_1.2um_k141_78854 | 4390 | 4578 | + | hypothetical protein | hypothetical protein |
| LC_50m_1.2um_k141_78854_4 | LC_50m_1.2um_k141_78854 | 4740 | 5099 | - | hypothetical protein | hypothetical protein |
| LC_50m_1.2um_k141_78854_5 | LC_50m_1.2um_k141_78854 | 5136 | 7790 | - | hypothetical protein | hypothetical protein |
| LC_50m_1.2um_k141_78854_6 | LC_50m_1.2um_k141_78854 | 7929 | 8459 | - | hypothetical protein | hypothetical protein |
| LC_50m_1.2um_k141_21225_1 | LC_50m_1.2um_k141_21225 | 1 | 654 | - | hypothetical protein | hypothetical protein |
| LC_50m_1.2um_k141_21225_2 | LC_50m_1.2um_k141_21225 | 760 | 2352 | - | RDRP | RDRP |
| LC_50m_1.2um_k141_21225_3 | LC_50m_1.2um_k141_21225 | 2356 | 3135 | - | hypothetical protein | hypothetical protein |
| LC_50m_1.2um_k141_21225_4 | LC_50m_1.2um_k141_21225 | 3538 | 6324 | - | hypothetical protein | hypothetical protein |
| LC_50m_1.2um_k141_21225_5 | LC_50m_1.2um_k141_21225 | 6325 | 6969 | - | hypothetical protein | hypothetical protein |
| LC_50m_1.2um_k141_21225_6 | LC_50m_1.2um_k141_21225 | 7003 | 7635 | - | hypothetical protein | hypothetical protein |
| LC_50m_1.2um_k141_394034_1 | LC_50m_1.2um_k141_394034 | 1314 | 1421 | + | hypothetical protein | hypothetical protein |
| LC_50m_1.2um_k141_394034_2 | LC_50m_1.2um_k141_394034 | 1746 | 2198 | + | RDRP | RDRP |
| LC_50m_1.2um_k141_394034_3 | LC_50m_1.2um_k141_394034 | 2653 | 3105 | + | hypothetical protein | hypothetical protein |
| LC_50m_1.2um_k141_394034_4 | LC_50m_1.2um_k141_394034 | 3594 | 3671 | + | hypothetical protein | hypothetical protein |
| LC_coassembly_k141_2619831_1 | LC_coassembly_k141_2619831 | 344 | 925 | - | hypothetical protein | hypothetical protein |
| LC_coassembly_k141_2619831_2 | LC_coassembly_k141_2619831 | 1574 | 2077 | - | hypothetical protein | hypothetical protein |
| LC_coassembly_k141_2619831_3 | LC_coassembly_k141_2619831 | 2849 | 2968 | - | hypothetical protein | hypothetical protein |
| LC_coassembly_k141_2619831_4 | LC_coassembly_k141_2619831 | 3014 | 3877 | - | RDRP | RDRP |
| LC_coassembly_k141_2619831_5 | LC_coassembly_k141_2619831 | 4004 | 4660 | - | hypothetical protein | hypothetical protein |
| LC_coassembly_k141_2619831_6 | LC_coassembly_k141_2619831 | 5042 | 5968 | - | hypothetical protein | hypothetical protein |
| LC_coassembly_k141_2619831_7 | LC_coassembly_k141_2619831 | 5990 | 6184 | - | hypothetical protein | hypothetical protein |
| LC_coassembly_k141_2619831_8 | LC_coassembly_k141_2619831 | 6335 | 6730 | - | hypothetical protein | hypothetical protein |
| LC_coassembly_k141_2619831_9 | LC_coassembly_k141_2619831 | 7022 | 7219 | - | hypothetical protein | hypothetical protein |
| LC_coassembly_k141_2619831_10 | LC_coassembly_k141_2619831 | 7562 | 7786 | - | hypothetical protein | hypothetical protein |
| LC_coassembly_k141_2619831_11 | LC_coassembly_k141_2619831 | 8135 | 8599 | - | hypothetical protein | hypothetical protein |
| LC_coassembly_k141_2619831_12 | LC_coassembly_k141_2619831 | 8747 | 9067 | - | hypothetical protein | hypothetical protein |
| LC_coassembly_k141_2619831_13 | LC_coassembly_k141_2619831 | 9383 | 9709 | - | hypothetical protein | hypothetical protein |
| LC_coassembly_k141_2619831_14 | LC_coassembly_k141_2619831 | 10004 | 10342 | - | hypothetical protein | hypothetical protein |
| LC_coassembly_k141_2619831_15 | LC_coassembly_k141_2619831 | 10682 | 11233 | - | hypothetical protein | hypothetical protein |
| LC_50m_1.2um_k141_194815_1 | LC_50m_1.2um_k141_194815 | 2 | 184 | + | hypothetical protein | hypothetical protein |
| LC_50m_1.2um_k141_194815_2 | LC_50m_1.2um_k141_194815 | 530 | 1540 | - | hypothetical protein | hypothetical protein |
| LC_50m_1.2um_k141_194815_3 | LC_50m_1.2um_k141_194815 | 1622 | 3427 | - | RDRP | RDRP |
| LC_50m_3um_k141_469426_1 | LC_50m_3um_k141_469426 | 1 | 951 | + | hypothetical protein | hypothetical protein |
| LC_50m_3um_k141_469426_2 | LC_50m_3um_k141_469426 | 1174 | 1809 | + | hypothetical protein | hypothetical protein |
| LC_50m_3um_k141_469426_3 | LC_50m_3um_k141_469426 | 1936 | 3222 | + | RDRP | RDRP |
| LC_50m_3um_k141_189961_1 | LC_50m_3um_k141_189961 | 1 | 222 | - | hypothetical protein | hypothetical protein |
| LC_50m_3um_k141_189961_2 | LC_50m_3um_k141_189961 | 289 | 2619 | - | RDRP | RDRP |
| LC_50m_3um_k141_189961_3 | LC_50m_3um_k141_189961 | 2761 | 4731 | - | hypothetical protein | hypothetical protein |
| LC_50m_3um_k141_189961_4 | LC_50m_3um_k141_189961 | 4738 | 5214 | - | hypothetical protein | hypothetical protein |
| LC_50m_3um_k141_189961_5 | LC_50m_3um_k141_189961 | 5236 | 5607 | - | hypothetical protein | hypothetical protein |
| LC_50m_3um_k141_189961_6 | LC_50m_3um_k141_189961 | 5632 | 7050 | - | hypothetical protein | hypothetical protein |
| LC_coassembly_k141_1689211_1 | LC_coassembly_k141_1689211 | 2 | 127 | + | hypothetical protein | hypothetical protein |
| LC_coassembly_k141_1689211_2 | LC_coassembly_k141_1689211 | 261 | 1787 | + | RDRP | RDRP |
| LC_coassembly_k141_1689211_3 | LC_coassembly_k141_1689211 | 1822 | 2073 | + | hypothetical protein | hypothetical protein |
| LC_50m_3um_k141_156190_1 | LC_50m_3um_k141_156190 | 94 | 273 | + | hypothetical protein | hypothetical protein |
| LC_50m_3um_k141_156190_2 | LC_50m_3um_k141_156190 | 625 | 3345 | - | RDRP | RDRP |
| LC_50m_3um_k141_156190_3 | LC_50m_3um_k141_156190 | 3406 | 3630 | - | hypothetical protein | hypothetical protein |
| LC_50m_3um_k141_156190_4 | LC_50m_3um_k141_156190 | 3652 | 5073 | - | hypothetical protein | hypothetical protein |
| LC_50m_3um_k141_156190_5 | LC_50m_3um_k141_156190 | 5206 | 6543 | - | hypothetical protein | hypothetical protein |
| LC_50m_3um_k141_385349_1 | LC_50m_3um_k141_385349 | 172 | 993 | + | hypothetical protein | hypothetical protein |
| LC_50m_3um_k141_385349_2 | LC_50m_3um_k141_385349 | 1074 | 2999 | + | RDRP | RDRP |
| LC_50m_3um_k141_385349_3 | LC_50m_3um_k141_385349 | 3315 | 4136 | + | hypothetical protein | hypothetical protein |
| LC_50m_3um_k141_385349_4 | LC_50m_3um_k141_385349 | 4217 | 6142 | + | RDRP | RDRP |
| LC_50m_3um_k141_238018_1 | LC_50m_3um_k141_238018 | 3 | 764 | + | hypothetical protein | hypothetical protein |
| LC_50m_3um_k141_238018_2 | LC_50m_3um_k141_238018 | 798 | 3539 | + | hypothetical protein | hypothetical protein |
| LC_50m_3um_k141_238018_3 | LC_50m_3um_k141_238018 | 3660 | 4241 | + | hypothetical protein | hypothetical protein |
| LC_50m_3um_k141_238018_4 | LC_50m_3um_k141_238018 | 4527 | 4919 | + | hypothetical protein | hypothetical protein |
| LC_50m_3um_k141_238018_5 | LC_50m_3um_k141_238018 | 4950 | 6521 | + | RDRP | RDRP |
| LC_50m_3um_k141_238018_6 | LC_50m_3um_k141_239166 | 3 | 2612 | - | RDRP | RDRP |
| LC_50m_3um_k141_238018_7 | LC_50m_3um_k141_239166 | 2862 | 3326 | - | hypothetical protein | hypothetical protein |
| LC_50m_3um_k141_238018_8 | LC_50m_3um_k141_239166 | 3345 | 6395 | - | hypothetical protein | hypothetical protein |
| LC_50m_3um_k141_238018_9 | LC_50m_3um_k141_239166 | 6534 | 7067 | - | hypothetical protein | hypothetical protein |
| LC_50m_3um_k141_468226_1 | LC_50m_3um_k141_468226 | 3 | 554 | + | hypothetical protein | hypothetical protein |
| LC_50m_3um_k141_468226_2 | LC_50m_3um_k141_468226 | 693 | 1337 | + | hypothetical protein | hypothetical protein |
| LC_50m_3um_k141_468226_3 | LC_50m_3um_k141_468226 | 1338 | 3743 | + | hypothetical protein | hypothetical protein |
| LC_50m_3um_k141_468226_4 | LC_50m_3um_k141_468226 | 3762 | 4226 | + | hypothetical protein | hypothetical protein |
| LC_50m_3um_k141_468226_5 | LC_50m_3um_k141_468226 | 4434 | 4676 | - | hypothetical protein | hypothetical protein |
| LC_50m_3um_k141_468226_6 | LC_50m_3um_k141_468226 | 4767 | 7202 | + | RDRP | RDRP |
| LC_50m_3um_k141_325294_1 | LC_50m_3um_k141_325294 | 475 | 3024 | + | RDRP | RDRP |
| LC_50m_3um_k141_325294_2 | LC_50m_3um_k141_325294 | 3011 | 3202 | + | hypothetical protein | hypothetical protein |
| LC_50m_3um_k141_325294_3 | LC_50m_3um_k141_325294 | 3338 | 3424 | + | hypothetical protein | hypothetical protein |
| LC_80m_0.2um_k141_102393_1 | LC_80m_0.2um_k141_102393 | 1 | 2544 | + | hypothetical protein | hypothetical protein |
| LC_80m_0.2um_k141_102393_2 | LC_80m_0.2um_k141_102393 | 2665 | 3246 | + | hypothetical protein | hypothetical protein |
| LC_80m_0.2um_k141_102393_3 | LC_80m_0.2um_k141_102393 | 3532 | 3924 | + | hypothetical protein | hypothetical protein |
| LC_80m_0.2um_k141_102393_4 | LC_80m_0.2um_k141_102393 | 3955 | 5352 | + | RDRP | RDRP |
| LC_80m_0.2um_k141_456306_1 | LC_80m_0.2um_k141_456306 | 202 | 2364 | - | hypothetical protein | hypothetical protein |
| LC_80m_0.2um_k141_456306_2 | LC_80m_0.2um_k141_456306 | 2431 | 3732 | - | RDRP | RDRP |
| LC_80m_0.2um_k141_456306_3 | LC_80m_0.2um_k141_456306 | 3889 | 4299 | - | hypothetical protein | hypothetical protein |
| LC_80m_0.2um_k141_456306_4 | LC_80m_0.2um_k141_456306 | 4360 | 4695 | - | hypothetical protein | hypothetical protein |
| LC_80m_0.2um_k141_456306_5 | LC_80m_0.2um_k141_456306 | 4702 | 6324 | - | hypothetical protein | hypothetical protein |
| LC_80m_0.2um_k141_456306_6 | LC_80m_0.2um_k141_456306 | 6625 | 7560 | - | hypothetical protein | hypothetical protein |
| LC_80m_0.2um_k141_456306_7 | LC_80m_0.2um_k141_456306 | 7684 | 8220 | - | hypothetical protein | hypothetical protein |
| LC_coassembly_k141_1539147_1 | LC_coassembly_k141_1539147 | 1 | 1602 | - | capsid/coat protein | capsid/coat protein |
| LC_coassembly_k141_1539147_2 | LC_coassembly_k141_1539147 | 1934 | 4015 | - | RDRP | RDRP |
| LC_coassembly_k141_1539147_3 | LC_coassembly_k141_1539147 | 4088 | 4342 | - | hypothetical protein | hypothetical protein |
| LC_coassembly_k141_1148680_1 | LC_coassembly_k141_1148680 | 382 | 609 | - | hypothetical protein | hypothetical protein |
| LC_coassembly_k141_1148680_2 | LC_coassembly_k141_1148680 | 664 | 1209 | - | hypothetical protein | hypothetical protein |
| LC_coassembly_k141_1148680_3 | LC_coassembly_k141_1148680 | 1303 | 1926 | - | capsid/coat protein | capsid/coat protein |
| LC_coassembly_k141_1148680_4 | LC_coassembly_k141_1148680 | 1996 | 4038 | - | RDRP | RDRP |
| LC_coassembly_k141_1148680_5 | LC_coassembly_k141_1148680 | 4057 | 4497 | - | hypothetical protein | hypothetical protein |
| LC_coassembly_k141_1148680_6 | LC_coassembly_k141_1148680 | 4597 | 4815 | + | hypothetical protein | hypothetical protein |
| LC_coassembly_k141_1148680_7 | LC_coassembly_k141_1148680 | 5146 | 5562 | - | hypothetical protein | hypothetical protein |
| LC_coassembly_k141_1148680_8 | LC_coassembly_k141_1148680 | 5758 | 6285 | + | hypothetical protein | hypothetical protein |
| LC_coassembly_k141_1148680_9 | LC_coassembly_k141_1148680 | 6317 | 6592 | + | hypothetical protein | hypothetical protein |
| LC_coassembly_k141_1148680_10 | LC_coassembly_k141_1148680 | 6589 | 6861 | - | hypothetical protein | hypothetical protein |
| LC_coassembly_k141_1148680_11 | LC_coassembly_k141_1148680 | 6925 | 7251 | - | hypothetical protein | hypothetical protein |
| LC_coassembly_k141_1148680_12 | LC_coassembly_k141_1148680 | 7339 | 7575 | - | hypothetical protein | hypothetical protein |
| LC_coassembly_k141_1148680_13 | LC_coassembly_k141_1148680 | 8242 | 8730 | - | hypothetical protein | hypothetical protein |
| LC_coassembly_k141_1148680_14 | LC_coassembly_k141_1148680 | 9346 | 9666 | - | hypothetical protein | hypothetical protein |
| LC_coassembly_k141_1148680_15 | LC_coassembly_k141_1148680 | 9757 | 10038 | - | hypothetical protein | hypothetical protein |
| LC_coassembly_k141_1148680_16 | LC_coassembly_k141_1148680 | 10147 | 10695 | - | hypothetical protein | hypothetical protein |
| LC_coassembly_k141_1148680_17 | LC_coassembly_k141_1148680 | 10960 | 11085 | - | hypothetical protein | hypothetical protein |
| LC_coassembly_k141_1148680_18 | LC_coassembly_k141_1148680 | 11161 | 11802 | - | hypothetical protein | hypothetical protein |
| LC_coassembly_k141_1148680_19 | LC_coassembly_k141_1148680 | 11821 | 12057 | + | hypothetical protein | hypothetical protein |
| LC_80m_0.2um_k141_81878_1 | LC_80m_0.2um_k141_81878 | 3 | 635 | - | hypothetical protein | hypothetical protein |
| LC_80m_0.2um_k141_81878_2 | LC_80m_0.2um_k141_81878 | 702 | 2342 | - | RDRP | RDRP |
| LC_80m_0.2um_k141_81878_3 | LC_80m_0.2um_k141_81878 | 2343 | 3092 | - | hypothetical protein | hypothetical protein |
| LC_80m_0.2um_k141_81878_4 | LC_80m_0.2um_k141_81878 | 3174 | 7421 | - | hypothetical protein | hypothetical protein |
| LC_80m_10um_k141_267292_1 | LC_80m_10um_k141_267292 | 1 | 1776 | - | RDRP | RDRP |
| LC_80m_10um_k141_267292_2 | LC_80m_10um_k141_267292 | 1827 | 2834 | - | hypothetical protein | hypothetical protein |
| LC_80m_10um_k141_267292_3 | LC_80m_10um_k141_267292 | 2884 | 3264 | - | hypothetical protein | hypothetical protein |
| LS_0.5m_1.2um_k141_463252_1 | LS_0.5m_1.2um_k141_463252 | 2 | 100 | + | hypothetical protein | hypothetical protein |
| LS_0.5m_1.2um_k141_463252_2 | LS_0.5m_1.2um_k141_463252 | 107 | 574 | - | capsid/coat protein | capsid/coat protein |
| LS_0.5m_1.2um_k141_463252_3 | LS_0.5m_1.2um_k141_463252 | 697 | 1545 | - | hypothetical protein | hypothetical protein |
| LS_0.5m_1.2um_k141_463252_4 | LS_0.5m_1.2um_k141_463252 | 1655 | 2842 | - | Helper-component-proteinase | other viral proteins |
| LS_0.5m_1.2um_k141_463252_5 | LS_0.5m_1.2um_k141_463252 | 2870 | 9538 | - | RDRP | RDRP |
| LS_0.5m_1.2um_k141_463252_6 | LS_0.5m_1.2um_k141_463252 | 9641 | 9850 | + | hypothetical protein | hypothetical protein |
| LS_0.5m_3um_k141_388117_1 | LS_0.5m_3um_k141_388117 | 41 | 508 | - | capsid/coat protein | capsid/coat protein |
| LS_0.5m_3um_k141_388117_2 | LS_0.5m_3um_k141_388117 | 565 | 1446 | - | hypothetical protein | hypothetical protein |
| LS_0.5m_3um_k141_388117_3 | LS_0.5m_3um_k141_388117 | 1538 | 2755 | - | Helper-component-proteinase | other viral proteins |
| LS_0.5m_3um_k141_388117_4 | LS_0.5m_3um_k141_388117 | 2783 | 9592 | - | RDRP | RDRP |
| LS_coassembly_k141_1033411_1 | LS_coassembly_k141_1033411 | 291 | 7100 | + | RDRP | RDRP |
| LS_coassembly_k141_1033411_2 | LS_coassembly_k141_1033411 | 7128 | 8345 | + | Helper-component-proteinase | other viral proteins |
| LS_coassembly_k141_1033411_3 | LS_coassembly_k141_1033411 | 8437 | 9318 | + | hypothetical protein | hypothetical protein |
| LS_coassembly_k141_1033411_4 | LS_coassembly_k141_1033411 | 9375 | 9842 | + | capsid/coat protein | capsid/coat protein |
| LC_50m_3um_k141_133114_1 | LC_50m_3um_k141_133114 | 62 | 538 | - | capsid/coat protein | capsid/coat protein |
| LC_50m_3um_k141_133114_2 | LC_50m_3um_k141_133114 | 587 | 1471 | - | hypothetical protein | hypothetical protein |
| LC_50m_3um_k141_133114_3 | LC_50m_3um_k141_133114 | 1443 | 2633 | - | Helper-component-proteinase | other viral proteins |
| LC_50m_3um_k141_133114_4 | LC_50m_3um_k141_133114 | 2657 | 9511 | - | RDRP | RDRP |
| LC_coassembly_k141_1476401_1 | LC_coassembly_k141_1476401 | 88 | 1731 | - | hypothetical protein | hypothetical protein |
| LC_coassembly_k141_1476401_2 | LC_coassembly_k141_1476401 | 1728 | 4151 | - | hypothetical protein | hypothetical protein |
| LC_coassembly_k141_1476401_3 | LC_coassembly_k141_1476401 | 4218 | 5510 | - | Peptidase-A6-family-protein | other viral proteins |
| LC_coassembly_k141_1476401_4 | LC_coassembly_k141_1476401 | 5458 | 6642 | - | hypothetical protein | hypothetical protein |
| LC_coassembly_k141_2580714_1 | LC_coassembly_k141_2580714 | 82 | 5295 | + | RDRP | RDRP |
| LC_coassembly_k141_2580714_2 | LC_coassembly_k141_2580714 | 5723 | 7210 | + | Peptidase-family-A21-protein | other viral proteins |
| LC_coassembly_k141_196466_1 | LC_coassembly_k141_196466 | 194 | 673 | - | capsid/coat protein | capsid/coat protein |
| LC_coassembly_k141_196466_2 | LC_coassembly_k141_196466 | 676 | 1482 | - | Viral-movement-protein-(MP) | other viral proteins |
| LC_coassembly_k141_196466_3 | LC_coassembly_k141_196466 | 1466 | 2923 | - | RDRP | RDRP |
| LC_coassembly_k141_196466_4 | LC_coassembly_k141_196466 | 2966 | 6307 | - | Replicase-large-subunit | other viral proteins |
| LC_0.5m_10um_k141_124243_1 | LC_0.5m_10um_k141_124243 | 2 | 205 | + | Viral methyltransferase | other viral proteins |
| LC_0.5m_10um_k141_124243_2 | LC_0.5m_10um_k141_124243 | 145 | 2058 | + | Replicase-large-subunit | other viral proteins |
| LC_0.5m_10um_k141_124243_3 | LC_0.5m_10um_k141_124243 | 2101 | 3558 | + | RDRP | RDRP |
| LC_0.5m_10um_k141_124243_4 | LC_0.5m_10um_k141_124243 | 3542 | 4348 | + | Viral-movement-protein-(MP) | other viral proteins |
| LC_0.5m_10um_k141_124243_5 | LC_0.5m_10um_k141_124243 | 4351 | 4830 | + | capsid/coat protein | capsid/coat protein |
| LS_coassembly_k141_947790_1 | LS_coassembly_k141_947790 | 63 | 1148 | + | capsid/coat protein | capsid/coat protein |
| LS_coassembly_k141_947790_2 | LS_coassembly_k141_947790 | 1162 | 1554 | + | head-protein | other viral proteins |
| LS_coassembly_k141_947790_3 | LS_coassembly_k141_947790 | 1662 | 3038 | + | RDRP | RDRP |
| LS_coassembly_k141_950304_1 | LS_coassembly_k141_950304 | 230 | 595 | - | hypothetical protein | hypothetical protein |
| LS_coassembly_k141_950304_2 | LS_coassembly_k141_950304 | 14055 | 14396 | - | hypothetical protein | hypothetical protein |
| LS_coassembly_k141_950304_3 | LS_coassembly_k141_950304 | 14560 | 15996 | - | nucleoprotein | other viral proteins |
| LS_coassembly_k141_950304_4 | LS_coassembly_k141_950304 | 725 | 7069 | - | RDRP | RDRP |
| LS_coassembly_k141_950304_5 | LS_coassembly_k141_950304 | 7168 | 7383 | - | hypothetical protein | hypothetical protein |
| LS_coassembly_k141_950304_6 | LS_coassembly_k141_950304 | 7464 | 9452 | - | Glycoprotein- | other viral proteins |
| LS_coassembly_k141_950304_7 | LS_coassembly_k141_950304 | 9650 | 10363 | - | hypothetical protein | hypothetical protein |
| LS_coassembly_k141_950304_8 | LS_coassembly_k141_950304 | 10511 | 11227 | - | hypothetical protein | hypothetical protein |
| LS_coassembly_k141_950304_9 | LS_coassembly_k141_950304 | 11371 | 12291 | - | hypothetical protein | hypothetical protein |
| LS_coassembly_k141_950304_10 | LS_coassembly_k141_950304 | 12382 | 13509 | - | hypothetical protein | hypothetical protein |
| LS_coassembly_k141_950304_11 | LS_coassembly_k141_950304 | 13606 | 14010 | - | hypothetical protein | hypothetical protein |
| LC_coassembly_k141_1176970_1 | LC_coassembly_k141_1176970 | 180 | 617 | - | hypothetical protein | hypothetical protein |
| LC_coassembly_k141_1176970_2 | LC_coassembly_k141_1176970 | 617 | 2908 | - | hypothetical protein | hypothetical protein |
| LC_coassembly_k141_1176970_3 | LC_coassembly_k141_1176970 | 2905 | 3777 | - | packaging-NTPase-P4 | other viral proteins |
| LC_coassembly_k141_1176970_4 | LC_coassembly_k141_1176970 | 3777 | 5606 | - | RDRP | RDRP |
| LC_coassembly_k141_1176970_5 | LC_coassembly_k141_1176970 | 5603 | 6208 | - | hypothetical protein | hypothetical protein |
| LC_coassembly_k141_1176970_6 | LC_coassembly_k141_1176970 | 6593 | 6733 | - | hypothetical protein | hypothetical protein |
| LS_coassembly_k141_409312_1 | LS_coassembly_k141_409312 | 345 | 3233 | - | RDRP | RDRP |
| LS_coassembly_k141_409312_2 | LS_coassembly_k141_409312 | 3264 | 6491 | - | structural-polyprotein | other viral proteins |
| LS_0.5m_10um_k141_12963_1 | LS_0.5m_10um_k141_12963 | 285 | 881 | - | capsid/coat protein | capsid/coat protein |
| LS_0.5m_10um_k141_12963_2 | LS_0.5m_10um_k141_12963 | 853 | 2079 | - | RDRP | RDRP |
| LS_0.5m_10um_k141_12963_3 | LS_0.5m_10um_k141_12963 | 2079 | 4043 | - | Peptidase-S39 | other viral proteins |
| LS_0.5m_10um_k141_12963_4 | LS_0.5m_10um_k141_12963 | 3985 | 4347 | - | hypothetical protein | hypothetical protein |
| LC_coassembly_k141_86982_1 | LC_coassembly_k141_86982 | 32 | 1582 | + | Peptidase-S39 | other viral proteins |
| LC_coassembly_k141_86982_2 | LC_coassembly_k141_86982 | 1747 | 2802 | + | RDRP | RDRP |
| LC_coassembly_k141_86982_3 | LC_coassembly_k141_86982 | 3258 | 4103 | + | capsid/coat protein | capsid/coat protein |
| LC_coassembly_k141_2159845_1 | LC_coassembly_k141_2159845 | 135 | 497 | + | hypothetical protein | hypothetical protein |
| LC_coassembly_k141_2159845_2 | LC_coassembly_k141_2159845 | 439 | 2403 | + | Peptidase-S39 | other viral proteins |
| LC_coassembly_k141_2159845_3 | LC_coassembly_k141_2159845 | 2403 | 3629 | + | RDRP | RDRP |
| LC_coassembly_k141_2159845_4 | LC_coassembly_k141_2159845 | 3601 | 4197 | + | capsid/coat protein | capsid/coat protein |
| LS_coassembly_k141_389387_1 | LS_coassembly_k141_389387 | 60 | 1346 | - | RDRP | RDRP |
| LS_coassembly_k141_389387_2 | LS_coassembly_k141_389387 | 1349 | 2761 | - | Peptidase-S39 | other viral proteins |
| LC_20m_0.2um_k141_269092_1 | LC_20m_0.2um_k141_269092 | 358 | 7212 | + | RDRP | RDRP |
| LC_20m_0.2um_k141_269092_2 | LC_20m_0.2um_k141_269092 | 7236 | 8426 | + | Helper-component-proteinase | other viral proteins |
| LC_20m_0.2um_k141_269092_3 | LC_20m_0.2um_k141_269092 | 8398 | 9282 | + | hypothetical protein | hypothetical protein |
| LC_20m_0.2um_k141_269092_4 | LC_20m_0.2um_k141_269092 | 9331 | 9807 | + | capsid/coat protein | capsid/coat protein |
| LC_20m_0.2um_k141_269092_5 | LC_20m_0.2um_k141_269092 | 9919 | 11082 | + | bifunctional uroporphyrinogen-III synthetase/uroporphyrin-III C-methyltransferase | AMGs |
| LC_20m_0.2um_k141_269092_6 | LC_20m_0.2um_k141_269092 | 11079 | 11882 | + | heme biosynthesis-associated TPR protein | AMGs |
| LC_coassembly_k141_945938_1 | LC_coassembly_k141_945938 | 1 | 174 | + | ATP synthase CF1 alpha subunit | AMGs |
| LC_coassembly_k141_945938_2 | LC_coassembly_k141_945938 | 735 | 3740 | + | RDRP | RDRP |
| LC_coassembly_k141_945938_3 | LC_coassembly_k141_945938 | 3993 | 5045 | + | hypothetical protein | hypothetical protein |
| LC_20m_3um_k141_357943_1 | LC_20m_3um_k141_357943 | 90 | 1781 | - | RDRP | RDRP |
| LC_20m_3um_k141_357943_2 | LC_20m_3um_k141_357943 | 1833 | 3779 | - | hypothetical protein | hypothetical protein |
| LC_20m_3um_k141_357943_3 | LC_20m_3um_k141_357943 | 3927 | 4802 | - | hypothetical protein | hypothetical protein |
| LC_20m_3um_k141_357943_4 | LC_20m_3um_k141_357943 | 4981 | 5715 | - | glycerophosphoryl diester phosphodiesterase | AMGs |
| LC_20m_3um_k141_357943_5 | LC_20m_3um_k141_357943 | 5836 | 7161 | - | capsid/coat protein | capsid/coat protein |
| LC_coassembly_k141_1021355_1 | LC_coassembly_k141_1021355 | 130 | 1443 | + | APIS022_anti-prokaryotic immune system protein | AMGs |
| LC_coassembly_k141_1021355_2 | LC_coassembly_k141_1021355 | 1819 | 3411 | + | hypothetical protein | hypothetical protein |
| LC_coassembly_k141_1021355_3 | LC_coassembly_k141_1021355 | 3427 | 4542 | + | hypothetical protein | hypothetical protein |
| LC_coassembly_k141_1021355_4 | LC_coassembly_k141_1021355 | 4807 | 6225 | + | hypothetical protein | hypothetical protein |
| LC_coassembly_k141_1021355_5 | LC_coassembly_k141_1021355 | 6247 | 7020 | + | RDRP | RDRP |
| LC_coassembly_k141_1021355_6 | LC_coassembly_k141_1021355 | 7117 | 7782 | + | hypothetical protein | hypothetical protein |
| LC_0.5m_1.2um_k141_396381_1 | LC_0.5m_1.2um_k141_396381 | 92 | 271 | + | hypothetical protein | hypothetical protein |
| LC_0.5m_1.2um_k141_396381_2 | LC_0.5m_1.2um_k141_396381 | 623 | 2758 | - | hypothetical protein | hypothetical protein |
| LC_0.5m_1.2um_k141_396381_3 | LC_0.5m_1.2um_k141_396381 | 2855 | 3628 | - | RDRP | RDRP |
| LC_0.5m_1.2um_k141_396381_4 | LC_0.5m_1.2um_k141_396381 | 3650 | 5068 | - | hypothetical protein | hypothetical protein |
| LC_0.5m_1.2um_k141_396381_5 | LC_0.5m_1.2um_k141_396381 | 5333 | 6448 | - | hypothetical protein | hypothetical protein |
| LC_0.5m_1.2um_k141_396381_6 | LC_0.5m_1.2um_k141_396381 | 6464 | 8056 | - | hypothetical protein | hypothetical protein |
| LC_0.5m_1.2um_k141_396381_7 | LC_0.5m_1.2um_k141_396381 | 8432 | 9745 | - | APIS022_anti-prokaryotic immune system protein | AMGs |
| LC_coassembly_k141_2670924_1 | LC_coassembly_k141_2670924 | 2 | 148 | + | ribosomal protein L1 | AMGs |
| LC_coassembly_k141_2670924_2 | LC_coassembly_k141_2670924 | 181 | 1251 | + | RDRP | RDRP |
| LS_coassembly_k141_232625_1 | LS_coassembly_k141_232625 | 5 | 1072 | - | RDRP | RDRP |
| LS_coassembly_k141_232625_2 | LS_coassembly_k141_232625 | 1630 | 1920 | - | C-terminal domain of Cytochrome c Oxidase subunit II | AMGs |
| LC_20m_3um_k141_102044_1 | LC_20m_3um_k141_102044 | 3 | 497 | + | 345341.KUTG_04140 | AMGs |
| LC_20m_3um_k141_102044_2 | LC_20m_3um_k141_102044 | 460 | 1953 | - | RDRP | RDRP |

| **Table S7. Detail information about host annotation for RNA vOTUs from Lake Nam Co.** | | | | | | | | | | | | | |
| --- | --- | --- | --- | --- | --- | --- | --- | --- | --- | --- | --- | --- | --- |
| **Contig** | **Phylum** | **Class** | **Order** | **Family** | **Kingdom  of host** | **Phylum  of host** | **Class of  host** | **Order of  host** | **Family  of host** | **Genus  of host** | **Identification  method** | **^1^Confidence  score** | **^2^E-value** |
| LC_coassembly_k141_2104764 | *Kitrinoviricota* | *Alsuviricetes* | *Martellivirales* | *Virgaviridae* | Eukaryota | *Plantae* | *Magnoliopsida* | *Solanales* | *Solanaceae* | *Capsicum* | Related virus | NA | NA |
| LC_coassembly_k141_2315390 | *Lenarviricota* | *Leviviricetes* | *Norzivirales* | *Atkinsviridae* | Bacteria | NA | NA | NA | NA | NA | Related virus | NA | NA |
| LC_coassembly_k141_1805655 | *Lenarviricota* | *Leviviricetes* | *Norzivirales* | *Fiersviridae* | Bacteria | NA | NA | NA | NA | NA | Related virus | NA | NA |
| LC_coassembly_k141_635711 | *Lenarviricota* | *Leviviricetes* | *Timlovirales* | *Steitzviridae* | Bacteria | NA | NA | NA | NA | NA | Related virus | NA | NA |
| LC_coassembly_k141_1177772 | *Lenarviricota* | *Leviviricetes* | *Norzivirales* | *Atkinsviridae* | Bacteria | NA | NA | NA | NA | NA | Related virus | NA | NA |
| LC_coassembly_k141_2772377 | *Kitrinoviricota* | *Alsuviricetes* | *o.0055* | *Endornaviridae* | Eukaryota | *Plantae* | *Magnoliopsida* | *Poales* | *Poaceae* | *Oryza* | Related virus | NA | NA |
| LC_coassembly_k141_1271871 | *Lenarviricota* | *Leviviricetes* | *Timlovirales* | *Steitzviridae* | Bacteria | NA | NA | NA | NA | NA | Related virus | NA | NA |
| LC_coassembly_k141_1394068 | *Pisuviricota* | *Pisoniviricetes* | *Picornavirales* | *Picornaviridae* | Eukaryota | *Animalia* | *Mammalia* | *Primates* | *Hominidae* | *Homo* | Related virus | NA | NA |
| LC_coassembly_k141_2477916 | *Duplornaviricota* | *Chrymotiviricetes* | *Ghabrivirales* | *f.0283.base-Megabirna* | Bacteria | *Bacteroidota* | *Bacteroidia* | *Marinilabiliales* | *Marinifilaceae* | *Labilibaculum* | iPHoP-RF | 96.1 | NA |
| LC_coassembly_k141_1817896 | *Lenarviricota* | *Leviviricetes* | *Norzivirales* | *Atkinsviridae* | Bacteria | NA | NA | NA | NA | NA | Related virus | NA | NA |
| LC_coassembly_k141_196466 | *Kitrinoviricota* | *Alsuviricetes* | *Martellivirales* | *Virgaviridae* | Eukaryota | *Plantae* | *Magnoliopsida* | *Solanales* | *Solanaceae* | *Nicotiana* | Related virus | NA | NA |
| LC_coassembly_k141_1968509 | *Lenarviricota* | *Leviviricetes* | *Norzivirales* | *Atkinsviridae* | Bacteria | NA | NA | NA | NA | NA | Related virus | NA | NA |
| LC_coassembly_k141_2570745 | *Negarnaviricota* | *NA* | *NA* | *NA* | Bacteria | *Pseudomonadota* | *Alphaproteobacteria* | *Rickettsiales* | *Midichloriaceae* | *SZAS-26* | iPHoP-RF | 92.1 | NA |
| LC_coassembly_k141_2541618 | *Lenarviricota* | *Leviviricetes* | *Timlovirales* | *Steitzviridae* | Bacteria | NA | NA | NA | NA | NA | Related virus | NA | NA |
| LC_coassembly_k141_2783551 | *Duplornaviricota* | *Chrymotiviricetes* | *Ghabrivirales* | *Totiviridae* | Bacteria | *Actinomycetota* | *Acidimicrobiia* | *Acidimicrobiales* | *Ilumatobacteraceae* | *UBA2NA93* | iPHoP-RF | 91.4 | NA |
| LC_coassembly_k141_1763940 | *Duplornaviricota* | *Chrymotiviricetes* | *Ghabrivirales* | *f.0283.base-Megabirna* | Bacteria | *Bacteroidota* | *Bacteroidia* | *Bacteroidales* | *Bacteroidaceae* | *Prevotella* | iPHoP-RF | 92.1 | NA |
| LC_coassembly_k141_1675587 | *Pisuviricota* | *Duplopiviricetes* | *Durnavirales* | *f.0102.base-Curvula* | Bacteria | *Bacteroidota* | *Bacteroidia* | *Flavobacteriales* | *Flavobacteriaceae* | *Yeosuana* | iPHoP-RF | 91.1 | NA |
| LC_coassembly_k141_864888 | *Lenarviricota* | *Leviviricetes* | *Norzivirales* | *f.0363* | Bacteria | NA | NA | NA | NA | NA | Related virus | NA | NA |
| LC_coassembly_k141_1827790 | *Lenarviricota* | *Leviviricetes* | *Norzivirales* | *Fiersviridae* | Bacteria | NA | NA | NA | NA | NA | Related virus | NA | NA |
| LC_coassembly_k141_2488396 | *Lenarviricota* | *Howeltoviricetes* | *Cryppavirales* | *Mitoviridae* | Bacteria | *Bacteroidota* | *Bacteroidia* | *Flavobacteriales* | *Flavobacteriaceae* | *Nonlabens* | iPHoP-RF | 97.1 | NA |
| LC_coassembly_k141_868598 | *Pisuviricota* | *Duplopiviricetes* | *Durnavirales* | *Picobirnaviridae* | Bacteria | NA | NA | NA | NA | NA | Related virus | NA | NA |
| LC_coassembly_k141_2371561 | *Lenarviricota* | *Amabiliviricetes* | *Wolframvirales* | *Narnaviridae* | Bacteria | *Bacillota* | *Bacteroidales* | *Desulfitobacteriales* | *Desulfitobacteriaceae* | *Desulfosporosinus* | iPHoP-RF | 92.4 | NA |
| LS_coassembly_k141_1183900 | *Lenarviricota* | *Leviviricetes* | *Norzivirales* | *f.0361.base-Solspi* | Bacteria | NA | NA | NA | NA | NA | Related virus | NA | NA |
| LS_coassembly_k141_656443 | *Lenarviricota* | *Leviviricetes* | *Norzivirales* | *Fiersviridae* | Bacteria | NA | NA | NA | NA | NA | Related virus | NA | NA |
| LS_coassembly_k141_542386 | *Lenarviricota* | *Leviviricetes* | *Timlovirales* | *Steitzviridae* | Bacteria | NA | NA | NA | NA | NA | Related virus | NA | NA |
| LS_coassembly_k141_543237 | *Lenarviricota* | *Leviviricetes* | *Norzivirales* | *Fiersviridae* | Bacteria | NA | NA | NA | NA | NA | Related virus | NA | NA |
| LS_coassembly_k141_357820 | *Pisuviricota* | *c.0013* | *o.0025.base-Sobeli* | *f.0076* | Bacteria | *Bacteroidota* | *Bacteroidia* | *Bacteroidales* | *Bacteroidaceae* | *Prevotella* | CRISPR-Cas evidence from iPHoP | 91.3 | 4.56E-06 |
| LS_coassembly_k141_187532 | *Pisuviricota* | *Pisoniviricetes* | *Picornavirales* | *Iflaviridae* | Eukaryota | *Animalia* | *Insecta* | *Hemiptera* | *Aphididae* | *Brevicoryne* | Related virus | NA | NA |
| LS_coassembly_k141_287825 | *Negarnaviricota* | *Monjiviricetes* | *Mononegavirales* | *Rhabdoviridae* | Eukaryota | *Animalia* | *Insecta* | *Hemiptera* | NA | NA | Related virus | NA | NA |
| LS_coassembly_k141_259450 | *Lenarviricota* | *Leviviricetes* | *Norzivirales* | *Fiersviridae* | Bacteria | NA | NA | NA | NA | NA | Related virus | NA | NA |
| LS_coassembly_k141_631699 | *Lenarviricota* | *Leviviricetes* | *Timlovirales* | *Steitzviridae* | Bacteria | NA | NA | NA | NA | NA | Related virus | NA | NA |
| LS_coassembly_k141_232625 | *Pisuviricota* | *Duplopiviricetes* | *Durnavirales* | *f.0102.base-Curvula* | Eukaryota | *Oomycota* | *Oomycetes* | *Saprolegniales* | *Saprolegniaceae* | *Achlya* | Host gene | NA | 5.98E-50 |
| LS_coassembly_k141_4481 | *Lenarviricota* | *Leviviricetes* | *Norzivirales* | *Solspiviridae* | Bacteria | NA | NA | NA | NA | NA | Related virus | NA | NA |
| LS_coassembly_k141_1274756 | *Pisuviricota* | *Duplopiviricetes* | *Durnavirales* | *f.0102.base-Curvula* | Bacteria | *Bacillota* | *Bacilli* | *RF39* | *UBA660* | *RGIG8386* | iPHoP-RF | 97.1 | NA |
| LS_coassembly_k141_447615 | *Lenarviricota* | *Leviviricetes* | *Norzivirales* | *Fiersviridae* | Bacteria | NA | NA | NA | NA | NA | Related virus | NA | NA |
| LS_coassembly_k141_947790 | *Lenarviricota* | *Leviviricetes* | *Norzivirales* | *Fiersviridae* | Bacteria | NA | NA | NA | NA | NA | Related virus | NA | NA |
| LS_coassembly_k141_63722 | *Lenarviricota* | *Leviviricetes* | *Norzivirales* | *Fiersviridae* | Bacteria | NA | NA | NA | NA | NA | Related virus | NA | NA |
| LS_coassembly_k141_379730 | *Lenarviricota* | *Leviviricetes* | *Norzivirales* | *f.0368.base-Atkins* | Bacteria | NA | NA | NA | NA | NA | Related virus | NA | NA |
| LS_coassembly_k141_8864 | *Lenarviricota* | *Leviviricetes* | *Norzivirales* | *Fiersviridae* | Bacteria | NA | NA | NA | NA | NA | Related virus | NA | NA |
| LS_coassembly_k141_1121902 | *Lenarviricota* | *Leviviricetes* | *Norzivirales* | *Fiersviridae* | Bacteria | NA | NA | NA | NA | NA | Related virus | NA | NA |
| LS_coassembly_k141_152525 | *Lenarviricota* | *Leviviricetes* | *Timlovirales* | *Steitzviridae* | Bacteria | NA | NA | NA | NA | NA | Related virus | NA | NA |
| LS_coassembly_k141_808804 | *Lenarviricota* | *Leviviricetes* | *Norzivirales* | *Fiersviridae* | Bacteria | NA | NA | NA | NA | NA | Related virus | NA | NA |
| LS_coassembly_k141_795198 | *Pisuviricota* | *Stelpaviricetes* | *Patatavirales* | *Potyviridae* | Eukaryota | *Plantae* | *Magnoliopsida* | *Fabales* | *Fabaceae* | *Vigna* | Related virus | NA | NA |
| LS_coassembly_k141_709523 | *Lenarviricota* | *Leviviricetes* | *Norzivirales* | *Fiersviridae* | Bacteria | NA | NA | NA | NA | NA | Related virus | NA | NA |
| LS_coassembly_k141_638690 | *Lenarviricota* | *Leviviricetes* | *Timlovirales* | *Steitzviridae* | Bacteria | NA | NA | NA | NA | NA | Related virus | NA | NA |
| LS_coassembly_k141_210683 | *Kitrinoviricota* | *Alsuviricetes* | *o.0055* | *Endornaviridae* | Eukaryota | *Plantae* | *Magnoliopsida* | *Cucurbitales* | *Cucurbitaceae* | *Cucumis* | Related virus | NA | NA |
| LS_coassembly_k141_824362 | *Lenarviricota* | *Leviviricetes* | *Timlovirales* | *Steitzviridae* | Bacteria | NA | NA | NA | NA | NA | Related virus | NA | NA |
| LS_coassembly_k141_981221 | *Lenarviricota* | *Leviviricetes* | *Norzivirales* | *f.0361.base-Solspi* | Bacteria | NA | NA | NA | NA | NA | Related virus | NA | NA |
| LS_coassembly_k141_1166806 | *Lenarviricota* | *Leviviricetes* | *Timlovirales* | *Steitzviridae* | Bacteria | NA | NA | NA | NA | NA | Related virus | NA | NA |
| LS_coassembly_k141_169245 | *Lenarviricota* | *Leviviricetes* | *Norzivirales* | *f.0367.base-Atkins* | Bacteria | NA | NA | NA | NA | NA | Related virus | NA | NA |
| LS_coassembly_k141_839853 | *Lenarviricota* | *Leviviricetes* | *Timlovirales* | *Steitzviridae* | Bacteria | NA | NA | NA | NA | NA | Related virus | NA | NA |
| LS_coassembly_k141_1338665 | *Lenarviricota* | *Leviviricetes* | *Norzivirales* | *f.0368.base-Atkins* | Bacteria | NA | NA | NA | NA | NA | Related virus | NA | NA |
| LS_coassembly_k141_839961 | *Lenarviricota* | *Leviviricetes* | *Norzivirales* | *Fiersviridae* | Bacteria | NA | NA | NA | NA | NA | Related virus | NA | NA |
| LS_coassembly_k141_112997 | *Lenarviricota* | *Leviviricetes* | *Norzivirales* | *f.0368.base-Atkins* | Bacteria | NA | NA | NA | NA | NA | Related virus | NA | NA |
| LS_coassembly_k141_370211 | *Lenarviricota* | *Leviviricetes* | *Timlovirales* | *Steitzviridae* | Bacteria | NA | NA | NA | NA | NA | Related virus | NA | NA |
| LS_coassembly_k141_1097885 | *Pisuviricota* | *Pisoniviricetes* | *Picornavirales* | *Marnaviridae* | Archaea | *Methanobacteriota* | *Methanobacteria* | *Methanobacteriales* | *Methanothermobacteraceae* | *Methanothermobacter* | iPHoP-RF | 94.7 | NA |
| LC_0.5m_10um_k141_124243 | *Kitrinoviricota* | *Alsuviricetes* | *Martellivirales* | *Virgaviridae* | Eukaryota | *Plantae* | *Magnoliopsida* | *Solales* | *Solaceae* | *Nicotiana* | Related virus | NA | NA |
| LC_20m_0.2um_k141_269092 | *Kitrinoviricota* | *Alsuviricetes* | *Hepelivirales* | *Benyviridae* | Bacteria | *Pseudomonadota* | *Betaproteobacteria* | *Burkholderiales* | *Comamonadaceae* | *Hydrogenophaga* | Host gene | 93.6 | 2.25E-83 |
| LC_20m_0.2um_k141_261093 | *Duplornaviricota* | *Chrymotiviricetes* | *Ghabrivirales* | *f.0295* | Bacteria | *Bacillota* | *Bacilli* | *RFN20* | *CAG-826* | *RUG131* | iPHoP-RF | 93.1 | NA |
| LC_20m_0.2um_k141_260174 | *Pisuviricota* | *Pisoniviricetes* | *Picornavirales* | *f.0028.base-Dicistro* | Bacteria | *Bacteroidota* | *Bacteroidia* | *Flavobacteriales* | *Flavobacteriaceae* | *Gelidibacter* | iPHoP-RF | 97.1 | NA |
| LC_20m_10um_k141_27699 | *Duplornaviricota* | *Chrymotiviricetes* | *Ghabrivirales* | *f.0296* | Bacteria | *Bacteroidota* | *Bacteroidia* | *Bacteroidales* | *Paludibacteraceae* | *UBA1181* | iPHoP-RF | 90.8 | NA |
| LC_20m_10um_k141_92587 | *Duplornaviricota* | *Chrymotiviricetes* | *Ghabrivirales* | *f.0283.base-Megabirna* | Bacteria | *Bacteroidota* | *Bacteroidia* | *Bacteroidales* | *Paludibacteraceae* | *Paludibacter* | iPHoP-RF | 94.7 | NA |
| LC_20m_10um_k141_33231 | *Duplornaviricota* | *Chrymotiviricetes* | *Ghabrivirales* | *f.0296* | Bacteria | *Pseudomonadota* | *Gammaproteobacteria* | *GCA-002705445* | *GCA-002716945* | *CACLCVNA1* | iPHoP-RF | 95.1 | NA |
| LC_20m_3um_k141_102044 | *Pisuviricota* | *Duplopiviricetes* | *Durnavirales* | *f.0111* | Bacteria | *Actinomycetota* | *Actinomycetes* | *Mycobacteriales* | *Mycobacteriaceae* | *Mycobacterium* | Host gene | NA | 8.66E-11 |
| LC_20m_3um_k141_555582 | *Pisuviricota* | *Duplopiviricetes* | *Durnavirales* | *Picobirnaviridae* | Bacteria | NA | NA | NA | NA | NA | Related virus | NA | NA |
| LC_20m_3um_k141_494413 | *Lenarviricota* | *Howeltoviricetes* | *Cryppavirales* | *Mitoviridae* | Bacteria | *Bacteroidota* | *Bacteroidia* | *Flavobacteriales* | *Flavobacteriaceae* | *Muricauda* | iPHoP-RF | 90.5 | NA |
| LC_20m_3um_k141_11485 | *Duplornaviricota* | *Chrymotiviricetes* | *Ghabrivirales* | *f.0286* | Bacteria | *Pseudomonadota* | *Gammaproteobacteria* | *Burkholderiales* | *Gallionellaceae* | *Gallionella* | iPHoP-RF | 91.4 | NA |
| LC_50m_1.2um_k141_154296 | *Kitrinoviricota* | *c.0034* | *o.0045* | *f.0176* | Bacteria | *Pseudomonadota* | *Gammaproteobacteria* | *Pseudomonadales* | *Pseudomonadaceae* | *Pseudomonas* | iPHoP-RF | 92.4 | NA |
| LC_80m_0.2um_k141_400659 | *Lenarviricota* | *Leviviricetes* | *Norzivirales* | *Atkinsviridae* | Bacteria | NA | NA | NA | NA | NA | Related virus | NA | NA |
| LS_0.5m_0.2um_k141_202953 | *Lenarviricota* | *Leviviricetes* | *Norzivirales* | *Fiersviridae* | Bacteria | NA | NA | NA | NA | NA | Related virus | NA | NA |
| LS_0.5m_0.2um_k141_97488 | *Lenarviricota* | *Leviviricetes* | *Norzivirales* | *f.0361.base-Solspi* | Bacteria | NA | NA | NA | NA | NA | Related virus | NA | NA |
| LS_0.5m_0.2um_k141_180478 | *Lenarviricota* | *Leviviricetes* | *Norzivirales* | *Solspiviridae* | Bacteria | NA | NA | NA | NA | NA | Related virus | NA | NA |
| LS_0.5m_1.2um_k141_16205 | *Lenarviricota* | *Leviviricetes* | *Timlovirales* | *Steitzviridae* | Bacteria | NA | NA | NA | NA | NA | Related virus | NA | NA |
| LS_0.5m_1.2um_k141_197594 | *Lenarviricota* | *Leviviricetes* | *Norzivirales* | *Fiersviridae* | Bacteria | NA | NA | NA | NA | NA | Related virus | NA | NA |
| LS_0.5m_1.2um_k141_627694 | *Lenarviricota* | *Leviviricetes* | *Norzivirales* | *Fiersviridae* | Bacteria | NA | NA | NA | NA | NA | Related virus | NA | NA |
| LS_0.5m_1.2um_k141_221766 | *Lenarviricota* | *Leviviricetes* | *Norzivirales* | *f.0361.base-Solspi* | Bacteria | NA | NA | NA | NA | NA | Related virus | NA | NA |
| LS_0.5m_1.2um_k141_623158 | *Lenarviricota* | *Leviviricetes* | *Norzivirales* | *f.0361.base-Solspi* | Bacteria | NA | NA | NA | NA | NA | Related virus | NA | NA |
| LS_0.5m_1.2um_k141_96485 | *Lenarviricota* | *Leviviricetes* | *Norzivirales* | *f.0361.base-Solspi* | Bacteria | NA | NA | NA | NA | NA | Related virus | NA | NA |
| LS_0.5m_1.2um_k141_585706 | *Lenarviricota* | *Leviviricetes* | *Norzivirales* | *Fiersviridae* | Bacteria | NA | NA | NA | NA | NA | Related virus | NA | NA |
| LS_0.5m_1.2um_k141_346045 | *Lenarviricota* | *Leviviricetes* | *Timlovirales* | *Steitzviridae* | Bacteria | NA | NA | NA | NA | NA | Related virus | NA | NA |
| LS_0.5m_3um_k141_360239 | *Lenarviricota* | *Leviviricetes* | *Norzivirales* | *Fiersviridae* | Bacteria | NA | NA | NA | NA | NA | Related virus | NA | NA |
| LS_0.5m_3um_k141_122486 | *Lenarviricota* | *Leviviricetes* | *Norzivirales* | *f.0368.base-Atkins* | Bacteria | NA | NA | NA | NA | NA | Related virus | NA | NA |
| LS_0.5m_3um_k141_37077 | *Lenarviricota* | *Leviviricetes* | *Norzivirales* | *f.0367.base-Atkins* | Bacteria | NA | NA | NA | NA | NA | Related virus | NA | NA |
| LC_coassembly_k141_1198831 | *Pisuviricota* | *Duplopiviricetes* | *Durnavirales* | *f.0111* | Bacteria | NA | NA | NA | NA | NA | CRISPR-Cas evidence from IMG | NA | 5.77E-31 |
| LC_80m_1.2um_k141_267137 | *Lenarviricota* | *Howeltoviricetes* | *Cryppavirales* | *Mitoviridae* | Bacteria | NA | NA | NA | NA | NA | CRISPR-Cas evidence from IMG | NA | 1.29E-08 |
| 1.Confidence score: The confidence score provided by the iPhop software | | | | | | | | | | | | | |
| 2.E-value: The e-value obtained from the CRISPR-Cas spacer or host gene alignment | | | | | | | | | | | | | |

| **Table S8. Detailed evidence regarding prokaryotic host annotation for RNA vOTUs from Lake Nam Co.** | | | | | | |
| --- | --- | --- | --- | --- | --- | --- |
| **Family** | **Viral contig** | **Host groups** | **Identification tools** | **^1^Confidence score** | **^2^E-value** | **Usage frequencies closest to Bacteria** |
| *Totiviridae* | LC_coassembly_k141_2783551 | Bacteria | iPHoP-RF | 91.4 | NA | AAUF/DUF |
| *Mitoviridae* | LC_coassembly_k141_2488396 | Bacteria | iPHoP-RF | 97.1 | NA | TUF |
|  | LC_20m_3um_k141_494413 | Bacteria | iPHoP-RF | 90.5 | NA | DUF/TUF |
|  | LC_80m_1.2um_k141_267137 | Bacteria | CRISPR-Cas spacer evidence from IMG | NA | 1.29E-08 | TUF |
| *Narnaviridae* | LC_coassembly_k141_2371561 | Bacteria | iPHoP-RF | 92.4 | NA | DUF/TUF |
| *Benyviridae* | LC_20m_0.2um_k141_269092 | Bacteria | Host gene | 93.6 | 2.25E-83 | AAUF/DUF/TUF |
| *f.0028* | LC_20m_0.2um_k141_260174 | Bacteria | iPHoP-RF | 97.1 | NA | DUF/TUF |
| *f.0076* | LS_coassembly_k141_357820 | Bacteria | CRISPR-Cas spacer evidence from iPHoP | 91.3 | 4.56E-06 | AAUF/DUF/TUF |
| *f.0102* | LC_coassembly_k141_1675587 | Bacteria | iPHoP-RF | 91.1 | NA | DUF/TUF |
|  | LS_coassembly_k141_1274756 | Bacteria | iPHoP-RF | 97.1 | NA | DUF |
| *f.0111* | LC_20m_3um_k141_102044 | Bacteria | Host gene | NA | 8.66E-11 | DUF/TUF |
|  | LC_coassembly_k141_1198831 | Bacteria | CRISPR-Cas spacer evidence from IMG | NA | 5.77E-31 | AAUF/DUF/TUF |
| *f.0176* | LC_50m_1.2um_k141_154296 | Bacteria | iPHoP-RF | 92.4 | NA | DUF/TUF |
| *f.0283* | LC_coassembly_k141_2477916 | Bacteria | iPHoP-RF | 96.1 | NA | TUF |
|  | LC_coassembly_k141_1763940 | Bacteria | iPHoP-RF | 92.1 | NA | DUF/TUF |
|  | LC_20m_10um_k141_92587 | Bacteria | iPHoP-RF | 94.7 | NA | DUF/TUF |
| *f.0286* | LC_20m_3um_k141_11485 | Bacteria | iPHoP-RF | 91.4 | NA | AAUF/DUF |
| *f.0295* | LC_20m_0.2um_k141_261093 | Bacteria | iPHoP-RF | 93.1 | NA | AAUF/TUF |
| *f.0296* | LC_20m_10um_k141_27699 | Bacteria | iPHoP-RF | 90.8 | NA | DUF |
|  | LC_20m_10um_k141_33231 | Bacteria | iPHoP-RF | 95.1 | NA | NA |
| *Marnaviridae* | LS_coassembly_k141_1097885 | Archaea/Bacteria | iPHoP-RF | 94.7/91.4 | NA | AAUF/DUF/TUF |
| 1.Confidence score: The confidence score provided by the iPhop software | | | | | | |
| 2.E-value: The e-value obtained from the CRISPR-Cas spacer or host gene alignment | | | | | | |

| **Table S9. Detailed information of AMGs and genes associated with anti-prokaryotic immune systems.** | | | | | | | | | | | |
| --- | --- | --- | --- | --- | --- | --- | --- | --- | --- | --- | --- |
| **ORF** | **Viral group (family-level)** | **Type** | **Name** | **Description** | **Identification  tools** | **E-value** | **Evidence_accession (cenote3)** | **Seed_ortholog (eggnog)** | **eggNOG_Ogs** | **Max_annot_lvl (eggnog)** | **COG category** |
| LC_0.5m_1.2um_k141_396381_7 | *f.0176* | Resistance | APIS022 | viral anti-prokaryotic immune system (APIS) proteins | Diamond BLASTx | 2.70E-04 | NA | NA | NA | NA | NA |
| LC_20m_0.2um_k141_269092_5 | *Benyviridae* | heme biosynthesis | hemX | putative uroporphyrinogen-III C-methyltransferase | eggNOG-mapper Cenote-Taker | 2.25E-83 3.17E-49 | PRK06975 | 365044.Pnap_3088 | COG2959@1\|root,COG2959@2\|Bacteria,1MY3A@1224\|Proteobacteria,2VMG1@28216\|Betaproteobacteria,4AARN@80864\|Comamonadaceae | 28216\|Betaproteobacteria | H |
| LC_20m_0.2um_k141_269092_6 | *Benyviridae* | heme biosynthesis | hemY | heme biosynthesis-associated TPR protein | eggNOG-mapper Cenote-Taker | 1.93E-65 2.88E-18 | TIGR00540 | 1458427.BAWN01000013_gene869 | COG3071@1\|root,COG3071@2\|Bacteria,1MU7A@1224\|Proteobacteria,2VK6P@28216\|Betaproteobacteria,4A9SX@80864\|Comamonadaceae | 28216\|Betaproteobacteria | H |
| LC_20m_3um_k141_102044_1 | *f.0111* | Unkonw | KUTG | unknown Actinobacteria protein | eggNOG-mapper | 8.66E-11 | NA | 345341.KUTG_04140 | 2DTR4@1\|root,33MCB@2\|Bacteria,2HAJT@201174\|Actinobacteria | 201174\|Actinobacteria | NA |
| LC_20m_3um_k141_357943_4 | *f.0167* | Lipid metabolism | ugpQ | glycerophosphoryl-diester-phosphodiesterase | Cenote-Taker | 2.71E-05 | PFAM:PF15254.5 | NA | NA | NA | NA |
| LC_coassembly_k141_1021355_1 | *f.0176* | Resistance | APIS022 | viral anti-prokaryotic immune system (APIS) proteins | Diamond BLASTx | 2.70E-04 | NA | NA | NA | NA | NA |
| LC_coassembly_k141_2670924_1 | *Dicistroviridae* | translation | rplA | Binds directly to 23S rRNA. The L1 stalk is quite mobile in the ribosome, and is involved in E site tRNA release | eggNOG-mapper Cenote-Taker | 1.13E-18 9.67E-16 | TIGR01169 | 880071.Fleli_2104 | COG0081@1\|root,COG0081@2\|Bacteria,4NEIC@976\|Bacteroidetes,47JJZ@768503\|Cytophagia | 976\|Bacteroidetes | J |
| LC_coassembly_k141_945938_1 | *Nam-Co_family_51* | energy generation | atpA | ATP synthase CF1 alpha subunit | Cenote-Taker | 4.85E-19 | CHL00059 | NA | NA | NA | NA |
| LS_coassembly_k141_232625_2 | *f.0102.base-Curvula* | energy generation | cyoA | Cytochrome C oxidase subunit II, periplasmic domain | eggNOG-mapper/Cenote-Taker | 5.98E-50 1.71E-47 | cd13912 | 70448.Q0P3F9 | COG1622@1\|root,KOG4767@2759\|Eukaryota,37T4F@33090\|Viridiplantae,34MHI@3041\|Chlorophyta | 3041\|Chlorophyta | C |
